# Prediction of metabolic dynamics through deep learning and high-throughput multiomics data

**DOI:** 10.1101/2025.05.27.656438

**Authors:** Jose Manuel Marti, Patrick C. Kinnunen, Mark Held, Azadeh Alikhani, Andrew Conley, Yan Chen, Tijana Radivojevic, Jennifer W. Gin, Apostolos Zournas, Rebecca M. Lennen, Hooman Hefzi, Mustafa Baniodeh, Peter Harrington, Mark Forrer, David Melis, Jeffrey Dietrich, William Holtz, Nick Ohler, Christopher J. Petzold, Hector Garcia Martin

## Abstract

Synthetic biology’s remarkable potential to tackle important societal problems is held back by our inability to predictably engineer biological systems. Here, we collected one of the largest public multiomics synthetic biology datasets generated to date, and used it to train a novel deep learning algorithm able to predict product and metabolic dynamics with great accuracy, starting to approach the predictive capabilities found in physics and chemistry. We were able to predict production time series with 90-99% accuracy, and final production with 96% accuracy. Further, we were able to produce good predictions for a majority of extracellular metabolites, and twenty different intracellular metabolites. These predictions were provided for a target of industrial relevance: a non-model yeast (*Pichia kudriavzevii*) engineered to produce large amounts of malonic acid, a desirable biomanufacturing target. This approach is generally applicable to any host, pathway, and product because all required knowledge is inferred from experimental data.

## Introduction

Synthetic biology has the alluring potential to provide enhanced-performance products, diminish supply chain disruptions, produce environmental benefits, address national security needs, and create new jobs and industries ^1–5^. Synthetic biology and biomanufacturing have contributed to the bioeconomy ^6,7^ by producing, e.g.: antimalarial and anticancer drugs ^8,9^, effective treatments for leukemia ^1,10^, biological concrete ^11^, biodegradable packing materials ^12^, synthetic silk for clothing ^13^, meatless burgers that taste like meat ^14^, greenhouse gas capture solutions ^15,16^, and aviation fuels with improved energy density ^17^. In the future, synthetic biology is also expected to help enable a circular bioeconomy ^18^ that diverts our society from an overreliance on petrochemicals, a goal stated by the European Union ^19^, China ^5^, Japan ^20^, U.K. ^21^, and previous US administrations ^22^. Indeed, the demand for products that use biological processes and genetically modified microorganisms in place of traditional production methods is increasing, and is estimated to have a market of ∼$200B by 2040 if production costs can be significantly lowered ^23^.

However, the steep challenge of systematically predicting the outcomes of cell modifications as accurately as for physical or chemical systems, makes synthetic biology a costly and long process ^24–26^. For example, it took Amyris an estimated 130 to 575 person-years to make farnesene, and Dupont and Genencor about 15 years and 575 person-years to generate an economical process for 1,3-propanediol ^27,28^. This inability to predict biological systems’ behavior stands in stark contrast to the predictive capabilities available in physics, where computational fluid dynamics can produce accurate simulations of water ^29–31^, and professional Formula 1 drivers are routinely trained on simulators leveraging sophisticated physics engines ^32^; or in chemistry, where Density Functional Theory (DFT ^33^) calculations can guide the design of Li-ion batteries with enhanced power delivery ^34^.

The increasing abundance of multiomics data ^35–41^ has not yet truly revolutionized our ability to engineer biological systems, because it is difficult to digest these data to effectively produce actionable recommendations. The tremendous complexity of biological systems, combined with the large amount of data and data types, impedes efficient use of the traditional intuition-driven hypothesis generation and testing. This hurdle is further exacerbated when working with the non-model organisms that are often required for large-scale production because of their robustness and metabolic versatility. Hence, and despite recent advances, the “Learn” phase of the synthetic biology Design-Build-Test-Learn (DBTL) cycle is widely acknowledged as the weakest link in the process ^25,42^.

Machine learning (ML) can provide the predictive power synthetic biology needs to enable efficient biomanufacturing. While mechanistic mathematical models have made great strides in predicting phenotypes and providing guidance for synthetic biology, their limited predictive power inhibits their widespread adoption. Genome-scale models or kinetic models, for example, have been successfully used to improve titers, rates and yields (T.R.Y.) ^2,43^, and whole-cell models have been used to successfully predict the outcomes of novel experiments ^44,45^. Even so, improving these mechanistic models by leveraging mispredicted experimental data is still a manual process that can take months for each DBTL cycle. Among mechanistic models, kinetic models are particularly favored because they establish mechanistic relations between metabolic reaction rates, enzyme levels, and metabolite concentrations ^43^, and capture cellular physiology beyond the metabolically steady-state assumption of traditional genome-scale metabolic models based on mass balance principles ^46^. However, kinetic models take a long time to develop and calibrate ^43,46^, and they can show limited predictive power because essential mechanisms are only sparsely known ^47^. ML approaches, on the other hand, are designed to leverage experimental data to systematically improve predictions. Their fruitful application in biology has been recently demonstrated by Alphafold’s predictions of protein structure from sequence ^48^, as well as recent achievements in pathway design ^49^, enzyme annotation ^50^, and protein engineering ^51^, among others ^52^.

We have previously demonstrated an ML-based approach, “kinetic learning” ^47^, that improved on kinetic modeling ^43,46^, but required a significant amount of data to be effective. Kinetic learning “learns” the dynamics directly from training data time series, and was able to outperform in predictive power a classical Michaelis–Menten kinetic model ^47^. Furthermore, these kinetic learning models are easier to construct, and more systematic and generally applicable than kinetic models because all required knowledge (regulation, host effects… etc) is inferred from experimental data, instead of arduously gathered and introduced by domain experts. However, kinetic learning was tested with a small dataset: only two time-series (strains) for training and one for testing were available. As a consequence, predictions were often qualitative and showed very limited accuracy.

Here, we create a completely new version of kinetic learning by integrating deep neural networks, so it can efficiently leverage high-throughput proteomics and metabolomics, and use it to very accurately predict metabolic dynamics in a non-model yeast producing industrially relevant levels of renewable malonic acid (Fig. 1). Malonic acid is a desirable target within the circular bioeconomy because it is considered one of the top 30 value-added chemicals from biomass ^53^, and is a versatile “building-block” used for applications ranging from coatings ^54^ to fragrances ^55^ and pharmaceuticals ^56,57^ (Supplementary Note 1). For this purpose, we generated a dataset consisting of about 80,000 data points which, to our knowledge, is one of the largest public microbial bulk (non-single cell) multiomics datasets generated to this date (ca. 2025) for synthetic biology. We used these data to train the deep neural networks encoding a novel kinetic *deep* learning (KinDL) approach, and were able to predict production time series with 90-99% accuracy and final production with 96% accuracy. Further, we were able to produce good predictions for a majority of extracellular metabolites, and twenty different intracellular metabolites, exceeding results by previous kinetic models. The accuracy of these predictions surpasses that of previously published kinetic models, genome-scale models, whole cell models, or any other attempt to predict cell behavior that we are aware of. Importantly, the model producing these results was crafted automatically from data, without the need to be manually developed by experts in metabolism. We used the predictive power of KinDL to recommend genetic edits that increased malonic acid production in a high-producing non-model yeast, although reasons beyond the metabolic dynamics modeling (reproducibility, protein modulation) limited the final performance improvement. These results show that ML approaches, trained with sufficient multiomics data, can provide predictive power for synthetic biology starting to approach what is found in physics and chemistry, helping unlock its full potential to nourish the bioeconomy with novel renewable products.

**Figure 1:**
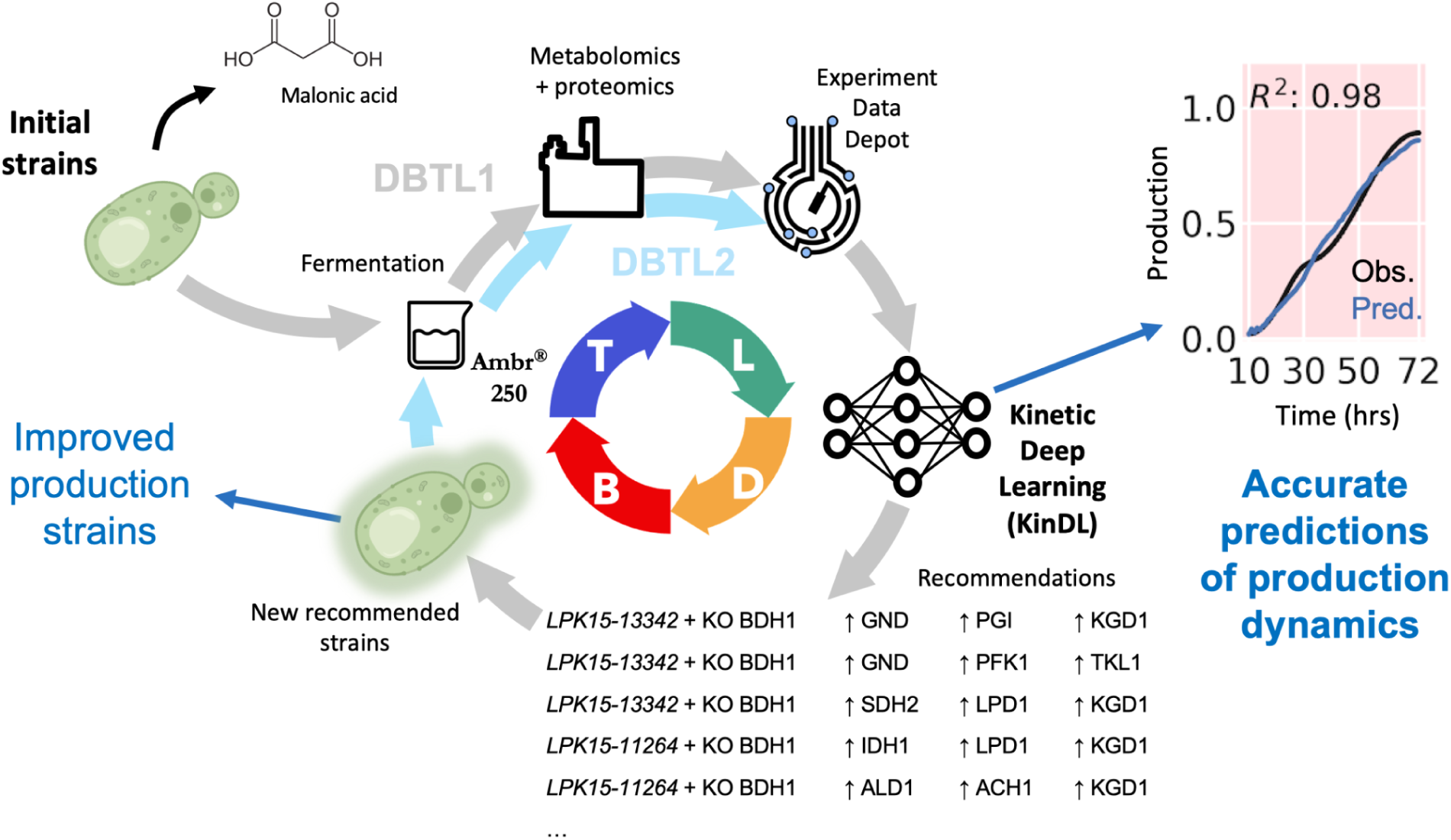
Deep learning leveraging large amounts of multiomics data can produce very accurate predictions of metabolism dynamics. In Design-Build-Test-Learn (DBTL) cycle 1 (in gray), twenty three initial *P. kudriavzevii* strains engineered to produce malonic acid (Supplementary Table 1) were cultured in the Sartorius Ambr250 automated fermentation platform, known to produce results that scale to higher volumes. Samples for intracellular metabolomics and proteomics, as well as extracellular metabolomics were obtained for eight time points, and the ensuing data stored in the Experiment Data Depot (EDD) database. These data were used to train a novel Kinetic Deep Learning (KinDL) algorithm to accurately predict the time series of Total Malonic Acid Formed (TMAF) and concentrations of extracellular and intracellular metabolites, from proteomics time series. Based on these predictive models, recommendations for genetic modifications (down regulations, up regulations or knockouts) that were predicted to improve malonic acid production were generated. In DBTL cycle 2 (blue) these recommendations were used to build twenty two new strains (Supplementary Table 2) that improved production over the control strain.

## Results

### Generating one of the largest multi-omics microbial synthetic biology datasets to date

We generated a dataset of about 80,000 data points for the non-model yeast *P. kudriavzevii* (Fig. 2) in order to train deep artificial neural networks (ANNs), notable for their need for large datasets to be truly effective ^58^. We produced these data from *P. kudriavzevii* (Pk) YB-431 strains engineered by Lygos to produce large amounts of malonic acid (in the range of 50-150 g/L) at demonstration scale ^59^ (Supplementary Note 1). These figures place this strain in the high production regime where further improvement efforts typically face diminishing returns. *P. kudriavzevii* is a host of industrial relevance because of its multi-stress tolerant physiology (used for, e.g., production of succinic and lactic acids by Bioamber and Cargill ^60–62)^, and is an unconventional non-model Crabtree-negative yeast that exhibits very different carbon flux profiles from model yeasts. Hence, it presents significant challenges to the traditional process of intuition-driven hypothesis generation and testing for non *P. kudriavzevii* specialists.

**Figure 2:**
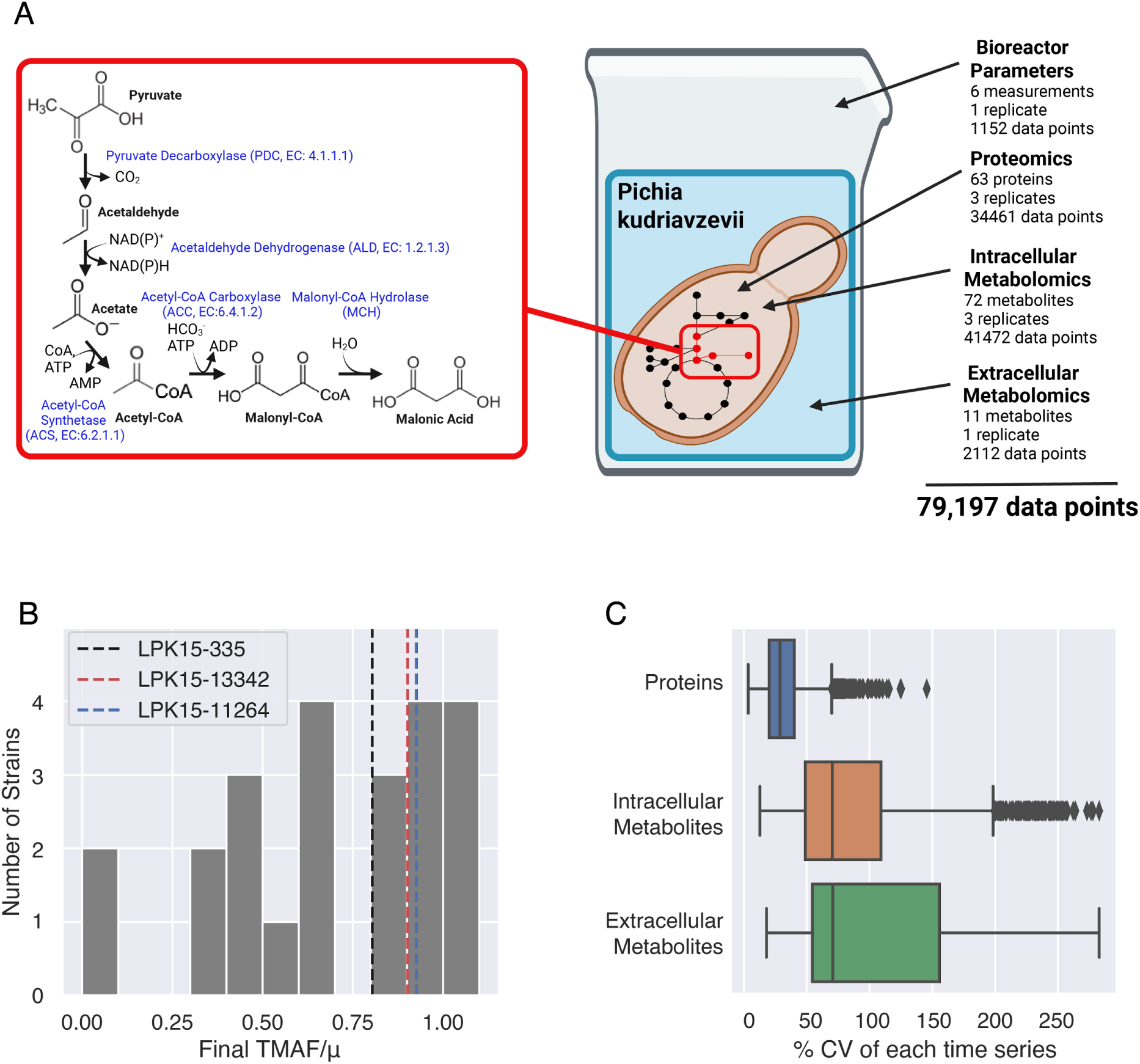
One of the largest multiomics datasets to date was collected to train the machine learning algorithms used to predict metabolic dynamics. **A)** The malonic acid production pathway involves five reactions (left) with a dependence on ATP and NAD(P)^+^ as cofactors, and was engineered in the non-model yeast *P. kudriavzevii*. We collected times series of eight time points for each the twenty three initial strains for proteomics data (63 proteins, Supplementary Table 3), intracellular metabolites (72 metabolites, Supplementary Table 4), extracellular metabolites (11 metabolites including carbon source, product and main excretion products, Supplementary Table 5), and bioreactor fermentation parameters (6 measurements including TMAF, pH, dissolved oxygen, etc, Supplementary Table 6), totaling around 80,000 data points, stored in EDD ^71^. Technical (Supplementary Figs 1, 2) and biological (Supplementary Figs 3, 4) replicates show good repeatability within a single fermentation run. **B)** The initial set of strains displayed a variety of production levels of Total Malonic Acid Formed (TMAF), so as to provide an informative training set. Production data was normalized by an undisclosed factor **μ** to preserve Lygos competitive advantage for an embargo period (see Data Availability). Strain LPK15-335 was used by Lygos as a baseline for this project (progenitor to a scaled-up strain), LPK15-11264 was the top production strain at the beginning of the project, and LPK15-13342 is the parental strain for the recommendations discussed below. **C)** Proteins tend to be more stable in time than metabolites, as shown by the distribution of coefficients of variation (cv, standard deviation over mean, specific examples available in Extended Data Fig. 1). This stability is a desirable characteristic as targets for implementation of recommendations. Boxes show the 25th, 50th, and 75th percentile of the data.

The data collected is one of the largest public bulk multi-omics synthetic biology datasets, and comprised proteomics and internal and external metabolomics for 23 different strains (Supplementary Table 1) and eight time points spanning 72 hours (Fig. 2A). The proteins and internal metabolites measured included those involved in the malonic acid pathway, as well as in central carbon metabolism, which supplies precursors to this pathway (Supplementary Tables 3 and 4). External metabolites measured included organic acids, previously observed *P. kudriavzevii* overflow metabolites (Supplementary Table 5), and the Total Malonic Acid Formed (TMAF). TMAF accounts for all malonic acid formed, including the portion removed from the system during sampling (see Methods). TMAF would later be the target to optimize by ML recommendations because it is a single figure that encompasses the most important industrial performance indicators: titer, rates and yield (T.R.Y.). This dataset is, to our knowledge, the largest public bulk (i.e., not single-cell) multi-omics dataset (63 proteins and 72 metabolites) to characterize a large amount (twenty three) of engineered strains of a non-model organism over many timepoints (eight). While several studies have used multi-omics to assess the effects of perturbations or interventions ^63–67^ (e.g., increased protein level after modifying a promoter) or to identify pathway bottlenecks ^36,68,69^, these datasets typically either capture only a few timepoints, compare a few strains, or study model organisms such as *E. coli* or *S. cerevisiae*.

A coarse-grained view of the data confirms that a wide variety of TMAF production values were explored (Fig. 2B) and that protein levels tend to be more stable as a function of time than metabolites (Fig. 2C). The twenty three strains used in the first DBTL cycle were extremely diverse so as to facilitate learning by the ML algorithm: they comprise everything from near wildtype strains that exhibit no malonic acid production (for which most of the glucose consumed is respired) to the top performing strain (at the start of this project), where the majority of the carbon was locked in the malonic acid pathway (Fig. 2B). Inspection of the proteomics and metabolomics data shows that protein concentrations are, in general, much more stable as a function of time (median coefficient of variation of 28.8%) than intracellular (median cv=70.5%) and extracellular metabolites (median cv=70.8%, Fig. 2C). This stability is a desirable characteristic as input features for the ML algorithm and targets to be acted upon (e.g., a doubling of protein expression is presumably easier to induce and check if it is stable over time).

Productively gathering, doing quality checks and analyzing such a large dataset required infrastructure and tools that are not often found in synthetic biology academic laboratories (see Methods). The tools provided by the DOE Agile BioFoundry ^70^ were critical to store experimental (Experiment Data Depot, EDD ^71,72^) and strain (Inventory of Composable Elements, ICE^73^) information. It would have been impossible to handle this amount of data and metadata (Fig. 2A) through, e.g., the exchange of MS Excel files which still represent the standard modus operandi in many synthetic biology projects ^74^, and are known to result in significant scientific errors ^75^ ^76^.

### Kinetic deep learning can predict pathway and product dynamics with great accuracy

Leveraging the large amounts of data collected, the deep neural networks used by KinDL (Fig. 3) were able to accurately predict product and metabolic dynamics for external metabolites and a large number of internal metabolites (Figs 4, 5, 6, Extended Data Fig. 2). To obtain these predictions, the algorithm was completely revamped from the original kinetic learning to a more effective approach that eliminated the need for integration (and the associated errors), and leveraged an ensemble of deep neural networks instead of the previously used traditional machine learning algorithms (Fig. 3). Moreover, despite being very large for synthetic biology standards, the dataset still needed to be augmented in order for deep learning to be effective, which we did through interpolation and using all replicates as instances (Fig. 3).

**Figure 3:**
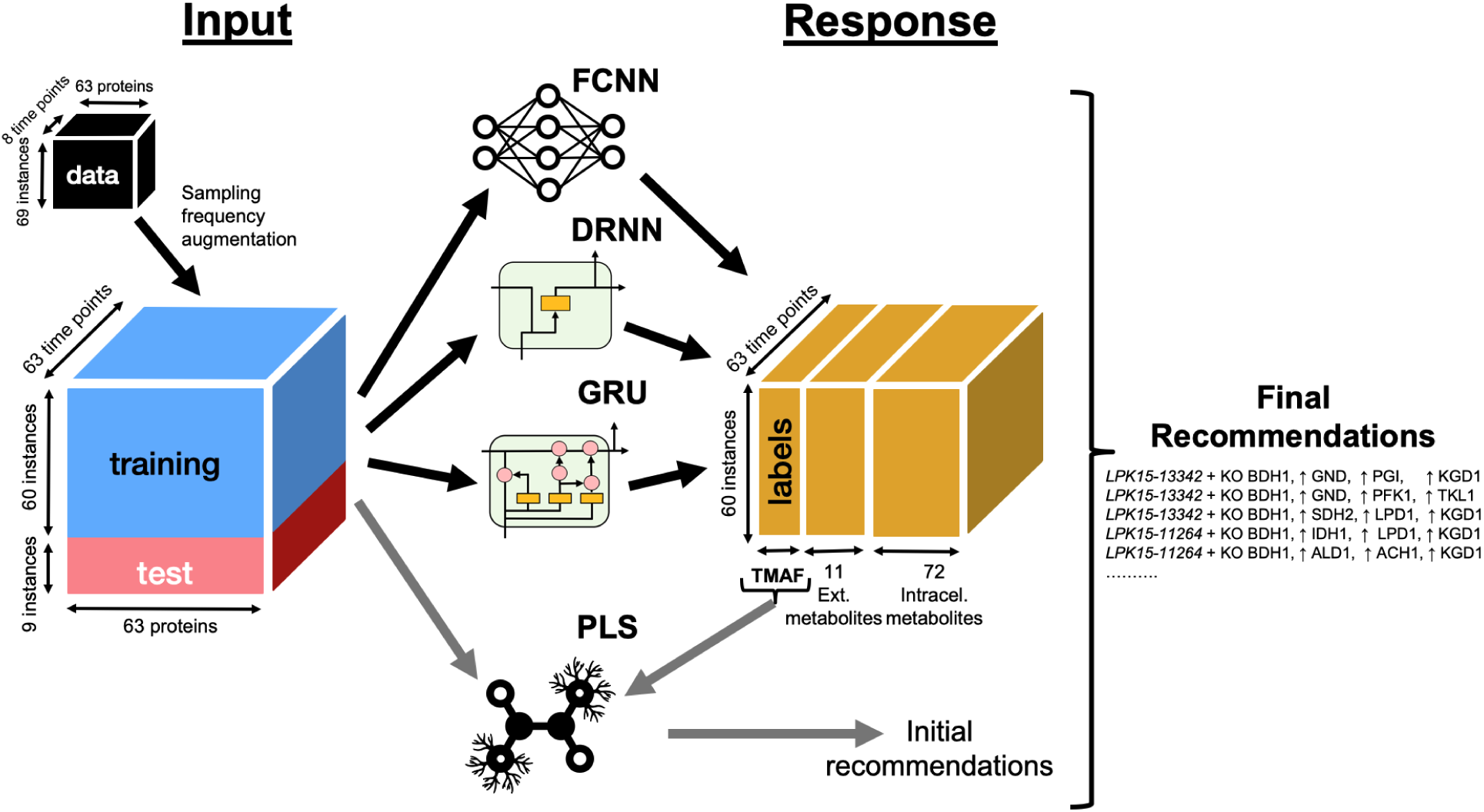
Deep learning enabled the accurate prediction of time series of production (TMAF) and metabolite concentrations from time series of protein concentrations. The input was the proteomics time series, and the responses to be predicted were the times series of: TMAF, extracellular metabolite concentrations, and intracellular metabolite concentrations. Whereas the original kinetic learning approach involved using ML to learn the rate of change of metabolites from proteomics and metabolites ^78^, kinetic deep learning directly maps proteins time series to final product and metabolite time series using a combination of four different methods: three deep neural networks (a fully connected neural network, FCNN, a deep recurrent neural network, DRNN, as well as a gated recurrent neural network, GRU, Supplementary Tables 7-10), and a partial least squares (PLS) model. We aggregated the predictions of the three neural networks to obtain a single final predicted time series by taking the weighted average of each individual model output, where weights were proportional to the inverse of the model accuracy. Our intention was for the deep hidden layers to capture core metabolism and for subsequent layers to use it to predict dynamics for each metabolite (Extended Data Fig. 3). Due to the very large number of possible recommendations, the PLS model provided a simple and fast approach to suggest recommendations, while the weighted average of deep neural networks provide much more accurate response predictions to check and correct these PLS recommendations. The increased predictive power of deep learning neural networks with respect to traditional methods comes at the expense of requiring large amounts of training data. We augmented our data (63 proteins and 72 metabolites measured at 8 time points in triplicate) as much as possible by 1) interpolating the eight measured time points to a total of 63 time points, and 2) by using all triplicates as training instances instead of just using averages. This approach allowed us to expand our training dataset from the 74,520 data points in the proteomics and metabolomics data (Fig. 2) to 586,845 data points. Data quality was checked and enforced by finding gaps in the time series, eliminating negative values, detecting and eliminating outliers, and uncovering premature time series ends. Instances refer here to combinations of strains and replicates (i.e., lines in the rest of this paper).

**Figure 4:**
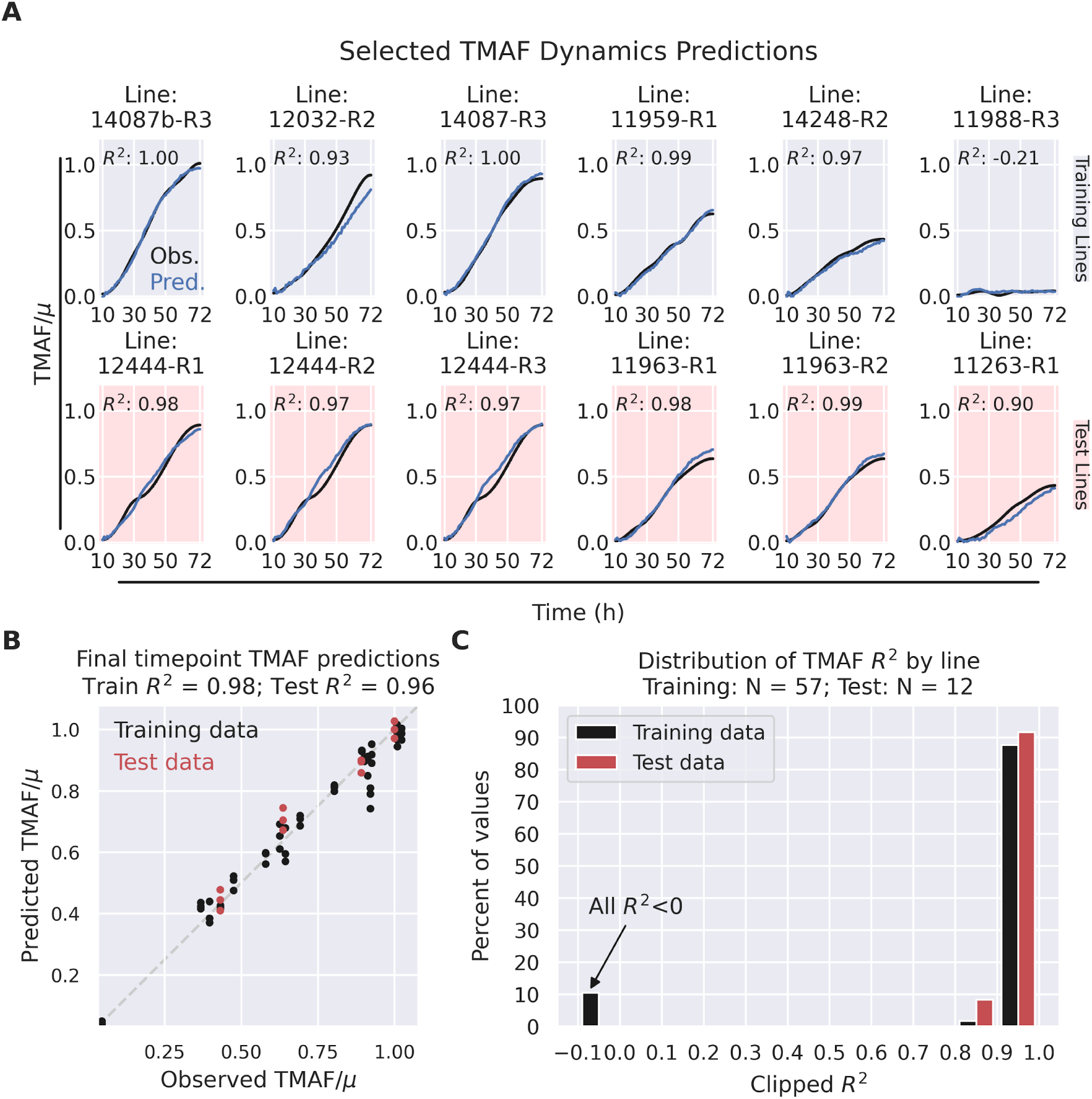
Kinetic Deep Learning can very accurately predict total product formed (TMAF). Time series of production were predicted with 90-99% accuracy, and final time points with 96% accuracy. **A)** Predictions and measured values of TMAF time series for a selection of representative lines (i.e., a combination of a strain and replicate^71^) showed excellent agreement (R^2^ ≥ 0.93 for training, R^2^ ≥ 0.90 for test). The top row (gray) shows training lines (used to train the algorithm), and bottom row (red) shows test lines (data that the algorithm has not seen before). Predictions for training data were excellent (R^2^ ≥ 0.93, explaining more than 93% of the data variance) for these lines, except for the few cases showing very low and relatively constant values of TMAF (e.g., last line in row, 11988-R3). Predictions for test data were also very accurate for these lines, explaining more than 90% of the data variance (R^2^ ≥ 0.90). Training and test lines were selected so as to showcase the full range of final TMAF values from high to low (Fig. 2B). Similar conclusions can be drawn for all twelve test lines (0.90 < R^2^ ≤ 0.99, 4 strains in triplicate totalling 96 data points, Extended Data Fig. 2). **B)** Predictions for final time points of TMAF were also extremely accurate: the model could explain 98% of the training data variance (R^2^ =0.98), and 96% of the test data variance (R^2^ =0.96). **C)** The distribution of R^2^ values for all the lines further confirmed that predictions are very accurate: R^2^ ≥ 0.90 for all test lines (Extended Data Fig. 2), showing that the model can explain more than 90% of the data variance. From these, ∼91% of test lines show R^2^ > 0.90, and 9% show 0.80 < R^2^ ≤ 0.90 (R^2^=0.9, actually). For the sake of visualization, all values of R^2^ below 0.1 are shown as 0.1. All R^2^ values below 0 are for strains where TMAF had very low variation (e.g., 11988-R3 in A), above).

Predictions for time series of total product formed (TMAF) were very precise (Fig. 4), predicting test data with 90-99% accuracy (coefficient of determination 0.99 ≥ R^2^ ≥ 0.90, see Methods). Predictions for TMAF training lines time series were excellent, explaining 98% of the data variance or more (R^2^ ≥ 0.98) for a majority (88%) of cases (Extended Data Fig. 2). The only exceptions were the few cases showing very low and relatively constant values (e.g., line 11988-R3 in Fig. 4A). Here, “lines” represent a combination of a strain and replicate ^71^ (e.g., 11988-R3 is the third replicate of strain 11988). It is the test line predictions, however, that truly show the generalizability of the algorithm’s predictive power, since it is data the algorithm has not “seen” before. For these test lines, the TMAF time series predictions were excellent, explaining 90% or more of the data (R^2^ ≥ 0.90) for all cases (Extended Data Fig. 2), boding well for the future recommendations. We also found that using fewer proteins as input only minimally degraded predictions, a useful piece of information given the high cost of multi-omics data (Extended Data Fig. 7). We identified the ten most important input features (proteins) through SHapley Additive exPlanations (SHAP) analysis ^77^ and found them to be proteins related to redox balance (mostly dehydrogensases, Extended Data Fig. 7A). We tested the predictive power of KinDL when using only these ten proteins as inputs, and found that TMAF final timepoint predictions deteriorated minimally for the final time points (test data R^2^= 0.86 vs 0.96, mean absolute error 0.04 vs 0.03) when compared with predictions using the full set of 63 proteins (Fig. 4B and Extended Data Fig. 7B). The TMAF time series predictions seemed equally excellent for ten and sixty three proteins (similar distribution of R^2^ for both cases, Fig. 4C and Extended Data Fig. 7C). However, the identity of the ten most informative proteins can only be found *a posteriori* in this manner, and hence are only practically useful for follow-up DBTL cycles (not initial ones).

Final timepoint TMAF predictions were also extremely accurate, explaining 96% of the data for test lines (R^2^ = 0.96) and 98% for training lines (R^2^ = 0.98, Fig. 4B). Interestingly, we found that the use of time series produced more accurate predictions than using only final time points (Extended Data Fig. 6), indicating a way in which phenotype datasets in synthetic biology can be augmented without the need to build new strains. A comparison with the Automated Recommendation Tool (ART), a ML tool successfully used in the past to predict single response points ^49,78,79^, shows that single point predictions are much less accurate than those we obtained by leveraging the time series (test data R^2^ = 0.60 from ART vs 0.96 from KinDL, Fig. 4B and Extended Data Fig. 6). Given the high-throughput proteomics pipelines that are starting to become available ^39^, collecting time series of data can be an effective way to enlarge the training dataset and improve the accuracy of ML predictions, since strain construction is still often a bottleneck in many synthetic biology projects.

Predictions for the extracellular metabolic time series were less accurate than for TMAF (Fig. 5), but still showed good predictions (test R^2^ ≥ 0.6) for a majority of metabolites. Indeed, most (8/11=73%) external metabolites (glucose, arabitol, malonate, uracil, trehalose, glycerine, citrate, and pyruvate) showed good predictions, explaining more than 60% of the test data variance (test R^2^ ≥ 0.6, Fig. 5C). Moreover, about half of the metabolites (45%) were very well predicted, explaining more than 80% of the test data variance (test R^2^ ≥ 0.8, Fig. 5C). In particular, the excellent predictions for extracellular malonate (R^2^ = 0.93, Fig. 5A) seemed to explain the high quality of TMAF predictions discussed above.

**Figure 5:**
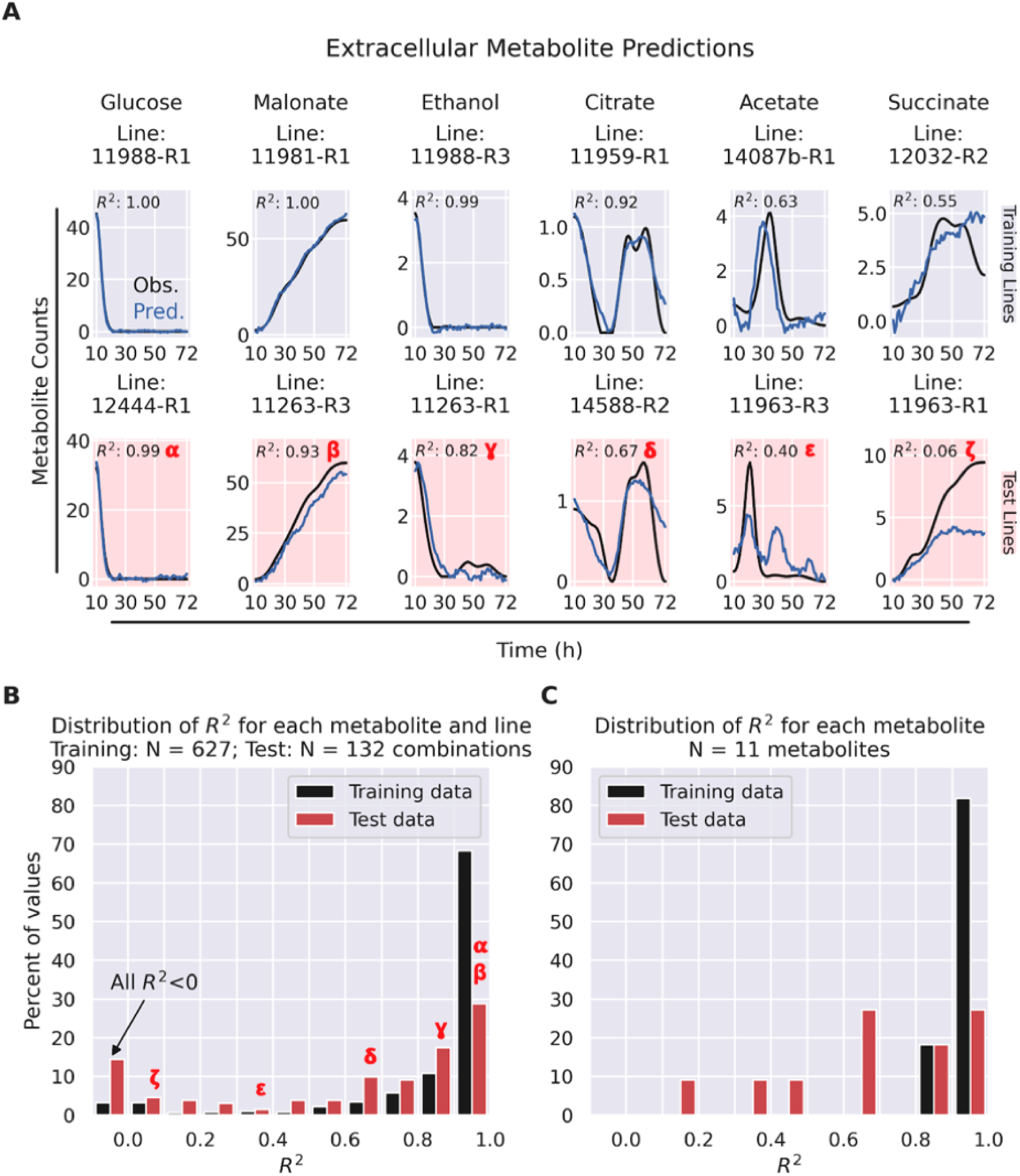
Kinetic Deep Learning provides good predictions of time series of extracellular metabolite concentrations. Extracellular metabolite dynamics were well predicted (R^2^ > 0.60 for 73% of metabolites), but less so than final product dynamics (Fig. 4). **A)** Representative extracellular metabolite dynamics for a variety of prediction performances (test R^2^ from 0.99 to 0.05) are shown in the upper panel for glucose, malonate, ethanol, citrate, acetate, and succinate. The relative abundance of each of these R^2^ values is shown in the histogram in the next panel through greek letters (α,β,γ,δ,ε,ζ). Predictions for all metabolites and lines can be found in Supplementary Material. **B)** The distribution of R^2^ values for each metabolite and line (i.e. no line aggregation) shows that predictions are quite accurate: in training, most combinations of metabolites and lines (88%) displayed R^2^ > 0.60, and in test, 65% of combinations showed R^2^ > 0.60. The leftmost bar represents all R^2^ values below 0. **C)** The distribution of R^2^ values for each metabolite aggregated across all lines similarly show good predictions across all metabolites: a majority of test data metabolites (73%) show R^2^ > 0.60 (100% for training data).

Predictions for intracellular metabolite time series showed a wider scope of results, ranging from excellent to poor, and a larger difference between training and test sets, but still showed good performance for twenty different metabolites (Fig. 6). Training lines showed a large majority of metabolites (76%) being well predicted (R^2^ ≥ 0.6). Test lines displayed only 28% of metabolites being well predicted (R^2^ ≥ 0.6), 24% showing passable predictions (0.6 > R^2^ ≥ 0.4), and half (48%) being poorly predicted (R^2^ < 0.4, Fig. 6C). Well predicted intracellular metabolites in the test set included glucose and malonate, directly involved in the engineered pathway, as well as NADP^+^ and coenzyme A, cofactors with global metabolic roles (Supplementary Table 11). Although extracellular metabolites are better predicted than intracellular ones, suggesting that higher concentration metabolites are perhaps better predicted, we do not see a dependence of errors on concentration (Extended Data Fig. 4). The prediction failures do not seem to have a single cause (Extended Data Fig. 5). To improve accuracy, we expect larger training sets than the nineteen strains used here are needed, as our previous tests with synthetic data have shown ^47^. In any case, the twenty well predicted metabolites represent more well predicted metabolites than previous kinetic modeling work ^80–85^.

**Figure 6:**
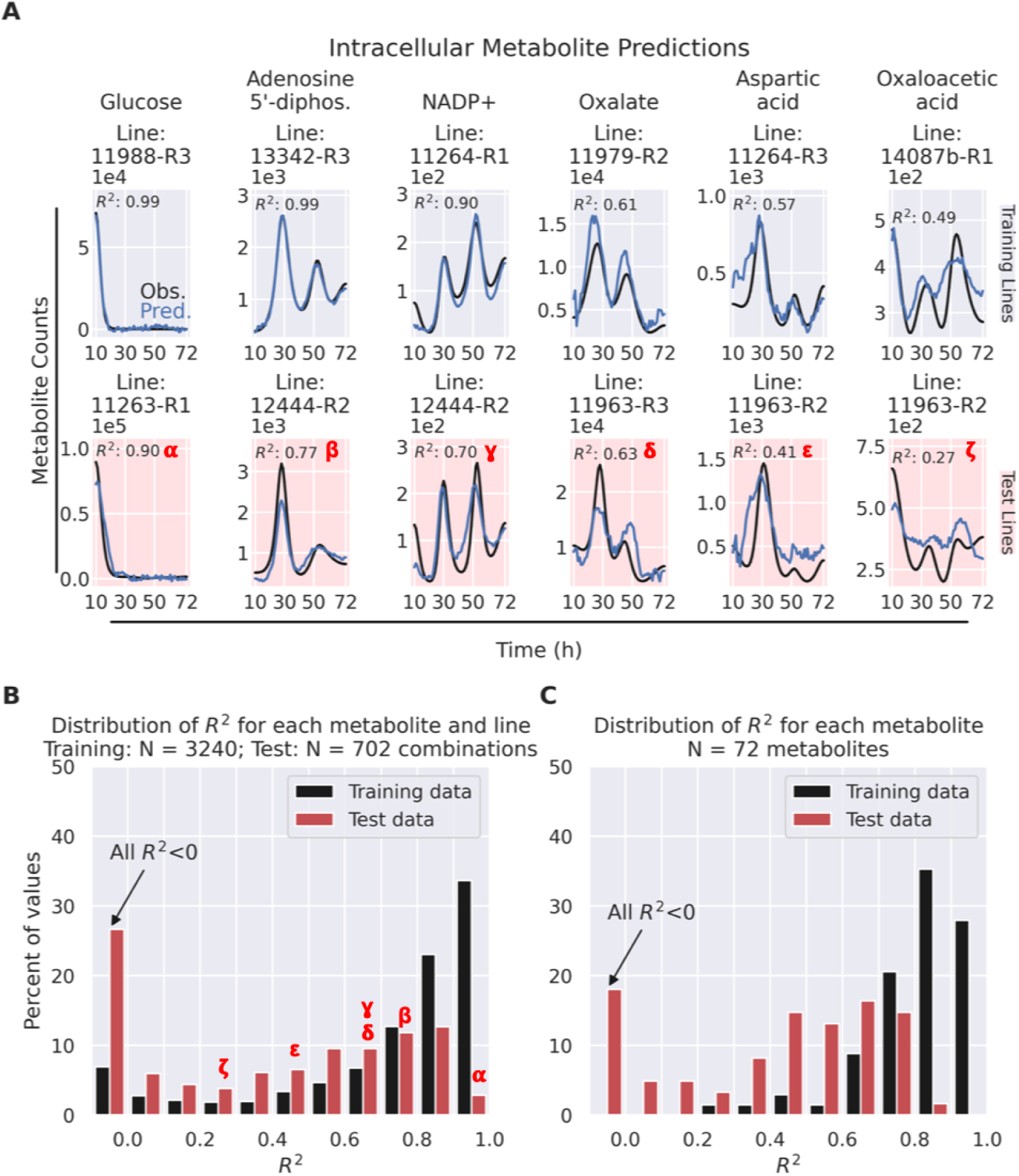
Predictions for intracellular metabolite time series showed a wide scope of results, ranging from excellent to poor. Although internal metabolite concentration time series were predicted in general with less accuracy than external metabolites (Fig. 5) or final product (Fig. 4), twenty metabolites were well predicted (R^2^ > 0.60 in test dataset). **A)** Here we show representative metabolic dynamics for several prediction performances (test R^2^ from 0.90 to 0.27), pertaining to glucose, ADP, NADP+, oxalate, aspartic acid, and oxaloacetic acid. The relative abundance of each of these R^2^ values is shown in the histogram in the next panel through greek letters (α,β,γ,δ,ε,ζ). Predictions for all metabolites and lines can be found in Supplementary Material. **B)** The distribution of R^2^ values for each combination of metabolite and line (i.e., no aggregation of different lines) shows that a sizable fraction of combinations (76% for training, 37% for test) are well predicted (R^2^ > 0.60). The leftmost bar represents all R^2^ values below 0. **C)** The distribution of R^2^ values for each metabolite aggregated across all lines show that a similar fraction of metabolites (88% for training, 28% for test) are well predicted (R^2^ > 0.60). This 28% of well predicted metabolites are twenty metabolites that represent more well predicted metabolites than any previous kinetic modeling work. For all panels, only strain-metabolite combinations below a maximum variability threshold and above a minimum detection limit are shown (Supplementary Fig. 2).

These prediction accuracies for the time series of the final product and extracellular metabolites surpass the latest and best kinetic modeling results ^80–83,86^, even for more amenable cell-free cases ^84,85^. These results do not predict product time series or final time points with the same accuracy (R^2^≥ 0.9 for time series, R^2^ = 0.96 for final time points), and do not provide good predictions for such a large number of metabolites (R^2^≥ 0.6 for eight extracellular metabolites and twenty intracellular metabolites). The comparison between observations and predictions is often not quantitatively assessed, and when performed it is often only through correlation coefficients ^80,81^, a much less stringent metric than the coefficient of determination ^87^ R^2^ we have used here (see Methods). Interestingly, the nineteen strains used in training to produce our results represent much fewer than originally estimated (∼100) to obtain this accuracy ^47^.

### Machine learning recommendations improve production over control strain

We exploited KinDL’s ability to accurately predict TMAF to generate recommendations that resulted in large production increases over the control strain, but limited increases over DBTL1 (Extended Data Fig. 8). To generate these recommendations, we tailored the use of the predictive model to the genetic interventions that could, in principle, be easily created in the Lygos strain building process: one gene knockout (KO) and up to three genes to either down or upregulate. Within these constraints, thousands of recommendations were produced, which were eventually reduced to a final set of 22 strains to be built (Supplementary Table 2), based on the feasibility of construction and the predicted TMAF improvement.

The recommended strains resulted in improvements of ∼20-40% and ∼330-400% with respect to the control strain in two ensuing DBTL cycles (DBTL2.1 and DBTL2.2), but the increase with respect to DBTL1 remained ∼11% because of the variability of the control strain performance (Extended Data Fig. 8). While new strains based on strain LPK15-13342 resulted in TMAF increases of ∼20-40% in an initial version of DBTL2 (DBTL2.1), this parental strain (used as control) only produced two thirds of the TMAF produced in DBTL1. This performance difference resulted in a minor TMAF improvement (8.7%) with respect to DBTL1 (Extended Data Fig. 8). This reproducibility issue prompted a second version of DBTL2 (DBTL2.2), with the expectation of a better control strain performance. Nonetheless, while the newly built strains were able to increase TMAF by ∼330-400% with respect to the control strain (Extended Data Fig. 9), this strain never reached the DBTL1 performance, resulting again in only minor improvements (11.2%) over DBTL1. This distinct variability in production results may be due to a genomic instability, or other effects due to the long time elapsed between DBTL1 and DBTL2.1 and DBTL2.2 (see Supplementary Note 2). This variability highlights the difficulties in ensuring reproducibility for the collection of the large datasets needed for ML, adding evidence to the increasing worry about science reproducibility ^88,89^. Cloud labs ^90,91^ and self-driving labs ^92–94^ can perhaps help ameliorate these problems by heavily leveraging automation in ways that significantly increase DBTL speed and improve reproducibility.

A further challenge possibly preventing the recommended strains from reaching the predicted performance was the inability to match target protein expression levels in engineered strains (Extended Data Fig. 10). While the BDH1 knockout was achieved on virtually all strains, and several of the desired upregulations were attained, no combinations of upregulations reached the recommended levels of expression (challenging our assumption that doubling gene copies would double the corresponding protein expression). This challenge is common in the metabolic engineering of non-model organisms, for which extensive gene regulatory tool sets are lacking, and highlights the importance of developing effective methods to regulate protein expression accurately, as previously discussed ^52,95–97^.

Despite these hurdles, involving issues beyond the modeling of metabolic dynamics, the recommendations were able to improve production in an already high-producing non-model yeast, a non-trivial task. At high production levels, further improvement efforts typically face diminishing returns even in the hands of seasoned metabolic engineers because most of the biological hypotheses guiding the traditional metabolic engineering process have often been exhausted, and they rely on deep metabolic knowledge that is scarce for non-model organisms (behaving very differently from the widely studied model organisms). Moreover, these recommendations were produced by the algorithm almost instantaneously compared with the time it would require a metabolism expert to generate equivalent recommendations.

## Discussion

We have collected one of the largest public microbial multiomics synthetic biology datasets generated to date (Figs. 1-2), and used it to train a completely revamped version of kinetic learning (KinDL), that leverages deep learning for significantly improved performance (Fig. 3). The KinDL algorithm exceeded the accuracy of previous mechanistic kinetic models ^43,80,81,84,86^ in predicting time series, arguably the most challenging type of prediction. KinDL was able to predict time series of production with 90-99% accuracy, and final time points with 96% accuracy (Fig. 4). Further, KinDL produced good predictions (R^2^≥ 0.6) for a majority of extracellular metabolites (7/11), and twenty different intracellular metabolites (Fig. 5 and 6). These results were produced for a strain and conditions which are very relevant to industrial biomanufacturing: a non-model organism (*P. kudriavzevii*) that is often used for large-scale biomanufacturing, synthesis of a desirable bioproduct (malonic acid) at the large quantities (50-150g/L) that are necessary for commercial relevance and for which improvement efforts typically face enhanced difficulty, and a culturing environment (Sartorius Ambr250) that is known to produce results extrapolatable to large scales. The recommendations provided by this model were able to improve production in this high-producing non-model yeast, a non-trivial task even for a seasoned metabolic engineer.

This ML approach is generally applicable to any product, pathway or host, given enough data, and holds the promise of medical applications. Since KinDL infers the predictive model directly from the data, it avoids arduously creating a model from literature, a significant bottleneck in the field ^43^ ^46^. This approach has been previously shown to work for *E. coli* ^47^, and here for *P. kudriavzevii,* a non-model organism. The generality of the approach suggests that accurate predictions for time series of metabolic dynamics in mammalian cells are achievable if enough multiomics data is available. These predictions would have interesting medical applications, since several afflictions are now viewed as metabolic diseases ^98,99^.

These results show that the use of ML provides an alternative, effective, and scalable way to leverage large multiomics datasets to predict cell behavior, a fundamental question in biology ^25,44,100–102^ with important practical applications ^2–4,24,26^. We have shown that it is possible to accurately predict metabolic dynamics through data-driven (as opposed to mechanistic) approaches using a relatively small amount of data: time series for just ∼20 strains. This amount of engineered strains is much less than we originally estimated ^47^, and lies within the reach of modern biofoundries, cloud labs, and industry, and several academic labs. We expect that improved prediction accuracy can be obtained using data for more than ∼20 strains, based on previous calculations and observed out-of-distribution failure modes. An orthogonal way to improve KinDL may involve merging data-driven and mechanistic strategies to combine the advantages of both approaches: predictive power and mechanistic insight. Whereas the traditional molecular biology expectation has been to find mechanisms first and then use them to enable predictions (through, e.g., kinetic models), this approach has been particularly inefficient in making predictions of complex systems such as cells. Perhaps a more fruitful approach would involve seeking accurate predictions first and then determining causal mechanisms. Possible approaches to this end include causal structural models ^103^ and closed mathematical forms obtained through Bayesian inference^104^.

The use of this type of ML-guided approaches can be the disruptive advance needed to shorten the 366 years we estimate are needed to enable a bioeconomy. Indeed, we estimate that, at current discovery rates, creating a bioeconomy by replacing the ∼3500 high volume chemicals used nowadays ^105^, derived mainly from petrochemicals, would take on the order of ∼366 years (75 - 2390 years, see “Back-of-the envelope estimate of the time to establish a bioeconomy” in Methods). With rising concerns of microplastic presence in human organs ^106^, a temperature anomaly growing by the year ^107^, and stiff competition in biomanufacturing ^108^, 366 years seems unacceptably long.

In sum, we believe that ML approaches like KinDL have the potential to radically change how Design-Build-Test-Learn cycles are used in industry and academia. This change is bound to significantly reduce the development time of new synthetic biology products, paving the way for a bioeconomy that provides enhanced-performance products, diminished supply chain disruptions, environmental benefits, solutions to national security needs, and new jobs and industries.

## Methods

### Strain construction

#### Base strain and initial strains

Initial engineered malonic acid production strains were derived from *Pichia kudriavzevii* (Pk) YB-431 from the U.S. Department of Agriculture ARS strain collection ^109,110^. These strains possess a codon-optimized Cas9 gene from *Streptococcus pyogenes* in an expression cassette under a Pk TDH1 promoter and *Saccharomyces cerevisiae* (Sc) TEF1 terminator, integrated heterozygously in single copy in the diploid genome of Pk in the ADH3b locus, and are histidine auxotrophs due to lack of HIS3. The strain descriptions for these DBTL1 strains (Fig. 1) can be found in Supplementary Table 1 and Supplementary Note 1.

#### Strain engineering to instantiate recommendations

The initial strains from DBTL1 were further engineered according to the recommendations provided by the machine learning approach (Fig. 1, Supplementary Table 2, Extended Data Fig. 8), which involved combinations of knockouts and overexpression targets. We intended to use integration cassettes and modify promoters as needed, in practical terms limiting recommendations to combinations of one knockout (where the cassette would integrate) and up to three up or down regulations (modifying promoter strength), with constraints as described below in “Supported modifications”. For knockouts, a cassette was to be integrated at the gene locus to be knocked out, eliminating its expression. For upregulations combined with the knockout, copies of the genes were to be added to the cassette. Downregulations were to be produced through promoter strength alteration. Hence, we expected knockouts to eliminate protein expression and upregulations to approximately double protein expression (since the gene copies were to be effectively doubled), whereas downregulations were expected to approximately halve protein expression through the characterized activities of selected promoters. Given these constraints, all recommendations produced involved a single knockout target and three overexpression targets (Supplementary Table 2). Integration cassettes were hence designed with the 3 overexpression targets under the Pk ENO1 promoter and Sc TPI1 terminator, Pk FBA1 promoter and Sc PYC2 terminator, Pk GPM1 promoter and Sc HXT1 terminator, respectively, in addition to containing an internal lox66/lox71 site flanked Pk HIS3 gene under a Pk TEF1 promoter and Sc TDH3 terminator as a selection marker. Each individual expression unit was cloned into independent vectors. Two guide RNA sequences targeting the integration site in the gene to be deleted were cloned into an expression cassette in a template vector. The guide RNA sequence was fused to a cRNA sequence and the coding RNA was flanked by a Hammerhead-HDV ribozyme system ^111^, and under control of a Pk TDH1 promoter and Sc TEF1 promoter. The full gRNA cassettes were amplified as linear DNA fragments. For in vivo assembly, the 3 gene overexpression cassette plus lox66/lox71-flanked Pk HIS3 cassette were amplified as 4 separate linear DNA fragments from plasmid templates, with 30-81 bp homology to neighboring fragments or the sequence immediately flanking the integration locus introduced via the oligonucleotide primers. All 6 linear DNA fragments were combined and transformed into Pk using a standard method ^112^, and cells were plated on CSM-His agar to select for integrants. Colonies were replated on YPD agar and screened by PCR to identify homozygous (two copy) integrants.

The final twenty two strains generated can be found in Supplementary Table 2.

### Strain fermentation

A seed train was used to build biomass for fermentation. The strain was first plated on YP + 2% glucose agar plates and incubated at 30°C until single colonies formed. Using a sterile inoculation loop, a single colony was inoculated into a 250 mL baffled flask containing 50 mL of YP + 2% glucose. The seed culture was grown for 18-24 h before 3 mL of culture was inoculated into the bioreactor vial sterile syringe and needle through the headplate.

Strains were then cultured in the Ambr® 250 High Throughput bioreactor system (Sartorius Stedim ^113^), a platform widely known to produce results representative of those found at industrial scales. Twenty four single-use Microbial vessels (Sartorius PN: 001-5G13) with two rushton impellers, pH probes and dissolved oxygen spot readers were used for this experiment. A maximum oxygen transfer rate (OTR) of 200 mmol/L/hr was targeted, achieved by setting the stir rate to 3000 rpm and the air flow to 100 mL/min. The temperature was maintained at 30°C. As volume increased over the course of the run the OTR was automatically maintained, first by increasing the stir rate and then by increasing the percentage of oxygen in the gas flow up to 30% through oxygen supplementation. The pH of the culture was maintained through automatic addition of 5 M ammonium hydroxide (no acid was used). Foaming was controlled through the addition of a 5% Struktol SB2121 in 120 proof ethanol. Proprietary defined batch and feed media containing glucose, salts, trace metals and vitamins were used in this experiment. A fed batch feeding strategy was carried out through the automatic addition of feed in a pulse fed manner. Feed pulses were automatically added when proprietary conditions were met.

### Multiomics data collection, quality check, and augmentation

#### Sampling

Samples were taken at eight different time points: 10, 22, 28, 36, 46, 52, 60, 72 hrs after inoculation of the bioreactor. The volume of the samples changed according to the cell density (higher ODs in later times required less volume to get the same signal), such that volume (ml) times the OD equaled approximately 15 (see below).

#### Proteomics

Cells (about 15 OD x mL units) from samples were harvested in triplicate by centrifugation at 5,000 g for 3 min and kept at -80°C until sample preparation. Pk cells were lysed and protein extracted as described previously ^39^. Briefly, the protocol consists of Zymolyase treatment, cell lysis, protein precipitation, protein resuspension, protein quantification, and normalization of protein concentration followed by standard bottom-up proteomic procedures of reducing and blocking cysteine residues and tryptic digestion. Peptide samples were analyzed on an Agilent 1290 UHPLC system coupled to an Agilent 6460 triple quadrupole MS (Agilent Technologies) as described previously ^114^. Briefly, peptide samples were loaded onto an Ascentis® ES-C18 Column (Sigma–Aldrich) and were eluted from the column using a 10 minute gradient from 95% solvent A (0.1 % FA in H2O) and 5% solvent B (0.1% FA in ACN) to 60% solvent A and 40% solvent B. Eluted peptides were introduced to the MS using a Jet Stream source (Agilent Technologies) operating in positive-ion mode. The data, quantified as relative “counts” signal detected by the Agilent High Energy Dynode detector, were acquired with Agilent MassHunter Data Acquisition software operating in dynamicMRM mode. MRM transitions for the targeted proteins were generated by Skyline software (MacCoss Lab Software) and selection criteria excluded peptides with Met/Cys residues, tryptic peptides followed by additional cut sites (KK/RR), and peptides with proline adjacent to K/R cut sites. The data and Skyline methods are available on Panoramaweb (https://panoramaweb.org/Marti_et_al_prediction_metabolic_dynamics deep_learning.url) and data are available via ProteomeXchange with identifier PXD059887.

#### Metabolomics

##### Intracellular metabolites

Between 0.4-1.2 mL of culture (with smaller volumes as cell density increased later in the fermentation) were collected into pre-chilled 15 mL Falcon tubes containing 3x the culture volume of methanol in an isopropanol/dry ice bath. Samples were quickly mixed by inverting and centrifuged at 4000 rpm and -10°C in a Beckman Allegra X-15R centrifuge equipped with a swinging bucket rotor. The supernatant was discarded and cell pellets were stored at -80°C until extraction. For extraction, 0.5 mL of 7:3 (v:v) methanol:water (LC/MS grade) was added to the cell pellets and samples were mixed by vortexing for 3 minutes. Three technical triplicate samples of 0.5 mL for each strain/sampling point were transferred into a pre-chilled (in a dry ice/isopropanol bath) Cryo-Adapter for 2210 titer plates (SPEX SamplePrep) containing a 96 well deepwell plate and sealed with a Cap-Mat for 2210 titer plate (SPEX SamplePrep). The pre-chilled accessories and deepwell plate were then fit into a FastPrep96 bead beater (MP Biomedicals) for 12 cycles at 1800 rpm, 20 seconds per cycle and 1 minute between cycles in a dry ice/isopropanol bath. After bead beating, the plate was centrifuged at 4000 rpm for 2 min at -10°C, and 200 uL of supernatant was transferred to a 96 well 0.22 um filter plate over a Waters HPLC collection plate. This was centrifuged again at 4000 rpm for 10 minutes at -10°C, foil sealed using an Agilent PlateLok plate sealer, and stored at -80°C until analysis. Intracellular metabolite samples were analyzed on an Agilent 1290 UHPLC system coupled to an Agilent 6460 triple quadrupole MS (Agilent Technologies). Briefly, metabolite samples were loaded onto an Agilent HILIC Column (Agilent) at initial composition of 5% solvent A (10 mM Ammonium acetate at pH 9.0 in H2O) and 95% solvent B (10 mM Ammonium acetate at pH 9.0 in ACN). Metabolites were eluted from the column using a 2.9 minute gradient by ramping up solvent A to 25% in 2.5 minutes and further increasing to 50% in 0.4 minutes. Eluted peptides were introduced to the MS using a Jet Stream source (Agilent Technologies) operating in negative-ion mode. The data, quantified as relative “counts” signal detected by the Agilent High Energy Dynode detector, were acquired with Agilent MassHunter Data Acquisition software operating in dynamicMRM mode. MRM transitions for the targeted proteins were generated by Skyline software (MacCoss Lab Software). The data and Skyline methods are available on Panoramaweb (https://panoramaweb.org/Marti_et_al_prediction_metabolic_dynamics deep_learning_metabolites.url).

##### Extracellular metabolites

Samples were prepared by obtaining supernatant via centrifugation, filtered at 0.2-0.4 um, followed by 40-fold or 10-fold dilution in 0.1N HCl, prior to analysis. An Agilent 1200 HPLC instrument using isocratic mobile phase consisting of 5.5 mM H2SO4 in DI water at 0.8 ml / min was used for this method. The HPLC column was a Bio-Rad Aminex HPX-87H (7.8 mm I.D. x 300 mm) with detection by both RID and UV. The HPLC method was used to quantify malonic acid, D-glucose, D-trehalose, L-arabitol, citrate, pyruvate, erythritol, succinate, glycerol, acetate and ethanol from fermentation broths and similar liquid samples. This method is suitable for quantitation over the range of 0.5 - 50 mM for each target analyte. Because of time and budget limitations, extracellular metabolites were measured in single replicates.

The Total Malonic Acid Formed (TMAF) accounts for all the malonic acid produced in the fermentation run. Because this experiment was conducted in an AMBR250 system(™), where sample volumes are relatively large compared to the working volume, the amount of product removed from the system through sampling needed to be accounted for. This is analogous to the difference between a batch fermenter and a chemostat fermenter which operate at the same volume and reach the same titer after a certain time: although the titers will be the same from that time on, the total product produced by the chemostat will increase over time (due to the extracted broth) while the total product produced by the batch fermenter will remain constant. For calculating the TMAF we adjusted the working volume using a mass balance approach at each time, which changed due to volume increases (due to, e.g., carbon source input or pH corrections) or decreases (due to sampling or volume removal to avoid going beyond a maximum working volume). The concentration of malonic acid (measured as indicated above) times the volume extracted constituted the product removed and was added to the product in the broth for the final point (= volume times the product concentration). TMAF results showed very good reproducibility, so measurements were done in single replicates. TMAF measurements in DBTL2.1 and 2.2 confirmed this excellent reproducibility when measured in triplicate and were used to estimate error bars in DBTL1 (Extended Data Fig. 8).

#### Database data upload and quality check (QC)

Proteomics and metabolomics data were uploaded to the Experiment Data Depot (EDD ^71^) for centralized storage and facilitation of the subsequent data quality check and analysis ^72^. We implemented a basic data version control system to detect changes in the different versions of the dataset that were uploaded to EDD. We performed data integrity and data quality checks and cleaned the data before using it to train machine learning algorithms. We looked for negative values due to calibration tolerances (found only 105 out of a total of 77593 records) and set them to zero. Any NaN (Not a Number) resulting from measurements under sensitivity were also set to zero. Outliers within a time series were found and eliminated so they could be filled in by the interpolator. We standardized the way to represent missing values as missing records and searched for missing data through different strains and over time. For each one of the multiomics datasets, we checked for potential duplicates in measurements, detected data gaps including premature end of the time series, and identified full missing datasets of strain, replicate, and sampling time. The data gaps were fixed by interpolation or extrapolation using the same methods used for data augmentation and described in the next section.

#### Data augmentation and normalization

The amount of collected data was at the lower limit of what deep learning can leverage for supervised learning. To enable the use of deep learning, we augmented the data in a similar way that we had previously done ^47^: by interpolation. In this way, we resampled the data from 8 to 63 timepoints (1 interpolated datapoint per hour from 10 hr to 72 hr). We found that a piecewise cubic polynomial interpolator, which is twice continuously differentiable, with boundary conditions imposed so that the first derivative at curve ends are zero, provided in our case the best results to minimize Runge’s phenomenon, in which edges see strong oscillations at the edges for the interpolation when using polynomial interpolation. To reduce the number and, especially, the magnitude of negative values resulting from the interpolation, we applied a logarithmic transformation, selecting log(1+x) as it was both numerically stable and efficiently implemented in NumPy ^115^. The transformation reduced the number of negative values to roughly ⅔ and, more usefully, the minimum value moved up 5 orders of magnitude from below -40,000 to -0.8 (with most of the negative values then over -0.05), which allowed directly rounding them to zero with negligible effects regarding the interpolation quality. For each feature and target time series, we therefore applied a log(1+x) transformation, applied cubic-spline interpolation (pandas function df.interpolate, using the cubic spline method with ‘clamped’ boundary conditions, where the 1st derivative at curve ends are zero), and untransformed the data by applying an exp(x)+1 transformation. To this end, we contributed a cubic interpolator from Scipy ^116^, CubicSpline, to the pandas package ^117^, which is now available since the v.1.1 stable version of pandas. Because we only had one set of bioreactor measurements (including target variable TMAF) for each strain, but -omics measurements in triplicate, we used the same bioreactor measurements for all three replicates. The interpolation strategy, combined with using all replicates as training instances instead of just using averages, increased our training data from ∼80,000 data points to ∼500,000.

Finally, we normalized the feature and label data before using it for neural network and PLS modeling. For that purpose, we created a new class StdSet wrapping the Dataset class from TensorFlow’s data module with additional methods to apply or revert a min-max normalization with calculated parameters for the new data or with those previously calculated for the data of another instance of StdSet.

### Machine learning for predicting final product and metabolite concentration time series

To predict time series of production (TMAF) and metabolite concentrations from time series of protein concentrations we used three different architectures of deep neural networks ^118^ and a partial least squares model ^119^ (Fig. 3). We used a supervised learning approach, with input being the time series of proteomics data and, as response, the time series of TMAF or metabolite concentrations we were interested in predicting (Fig. 3). This line of action differs from our previous approach (kinetic learning^47^), which aimed to relearn the Michaelis-Menten kinetics (i.e. relationship between the metabolite change and the concentrations of proteins and metabolites). Instead we chose to directly map proteomics time series to metabolite time series, eliminating the need to integrate the ordinary differential equation (a source of significant error propagation). We restricted the input to proteomics data because they are relatively stable (Fig. 2) and also more actionable variables: through the wise choice of stronger or weaker promoters, it is, in principle, possible to qualitatively change protein concentration as desired (i.e., increase or decrease), whereas we had no easy way to change the metabolite concentrations in a prescribed manner. We used three different architectures of deep neural networks ^118^ and a partial least squares model ^119^ to predict response from input features (Fig. 3). The PLS model represented a simple and fast approach to suggest recommendations, while the weighted average of deep neural networks provided much more accurate response predictions to check and correct PLS recommendations. The PLS regression model was trained to predict final TMAF from proteomics data, using the scikit-learn implementation ^120^. The first basis vector from the PLS regression is the direction that explains the most covariance between the proteomics and TMAF data, with subsequent vectors explaining progressively less covariance. The initial codebase used to produce the fits and recommendations was consolidated into a clean library: the KinDL library that can be found in the Code Availability section.

#### Artificial neural network architecture

Our ensemble of neural networks consisted of 1) a network consisting of densely (fully) connected layers (FCNN); 2) a network of connected recurrent neural network layers (DRNN); and 3) a network of connected gated recurrent unit (GRU) layers. The input to each neural network was the same (Fig. 3): a time-series of proteomics data paired with either a single time series of TMAF (for TMAF predictions) or time series of all metabolites (for metabolite predictions). Our intention was for the deep hidden layers to capture core metabolism and for subsequent layers to use this to predict each metabolite’s dynamics (Extended Data Fig. 3). All proteomics, metabolomics, and TMAF time series were preprocessed as described previously (smoothed and interpolated from 8 measured time points to 69 interpolated time points). The individual neural networks are described in detail below, and their architecture and hyperparameters (number of hidden layers/nodes in each model, learning rate) are described in Supplementary Tables 7-10. All neural networks were trained for 100 epochs, with early stopping (patience = 10 epochs) and used the model parameters from the best performing epoch. We also iteratively reduced the learning rate by a factor of 10 for each model (see below for each model’s starting learning rate) during training if the loss plateaued for 5 epochs. All neural networks were created and trained using Keras TensorFlow v2.2 ^121^.

The FCNN model (Supplementary Table 7) consisted of an input layer, which converts the input data into a Keras tensor, a flattening layer which converts the input from a 2D matrix of time series data to a 1D vector, 4 hidden layers consisting of 276 (4*the number of time points) nodes with a rectified linear unit activation function, and an output layer with 69 nodes (one for each time point) and a linear activation function, which produces the predicted time series. The model was trained using the Adadelta optimizer with a learning rate of 1 and a rho value of 0.95. The mean squared error (MSE) between the predicted time series and the true time series was used as a loss function:

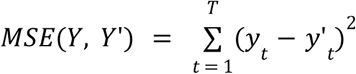

where Y is the observed time series, y_t_ is a single observed value in the time series, Y’ is the predicted data, and y’_t_ is a single predicted value in the time series

The DRNN model (Supplementary Table 8) consisted of an input layer, and no flattening layer was included because RNNs operate on the full time series. The input layer was connected to 3 keras.layers.SimpleRNN layers with 130 nodes each and a final simpleRNN layer with 27 nodes. The SimpleRNN layers all use the default tanh() activation function and return sequences. The final SimpleRNN layer feeds into a Dense layer which is distributed across the time series and has a linear activation, producing a predicted time series. The DRNN model was trained with the ADAM optimizer with a learning rate of 0.001 and the same MSE loss function used previously.

The GRU model (Supplementary Table 9) consisted of an input layer, and no flattening layer was included because the GRU nodes operate on the full time series. The input layer was connected to 3 keras.layers.GRU layers with 97 nodes each and a final simpleRNN layer with 46 nodes. The GRU layers all use the default tanh() activation function and return sequences. The final GRU layer feeds into a Dense layer which is distributed across the time series and has a linear activation, producing a predicted time series. The GRU model was trained with the ADAM optimizer with a learning rate of 0.001 and the same MSE loss function used previously.

For all models, when training to predict all metabolites, one dense output layer for each metabolite was used, providing the output for a specific metabolite time series. Our expectation was that deep hidden layers may capture a representation of core metabolism that can then be leveraged to predict the time series for extracellular and intracellular metabolites (Extended Data Fig. 3).

#### Hyperparameter optimization

Hyperparameter optimization was performed using the BayesianOptimization tuner from KerasTuner scalable hyperparameter optimization framework ^122^. The dataset was first pre-processed as described in the previous subsections. K-fold cross-validation (k=10) was directly implemented in TensorFlow using the StdSet class (see “Data augmentation and normalization” subsection) through a derived class wrapping Keras’ Model class. Using KerasTuner, a hypermodel was defined to explore specific ranges for various hyperparameter configurations.

The BayesianOptimization tuner was selected for its efficiency in exploring the hyperparameter space by building a probabilistic model of the objective function and selecting promising configurations based on previous evaluations using the mean squared error between observed and predicted time series for a validation dataset as the optimization metric. The search space included the number of layers (range: 1-3), units per hidden layer (range: 32-1024), and units per target layer (range: 16-128). The tuner performed 64 trials maximum with at most 3 executions per trial, iteratively adjusting the hyperparameters and evaluating model performance using k-fold cross-validation to ensure robust validation. The final model, configured with the optimal hyperparameters, was trained on the complete training dataset before evaluation on the test set to assess generalization performance. The optimal hyperparameters and training parameters are detailed in Supplementary Table 10.

#### Ensemble modeling

After training each ANN model, we aggregated the predictions from multiple models to generate a final prediction of the response (Fig. 3). The final version of the aggregation involved a weighted average of the three neural network predictions, in which the weight was the inverse of the training mean absolute error of each model.

#### Testing prediction accuracy via the coefficient of determination

Given a relatively low-data regime (for neural networks), we decided on an approximate 1:5 ratio between test and training data. The separation was at the strain level, with all the three technical replicates of each strain going into the training or the test set. The only two biological replicates in the dataset, LPK15_14087 and LPK15_14087b, were both assigned to the training set. The four test strains (Supplementary Table 1) were selected to cover a range of TMAF values.

We evaluated predictive accuracy by calculating the coefficient of determination ^87^ (R^2^) between predicted and measured time series for both training and test datasets (Fig. 4, 5, 6). For the TMAF we also calculated R^2^ for the final TMAF levels (Fig. 4B). The coefficient of determination R^2^ is the fraction of the variance in the response that is predictable from the input variables. Hence, it provides a single number that can be used to compare predictions across different datasets and prediction methods: a value close to one indicates very good predictions (almost all response variance predictable from the input variables), and values close to zero or negative indicate no predictive power. The values between zero and one have been shown to be more informative than other alternatives ^87^, and represent a more stringent option than using the correlation coefficient. Indeed, R^2^ corresponds to the square of the correlation coefficient ^87^ (for a linear regression): a correlation coefficient of 0.9 corresponds to an R^2^=0.81, and an R^2^=0.9 to a correlation coefficient of ∼0.95.

#### Calculating feature importance

We identified the ten most important input features (proteins) through SHapley Additive exPlanations (SHAP) analysis ^77^ and tested the predictive power of KinDL when using only those ten proteins as inputs (Extended Data Fig. 7). Shapley values are calculated by comparing the model output when a given feature is included or excluded (i.e., setting its value to the average of all observations). To calculate Shapley values for each timepoint and protein we used the shap.explainer() method from the SHAP package ^77^. To calculate Shapley values for individual proteins across the entire time series, we added up the Shapley values at each timepoint for each protein.

#### Predicting TMAF using single timepoint proteomics data

A comparison with the Automated Recommendation Tool (ART), a ML tool successfully used in the past to predict single response points ^49,78,79^, shows that single point predictions are much less accurate than those we obtained by leveraging the time series (test data R^2^ = 0.60 from ART vs 0.96 from KinDL, Fig. 4b and Extended Data Fig. 6). ART uses a linear ensemble model where the coefficients for each model are probability distributions, generating predictions which are themselves probabilistic. We used the proteomics measurements from the final timepoint as input (features) to predict the final TMAF (response), using the same train-test split as when we trained KinDL. Since the ART model is probabilistic, we calculated final predictions of TMAF/**μ** by taking the mean of the output distribution. Given the high-throughput proteomics pipelines that are starting to become available ^39^, these results suggest that collecting time series of data can be an effective way to enlarge the training dataset and improve the accuracy of ML predictions, since strain construction is still often a bottleneck.

### Recommendations

#### Supported genetic edits

To generate the recommendations for genetic edits the use of the predictive model had to be tailored to the genetic interventions that could be created in a high-throughput fashion by the strain building process in Lygos. Three basic modifications of protein expression were supported by the synthetic biology strain construction process: knockouts, downregulations, or upregulations. We assumed that upregulations would result in a doubling of the protein concentration (through the addition of an extra copy of the gene in the integration cassette), that a downregulation would halve it (through the choice of promoter of appropriate strength), and that a knockout would result in a complete lack of protein expression (see “Strain engineering to instantiate recommendations” above). Valid recommendations then involved combinations of one gene knockout (KO) and up to three genes to either down or upregulate, according to the following rules: (i) one knockout could be combined with up to three different upregulations; (ii) one downregulation could be combined with up to two different upregulations; and (iii) upregulations by themselves could be combined for up to three different proteins. Consequently, the simplest valid recommendations consisted of either a single knockout, a single downregulation, or a single upregulation, while the most complex recommendations involved one knockout combined with three upregulations. These combinatorial possibilities were implemented in the code using a general framework through a ’Plan’ class that implements a recursive data structure to represent all valid combinations. The implementation uses a ’Recommendation’ data class that contains one or more ’Modification’ objects following the rules defined by ’Plan’. Importantly, recommendation objects are order-independent with respect to their constituent modifications.

#### Generating recommendations

Despite the constraints above, the possible phase space for recommendations was too large to predict the result for all combinations in a practical amount of time. For this reason, we used the fast, but not very accurate, partial least squares (PLS, Fig. 3) model to rank which proteins were most likely to affect final TMAF, down selecting them to a reasonable number we could later easily enumerate. We then used the, slower but more accurate, ensemble model provided by the three ANNs (Fig. 3) to calculate the TMAF improvement for all possible combinations and ranked them accordingly. Specifically, we trained a PLS model with two components to predict TMAF from proteomics data, because the inclusion of three components only improved prediction accuracy minimally (R^2^ of 0.869 for three components vs 0.825 for two). Since both components were correlated with TMAF, we added the PLS weights together for each protein. Proteins were then ranked by their summed PLS weights, creating a sorted list from most positively to most negatively correlated with titer. The top 12 proteins from each extreme of this sorted list were selected for modification: those most positively correlated were considered for upregulation, while those most negatively correlated were considered for both downregulation and knockout.

After identifying candidate proteins, we modeled the effects of perturbations on DBTL1 strains by simulating the impact of protein modifications. We first simulated the effects of a combination by multiplying the targeted protein(s) from the measured data for that line by a factor *p*. Proteins were multiplied by factor *p*=0 for a knockout, *p*=0.5 for a downregulation, or *p*=2 for an upregulation (see “Strain construction” above). We then passed the modified protein inputs into all four predictive models (ANN, RNN, GRU, PLS) and obtained predicted TMAF dynamics from each. We calculated the final predicted TMAF dynamics by taking the weighted average of the three neural network predictions (see “Ensemble modeling” above). The final output of this calculation was the ratio of the final predicted TMAF for the genetic modification and the final TMAF for the base strain. This process was implemented through a recursive algorithm that evaluated all valid combinations of modifications (as per "Supported modifications"), while selecting proteins from the corresponding extremes of the correlation-sorted list (detailed in the previous paragraph) up to a maximum depth of 12 proteins from each extreme. The algorithm generated 26388 total predictions per base strain, each consisting of proposed genetic perturbations (e.g., one knockout combined with three upregulations, Fig. 1) and their predicted impact on final TMAF. For each strain, recommendations were the top 2500 of these predictions ranked by predicted TMAF improvement. From the recommendations for each base strain, Lygos metabolic engineers selected 22 strains for implementation based on expected build feasibility and strain diversity (Supplementary Table 2).

### Back-of-the-envelope estimate of the time to establish a bioeconomy

We estimated that establishing a bioeconomy that replaces the current high-production-volume (HPV) chemicals (i.e., manufactured in or imported into the United States in amounts equal to or greater than 500 MT/year ^105^) by renewable equivalents would take 366.5 years (with 75 - 2389.5 as the confidence intervals). To arrive at this number, we used a back-of-the envelope estimation approach. This approach is not expected to provide an exact number, but rather an order of magnitude estimate that can be used to rule out unrealistic options.

To obtain this number, we used the estimate of Wu *et al* ^105^ of 3574 HPV chemicals in the US. We multiplied this number by the number of average person-years to create a single renewable chemical, to find the total number of person-years it would take to replace all of them. The average person-years to synthesize a single renewable chemical is difficult to estimate, and we only have a few examples to do so: Dupont and Genencor needed 575 person years to develop and produce 1,3-propanediol ^27^, Amyris Biotechnologies took between 130 and 575 person years to make farnesene ^27^, and more than 150 person-years to produce artemisinin ^123^. That would give a range of 130 to 575 and an approximate average of 357.5 = (575+130+575+150)/4 person-years. We then divide the total number of person-years needed to produce a bioeconomy by the amount of persons working on this task. Again, this is a difficult number to estimate, and we used two different approaches: the total number of members of the Society for Industrial Microbiology and Biotechnology (SIMB ^124^), and a reasonable fraction of the total number of biological scientists in the US as provided by the U.S. bureau of labor statistics. We reasoned that the 850-1000 members of SBIM (personal communication) would be a good estimate for researchers willing and prepared to take on this challenge, given SIMB’s dedication to the advancement of microbiological sciences focusing on the practical applications of microbiology in industry. Due to the multiple assumptions involved in this estimate, we found an orthogonal way to do it: we assumed that 10% of biological scientists in the US would be available and trained for this challenge, should the need arise. This is likely an overestimate, since biology is a large discipline, covering a lot of fields from the study of human brain to oceanology. By May 2023, the U.S. bureau of labor statistics reported 61,220 biological scientists ^125^, 10% of which are 6,122. Hence, we estimated the number of researchers available for this task to be from 850 to 6122 people (average = 3486 = (850+6122)/2). The estimate for the number of years to establish a bioeconomy is then:

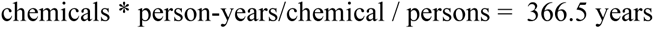

with confidence intervals given by 75 - 2389.5, obtained by using the worst and best case scenarios for the range of the numbers above: 75 = 3574*130 / 6122 and 2389.5 = 3574 * 575 / 850.

## Data Availability

The normalized experimental data for each cycle can be found in the Experimental Data Depot ^71^ in the following studies:

● DBTL1: https://public-edd.agilebiofoundry.org/s/beeps-lygos-ml-training-set-dec2019-2020-tmaf-614d/
● DBTL2.1: https://public-edd.agilebiofoundry.org/s/beeps-lygos-ml-cycle-2-dec2022-tmaf_normalize-a8bc/
● DBTL2.2: https://public-edd.agilebiofoundry.org/s/beeps-lygos-ml-cycle-2-feb2023-tmaf_normalize-0a77/

These data are also available as .csv files in the github repository (data/normalized_studies_from_edd/ folder) where code is made available (see “Code availability” section).

Production data were normalized by an undisclosed factor **μ** to preserve Lygos’ competitive advantage, but the terms of the ABF contract state that these data must eventually be publicly released (five years after data is produced) since this work was funded through public funds. Hence, the experimental data without normalization can be found in these EDD studies, which will be made publicly available on **2/2/28** upon the ending of the data embargo:

● DBTL1: https://public-edd.agilebiofoundry.org/s/beeps-lygos-ml-training-set-dec2019-2020-b223/
● DBTL2.1: https://public-edd.agilebiofoundry.org/s/beeps-lygos-ml-cycle-2-dec2022-7e33/
● DBTL2.2: https://public-edd.agilebiofoundry.org/s/beeps-lygos-ml-cycle-2-feb2023/

Freely available accounts on public-edd.agilebiofoundry.org are required to view and download these studies.

## Code Availability

KinDL (version 0.1) was developed in Python 3.11 and relies on the packages: keras-tuner 1.0.1, matplotlib 3.4.2, numpy 1.18.5, pandas 1.3.1, plotly 4.8.1, scikit-learn 0.24.2, scipy 1.7.1, seaborn 0.11.1, and tensorflow 2.2.3.

The KinDL library is released under a dual license that allows for free non-commercial use for academic institutions. A no-cost, one-year evaluation license is available to for-profit and nonprofit entities from Berkeley Lab. See https://github.com/AgileBioFoundry/KineticDeepLearning for software and licensing details.

## Acknowledgements

This material is based upon work supported by the U.S. Department of Energy’s Office of Energy Efficiency and Renewable Energy (EERE) under the Bio Energy Technology Office (BETO) and BioEnergy Engineering for Products Synthesis (BEEPS), Award Number DE-EE0008489. This work was also part of the DOE Agile BioFoundry (http://agilebiofoundry.org), supported by the U. S. Department of Energy, Energy Efficiency and Renewable Energy, Bioenergy Technologies Office, and is based upon work supported by the DOE Joint BioEnergy Institute (http://www.jbei.org), U. S. Department of Energy, Office of Science, Biological and Environmental Research Program, under Award Number DE-AC02-05CH11231 between Lawrence Berkeley National Laboratory and the U.S. Department of Energy. This work was also supported by the Basque Government through the BERC 2022–2025 program and by the Spanish Ministry of Science and Innovation MICINN (AEI): BCAM Severo Ochoa excellence accreditation CEX2021-001142-S.

Sandia National Laboratories is a multi-mission laboratory managed and operated by National Technology & Engineering Solutions of Sandia, LLC (NTESS), a wholly owned subsidiary of Honeywell International Inc., for the U.S. Department of Energy’s National Nuclear Security Administration (DOE/NNSA) under contract DE-NA0003525. This written work is authored by an employee of NTESS. The employee, not NTESS, owns the right, title and interest in and to the written work and is responsible for its contents. Any subjective views or opinions that might be expressed in the written work do not necessarily represent the views of the U.S. Government. The publisher acknowledges that the U.S. Government retains a non-exclusive, paid-up, irrevocable, world-wide license to publish or reproduce the published form of this written work or allow others to do so, for U.S. Government purposes. The DOE will provide public access to results of federally sponsored research in accordance with the DOE Public Access Plan.

## Author’s contributions

H.G.M., J.M.M., J.D., W.H. conceived the original idea. J.M.M., H.G.M., M.B., and P.H. developed the methodology, and designed the software. J.M.M., T.R., A.Z., and P.C.K. wrote the code, performed computer experiments, and analyzed results. M.H., A.A., C.J.P., H.H., R.M.L., A.C. and N.O. designed and coordinated experiments, and analyzed results. M.F. contributed to A.R.T. software maintenance and development. M.H., A.Z., A.C., Y.C., J.W.G., and D.M. performed the experiments. H.G.M., P.C.K., M.H., A.A., R.M.L., C.J.P, and J.M.M. wrote the paper.

## Extended Data Figures

**Extended Data Figure 1:**
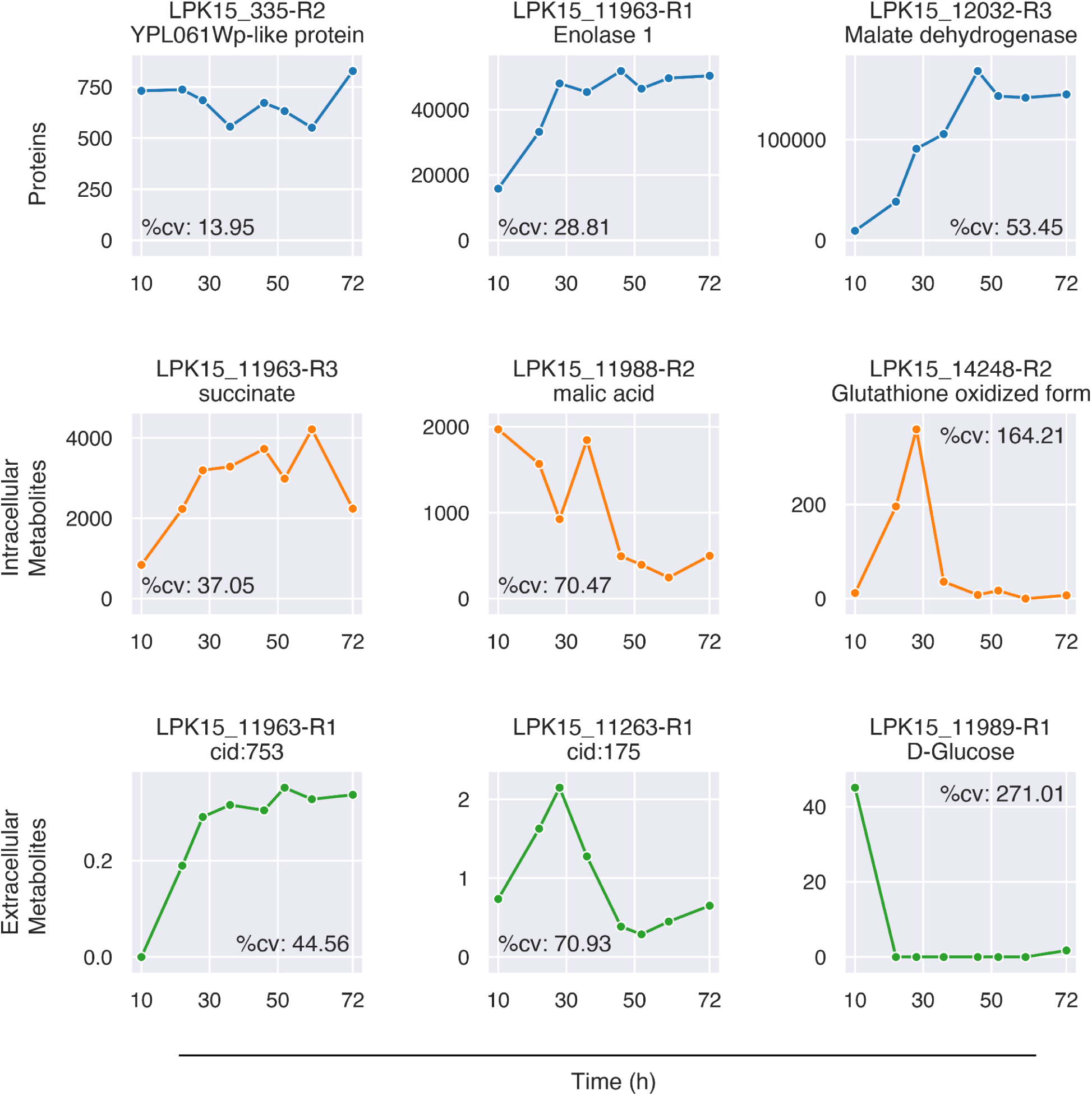
Specific examples of how proteins tend to be more stable in time than metabolites. Whereas Fig. 2 showed the distributions of coefficients of variation (cv, standard deviation over mean) for protein and intracellular/extracellular metabolites (analytes), here we show a few specific examples. We plotted the time series and corresponding coefficients of variation for several proteins and metabolites. For each analyte category (proteins, intra/extracellular metabolites), lines and analytes were sorted by their cv, and a low (10th percentile), median (50th percentile), and high (90th percentile) line-analyte combination are shown. In our nomenclature, “lines” represent a combination of a strain and replicate (e.g., LPK15_335-R2 involves the second replicate of strain LPK15_335). The lower cv for each percentile further shows that proteins tend to be more stable in time than metabolites.

**Extended Data Figure 2:**
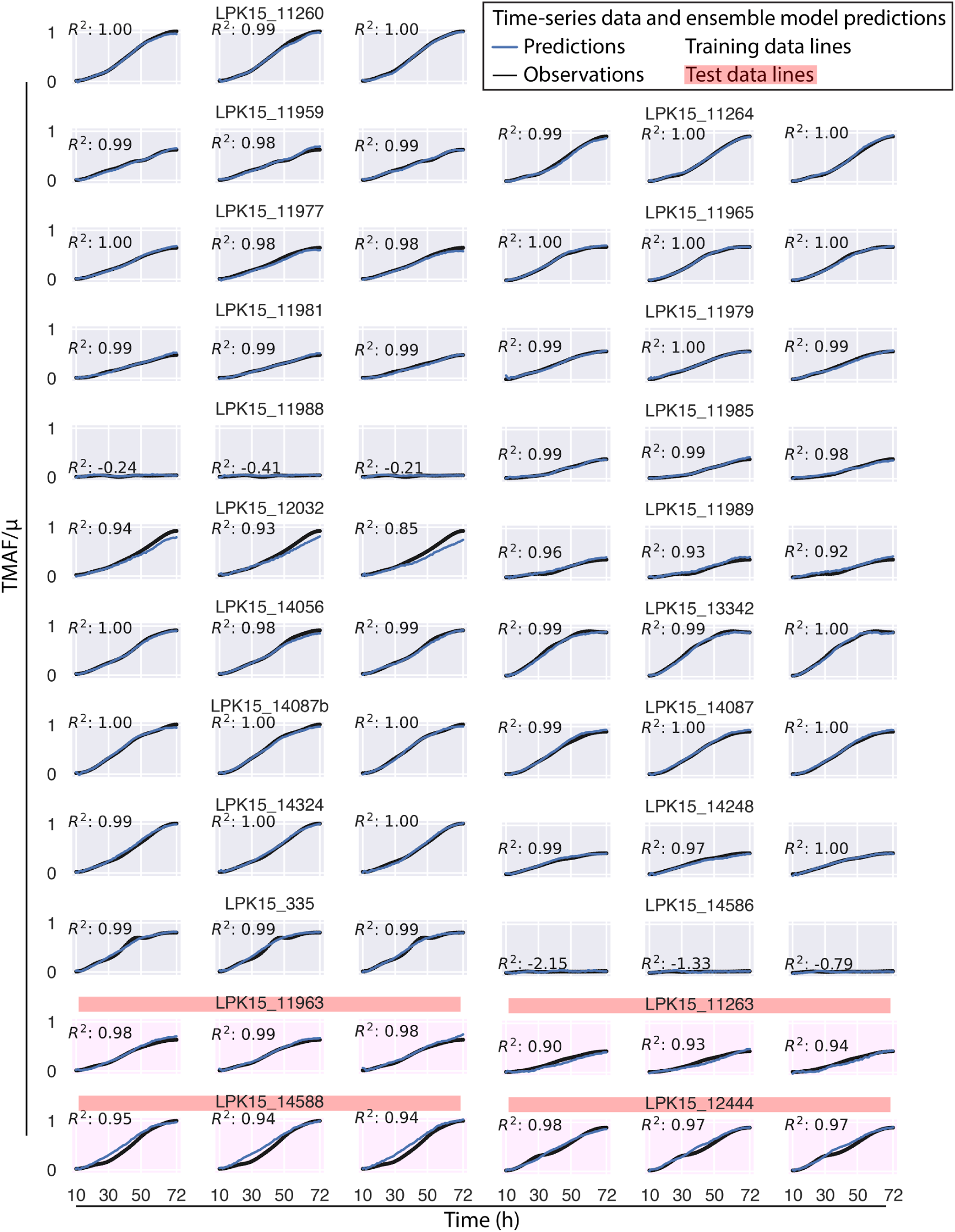
KinDL shows excellent prediction of Total Malonic Acid Formed (TMAF) times series for all lines. Predictions for training lines explain 98% of the data variance or more (R^2^≥ 0.98) for most cases (48/54 lines, 88%) and 85% of the data variance or more (R^2^≥ 0.85) for all cases, except for low almost constant values (for which R^2^ is not a good measure of predictive power). Predictions for test lines explain 90% to 99% of the data variance (0.90 ≤ R^2^ ≤ 0.99). Each of the three graphs for each line represents a replicate.

**Extended Data Figure 3:**
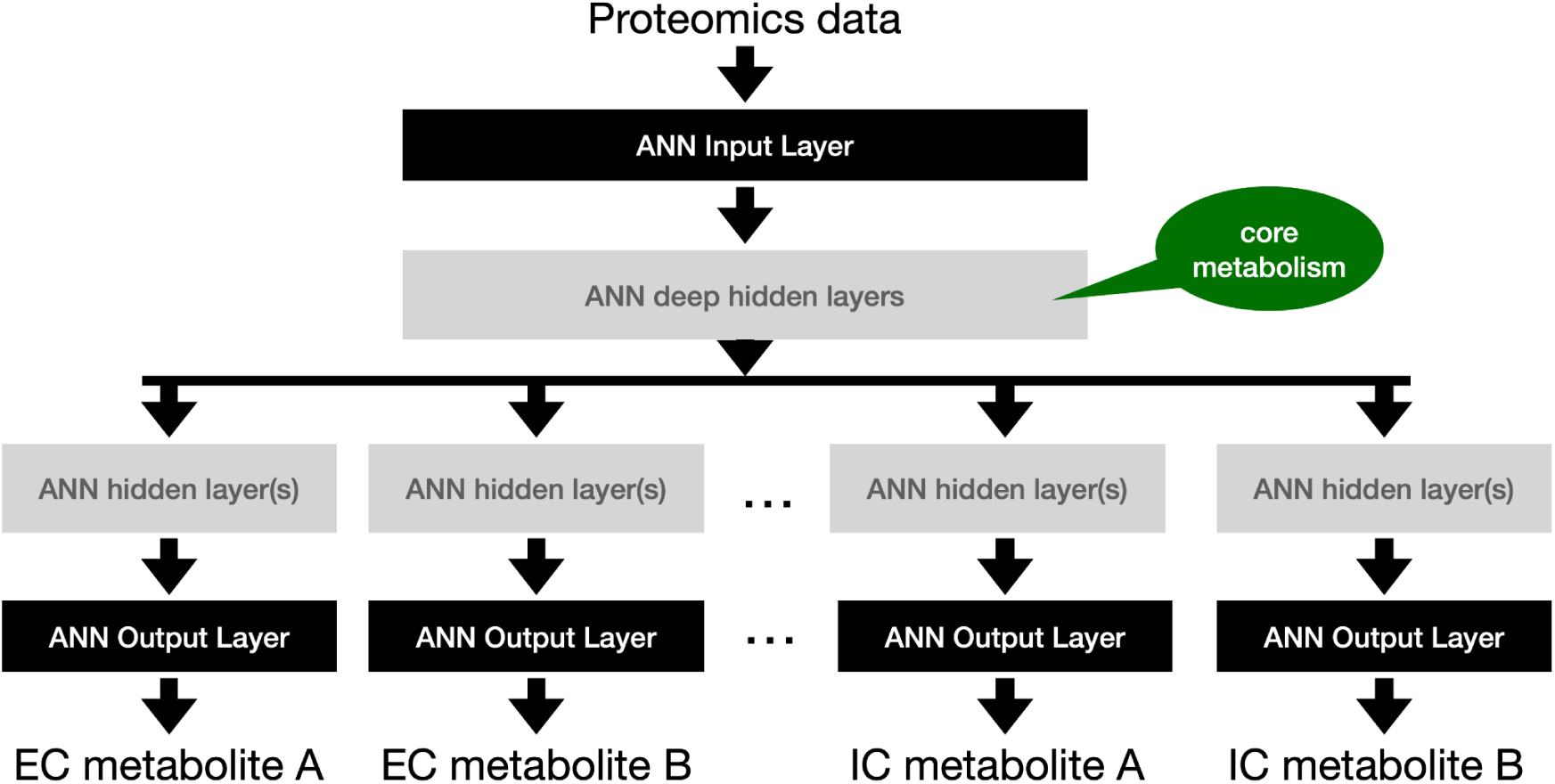
The neural networks used in this work were designed with the expectation that the ANN deep hidden layers would be able to capture a representation of core metabolism that can then be leveraged to predict the time series for extracellular and intracellular metabolites.

**Extended Data figure 4:**
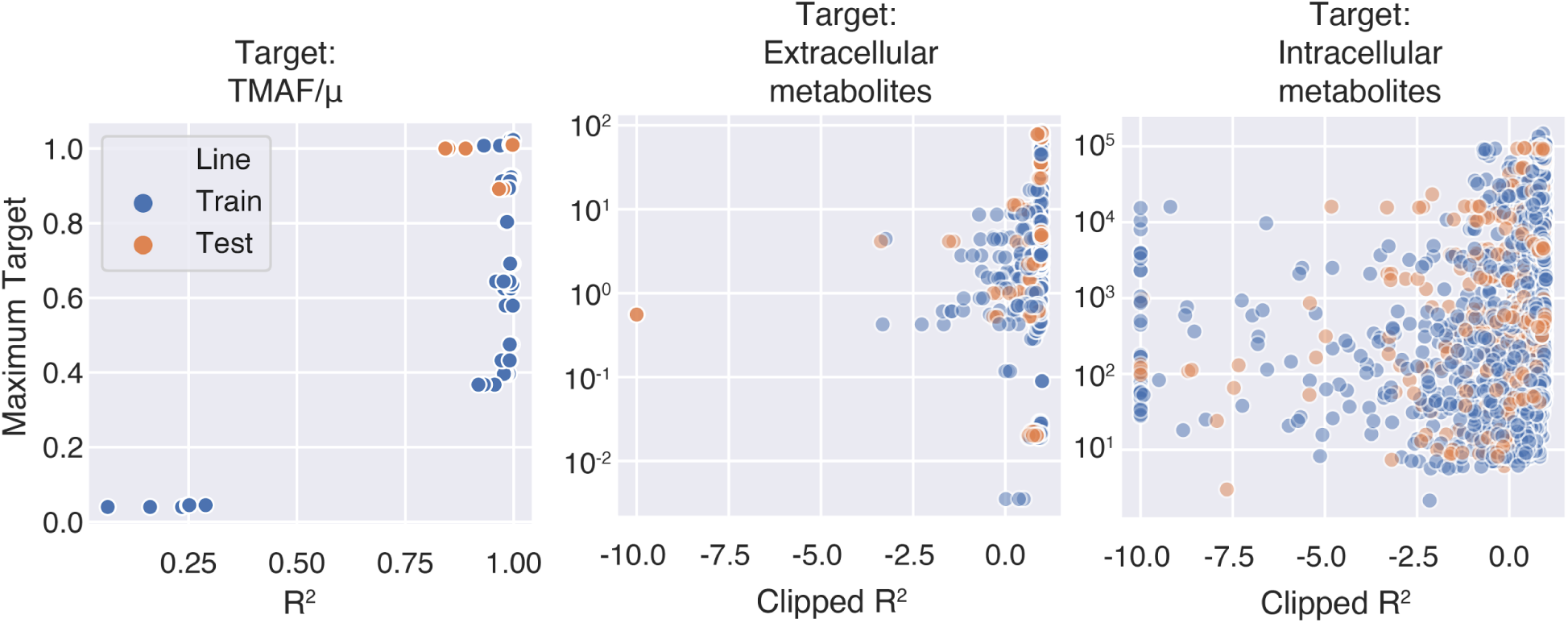
There is no dependence of prediction accuracy with concentration, except for TMAF. The y axis shows the maximum value of either TMAF/**μ**, or the counts for each extracellular or intracellular metabolite, whereas the x axis shows the corresponding value of R^2^ (or clipped R^2^, where all values below -10 are considered -10). All poor predictions for TMAF time series (low R^2^) are for low values of TMAF. Extracellular and intracellular metabolites, however, do not show that trend. Highly and low abundance metabolites are equally well or poorly predicted.

**Extended Data figure 5:**
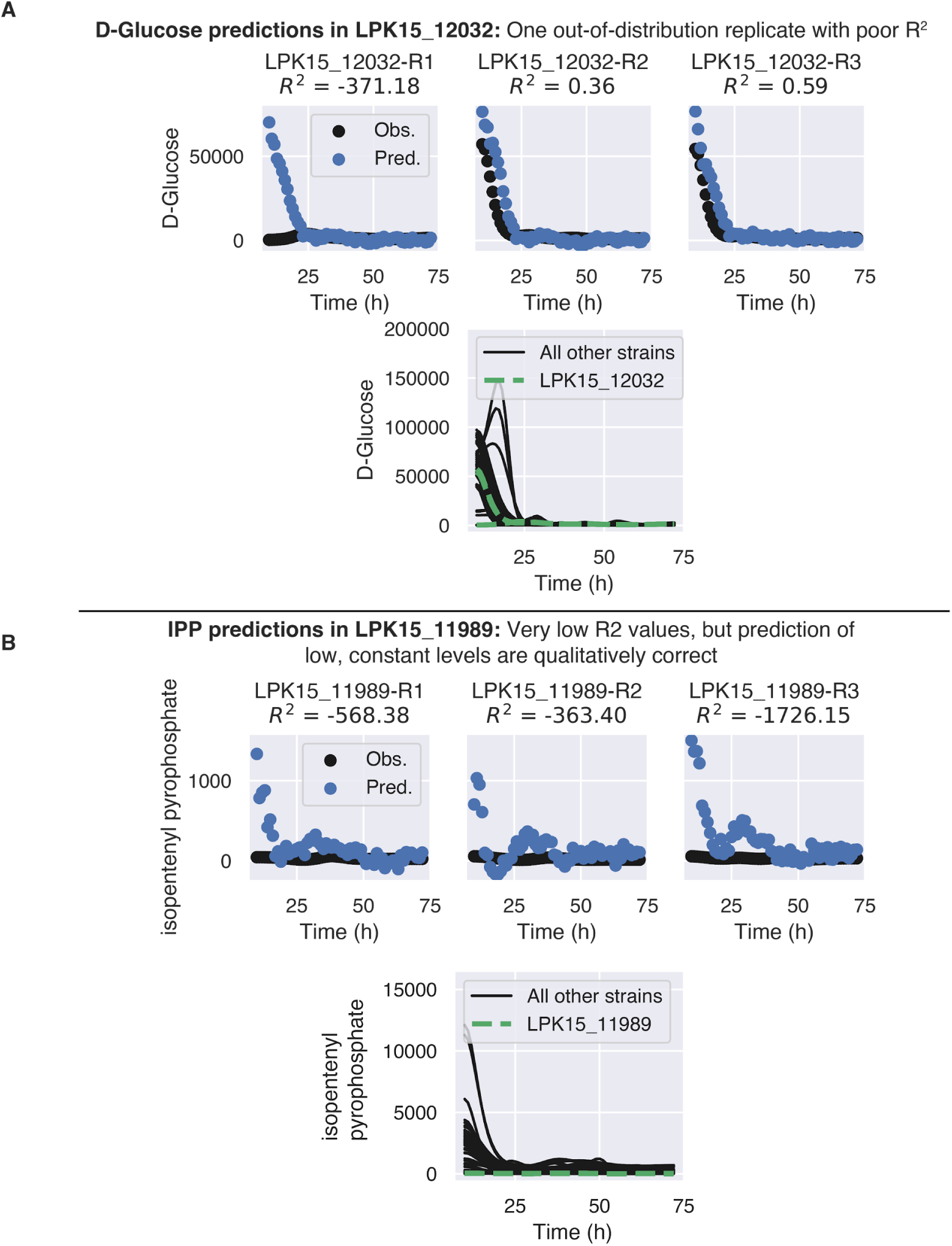
Examples of two specific cases in which predictions fail for different reasons. **A)** Out of distribution replicates: predictions for intracellular glucose in strain LPK15_12032 show that two of the replicates (R2 and R3) are very well predicted, but R1 is not because it has unexpectedly low glucose in the first measured timepoint. At the bottom graph we can see that the poorly predicted strain with a low starting intracellular glucose is out of distribution compared to other strains (in black). **B)** Low response values produce artificially low figures for R^2^: predictions for intracellular isopentenyl pyrophosphate (IPP) in strain LPK15_11989 show that predictions with extremely low R^2^ across all replicates are due to low, relatively constant, metabolite levels. The bottom graph demonstrates that the model predictions of very low IPP are qualitatively correct compared to other strains.

**Extended Data Figure 6:**
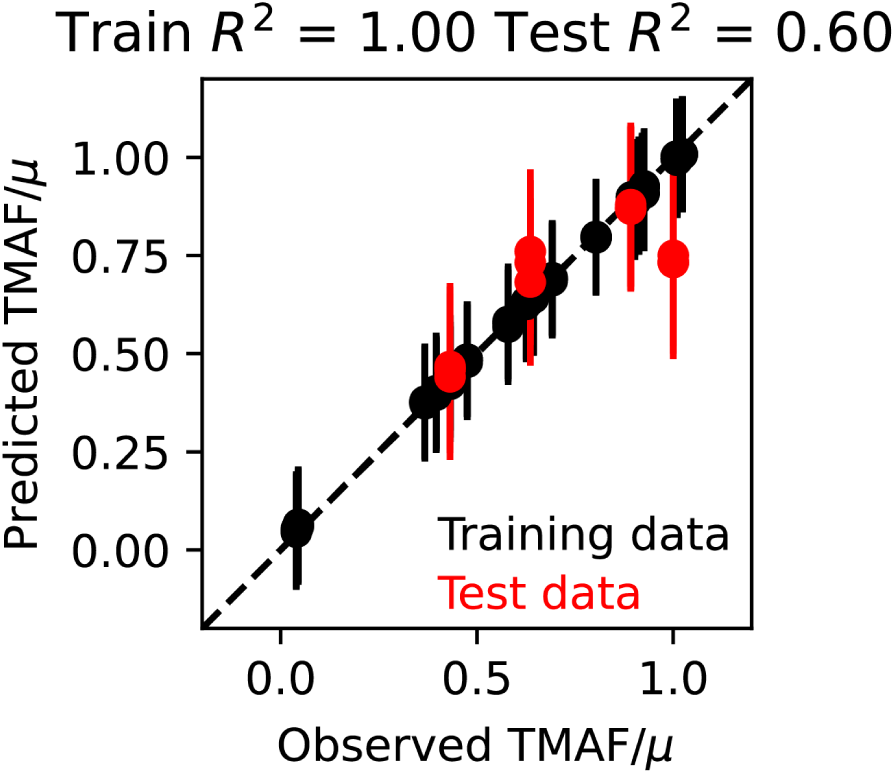
Predictions using final time points instead of the full time series provide worse results. We trained an ART model (see Methods) using the final time points of proteomics as input, and the final time points of TMAF/**μ** as response, for the same test set used for kinetic deep learning, and found the predictions to be less accurate (R^2^ = 0.60 from ART vs 0.96 from KinDL, Fig. 4B).

**Extended Data Figure 7:**
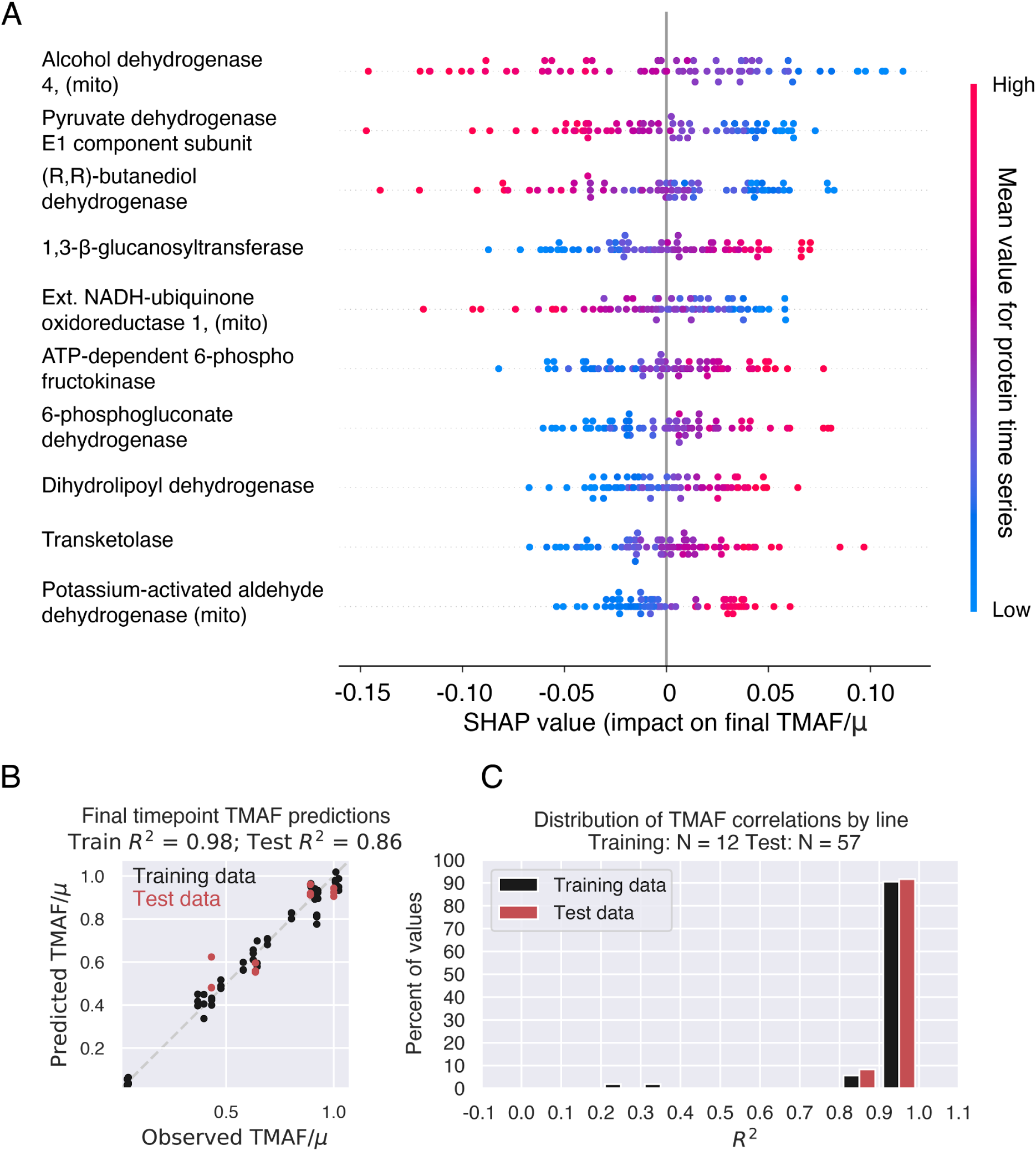
KinDL trained on only the 10 most important proteins produces similar predictions for full time series and final time points. After training KinDL on the full set of proteomics dynamics, we used SHAP analysis to identify the 10 most important proteins for predicting final TMAF production (see Methods). **A)** Distribution of SHAP values for the 10 most important proteins (one point for each final TMAF value), which are mostly dehydrogenases. **B)** Final point TMAF predictions based on the 10-protein model are worse than the full model by a small amount (in test data R^2^= 0.86 vs 0.96, Fig 4B). **C)** Predictions for TMAF time series for the 10-protein model are approximately the same as for the full model (similar distribution of R^2^ for both cases, Fig. 4C). However, the identity of the ten most informative proteins can only be found *a posteriori* in this manner, and hence can only be practically useful for follow-up DBTL cycles.

**Extended Data Figure 8:**
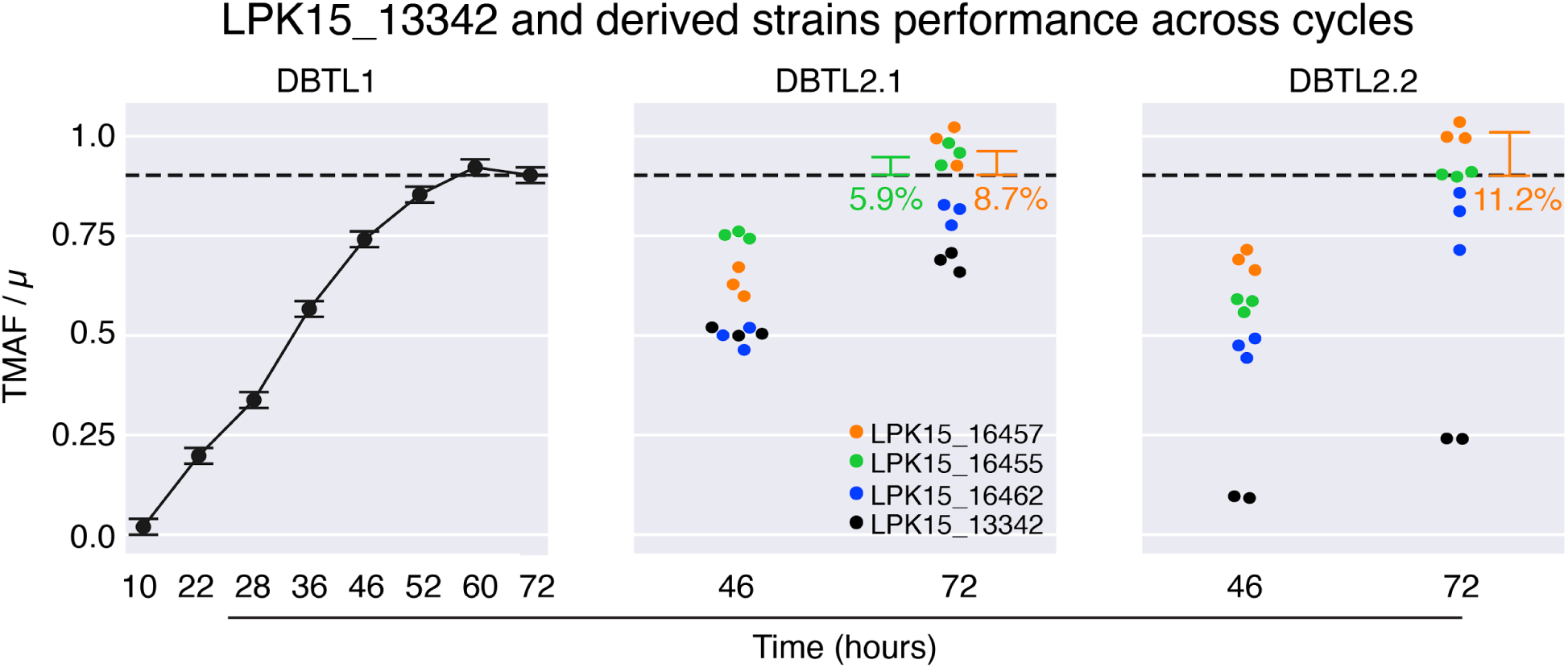
The recommendations provided by kinetic deep learning improved malonic acid production ∼20-440% with respect to the control, but reproducibility issues and complications in reaching desired protein expression targets limited improvement over cycle DBTL1. The recommendations provided by kinetic deep learning using strain LPK15_13342 as base (or control, in black), when implemented in new strains, resulted in TMAF increases of ∼20-40% at t = 72 hrs (see also Extended Data Fig. 9) with respect to the control for DBTL2.1. However, the TMAF for the control strain was only two thirds of that produced in DBTL1, resulting in minor improvements (∼8%) in production in DBTL2.1 with respect to DBTL1. A second attempt at DBTL2 (DBTL2.2) was hence performed, resulting in increases of ∼330-440% at t = 72 hrs (Extended Data Fig. 9) with respect to the control strain, but this control strain TMAF never reached the DBTL1 performance, resulting again in minor improvements over DBTL1 (∼11%). This large variability in results is not entirely surprising since DBTL2 was performed four years after DBTL1 by a different fermentation team (see Methods), and it adds evidence to the burgeoning worry about science reproducibility. Complications in reaching desired protein expression targets compounded this issue (Extended Data Fig. 10). Error bars for DBTL1 were estimated from TMAF results in DBTL2.1 and 2.2, since they were not initially measured due to the very good repeatability obtained by the Ambr 250.

**Extended Data Figure 9:**
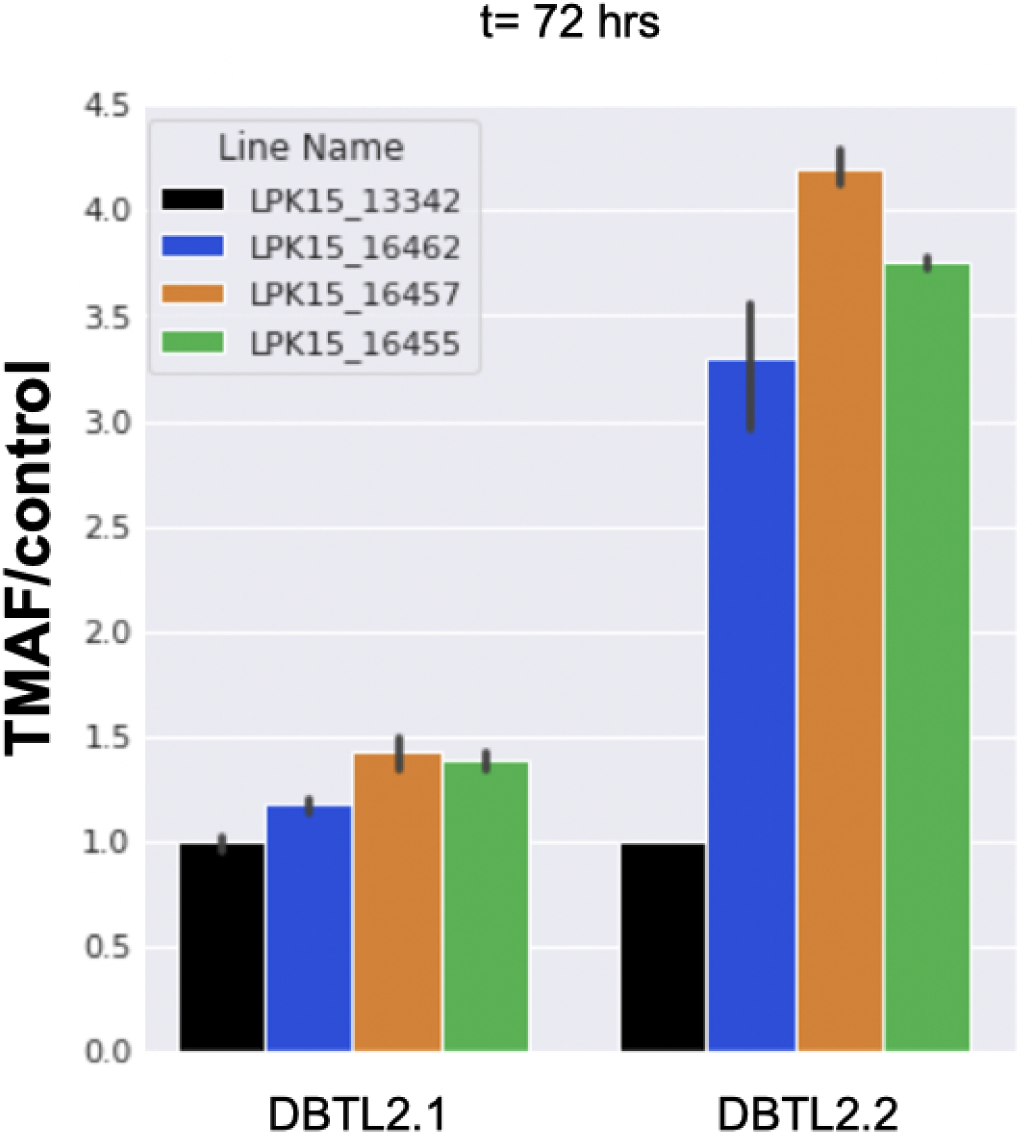
Recommendations in DBTL2 result in a maximum of ∼400% improvement in TMAF over the control strain. However, reproducibility issues in the control result in minimal improvements over DBTL1 (Extended Data Fig. 8).

**Extended Data figure 10:**
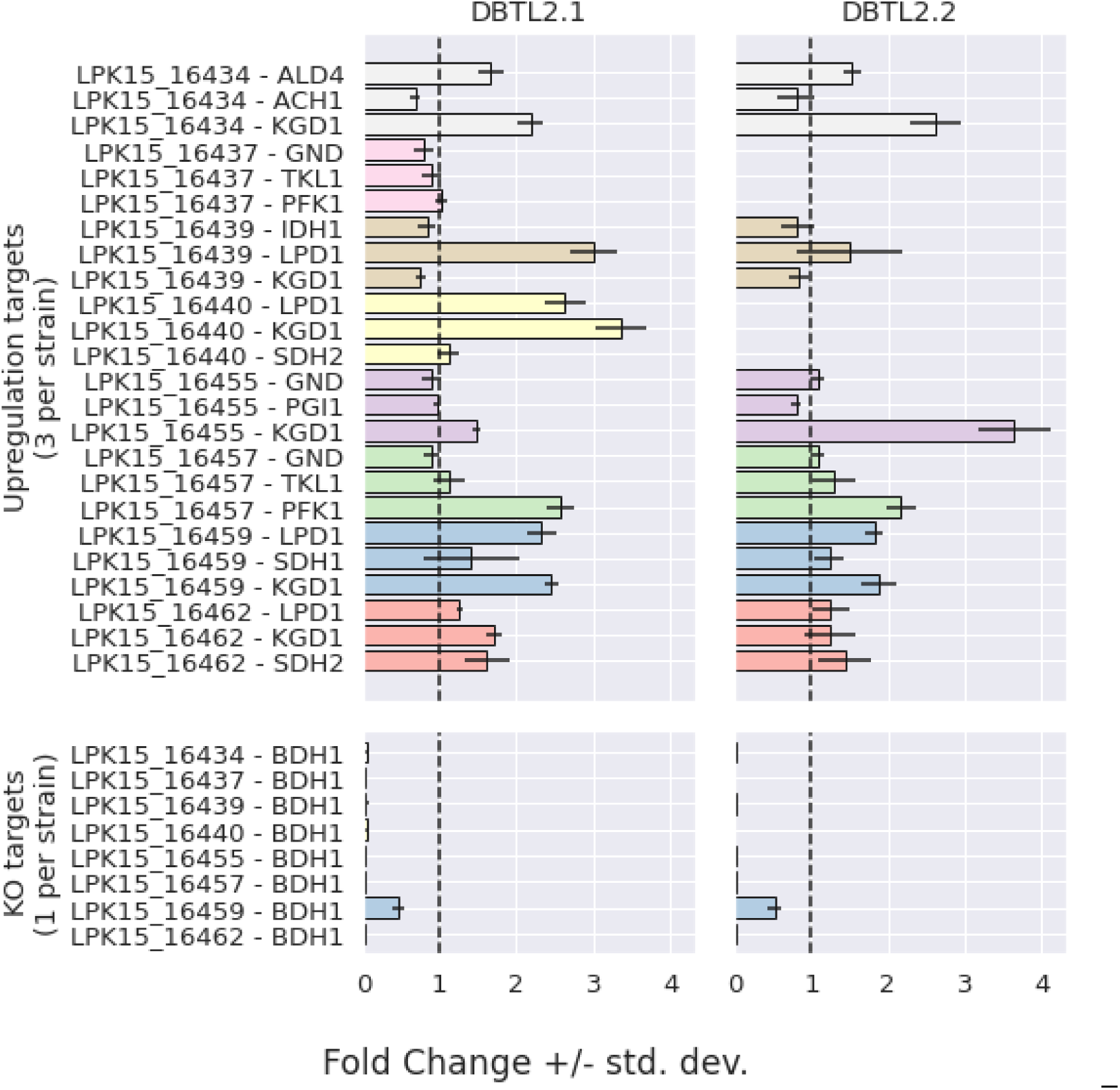
The fold changes of each targeted upregulation or knockout in DBTL2.1 and 2.2 strains compared to their parental strain show that the recommended protein levels were not met, generally speaking. For each strain, we calculated the mean and standard deviation of the strain and parental protein measurements for targeted proteins. We calculated fold change by dividing the strain mean by the parental mean and propagated error to calculate the standard deviation of the fold change. Whereas the BDH1 knockout was achieved on virtually all strains, and several of the desired upregulations were attained, none of the combinations of upregulations were realized at the prescribed levels (2x increase). The BDH1 knockout worked on all strains except one: LPK15_16459. Upregulation was obtained for ALD4, KGD1, LPD1, PFK1, and SDH2. PFK1 upregulation was attained at the prescribed 2x level, and so was LDP1 and KGD1, in most cases. However, GND, SDH2, and IDH1 were never attained at the prescribed level. Neither did all the recommended combinations (Supplementary Table 2). In the highest producing strain (LPK15_16457) only PFK1 was overexpressed, while GND and TKL were not. In the second highest producing strain (LPK15_16455) only KGD1 was overexpressed and only in DBTL2.2 to the levels expected by the recommendations. For the other strain that surpassed the control strain production (LPK15_16462), LPD1, KGD1, SDH2 were overexpressed but none of them at the desired 2x level. This inability to match targeted levels further stresses the importance of effective methods to regulate protein expression accurately, as we have discussed in the past ^52,95^.

