## Supplementary_information for "Prediction of metabolic dynamics through deep learning and high-throughput multiomics data"

### Supplementary Information for:
### Prediction of metabolic dynamics through deep learning and high-throughput multiomics data

Jose Manuel Marti[*,1,2,3], Patrick C. Kinnunen[*,1,2,3], Mark Held[*,4], Azadeh Alikhani[*,4], Andrew Conley[4], Yan Chen[1,2,3], Tijana Radivojevic[1,2,3], Jennifer W. Gin[1,2,3], Apostolos Zournas[1,2], Rebecca M. Lennen[4], Hooman Hefzi[4,γ], Mustafa Baniodeh[5], Peter Harrington[5], Mark Forrer[6], David Melis[4], Jeffrey Dietrich[4], William Holtz[4], Nick Ohler[4], Christopher J. Petzold[1,2,3], Hector Garcia Martin[1,2,3,7,†]

[1] Biological Systems and Engineering Division, Lawrence Berkeley National Laboratory, Berkeley, CA 94720, USA
[2] Department of Energy Agile BioFoundry, Emeryville, CA 94608, USA
[3] Joint BioEnergy Institute, Emeryville, CA 94608, USA
[4] Lygos, Inc, Hayward, CA 94545, USA
[5] NERSC, Lawrence Berkeley National Laboratory, Berkeley, CA 94720, USA
[6] Biomaterials and Biomanufacturing, Sandia National Laboratories, Livermore, California 94550, USA
[7] BCAM, Basque Center for Applied Mathematics, Bilbao 48009, Spain.

[γ] Present address: Department of Biotechnology and Biomedicine, Technical University of Denmark, Lyngby, Denmark

#### Supplementary Note 1: Malonate-producing strains

Malonic acid constitutes a desirable target to contribute to a circular bioeconomy because it is considered one of the top 30 value-added chemicals from biomass [1], and its production via petrochemistry is problematic. Producing malonic acid via petrochemistry requires the use of toxic materials (sodium cyanide, and halogenated precursors such as chloroacetic acid), an undesirable dependence for sustainable chemical production. Malonic acid (often in the form of a dimethyl or diethyl ester) is a versatile "building-block" chemical used for the formation of carbon-carbon-bonds via Michael addition chemistry [2], and is used in wide range of applications that span pharmaceuticals and a range of commercial industrial products [3,4]. For example, malonate is used in some coating applications as a catalytic activity modulator [5] and to control cross-linking [6]. Moreover, dimethyl malonate is a key building block for methyl dihydrojasmonate [7], one of the highest volume fragrance molecules produced [8]. Malonate is also used for synthesis of a range of pharmaceuticals including favipiravir and atorvastatin [9,10].

The industrial biotechnology company Lygos had engineered *Pichia kudriavzevii*, a non-model acid-tolerant yeast, to produce malonic acid in industrially relevant quantities (in the 50-150 g/L range for titer) at demonstration scale [11]. *P. kudriavzevii* is an unconventional non-model yeast that exhibits very different carbon flux profiles from model yeasts, as well as multi-stress tolerant physiology (salt, temperature, and acid), high growth rate, fast glucose consumption rate, and Biosafety Level 1 (BSL-1) status, making it an ideal host for manufacturing chemicals, and in particular organic acids such as malonic acid. That said, as is the case for most renewable biomanufacturing processes [12], improved malonic acid production levels can lower the cost and enhance the economic competitiveness of this sustainable bioproduction route. The twenty three strains used in the first DBTL cycle (Supplementary Table 1) were extremely diverse so as to facilitate learning by the ML algorithms: they comprise everything from near wildtype strains that exhibit no malonic acid production, and where most of the glucose consumed is respired, to Lygos' top performing strain (at the start of this project), where the majority of the carbon was locked in the malonic acid pathway (Fig. 2B).

#### Supplementary Note 2: Possible sources of strain performance variability

The high variability observed in the production performance of the control strain (Extended Data Fig. 8) can be due to several causes arising in the long interval between DBTL1 and DBTL2.1 and DBTL2.2. Perhaps the most likely is genomic instability, in part due to the presence of heterozygous integrations, which has been previously observed for some loci in strain LPK15-11264. Genomic instability can result from many difficult-to-track biological mutational events and can be enhanced in heavily engineered strains that often experience additional selective pressures. Ordinarily the impact of genomic instability is mitigated by always drawing from the same cryogenic stock of the organism, however occasionally working stocks need to be remade, and this occurred between DBTL2.1 and DBTL2.2 using an ancestral permanent cryogenic stock, likely leading to the controls not being genetically identical. These instabilities and operational practicalities highlight the difficulties in ensuring reproducibility for the collection of the large datasets needed for ML in general in synthetic biology. However, time-resolved full genome resequencing of individual colonies from cryogenic stocks was not available as a part of this study to confirm this hypothesis. Another possible source of variability is the lot-to-lot variability of media components over the 4 year span from DBTL1 to DBTL2.1 and DBTL2.2.

We consider this effect to be unlikely to be responsible for the magnitude of observed changes, according to our previous experience with these strains.

The long delay from DBTL1 to DBTL2.1 and DBTL2.2 (~ 4 years) was due to several circumstances outside of our direct control, and underscores the fact that fast DBTL cycles producing large amounts of reproducible multi-omics data are still a challenge in the field. Firstly, a fire in the LBNL lab at Emeryville Station East, where the proteomics and internal metabolomics data were to be measured, led to an initial delay of several months. A second difficulty was the COVID pandemic, which not only stopped work for several months, but had a long-lasting impact due to the shortage of supplies it created for the automated fermentation instrument (Sartorius Ambr250) that was key to this project. An extra impediment came in the form of the lead data scientist moving to a new position, a frequent event in the field due to the scarcity of data science and machine learning talent. Also, this project was finalized with a different Lygos crew than the one that started it. High staff turnover is a common occurrence in the biotech startup space, and this was exacerbated by previous delays. Each of these is a low probability event, but the combination of several tasks with low (but finite) probability of failure results in a complex task with a much increased probability of failure. Perhaps automation in the form of cloud labs or self-driving labs can facilitate this integration and lower the chance of process failure. Furthermore, in our case the significant effort needed to bring the control strain performance to the DBTL1 levels was precluded by Lygos' current business goals. This occurrence highlights the pros (industrially relevant strains, commercially relevant products, generally faster turnarounds) and cons (less ability to follow scientific leads, tighter timelines) of scientific collaboration with industrial partners.

Supplementary Figure 1

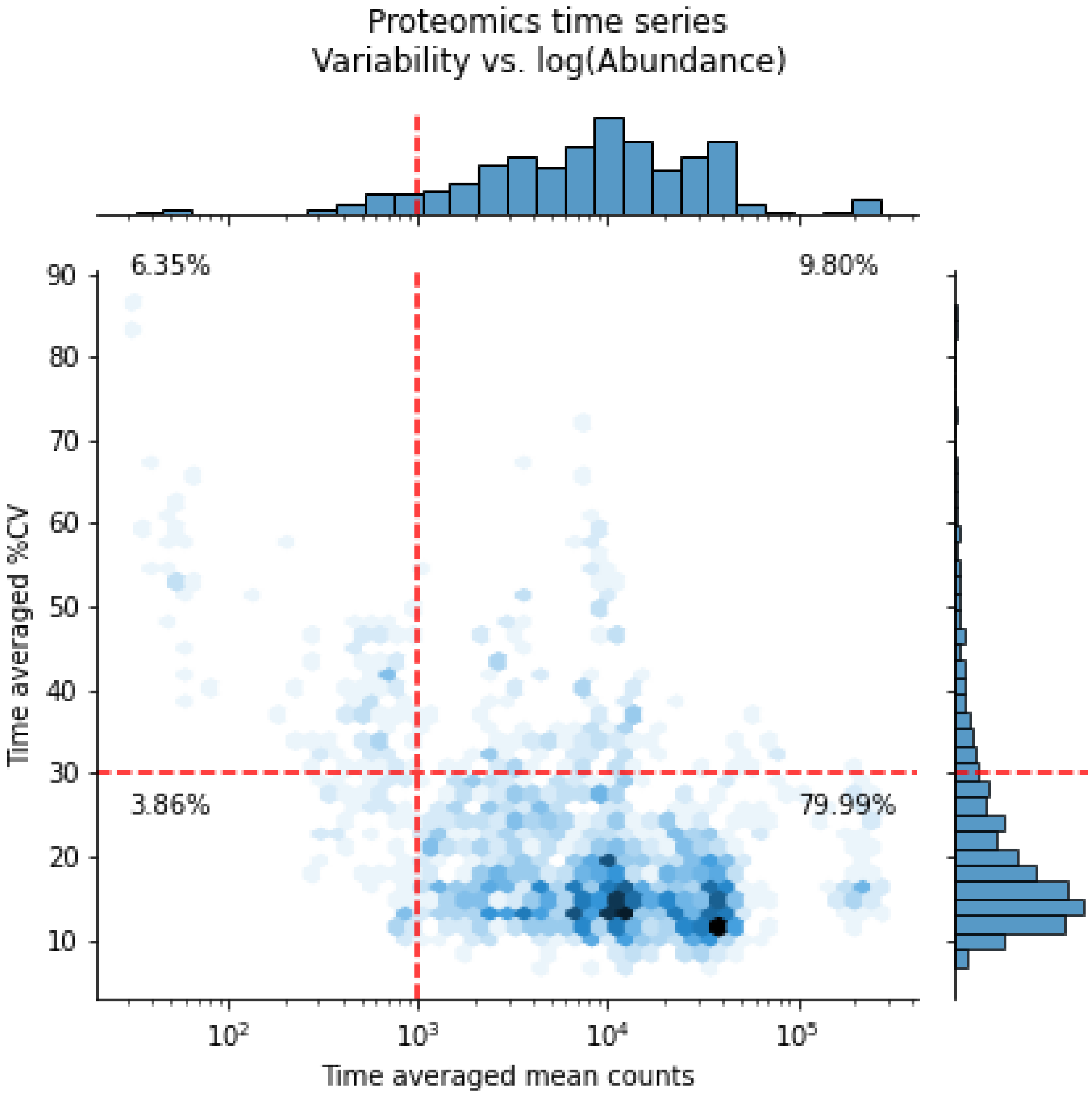


**Supplementary Figure 1: Technical replicates for proteomics data show good repeatability within the same fermentation run.** The coefficient of variation (cv) increases for low counts (protein abundances), but stays below 30% for proteins above a minimal threshold (1000) for a majority of the data (80% of the line/protein combinations).

Supplementary Figure 2

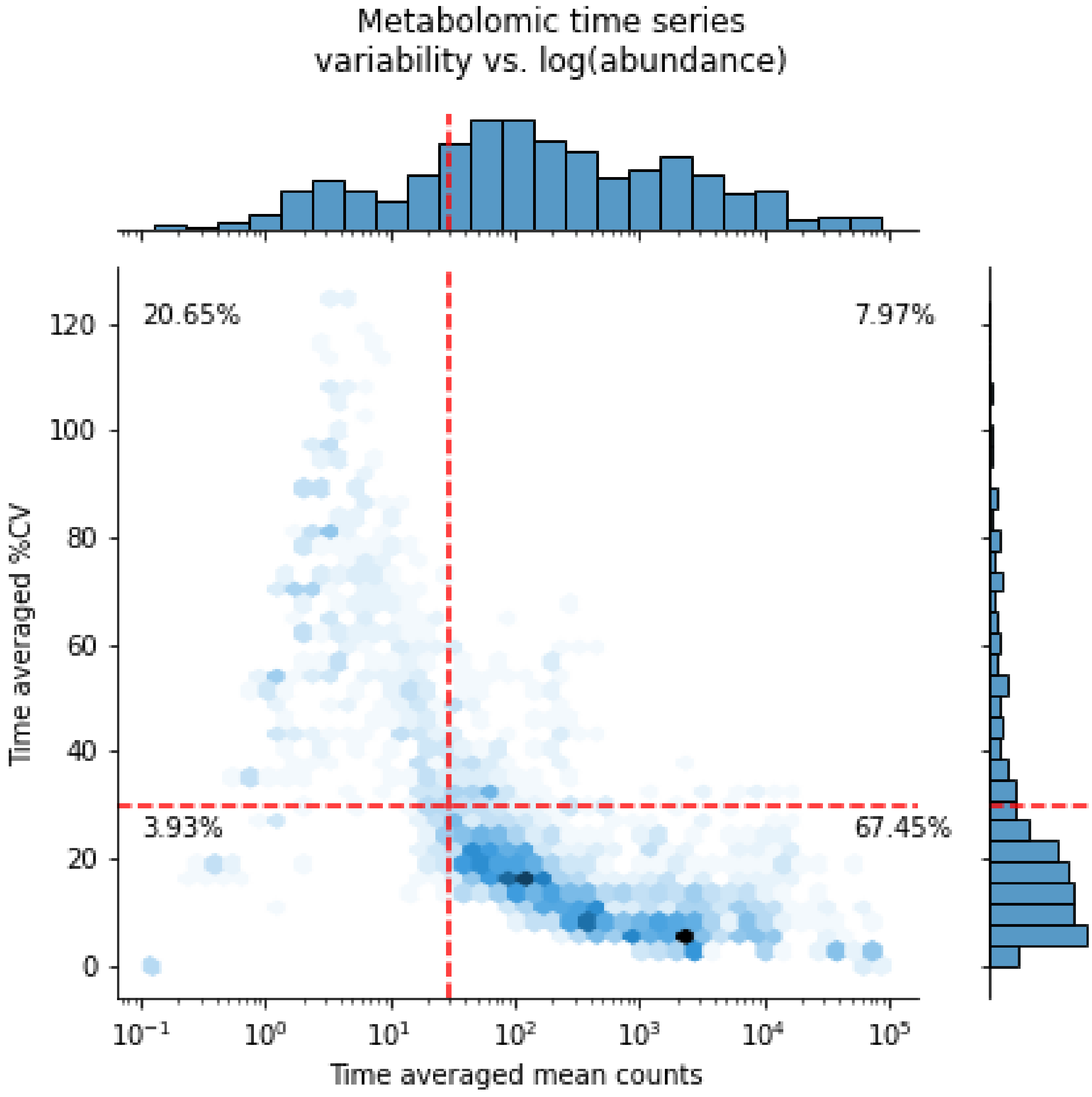


**Supplementary Figure 2: Technical replicates for metabolomics data show good repeatability within the same fermentation run.** The coefficient of variation (cv) increases for low counts (metabolite abundances), but stays below 30% for metabolites above a minimal threshold (30 counts) for a majority of the data (67% of the line/metabolite combinations). This analysis involves the 72 intracellular metabolites. TMAF showed extremely good technical (Extended Data Fig. 8) and biological repeatability (Supplementary Fig. 3) within a fermentation run. For the purpose of the x axis log-transformation, metabolite counts of zero were set to a value of half of the minimum non-zero counts in the dataset (to avoid undefined numbers).

#### Supplementary Figure 3

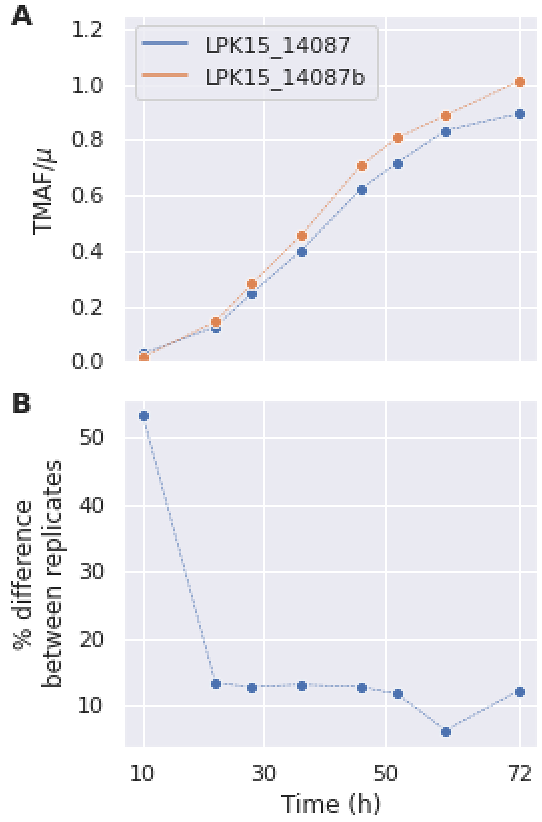


**Supplementary Figure 3: Biological replicates exhibited good repeatability for TMAF measurements within a single fermentation run.** The strain LPK15_14087 was run as a biological duplicate in DBTL1 (LPK15_14087b in Supplementary Table 1). **A)** Measurements of TMAF/$\mu$ for both replicates are very similar. **B)** The percent difference between biological replicates stays ~10%, except for the first timepoint (T = 10 hours) in which TMAF production was very low.

Supplementary Figure 4

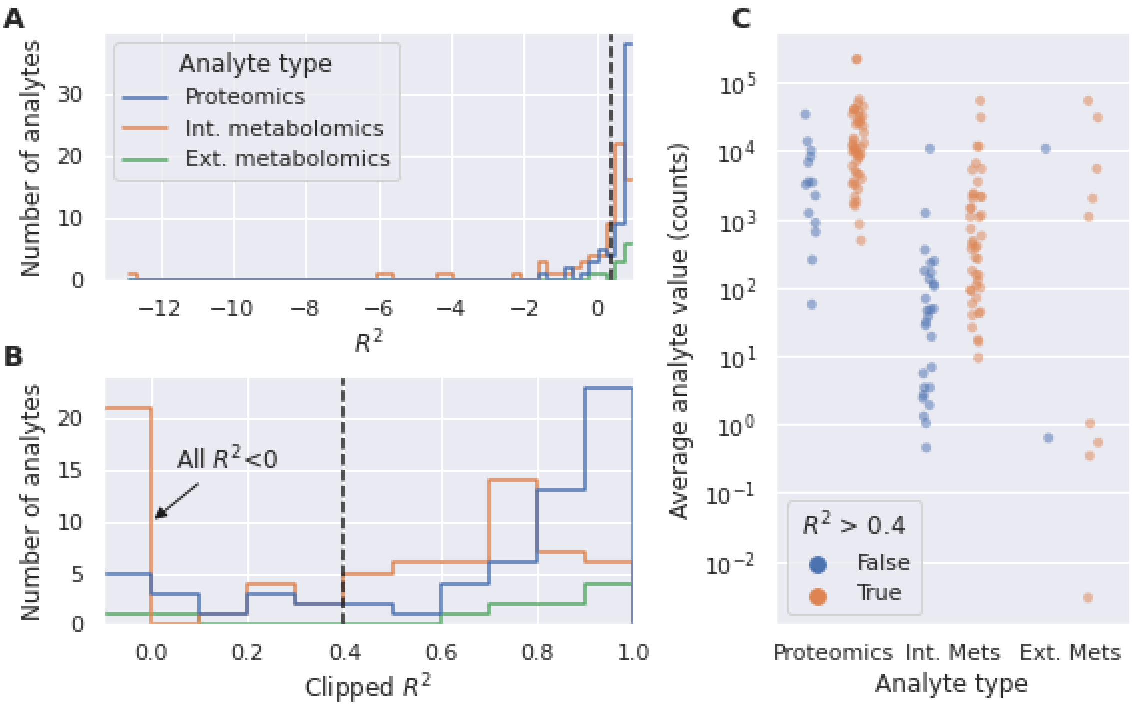


**Supplementary Figure 4: Biological replicates show good repeatability for proteomics and metabolomics data within a single fermentation run.** The distribution of $R^2$ values calculated between LPK15_14087 and LPK15_14087b (biological replicates) for each protein and metabolite show a large proportion of acceptable values ($R^2$>0.4) demonstrating good repeatability for biological replicates, and showing that proteomics and extracellular metabolite data are more repeatable than intracellular metabolomics data. **A)** Full distribution of $R^2$ values. **B)** Same as A), but using a clipped $R^2$, where all values below 0 are considered 0. **C)** Comparing metabolites and proteins with acceptable $R^2$ values ($R^2$ > 0.4) and low $R^2$ values ($R^2 \leq 0.4$) between replicates shows that low $R^2$ metabolites/proteins typically exhibit relatively low abundance.

#### Strain tables

| Strain ID | Description | Primary modifications from base strain | Train/ Test |
|---|---|---|---|
| *LPK15_335* | Base strain | Gen1 strain (post-evolution) | Train |
| LPK15_11959 | | adh3b^::pTDH1>Cas9 | Train |
| LPK15_11965 | | adh6c^::pTDH1>MCH | Train |
| LPK15_11963 | | adh6c^^::pTDH1>MCH | Test |
| LPK15_11977 | | pda1^^:: | Train |
| LPK15_11979 | | gpd1^^:: | Train |
| LPK15_11981 | | adh6b^::pTDH1>MtCAH-l1-PkACC1.m1 | Train |
| LPK15_11988 | | yat1^::pTDH1>PkACC1.m1/pPGK1>ScALD6/pFBA1>YlACS/pFBA1>ScPDC6 | Train |
| LPK15_11989 | | adh1^::pTDH1>PkACC1/pFBA1>DzAADH | Train |
| LPK15_11263 | | adh1^::pTDH1>PkACC1/pFBA1>ScALD6/pPGK1s>YlACS | Test |
| LPK15_11985 | | adh1^^::pTDH1>PkACC1/pFBA1>ScALD6/pPGK1>YlACS | Train |
| LPK15_11991 | | adh6b^::pTDH1>DzAADH | Train |
| LPK15_11260 | | yat1^^::pFBA1>ScPDC6 | Train |
| *LPK15_11264* | | adh3a^::pGPD1>GPD1/pPDA1>PDA1 | Train |
| LPK15_12032 | | adh3a^^::pGPD1>GPD1/pPDA1>PDA1 | Train |
| LPK15_12444 | | ura2^::pGPD1>GPD1/pPDA1>PDA1 | Test |
| *LPK15_13342* | | adh6b^^::pPGK1>Mph-2-C4T | Train |
| LPK15_14056 | | adh6b^^::pPGK1>A0A2J6S698 | Train |
| LPK15_14586 | | ach1^^:: | Train |
| LPK15_14588 | | yat1^^::pTDH1>PkACH1 | Test |
| LPK15_14248 | | Attempted deletion of acc1; sequencing indicates recombination event occurred between native ACC1 promoter and pTDH1 in an introduced extra copy of ACC1 | Train |
| LPK15_14087 | | Cas9^:: | Train |
| LPK15_14087b | Biological replicate of LPK15_ 14087 | Cas9^:: | Train |
| LPK15_14324 | | adh6b^^::pTDH1>MCH | Train |

**Supplementary Table 1: DBTL1 strains**. These strains were used as training data for the machine learning model, which produced the recommended strains in Supplementary Table 2. They include twenty three strains and a biological replicate of LPK15_14087 (LPK15_14087b). Strains in italics are the base strains for the strains shown in

Extended Data Figure 8. A single carat (^) indicates heterozygous integration and a double carat (^^) indicates homozygous integration into the preceding gene locus name. The integration cassette is described after the double colon (::), with “p” indicating a promoter (e.g. pTDH1 is a Pk TDH1 promoter) driving expression of the gene after it (e.g. pTDH1>ScPDC6 is the PDC6 gene from *Saccharomyces cerevisiae* under the Pk TDH1 promoter). Terminators are present in the cassettes but not indicated in the notation in the table. Expression of multiple gene expression units within a cassette are listed with a slash (/) separating them.

| Strain ID | Parental strain | Upregulated genes | Gene KO | ΔTMAF 2.1 46hr | ΔTMAF 2.1 72hr | ΔTMAF 2.2 46hr | ΔTMAF 2.2 72hr |
|---|---|---|---|---|---|---|---|
| *LPK15_11264* | | | | | | | |
| *LPK15_13342* | | | | | | | |
| LPK15_16434 | LPK15_11264 | ALD4, ACH1, KGD1 | BDH1 | 0.103 (29.9%) | 0.297 (53.6%) | 0.02 (6.8%) | 0.147 (23.5%) |
| LPK15_16437 | LPK15_11264 | GND, TKL1, PFK1 | BDH1 | 0.024 (7.0%) | 0.162 (29.2%) | N/A | N/A |
| LPK15_16439 | LPK15_11264 | IDH1, LPD1, KGD1 | BDH1 | 0.163 (47.4%) | 0.262 (47.3%) | -0.039 (-13.8%) | -0.256 (-41.1%) |
| LPK15_16440 | LPK15_11264 | LPD1, KGD1, SDH2 | BDH1 | -0.014 (-4.1%) | -0.035 (-6.4%) | N/A | N/A |
| LPK15_16441 | LPK15_335 | ALD4, ACH1, KGD1 | BDH1 | N/A | N/A | N/A | N/A |
| LPK15_16442 | LPK15_335 | GND, PGI1, KGD1 | BDH1 | N/A | N/A | N/A | N/A |
| LPK15_16443 | LPK15_335 | GND, TKL1, PFK1 | BDH1 | N/A | N/A | N/A | N/A |
| LPK15_16445 | LPK15_335 | LPD1, SDH1, KGD1 | BDH1 | N/A | N/A | N/A | N/A |
| LPK15_16449 | LPK15_11979 | GND, PGI1, KGD1 | BDH1 | N/A | N/A | N/A | N/A |
| LPK15_16452 | LPK15_11979 | LPD1, SDH1, KGD1 | BDH1 | N/A | N/A | N/A | N/A |
| LPK15_16454 | LPK15_11979 | LPD1, KGD1, SDH2 | BDH1 | N/A | N/A | N/A | N/A |
| LPK15_16455 | LPK15_13342 | GND, PGI1, KGD1 | BDH1 | 0.244 (48.1%) | 0.271 (39.5%) | 0.484 (514.1%) | 0.663 (275.4%) |
| LPK15_16457 | LPK15_13342 | GND, TKL1, PFK1 | BDH1 | 0.125 (24.6%) | 0.296 (43.2%) | 0.596 (633.1%) | 0.769 (319.4%) |
| LPK15_16459 | LPK15_13342 | LPD1, SDH1, KGD1 | BDH1 | -0.031 (-6.1%) | -0.012 (-1.8%) | 0.346 (367.5%) | 0.43 (178.6%) |
| LPK15_16462 | LPK15_13342 | LPD1, KGD1, SDH2 | BDH1 | -0.014 (-2.7%) | 0.122 (17.8%) | 0.376 (399.4%) | 0.554 (230.1%) |
| LPK15_16463 | LPK15_14324 | ALD4, ACH1, KGD1 | BDH1 | N/A | N/A | N/A | N/A |
| LPK15_16464 | LPK15_14324 | GND, PGI1, KGD1 | BDH1 | N/A | N/A | N/A | N/A |
| LPK15_16466 | LPK15_14324 | IDH1, LPD1, KGD1 | BDH1 | N/A | N/A | N/A | N/A |
| LPK15_16470 | LPK15_14588 | ALD4, ACH1, KGD1 | BDH1 | N/A | N/A | N/A | N/A |
| LPK15_16473 | LPK15_14588 | IDH1, LPD1, KGD1 | BDH1 | N/A | N/A | N/A | N/A |
| LPK15_16477 | LPK15_14588 | LPD1, KGD1, SDH2 | BDH1 | N/A | N/A | N/A | N/A |
| LPK15_16448 | LPK15_11979 | ALD4, ACH1, KGD1 | BDH1 | N/A | N/A | N/A | N/A |

**Supplementary Table 2: DBTL2 strains**. These strains include two control strains, in italics, plus twenty two recommendations obtained using KinDL predictions, to fulfill the 24 strains tested in each DBTL cycle (each Ambr250 has 24 flasks). Recommendations from KinDL took the form of a parental strain to modify, 3 gene upregulation targets, and one gene knockout target. ΔTMAF values are calculated as the difference in measured TMAF/μ between the means of the strain and its parental strain. The final recommendations involved knocking out butanediol dehydrogenase (BDH1), which represents a competing pathway for intracellular pyruvate) in all cases, and only upregulations. The upregulations targeted enzymes responsible for the glycolytic flux toward pyruvate and acetyl-coA (PGI1, PFK1, LPD1), as well as those influencing the redox balance (GND, TKL1, KGD1, IDH1, SDH1, SDH2). While none of these targets were surprising, the combinations predicted to improve prediction were difficult to guess *a priori.*

#### Measurements tables

| Uniprot ID | Enzyme name | Alias |
|---|---|---|
| A0A099NUP4 | Pyruvate dehydrogenase beta subunit | PDB1 |
| A0A099NWI1 | Succinate dehydrogenase [ubiquinone] cytochrome b small subunit, mitochondrial | SDH4 |
| A0A099NWM4 | 6-phosphogluconate dehydrogenase | GND |
| A0A099NX43 | Phosphoenolpyruvate carboxykinase | PCK1 |
| A0A099NXA0 | NADP+ Isocitrate dehydrogenase, mitochondrial | IDP1 |
| A0A099NXC1 | Adenylate kinase (cytoplasmic) | ADK1 |
| A0A099NXC2 | Aconitate hydratase, mitochondrial | ACO1 |
| A0A099NXI4 | Isocitrate dehydrogenase [NAD] subunit 1, mitochondrial | IDH1 |
| A0A099NY27 | Mitochondrial NADH:ubiquinone oxidoreductase subunit (Complex I) | NUBM |
| A0A099NYL7 | Enolase | ENO1 |
| A0A099NZA9 | Dihydrolipoyl dehydrogenase, mitochondrial | LPD1 |
| A0A099NZE5 | Mitochondrial NADH:ubiquinone oxidoreductase subunit (Complex I) | NUKM |
| A0A099P0J4 | Isocitrate dehydrogenase [NAD] subunit 2, mitochondrial | IDH2 |
| A0A099P156 | Mitochondrial NADH:ubiquinone oxidoreductase subunit (Complex I) | NUAM |
| A0A099P1I6 | Butanediol dehydrogenase | BDH1 |
| A0A099P1N2 | Mitochondrial alcohol dehydrogenase isozyme III | ADH1 |
| A0A099P2U5 | Mitochondrial NADH:ubiquinone oxidoreductase subunit (Complex I) | NUHM |
| A0A099P2X4 | Inorganic pyrophosphatase | IPP1 |
| A0A099P395 | Aldehyde dehydrogenase 5, mitochondrial | ALD4 |
| A0A099P3L2 | Transaldolase | TAL1 |
| A0A099P4E1 | External NADH-ubiquinone oxidoreductase 1, mitochondrial | NDE1 |
| A0A099P5C0 | Acetoacetyl-CoA thiolase | ERG10 |
| A0A099P5F4 | Isocitrate lyase | ICL1 |
| A0A099P5T7 | Isocitrate dehydrogenase [NADP] peroxisomal | IDP2 |
| A0A099P5Y4 | Mitochondrial NADH:ubiquinone oxidoreductase subunit (Complex I) | NUCM |
| A0A099P6G3 | Acetyl-coenzyme A synthetase 1 | ACS1 |
| A0A099P6W9 | Phosphoglycerate kinase | PGK1 |

| | | |
|---|---|---|
| A0A099P7E7 | Pyruvate decarboxylase isozyme 3 (major isoform?) | PDC1 |
| A0A099P8N3 | Succinate dehydrogenase [ubiquinone] flavoprotein subunit, mitochondrial | SDH1 |
| A0A1V2LF84 | NADP-dependent alcohol dehydrogenase 6 | ADH6b |
| A0A1V2LFH0 | Pyruvate kinase | PYK1 |
| A0A1V2LGF2 | Aldolase | FBA1 |
| A0A1V2LH63 | Succinyl-CoA ligase [ADP-forming] subunit beta, mitochondrial | LSC2 |
| A0A1V2LHP2 | glucose-6-phosphate isomerase | PGI1 |
| A0A1V2LHT6 | Trehalose-6-P synthase (synthase subunit) | TPS1 |
| A0A1V2LI03 | Transketolase | TKL1 |
| A0A1V2LI93 | Glyceraldehyde-3-phosphate dehydrogenase | TDH1 |
| A0A1V2LIM1 | Malate dehydrogenase, mitochondrial | MDH1 |
| A0A1V2LLS3 | Triosephosphate Isomerase | TPI2 |
| A0A1V2LM59 | Acetyl-coenzyme A synthetase 2 | ACS2 |
| A0A1V2LME1 | Phosphofructokinase beta | PFK2 |
| A0A1V2LN94 | Fatty acid synthase subunit beta | FAS1 |
| A0A1V2LP13 | Acetyl-CoA hydrolase | ACH1 |
| A0A1V2LP29 | Hexokinase | HXK1 |
| A0A1V2LPA5 | Dihydrolipoyllysine-residue succinyltransferase component of 2-oxoglutarate dehydrogenase complex, mitochondrial | KGD2 |
| A0A1V2LPQ8 | Potassium-activated aldehyde dehydrogenase, mitochondrial | ALD2b |
| A0A1V2LQC3 | Aldehyde dehydrogenase 5, mitochondrial | ALD6 |
| A0A1V2LQM1 | Fatty acid synthase subunit alpha | FAS2 |
| A0A1V2LR63 | Mitochondrial alcohol dehydrogenase isozyme III | ADH3b |
| A0A1V2LRW2 | 2-oxoglutarate dehydrogenase, mitochondrial | KGD1 |
| A0A1V2LRX1 | NAD-dependent malic enzyme, mitochondrial | MAE1 |
| A0A1V2LS36 | Aldehyde dehydrogenase | ALD3 |
| A0A1V2LSB2 | Glycogen synthase | GSY1 |
| A0A1V2LSR5 | Acetyl-CoA carboxylase | ACC1 |
| A0A1V2LT98 | Pyruvate carboxylase | PYC1 |
| A0A1V2LTH8 | Hexokinase 2 | HXK2 |

| | | |
|---|---|---|
| A0A1V2LTK2 | UDP-glucose pyrophosphorylase | UGP1 |
| A0A1V2LU08 | Succinate dehydrogenase [ubiquinone] iron-sulfur subunit, mitochondrial | SDH2 |
| A0A1V2LU20 | Phosphofructokinase alpha | PFK1 |
| B5VTA0 | ScALD6 | ScALD6 |
| F6AA82 | MCH | MCH |
| Q6C2Q5 | YlACS | YlACS |
| Q8NJ70 | Citrate synthase | CIT1 |

**Supplementary Table 3: List of measured proteins**

| Measured intracellular metabolites | | |
|---|---|---|
| Oxaloacetic Acid | D-glucose | Dihydroxyacetone phosphate |
| Oxalate | Fructose 6-phosphate | Cytidine 5’ diphosphate |
| NADP+ | Pyruvate | L-arginine |
| Succinyl-CoA | DL-glyceraldehyde 3-phosphate | FAD |
| Malonate | Trehalose 6-phosphate | NADH |
| L-tyrosine | glyoxylate | Biotin |
| L-glutamic acid | Malic acid | D-glucose 6-phosphate |
| Methylmalonic acid | Ribose 5-phosphate | Uridine 5’-diphosphate |
| Coenzyme A | Methylmalonyl-CoA | Deoxythymidine diphosphate |
| Trehalose | Succinate | 6-phosphogluconic acid |
| Cytidine Triphosphate | NADPH | 5’-cytidilic acid |
| Cis-aconitic acid | L-leucine | Guanosine triphosphate |
| L-methionine | 3-phosphoglycerate | D-arabinitol |
| Fumarate | acetylphosphate | Adenosine 5’-diphosphate |
| Lactic acid | cis-4-coumarate | D-Erythrose 4-phosphate |
| Sedoheptulose 7-phosphate | stearoyl-CoA | Propionyl-CoA |
| Oxidized glutathione | phosphoenolpyruvate | Deoxythymidine triphosphate |
| Isopentenyl pyrophosphate | β-D-fructose 1,6-bisphosphate | L-phenylalanine |
| R-mevalonate | L-aspartic acid | ATP |
| Thymidylic acid | Guanosine 5’-diphosphate | L-serine |
| Acetyl-CoA | L-histidine | Glutathione |
| Uridine 5’-triphosphate | Adenosine 5’ monophosphate | Nadide |
| 5’-guanylic acid | palmitoyl-CoA | |
| L-threonine | 2-ketoglutaric acid | |
| Uridine 5’-monophosphate | Malonyl CoA | |

**Supplementary Table 4: List of measured intracellular metabolites**.

| **Measured extracellular metabolites** |
|---|
| Pyruvate |
| Malonate |
| Ethanol |
| Citrate |
| Trehalose |
| Acetate |
| D-arabinitol |
| Glycerol |
| Uracil |
| Succinate |
| D-glucose |

**Supplementary Table 5: List of measured extracellular metabolites**.

| |
|---|
| Dry Cell Weight |
| Dissolved O2 |
| pH |
| CO2 |
| TMAF (Total Malonic Acid Formed) |

**Supplementary Table 6: List of measured bioreactor-related data.**

#### Neural network architecture tables

| Layer | Input Size | Output Size | Number of Nodes | Activation Function |
|---|---|---|---|---|
| Input | (n_batch, n_times, n_features) | (n_batch, n_times, n_features) | N/A | N/A |
| Flatten | (n_batch, n_times, n_features) | (n_batch, n_times * n_features) | N/A | N/A |
| Dense-1 | (n_batch, n_times * n_features) | (n_batch, 252) | 252 | ReLU |
| Dense-2 | (n_batch, 252) | (n_batch, 252) | 252 | ReLU |
| Dense-3 | (n_batch, 252) | (n_batch, 252) | 252 | ReLU |
| Dense-4 | (n_batch, 252) | (n_batch, 252) | 252 | ReLU |
| Dense-TMAF | (n_batch, 252) | (n_batch, n_times) | n_times | None |
| Dense-CO2 | (n_batch, 252) | (n_batch, n_times) | n_times | None |

**Supplementary Table 7:** Fully connected neural network (FCNN) description. Initial versions of the neural networks included $CO_2$ as a response to be predicted, but these predictions were dropped in all simulations because they were not used.

| Layer | Input Size | Output size | Number of Nodes | Activation Function |
|---|---|---|---|---|
| Input | (n_batch, n_times, n_features) | (n_batch, n_times, n_features) | N/A | N/A |
| RNN-1 | (n_batch, n_times, n_features) | (n_batch, n_times, 130) | 130 | Tanh |
| RNN-2 | (n_batch, n_times, 130) | (n_batch, n_times, 130) | 130 | Tanh |
| RNN-3 | (n_batch, n_times, 130) | (n_batch, n_times, 130) | 130 | Tanh |
| RNN–TMAF | (n_batch, n_times, 27) | (n_batch, n_times, 27) | 27 | Tanh |
| RNN–CO2 | (n_batch, n_times, 27) | (n_batch, n_times, 27) | 27 | Tanh |
| Time distributed Dense-TMAF | (n_batch, n_times) | (n_batch, n_times, 1) | 1 | None |
| Time distributed Dense-CO2 | (n_batch, n_times) | (n_batch, n_times, 1) | 1 | None |

**Supplementary Table 8:** Deep recurrent neural network (DRNN) description.

| Layer | Input Size | Output size | Number of Nodes | Activation Function |
|---|---|---|---|---|
| Input | (n_batch, n_times, n_features) | (n_batch, n_times, n_features) | N/A | N/A |
| GRU-1 | (n_batch, n_times, n_features) | (n_batch, n_times, 97) | 97 | Tanh |
| GRU-2 | (n_batch, n_times, 97) | (n_batch, n_times, 97) | 97 | Tanh |
| GRU-3 | (n_batch, n_times, 97) | (n_batch, n_times, 97) | 97 | Tanh |
| GRU–TMAF | (n_batch, n_times) | (n_batch, n_times, 46) | 46 | Tanh |
| GRU–CO2 | (n_batch, 252) | (n_batch, n_times, 46) | 46 | Tanh |
| Time distributed Dense-TMAF | (n_batch, 252) | (n_batch, n_times, 1) | 1 | None |
| Time distributed Dense-CO2 | (n_batch, 252) | (n_batch, n_times, 1) | 1 | None |

**Supplementary Table 9:** Recurrent neural network with GRU neurons (GRU) description.

| Model | Parameter | Value |
|---|---|---|
| Model 1 - ANN | Loss Function | Mean Squared Error |
| | Learning Rate | 0.001 |
| | Rho | 0.95 |
| | optimizer | Adadelta |
| | epochs | 100 |
| | Steps per epoch | 50 |
| Model 2 - Deep RNN | Loss Function | Mean Squared Error |
| | Learning Rate | 1.0 |
| | optimizer | Adam (default) |
| | epochs | 100 |
| | Steps per epoch | 50 |
| Model 3 - GRU | Loss function | Mean Squared Error |
| | Learning rate | 1.0 |
| | optimizer | Adam (default) |
| | epochs | 100 |
| | Steps per epoch | 50 |

**Supplementary Table 10:** Training hyperparameters for each model.

#### Predictions table

| Intracellular Metabolites showing $R^2 > 0.60$ | |
|---|---|
| Metabolite | $R^2$ |
| D-glucose | 0.836 |
| cis-Aconitic acid | 0.796 |
| Glutathione | 0.794 |
| 2-ketoglutaric acid | 0.759 |
| Adenosine 5' diphosphate | 0.758 |
| 5'-Guanylic acid | 0.745 |
| Trehalose | 0.741 |
| Coenzyme A | 0.734 |
| 3-phosphoglycerate | 0.730 |
| L-threonine | 0.728 |
| 5'-Cytidilic acid | 0.690 |
| NADP+ | 0.682 |
| malonate | 0.681 |
| D-Arabinitol | 0.672 |
| (R)-mevalonate | 0.654 |
| cis-4-coumarate | 0.650 |
| isopentenyl pyrophosphate | 0.650 |
| biotin | 0.644 |
| Glutathione oxidized form | 0.642 |
| Uridine 5'-diphosphate | 0.616 |

**Supplementary Table 11: Well-predicted intracellular metabolites.** Twenty intracellular metabolites displayed an $R^2>0.6$ (well predicted).

#### Predictions for all metabolites and lines.

The pages after the bibliography compare all intracellular and extracellular metabolite predictions and measured data for all strains. In each page, a single metabolite is presented, as indicated at the top of the page. Training strains are shown, and then test strains. Under each strain heading, the replicates are shown in order (R1, R2, and R3).

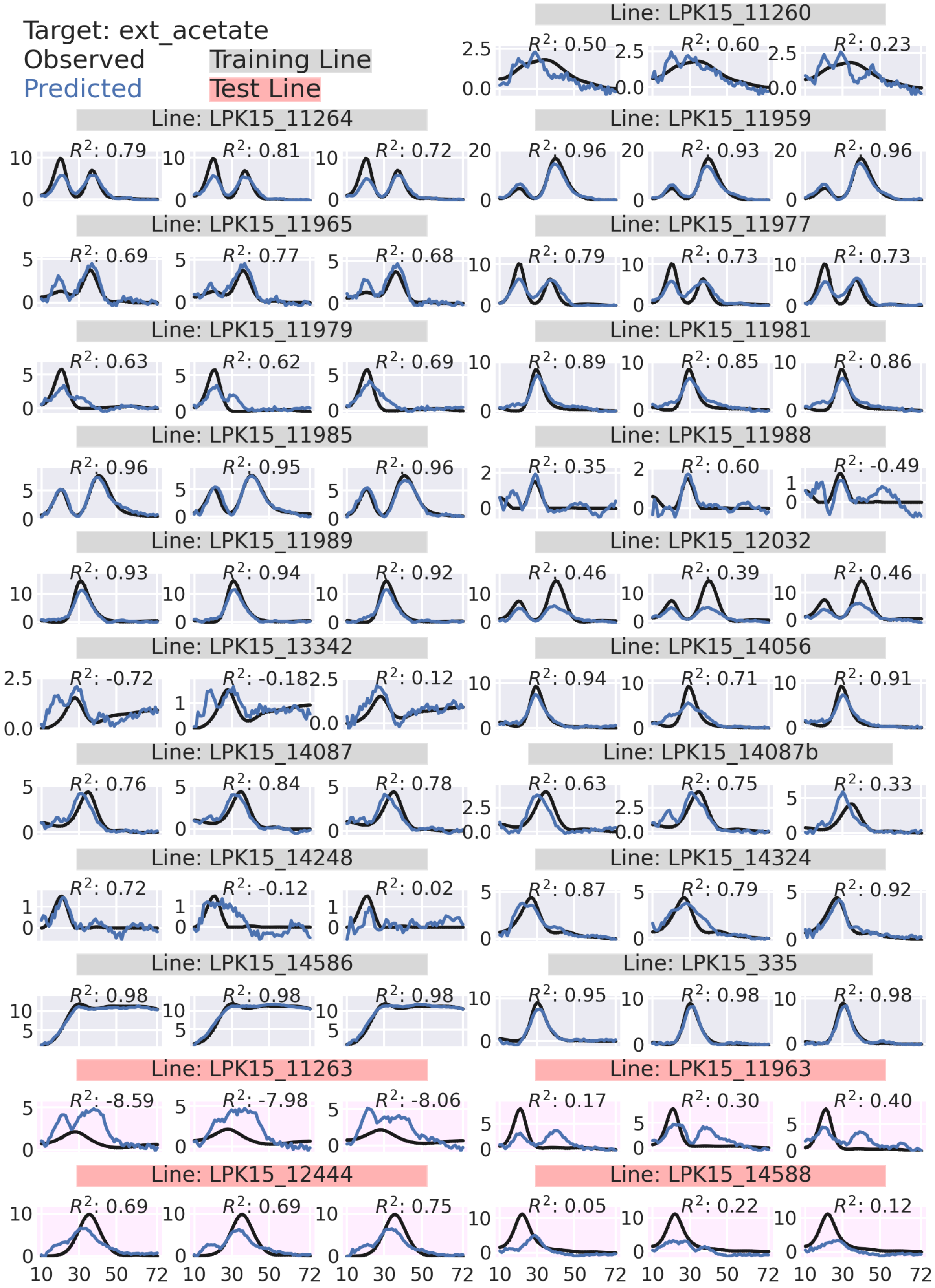

Target: ext_acetate
Observed
Predicted
Training Line
Test Line
Line: LPK15_11260
$R^2$: 0.50
$R^2$: 0.60
$R^2$: 0.23
Line: LPK15_11264
$R^2$: 0.79
$R^2$: 0.81
$R^2$: 0.72
Line: LPK15_11959
$R^2$: 0.96
$R^2$: 0.93
$R^2$: 0.96
Line: LPK15_11965
$R^2$: 0.69
$R^2$: 0.77
$R^2$: 0.68
Line: LPK15_11977
$R^2$: 0.79
$R^2$: 0.73
$R^2$: 0.73
Line: LPK15_11979
$R^2$: 0.63
$R^2$: 0.62
$R^2$: 0.69
Line: LPK15_11981
$R^2$: 0.89
$R^2$: 0.85
$R^2$: 0.86
Line: LPK15_11985
$R^2$: 0.96
$R^2$: 0.95
$R^2$: 0.96
Line: LPK15_11988
$R^2$: 0.35
$R^2$: 0.60
$R^2$: -0.49
Line: LPK15_11989
$R^2$: 0.93
$R^2$: 0.94
$R^2$: 0.92
Line: LPK15_12032
$R^2$: 0.46
$R^2$: 0.39
$R^2$: 0.46
Line: LPK15_13342
$R^2$: -0.72
$R^2$: -0.18
$R^2$: 0.12
Line: LPK15_14056
$R^2$: 0.94
$R^2$: 0.71
$R^2$: 0.91
Line: LPK15_14087
$R^2$: 0.76
$R^2$: 0.84
$R^2$: 0.78
Line: LPK15_14087b
$R^2$: 0.63
$R^2$: 0.75
$R^2$: 0.33
Line: LPK15_14248
$R^2$: 0.72
$R^2$: -0.12
$R^2$: 0.02
Line: LPK15_14324
$R^2$: 0.87
$R^2$: 0.79
$R^2$: 0.92
Line: LPK15_14586
$R^2$: 0.98
$R^2$: 0.98
$R^2$: 0.98
Line: LPK15_335
$R^2$: 0.95
$R^2$: 0.98
$R^2$: 0.98
Line: LPK15_11263
$R^2$: -8.59
$R^2$: -7.98
$R^2$: -8.06
Line: LPK15_11963
$R^2$: 0.17
$R^2$: 0.30
$R^2$: 0.40
Line: LPK15_12444
$R^2$: 0.69
$R^2$: 0.69
$R^2$: 0.75
Line: LPK15_14588
$R^2$: 0.05
$R^2$: 0.22
$R^2$: 0.12
10 30 50 72

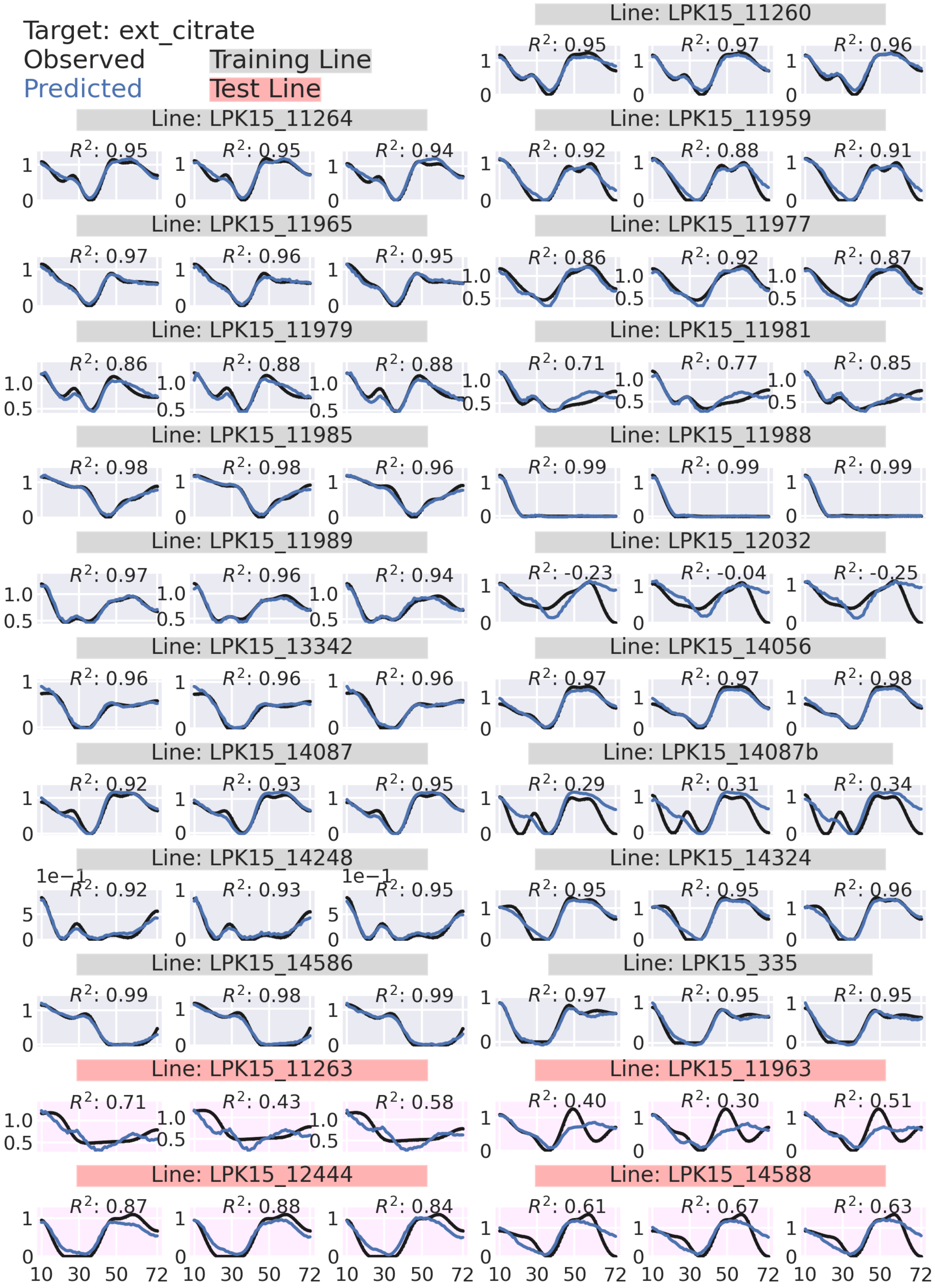

Target: ext_citrate
Observed
Predicted
Training Line
Test Line
Line: LPK15_11260
Line: LPK15_11264
Line: LPK15_11959
Line: LPK15_11965
Line: LPK15_11977
Line: LPK15_11979
Line: LPK15_11981
Line: LPK15_11985
Line: LPK15_11988
Line: LPK15_11989
Line: LPK15_12032
Line: LPK15_13342
Line: LPK15_14056
Line: LPK15_14087
Line: LPK15_14087b
Line: LPK15_14248
Line: LPK15_14324
Line: LPK15_14586
Line: LPK15_335
Line: LPK15_11263
Line: LPK15_11963
Line: LPK15_12444
Line: LPK15_14588

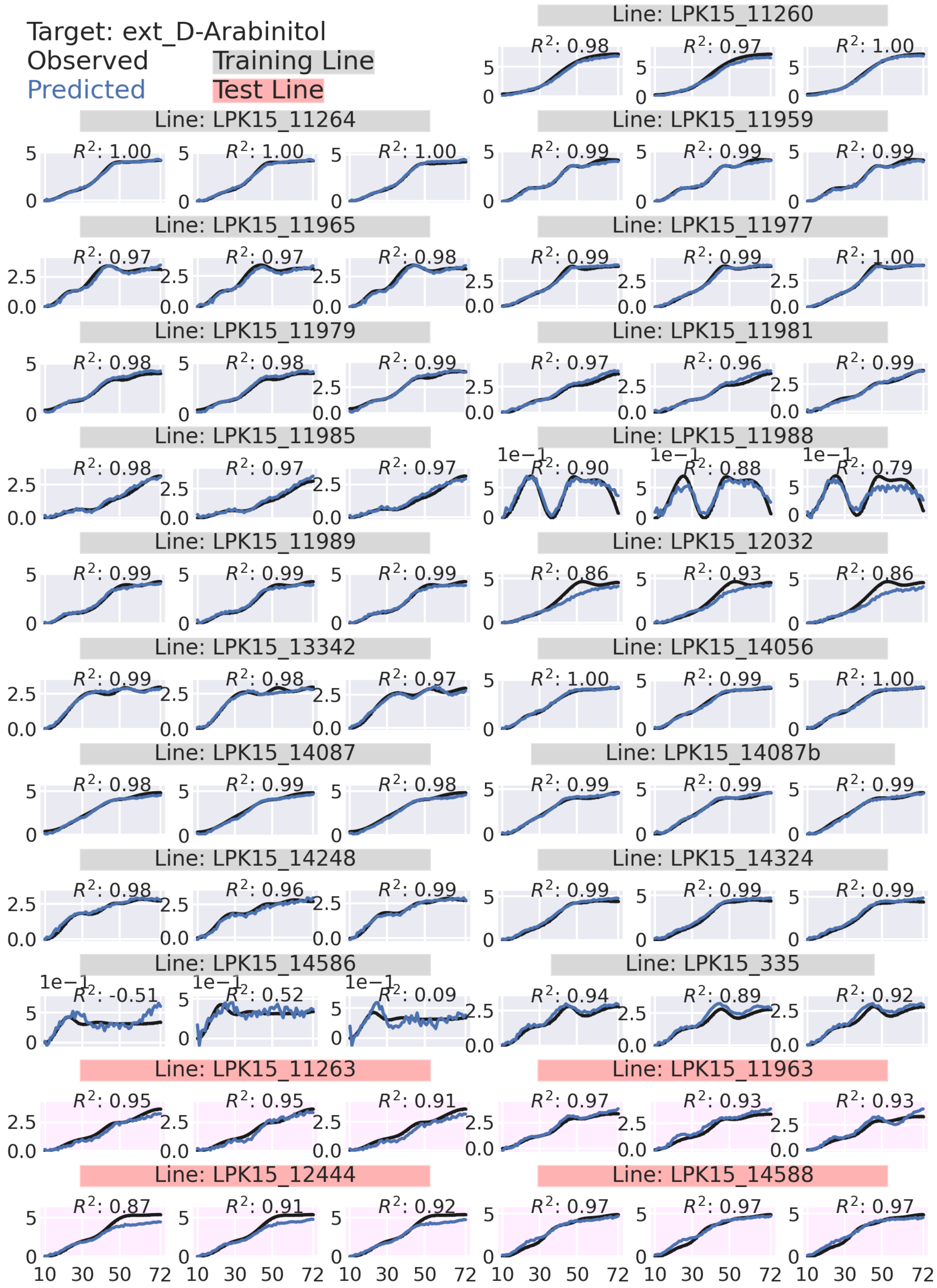
Target: ext_D-Arabinitol
Observed
Predicted
Training Line
Test Line
Line: LPK15_11260
$R^2$: 0.98
$R^2$: 0.97
$R^2$: 1.00
Line: LPK15_11264
$R^2$: 1.00
$R^2$: 1.00
$R^2$: 1.00
Line: LPK15_11959
$R^2$: 0.99
$R^2$: 0.99
$R^2$: 0.99
Line: LPK15_11965
$R^2$: 0.97
$R^2$: 0.97
$R^2$: 0.98
Line: LPK15_11977
$R^2$: 0.99
$R^2$: 0.99
$R^2$: 1.00
Line: LPK15_11979
$R^2$: 0.98
$R^2$: 0.98
$R^2$: 0.99
Line: LPK15_11981
$R^2$: 0.97
$R^2$: 0.96
$R^2$: 0.99
Line: LPK15_11985
$R^2$: 0.98
$R^2$: 0.97
$R^2$: 0.97
Line: LPK15_11988
$R^2$: 0.90
$R^2$: 0.88
$R^2$: 0.79
Line: LPK15_11989
$R^2$: 0.99
$R^2$: 0.99
$R^2$: 0.99
Line: LPK15_12032
$R^2$: 0.86
$R^2$: 0.93
$R^2$: 0.86
Line: LPK15_13342
$R^2$: 0.99
$R^2$: 0.98
$R^2$: 0.97
Line: LPK15_14056
$R^2$: 1.00
$R^2$: 0.99
$R^2$: 1.00
Line: LPK15_14087
$R^2$: 0.98
$R^2$: 0.99
$R^2$: 0.98
Line: LPK15_14087b
$R^2$: 0.99
$R^2$: 0.99
$R^2$: 0.99
Line: LPK15_14248
$R^2$: 0.98
$R^2$: 0.96
$R^2$: 0.99
Line: LPK15_14324
$R^2$: 0.99
$R^2$: 0.99
$R^2$: 0.99
Line: LPK15_14586
$R^2$: -0.51
$R^2$: 0.52
$R^2$: 0.09
Line: LPK15_335
$R^2$: 0.94
$R^2$: 0.89
$R^2$: 0.92
Line: LPK15_11263
$R^2$: 0.95
$R^2$: 0.95
$R^2$: 0.91
Line: LPK15_11963
$R^2$: 0.97
$R^2$: 0.93
$R^2$: 0.93
Line: LPK15_12444
$R^2$: 0.87
$R^2$: 0.91
$R^2$: 0.92
Line: LPK15_14588
$R^2$: 0.97
$R^2$: 0.97
$R^2$: 0.97
1e−1
10 30 50 72

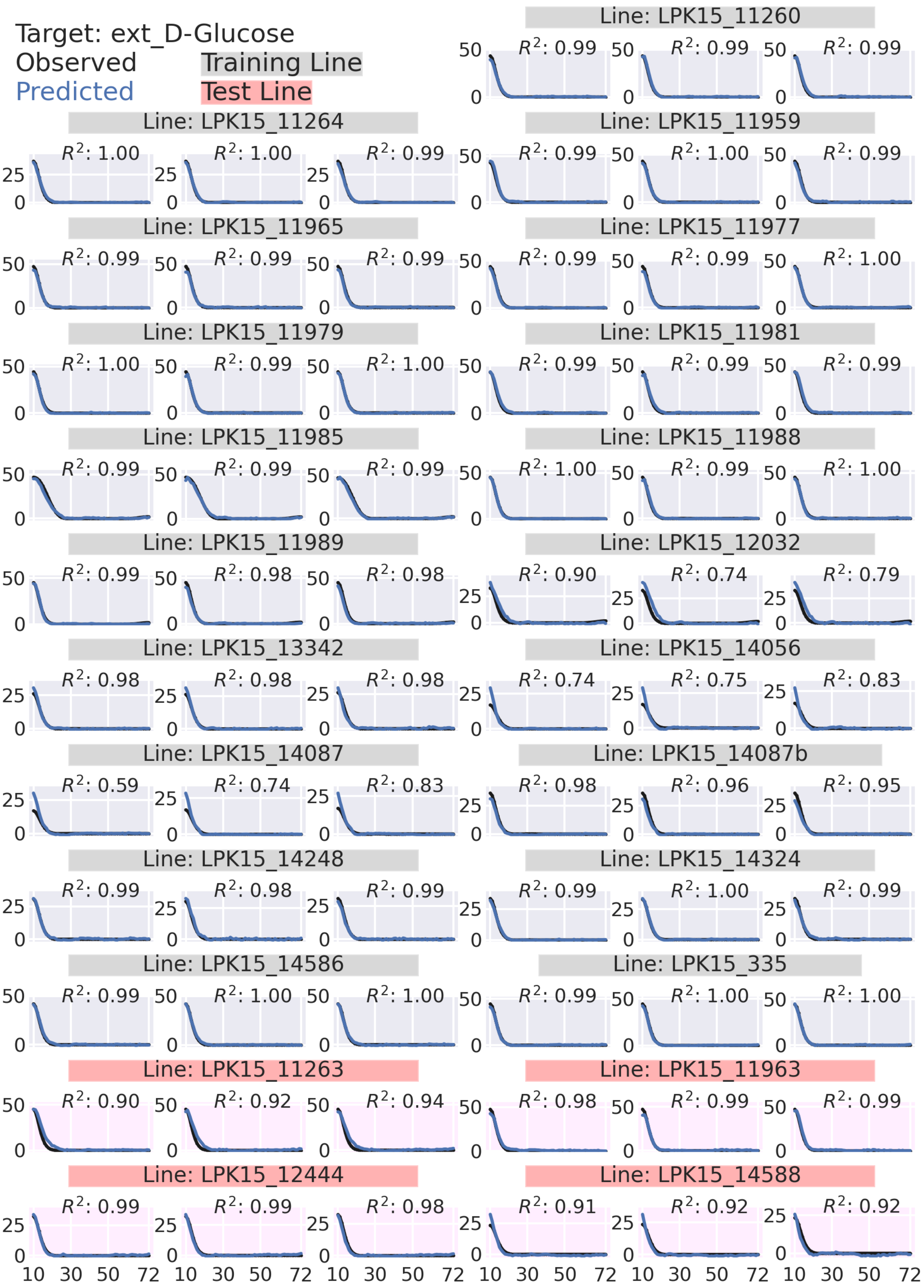
Target: ext_D-Glucose
Observed
Predicted
Training Line
Test Line
Line: LPK15_11260
Line: LPK15_11264
Line: LPK15_11959
Line: LPK15_11965
Line: LPK15_11977
Line: LPK15_11979
Line: LPK15_11981
Line: LPK15_11985
Line: LPK15_11988
Line: LPK15_11989
Line: LPK15_12032
Line: LPK15_13342
Line: LPK15_14056
Line: LPK15_14087
Line: LPK15_14087b
Line: LPK15_14248
Line: LPK15_14324
Line: LPK15_14586
Line: LPK15_335
Line: LPK15_11263
Line: LPK15_11963
Line: LPK15_12444
Line: LPK15_14588

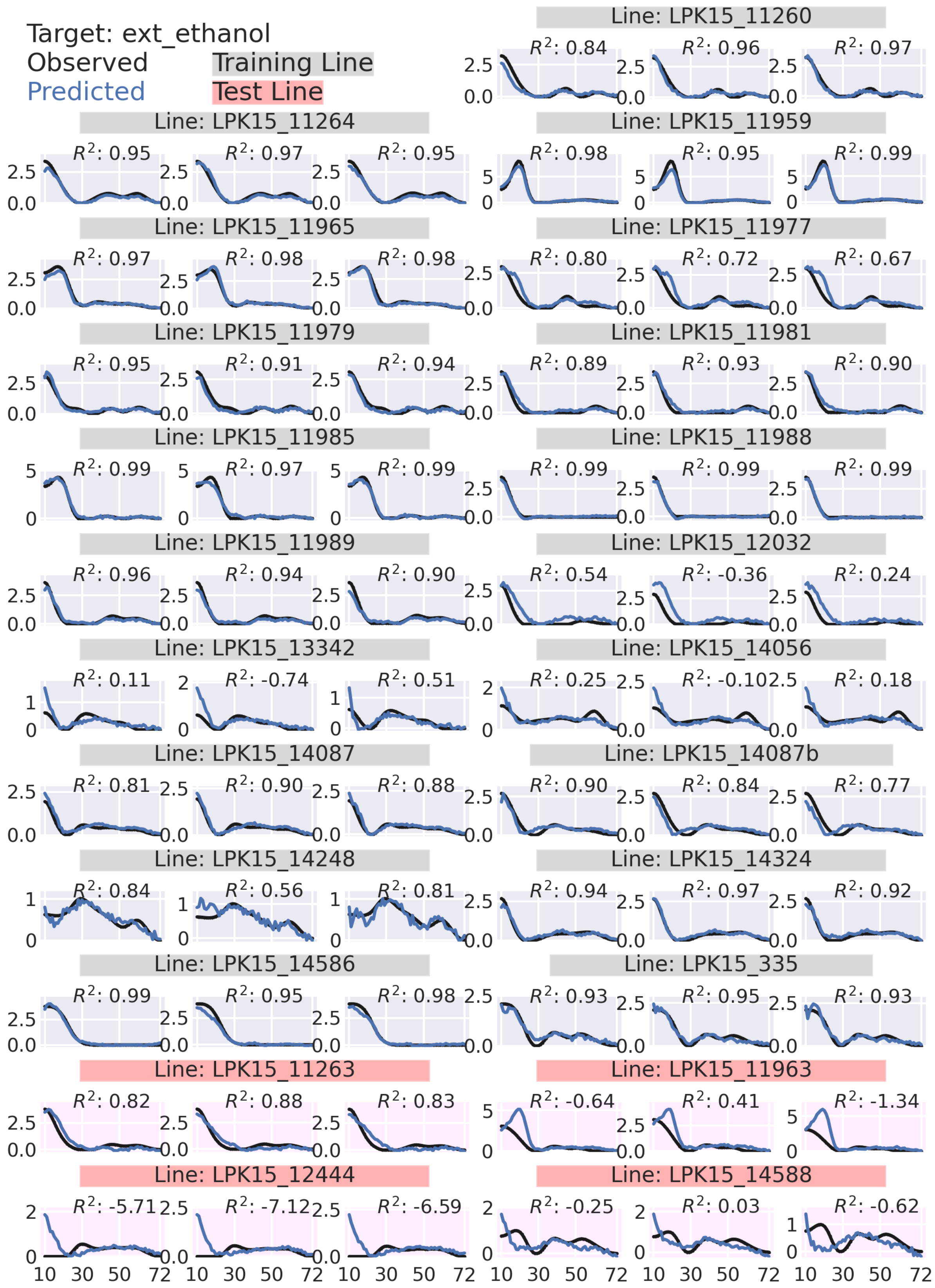
Target: ext_ethanol
Observed
Predicted
Training Line
Test Line
Line: LPK15_11260
Line: LPK15_11264
Line: LPK15_11959
Line: LPK15_11965
Line: LPK15_11977
Line: LPK15_11979
Line: LPK15_11981
Line: LPK15_11985
Line: LPK15_11988
Line: LPK15_11989
Line: LPK15_12032
Line: LPK15_13342
Line: LPK15_14056
Line: LPK15_14087
Line: LPK15_14087b
Line: LPK15_14248
Line: LPK15_14324
Line: LPK15_14586
Line: LPK15_335
Line: LPK15_11263
Line: LPK15_11963
Line: LPK15_12444
Line: LPK15_14588

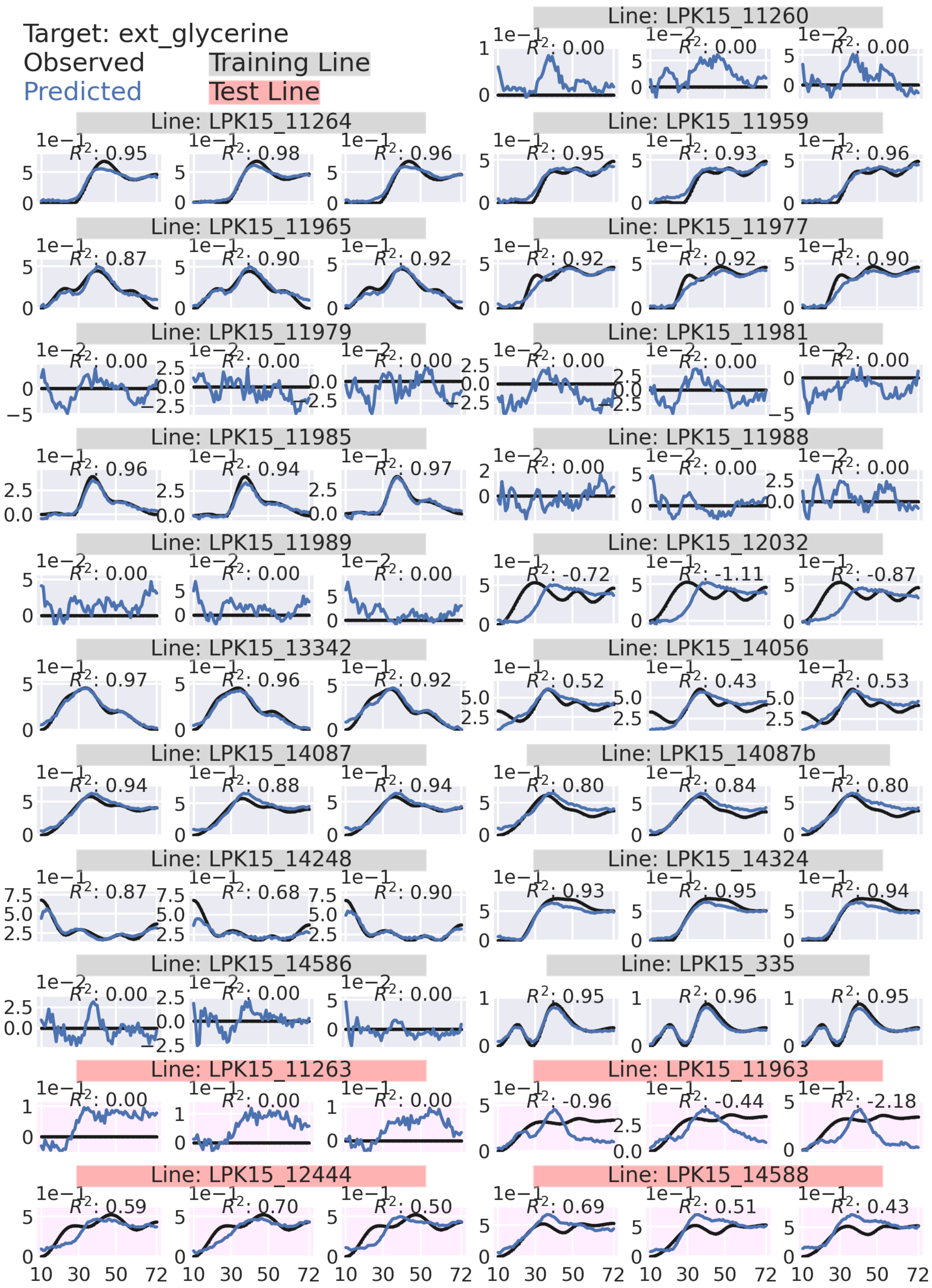

Target: ext_glycerine
Observed
Predicted
Training Line
Test Line
Line: LPK15_11260
Line: LPK15_11264
Line: LPK15_11959
Line: LPK15_11965
Line: LPK15_11977
Line: LPK15_11979
Line: LPK15_11981
Line: LPK15_11985
Line: LPK15_11988
Line: LPK15_11989
Line: LPK15_12032
Line: LPK15_13342
Line: LPK15_14056
Line: LPK15_14087
Line: LPK15_14087b
Line: LPK15_14248
Line: LPK15_14324
Line: LPK15_14586
Line: LPK15_335
Line: LPK15_11263
Line: LPK15_11963
Line: LPK15_12444
Line: LPK15_14588

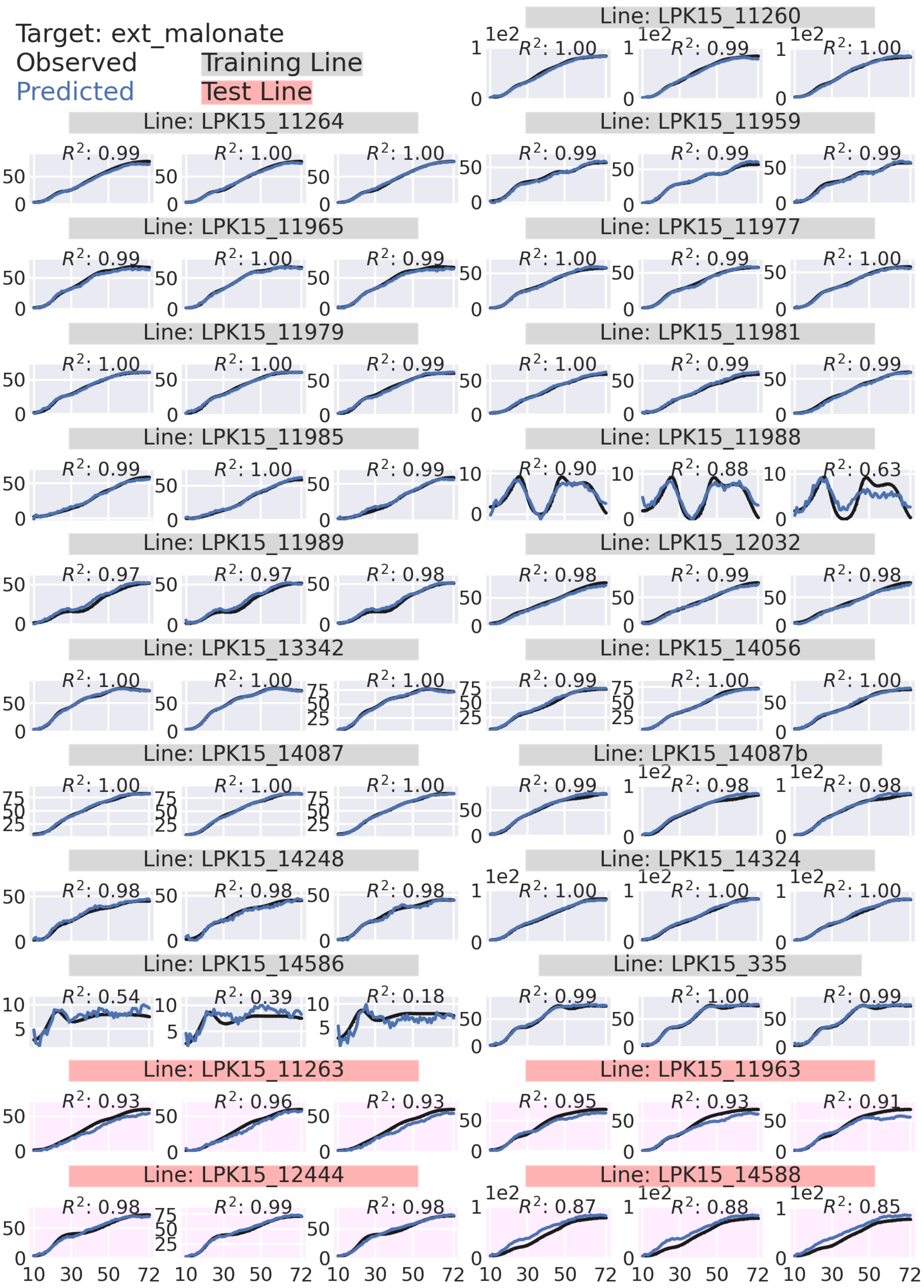

Target: ext_malonate
Observed
Predicted
Training Line
Test Line
Line: LPK15_11260
$R^2$: 1.00
$R^2$: 0.99
$R^2$: 1.00
Line: LPK15_11264
$R^2$: 0.99
$R^2$: 1.00
$R^2$: 1.00
Line: LPK15_11959
$R^2$: 0.99
$R^2$: 0.99
$R^2$: 0.99
Line: LPK15_11965
$R^2$: 0.99
$R^2$: 1.00
$R^2$: 0.99
Line: LPK15_11977
$R^2$: 1.00
$R^2$: 0.99
$R^2$: 1.00
Line: LPK15_11979
$R^2$: 1.00
$R^2$: 1.00
$R^2$: 0.99
Line: LPK15_11981
$R^2$: 1.00
$R^2$: 0.99
$R^2$: 0.99
Line: LPK15_11985
$R^2$: 0.99
$R^2$: 1.00
$R^2$: 0.99
Line: LPK15_11988
$R^2$: 0.90
$R^2$: 0.88
$R^2$: 0.63
Line: LPK15_11989
$R^2$: 0.97
$R^2$: 0.97
$R^2$: 0.98
Line: LPK15_12032
$R^2$: 0.98
$R^2$: 0.99
$R^2$: 0.98
Line: LPK15_13342
$R^2$: 1.00
$R^2$: 1.00
$R^2$: 1.00
Line: LPK15_14056
$R^2$: 0.99
$R^2$: 1.00
$R^2$: 1.00
Line: LPK15_14087
$R^2$: 1.00
$R^2$: 1.00
$R^2$: 1.00
Line: LPK15_14087b
$R^2$: 0.99
$R^2$: 0.98
$R^2$: 0.98
Line: LPK15_14248
$R^2$: 0.98
$R^2$: 0.98
$R^2$: 0.98
Line: LPK15_14324
$R^2$: 1.00
$R^2$: 1.00
$R^2$: 1.00
Line: LPK15_14586
$R^2$: 0.54
$R^2$: 0.39
$R^2$: 0.18
Line: LPK15_335
$R^2$: 0.99
$R^2$: 1.00
$R^2$: 0.99
Line: LPK15_11263
$R^2$: 0.93
$R^2$: 0.96
$R^2$: 0.93
Line: LPK15_11963
$R^2$: 0.95
$R^2$: 0.93
$R^2$: 0.91
Line: LPK15_12444
$R^2$: 0.98
$R^2$: 0.99
$R^2$: 0.98
Line: LPK15_14588
$R^2$: 0.87
$R^2$: 0.88
$R^2$: 0.85
10 30 50 72

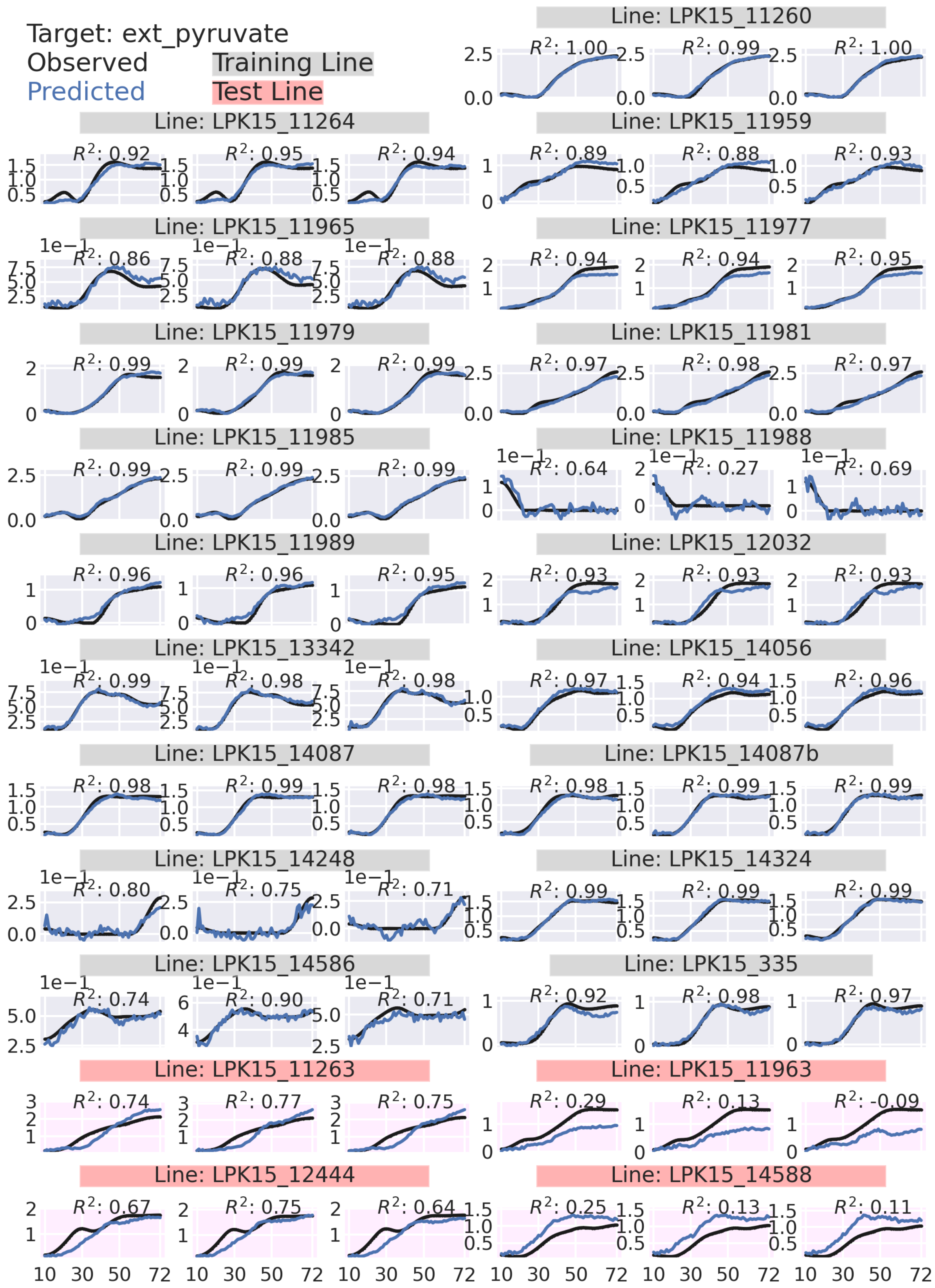

Target: ext_pyruvate
Observed
Predicted
Training Line
Test Line
Line: LPK15_11260
Line: LPK15_11264
Line: LPK15_11959
Line: LPK15_11965
Line: LPK15_11977
Line: LPK15_11979
Line: LPK15_11981
Line: LPK15_11985
Line: LPK15_11988
Line: LPK15_11989
Line: LPK15_12032
Line: LPK15_13342
Line: LPK15_14056
Line: LPK15_14087
Line: LPK15_14087b
Line: LPK15_14248
Line: LPK15_14324
Line: LPK15_14586
Line: LPK15_335
Line: LPK15_11263
Line: LPK15_11963
Line: LPK15_12444
Line: LPK15_14588

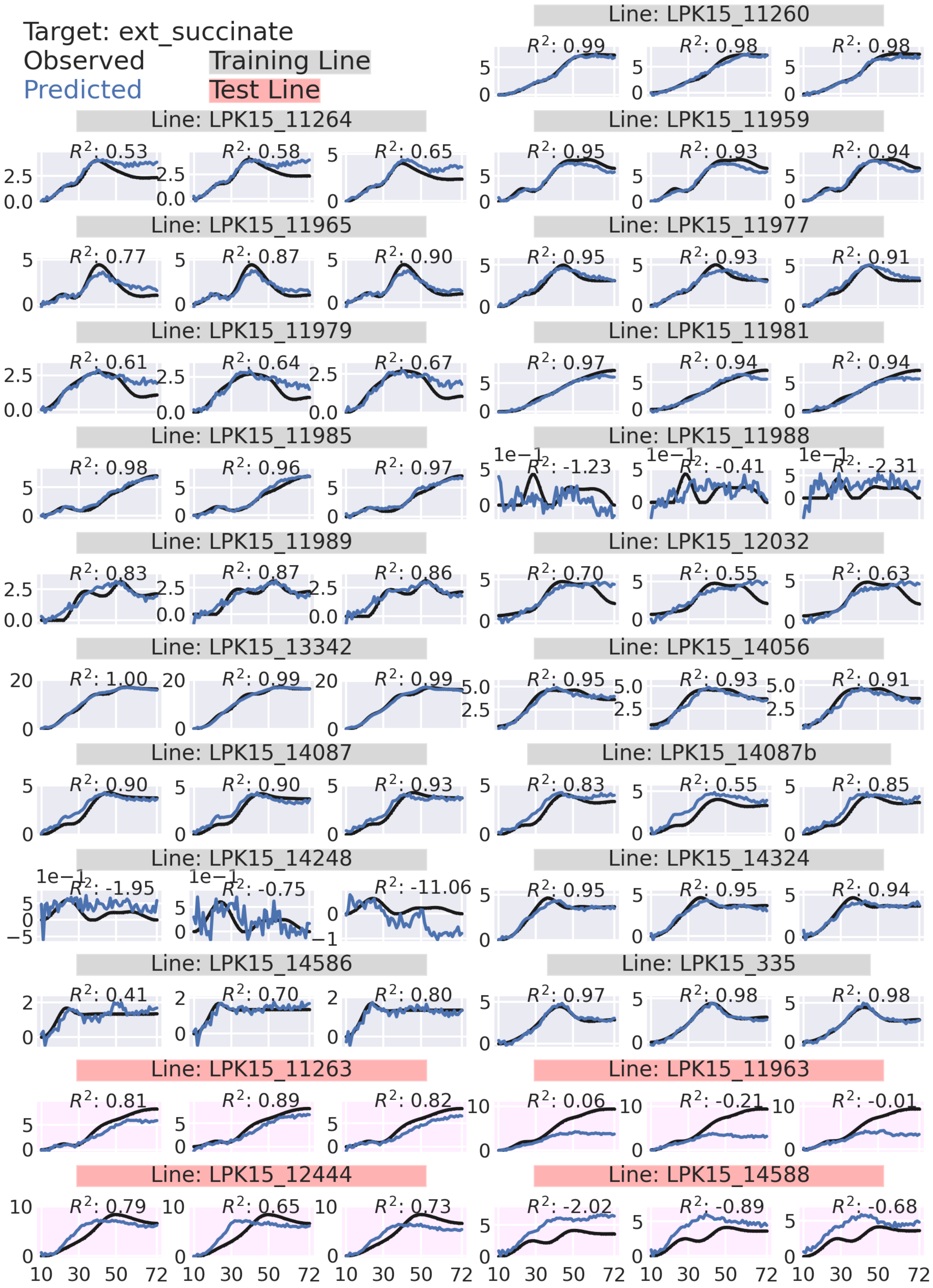
Target: ext_succinate
Observed
Predicted
Training Line
Test Line
Line: LPK15_11260
Line: LPK15_11264
Line: LPK15_11959
Line: LPK15_11965
Line: LPK15_11977
Line: LPK15_11979
Line: LPK15_11981
Line: LPK15_11985
Line: LPK15_11988
Line: LPK15_11989
Line: LPK15_12032
Line: LPK15_13342
Line: LPK15_14056
Line: LPK15_14087
Line: LPK15_14087b
Line: LPK15_14248
Line: LPK15_14324
Line: LPK15_14586
Line: LPK15_335
Line: LPK15_11263
Line: LPK15_11963
Line: LPK15_12444
Line: LPK15_14588

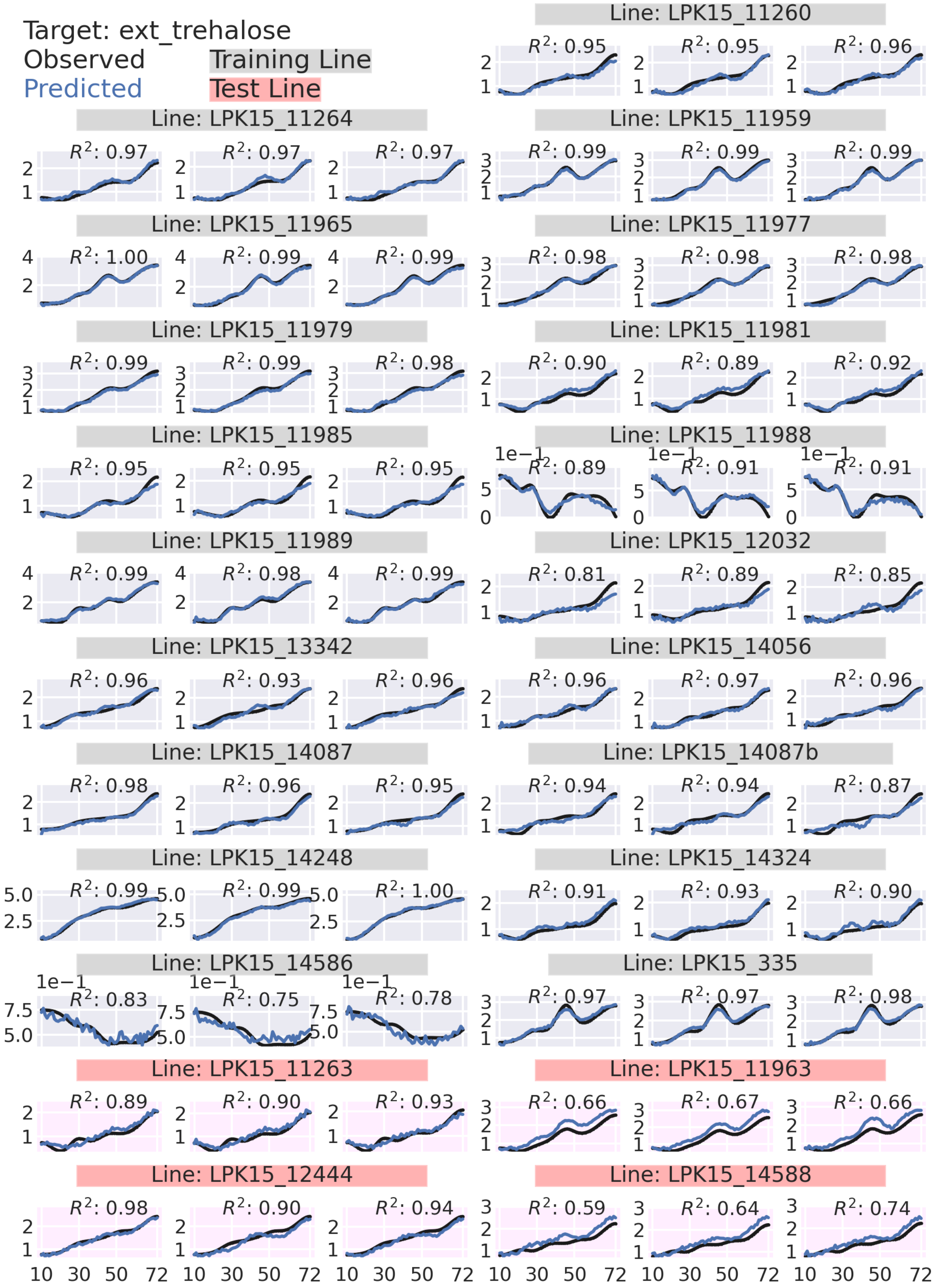
Target: ext_trehalose
Observed
Predicted
Training Line
Test Line
Line: LPK15_11260
Line: LPK15_11264
Line: LPK15_11959
Line: LPK15_11965
Line: LPK15_11977
Line: LPK15_11979
Line: LPK15_11981
Line: LPK15_11985
Line: LPK15_11988
Line: LPK15_11989
Line: LPK15_12032
Line: LPK15_13342
Line: LPK15_14056
Line: LPK15_14087
Line: LPK15_14087b
Line: LPK15_14248
Line: LPK15_14324
Line: LPK15_14586
Line: LPK15_335
Line: LPK15_11263
Line: LPK15_11963
Line: LPK15_12444
Line: LPK15_14588

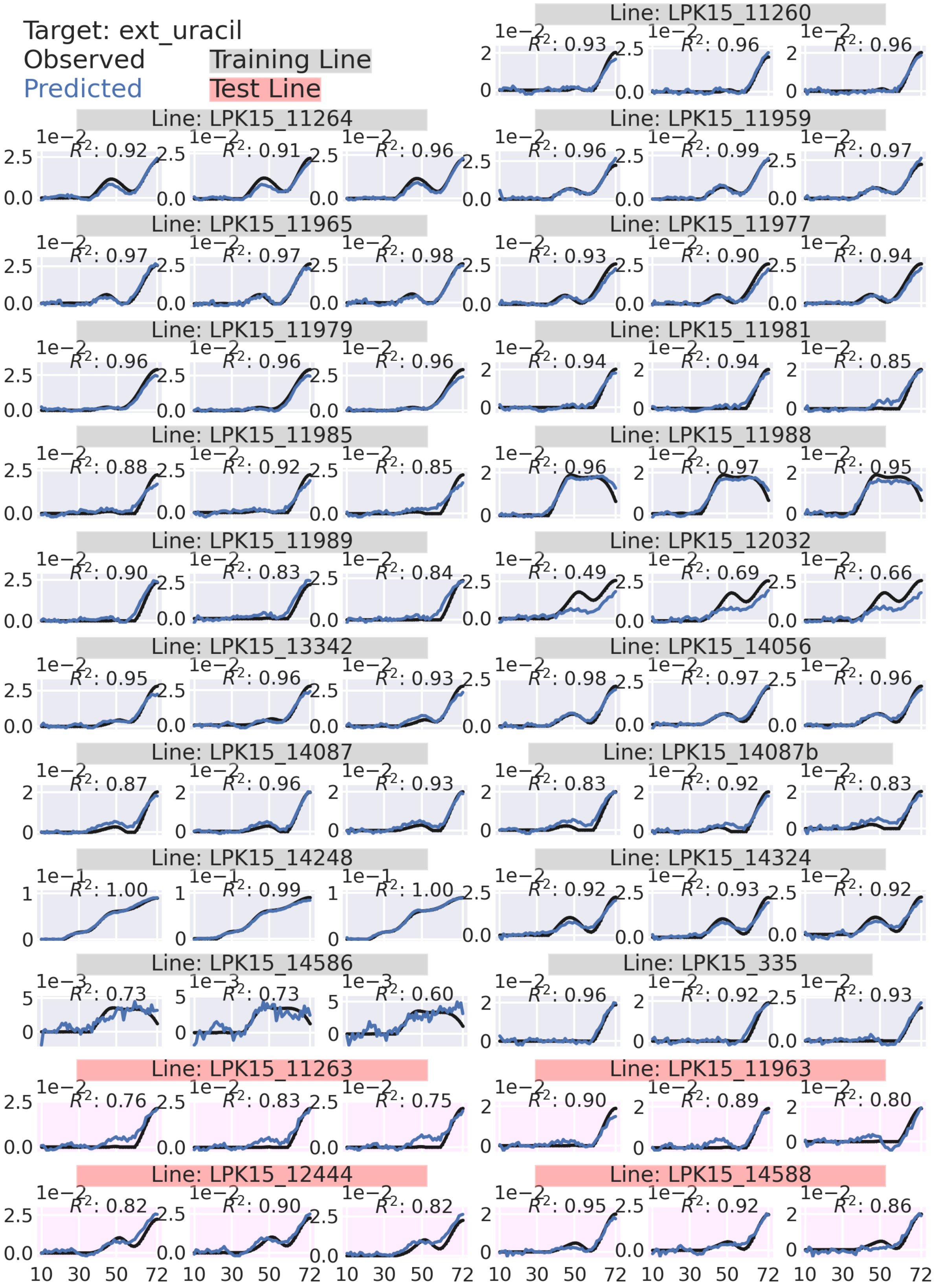
Target: ext_uracil
Observed
Predicted
Training Line
Test Line
Line: LPK15_11260
$R^2$: 0.93
$R^2$: 0.96
$R^2$: 0.96
Line: LPK15_11264
$R^2$: 0.92
$R^2$: 0.91
$R^2$: 0.96
Line: LPK15_11959
$R^2$: 0.96
$R^2$: 0.99
$R^2$: 0.97
Line: LPK15_11965
$R^2$: 0.97
$R^2$: 0.97
$R^2$: 0.98
Line: LPK15_11977
$R^2$: 0.93
$R^2$: 0.90
$R^2$: 0.94
Line: LPK15_11979
$R^2$: 0.96
$R^2$: 0.96
$R^2$: 0.96
Line: LPK15_11981
$R^2$: 0.94
$R^2$: 0.94
$R^2$: 0.85
Line: LPK15_11985
$R^2$: 0.88
$R^2$: 0.92
$R^2$: 0.85
Line: LPK15_11988
$R^2$: 0.96
$R^2$: 0.97
$R^2$: 0.95
Line: LPK15_11989
$R^2$: 0.90
$R^2$: 0.83
$R^2$: 0.84
Line: LPK15_12032
$R^2$: 0.49
$R^2$: 0.69
$R^2$: 0.66
Line: LPK15_13342
$R^2$: 0.95
$R^2$: 0.96
$R^2$: 0.93
Line: LPK15_14056
$R^2$: 0.98
$R^2$: 0.97
$R^2$: 0.96
Line: LPK15_14087
$R^2$: 0.87
$R^2$: 0.96
$R^2$: 0.93
Line: LPK15_14087b
$R^2$: 0.83
$R^2$: 0.92
$R^2$: 0.83
Line: LPK15_14248
$R^2$: 1.00
$R^2$: 0.99
$R^2$: 1.00
Line: LPK15_14324
$R^2$: 0.92
$R^2$: 0.93
$R^2$: 0.92
Line: LPK15_14586
$R^2$: 0.73
$R^2$: 0.73
$R^2$: 0.60
Line: LPK15_335
$R^2$: 0.96
$R^2$: 0.92
$R^2$: 0.93
Line: LPK15_11263
$R^2$: 0.76
$R^2$: 0.83
$R^2$: 0.75
Line: LPK15_11963
$R^2$: 0.90
$R^2$: 0.89
$R^2$: 0.80
Line: LPK15_12444
$R^2$: 0.82
$R^2$: 0.90
$R^2$: 0.82
Line: LPK15_14588
$R^2$: 0.95
$R^2$: 0.92
$R^2$: 0.86
1e−2
1e−1
1e−3
10 30 50 72

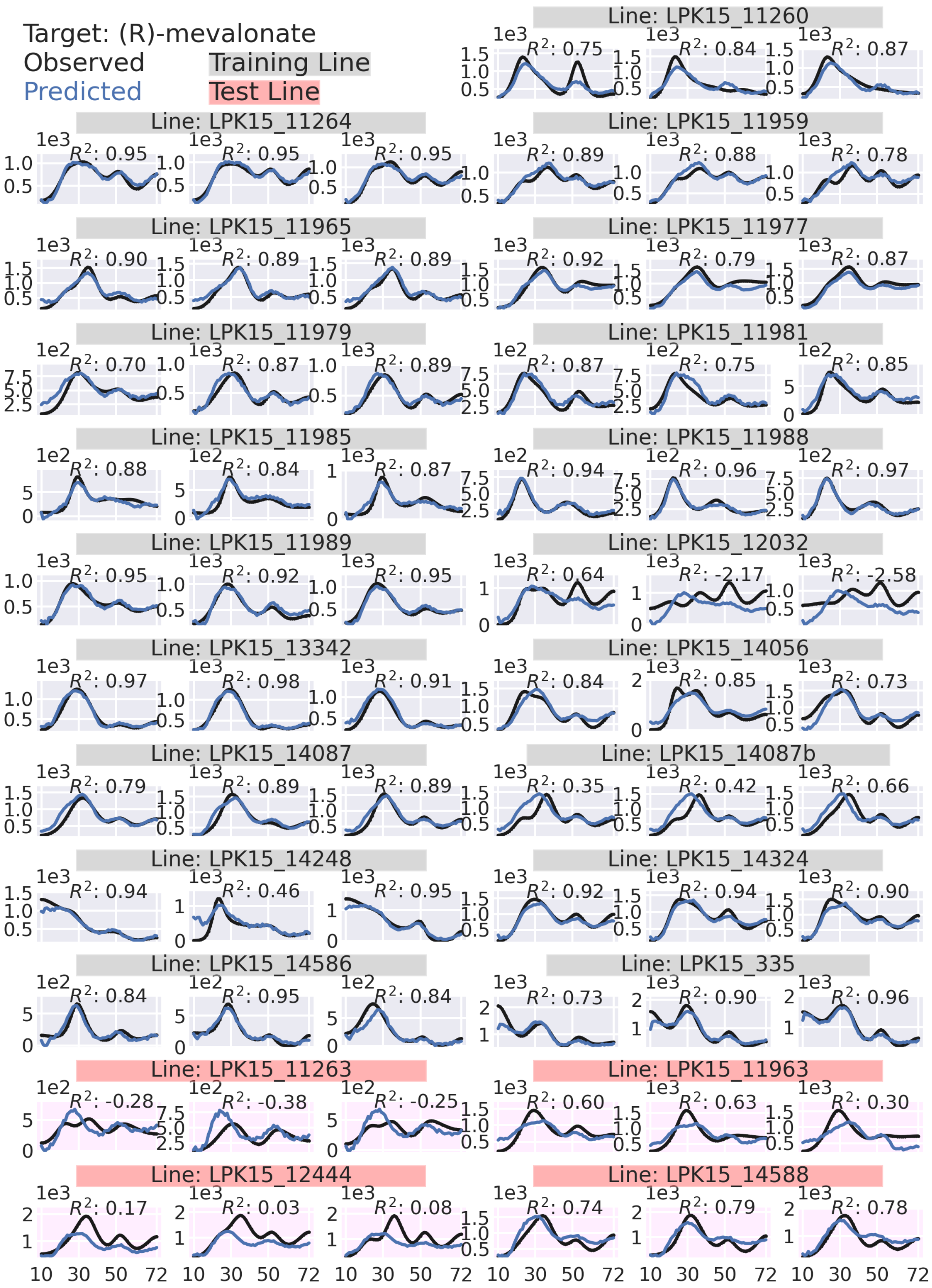
Target: (R)-mevalonate
Observed
Predicted
Training Line
Test Line
Line: LPK15_11260
R²: 0.75
R²: 0.84
R²: 0.87
Line: LPK15_11264
R²: 0.95
R²: 0.95
R²: 0.95
Line: LPK15_11959
R²: 0.89
R²: 0.88
R²: 0.78
Line: LPK15_11965
R²: 0.90
R²: 0.89
R²: 0.89
Line: LPK15_11977
R²: 0.92
R²: 0.79
R²: 0.87
Line: LPK15_11979
R²: 0.70
R²: 0.87
R²: 0.89
Line: LPK15_11981
R²: 0.87
R²: 0.75
R²: 0.85
Line: LPK15_11985
R²: 0.88
R²: 0.84
R²: 0.87
Line: LPK15_11988
R²: 0.94
R²: 0.96
R²: 0.97
Line: LPK15_11989
R²: 0.95
R²: 0.92
R²: 0.95
Line: LPK15_12032
R²: 0.64
R²: -2.17
R²: -2.58
Line: LPK15_13342
R²: 0.97
R²: 0.98
R²: 0.91
Line: LPK15_14056
R²: 0.84
R²: 0.85
R²: 0.73
Line: LPK15_14087
R²: 0.79
R²: 0.89
R²: 0.89
Line: LPK15_14087b
R²: 0.35
R²: 0.42
R²: 0.66
Line: LPK15_14248
R²: 0.94
R²: 0.46
R²: 0.95
Line: LPK15_14324
R²: 0.92
R²: 0.94
R²: 0.90
Line: LPK15_14586
R²: 0.84
R²: 0.95
R²: 0.84
Line: LPK15_335
R²: 0.73
R²: 0.90
R²: 0.96
Line: LPK15_11263
R²: -0.28
R²: -0.38
R²: -0.25
Line: LPK15_11963
R²: 0.60
R²: 0.63
R²: 0.30
Line: LPK15_12444
R²: 0.17
R²: 0.03
R²: 0.08
Line: LPK15_14588
R²: 0.74
R²: 0.79
R²: 0.78
10 30 50 72

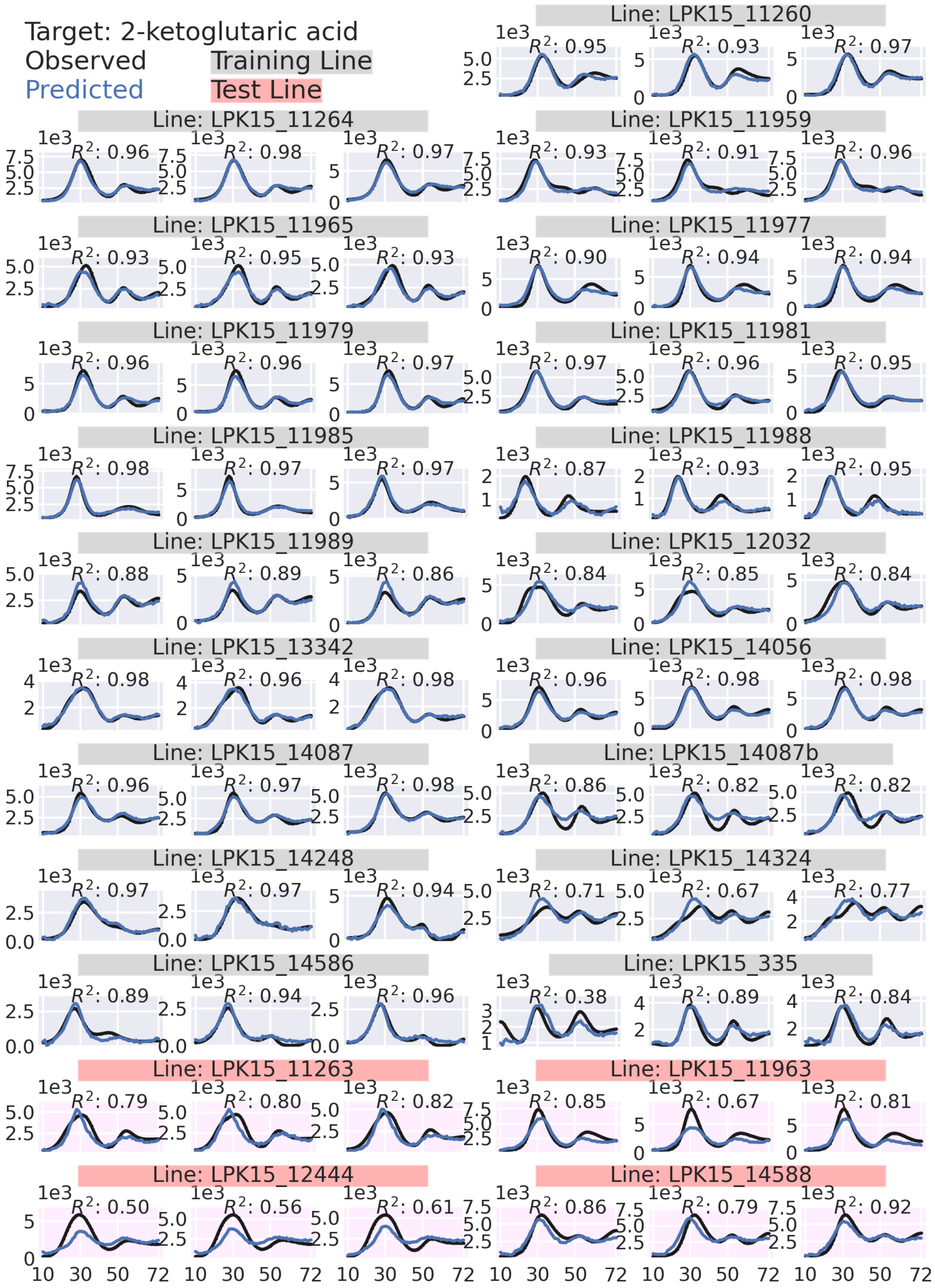
Target: 2-ketoglutaric acid
Observed
Predicted
Training Line
Test Line
Line: LPK15_11260
Line: LPK15_11264
Line: LPK15_11959
Line: LPK15_11965
Line: LPK15_11977
Line: LPK15_11979
Line: LPK15_11981
Line: LPK15_11985
Line: LPK15_11988
Line: LPK15_11989
Line: LPK15_12032
Line: LPK15_13342
Line: LPK15_14056
Line: LPK15_14087
Line: LPK15_14087b
Line: LPK15_14248
Line: LPK15_14324
Line: LPK15_14586
Line: LPK15_335
Line: LPK15_11263
Line: LPK15_11963
Line: LPK15_12444
Line: LPK15_14588
1e3
10 30 50 72

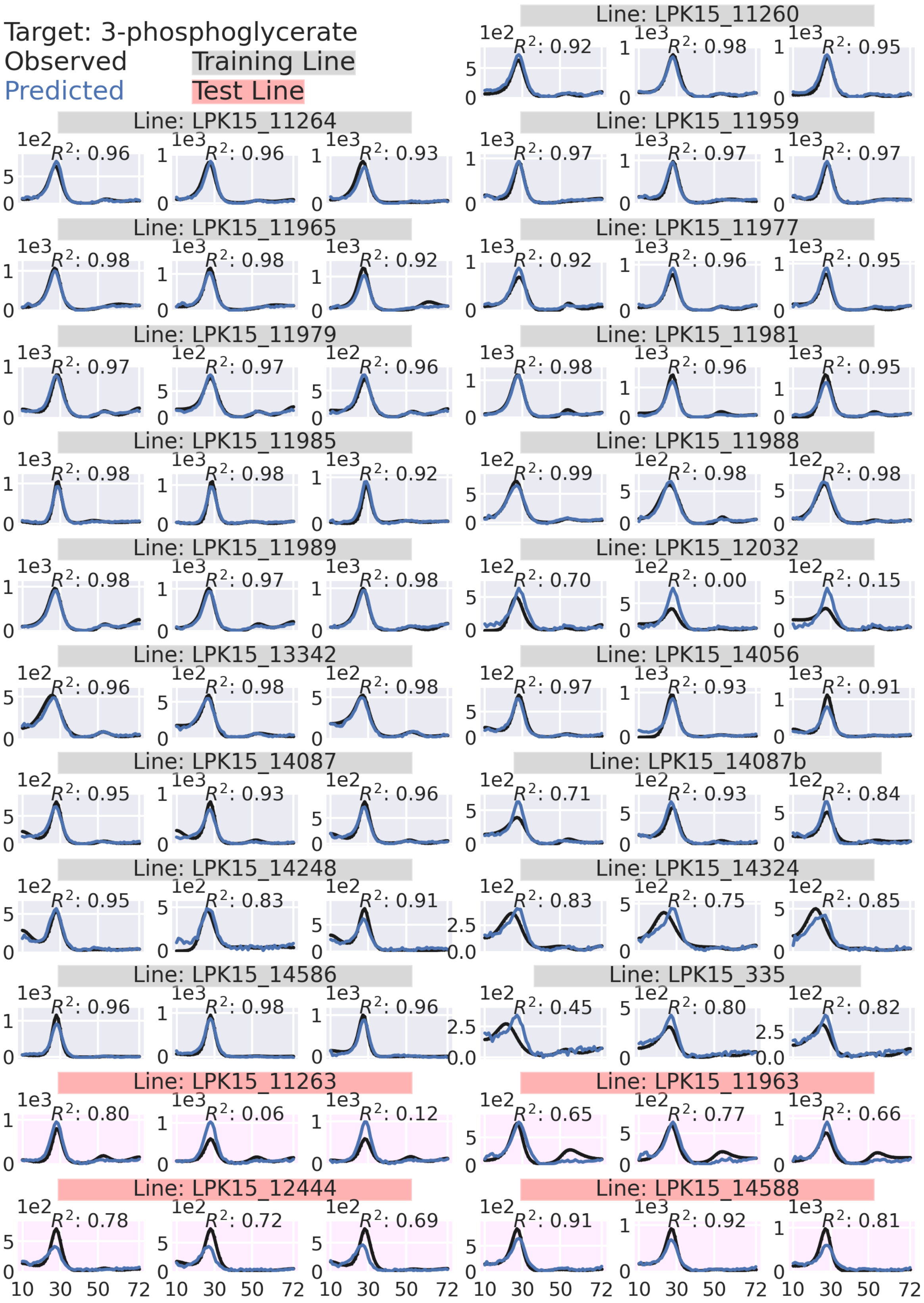

Target: 3-phosphoglycerate
Observed
Predicted
Training Line
Test Line
Line: LPK15_11260
$R^2$: 0.92
$R^2$: 0.98
$R^2$: 0.95
Line: LPK15_11264
$R^2$: 0.96
$R^2$: 0.96
$R^2$: 0.93
Line: LPK15_11959
$R^2$: 0.97
$R^2$: 0.97
$R^2$: 0.97
Line: LPK15_11965
$R^2$: 0.98
$R^2$: 0.98
$R^2$: 0.92
Line: LPK15_11977
$R^2$: 0.92
$R^2$: 0.96
$R^2$: 0.95
Line: LPK15_11979
$R^2$: 0.97
$R^2$: 0.97
$R^2$: 0.96
Line: LPK15_11981
$R^2$: 0.98
$R^2$: 0.96
$R^2$: 0.95
Line: LPK15_11985
$R^2$: 0.98
$R^2$: 0.98
$R^2$: 0.92
Line: LPK15_11988
$R^2$: 0.99
$R^2$: 0.98
$R^2$: 0.98
Line: LPK15_11989
$R^2$: 0.98
$R^2$: 0.97
$R^2$: 0.98
Line: LPK15_12032
$R^2$: 0.70
$R^2$: 0.00
$R^2$: 0.15
Line: LPK15_13342
$R^2$: 0.96
$R^2$: 0.98
$R^2$: 0.98
Line: LPK15_14056
$R^2$: 0.97
$R^2$: 0.93
$R^2$: 0.91
Line: LPK15_14087
$R^2$: 0.95
$R^2$: 0.93
$R^2$: 0.96
Line: LPK15_14087b
$R^2$: 0.71
$R^2$: 0.93
$R^2$: 0.84
Line: LPK15_14248
$R^2$: 0.95
$R^2$: 0.83
$R^2$: 0.91
Line: LPK15_14324
$R^2$: 0.83
$R^2$: 0.75
$R^2$: 0.85
Line: LPK15_14586
$R^2$: 0.96
$R^2$: 0.98
$R^2$: 0.96
Line: LPK15_335
$R^2$: 0.45
$R^2$: 0.80
$R^2$: 0.82
Line: LPK15_11263
$R^2$: 0.80
$R^2$: 0.06
$R^2$: 0.12
Line: LPK15_11963
$R^2$: 0.65
$R^2$: 0.77
$R^2$: 0.66
Line: LPK15_12444
$R^2$: 0.78
$R^2$: 0.72
$R^2$: 0.69
Line: LPK15_14588
$R^2$: 0.91
$R^2$: 0.92
$R^2$: 0.81
10 30 50 72

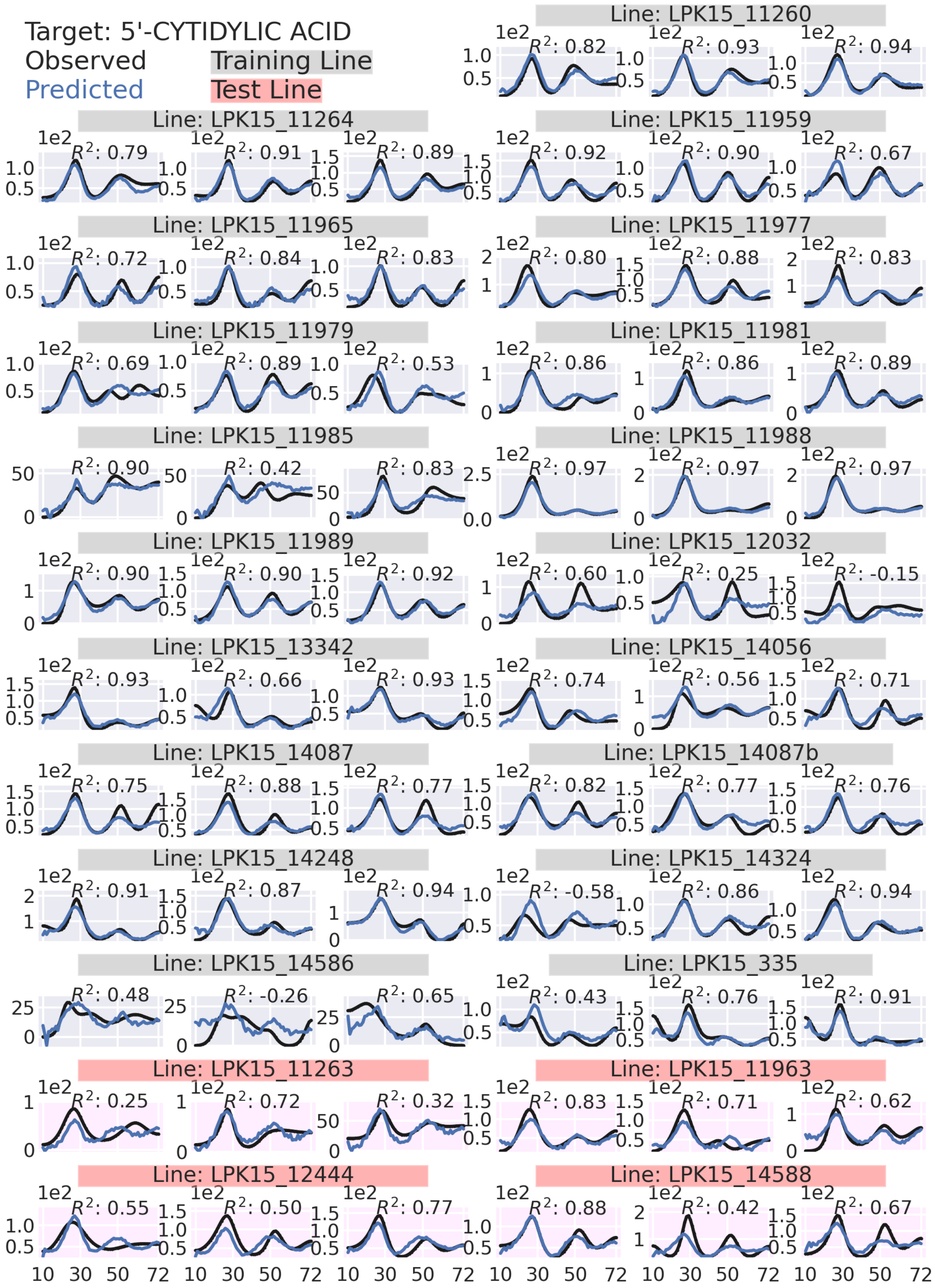

Target: 5'-CYTIDYLIC ACID
Observed
Predicted
Training Line
Test Line
Line: LPK15_11260
R²: 0.82
R²: 0.93
R²: 0.94
Line: LPK15_11264
R²: 0.79
R²: 0.91
R²: 0.89
Line: LPK15_11959
R²: 0.92
R²: 0.90
R²: 0.67
Line: LPK15_11965
R²: 0.72
R²: 0.84
R²: 0.83
Line: LPK15_11977
R²: 0.80
R²: 0.88
R²: 0.83
Line: LPK15_11979
R²: 0.69
R²: 0.89
R²: 0.53
Line: LPK15_11981
R²: 0.86
R²: 0.86
R²: 0.89
Line: LPK15_11985
R²: 0.90
R²: 0.42
R²: 0.83
Line: LPK15_11988
R²: 0.97
R²: 0.97
R²: 0.97
Line: LPK15_11989
R²: 0.90
R²: 0.90
R²: 0.92
Line: LPK15_12032
R²: 0.60
R²: 0.25
R²: -0.15
Line: LPK15_13342
R²: 0.93
R²: 0.66
R²: 0.93
Line: LPK15_14056
R²: 0.74
R²: 0.56
R²: 0.71
Line: LPK15_14087
R²: 0.75
R²: 0.88
R²: 0.77
Line: LPK15_14087b
R²: 0.82
R²: 0.77
R²: 0.76
Line: LPK15_14248
R²: 0.91
R²: 0.87
R²: 0.94
Line: LPK15_14324
R²: -0.58
R²: 0.86
R²: 0.94
Line: LPK15_14586
R²: 0.48
R²: -0.26
R²: 0.65
Line: LPK15_335
R²: 0.43
R²: 0.76
R²: 0.91
Line: LPK15_11263
R²: 0.25
R²: 0.72
R²: 0.32
Line: LPK15_11963
R²: 0.83
R²: 0.71
R²: 0.62
Line: LPK15_12444
R²: 0.55
R²: 0.50
R²: 0.77
Line: LPK15_14588
R²: 0.88
R²: 0.42
R²: 0.67
10 30 50 72

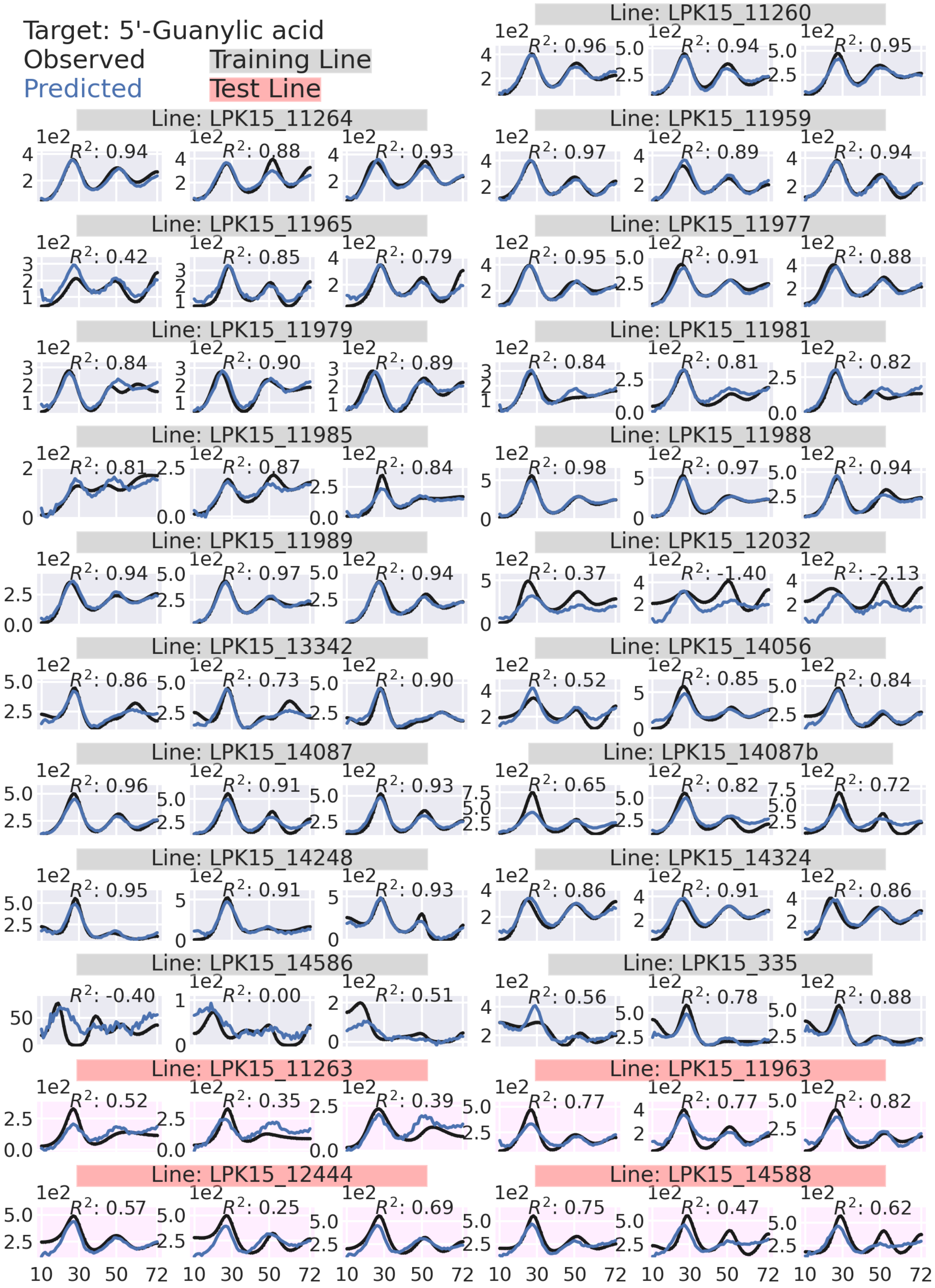

Target: 5'-Guanylic acid
Observed
Predicted
Training Line
Test Line
Line: LPK15_11260
R²: 0.96 R²: 0.94 R²: 0.95
Line: LPK15_11264
R²: 0.94 R²: 0.88 R²: 0.93
Line: LPK15_11959
R²: 0.97 R²: 0.89 R²: 0.94
Line: LPK15_11965
R²: 0.42 R²: 0.85 R²: 0.79
Line: LPK15_11977
R²: 0.95 R²: 0.91 R²: 0.88
Line: LPK15_11979
R²: 0.84 R²: 0.90 R²: 0.89
Line: LPK15_11981
R²: 0.84 R²: 0.81 R²: 0.82
Line: LPK15_11985
R²: 0.81 R²: 0.87 R²: 0.84
Line: LPK15_11988
R²: 0.98 R²: 0.97 R²: 0.94
Line: LPK15_11989
R²: 0.94 R²: 0.97 R²: 0.94
Line: LPK15_12032
R²: 0.37 R²: -1.40 R²: -2.13
Line: LPK15_13342
R²: 0.86 R²: 0.73 R²: 0.90
Line: LPK15_14056
R²: 0.52 R²: 0.85 R²: 0.84
Line: LPK15_14087
R²: 0.96 R²: 0.91 R²: 0.93
Line: LPK15_14087b
R²: 0.65 R²: 0.82 R²: 0.72
Line: LPK15_14248
R²: 0.95 R²: 0.91 R²: 0.93
Line: LPK15_14324
R²: 0.86 R²: 0.91 R²: 0.86
Line: LPK15_14586
R²: -0.40 R²: 0.00 R²: 0.51
Line: LPK15_335
R²: 0.56 R²: 0.78 R²: 0.88
Line: LPK15_11263
R²: 0.52 R²: 0.35 R²: 0.39
Line: LPK15_11963
R²: 0.77 R²: 0.77 R²: 0.82
Line: LPK15_12444
R²: 0.57 R²: 0.25 R²: 0.69
Line: LPK15_14588
R²: 0.75 R²: 0.47 R²: 0.62
10 30 50 72

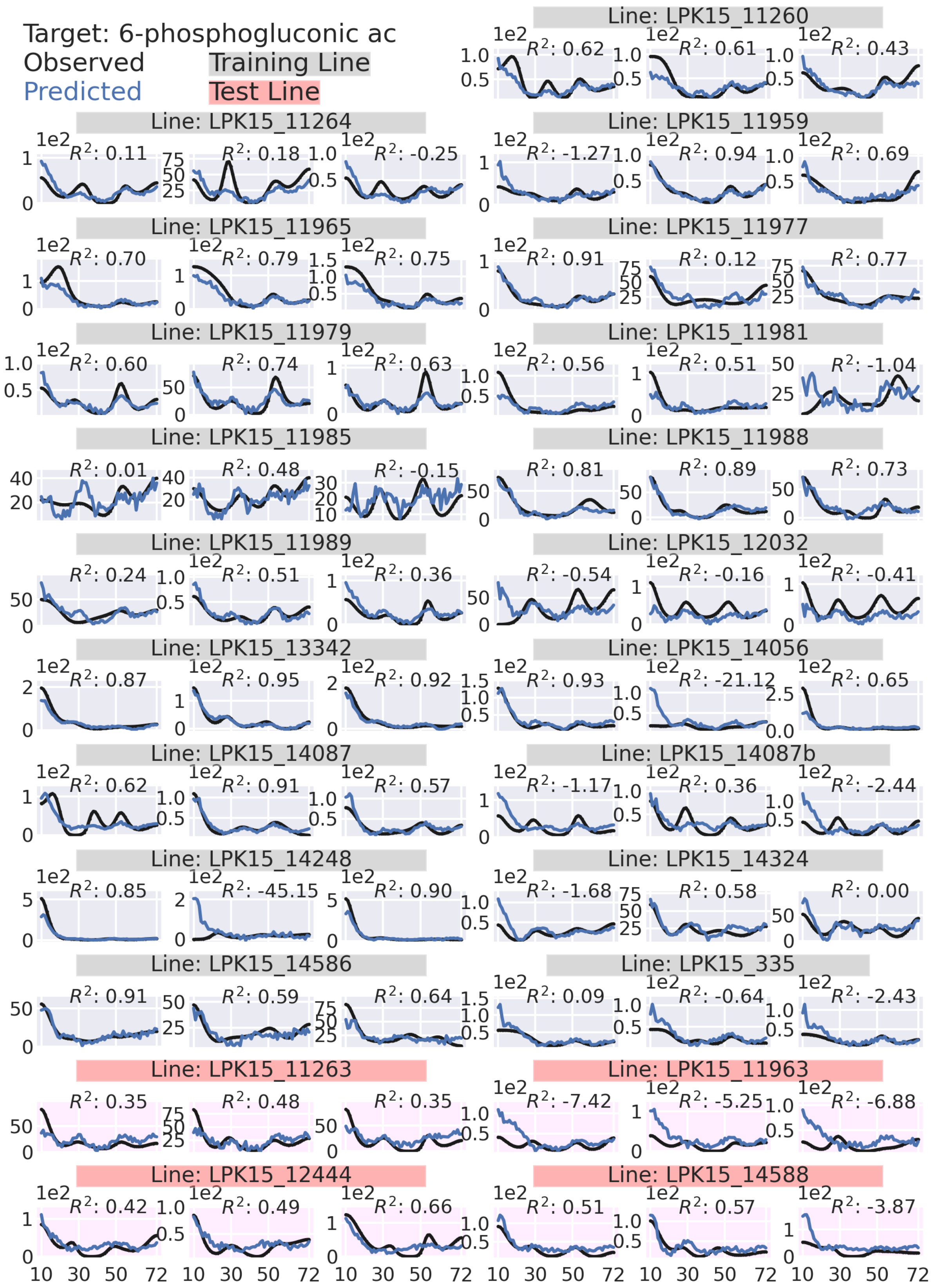

Target: 6-phosphogluconic ac
Observed
Predicted
Training Line
Test Line
Line: LPK15_11260
Line: LPK15_11264
Line: LPK15_11959
Line: LPK15_11965
Line: LPK15_11977
Line: LPK15_11979
Line: LPK15_11981
Line: LPK15_11985
Line: LPK15_11988
Line: LPK15_11989
Line: LPK15_12032
Line: LPK15_13342
Line: LPK15_14056
Line: LPK15_14087
Line: LPK15_14087b
Line: LPK15_14248
Line: LPK15_14324
Line: LPK15_14586
Line: LPK15_335
Line: LPK15_11263
Line: LPK15_11963
Line: LPK15_12444
Line: LPK15_14588

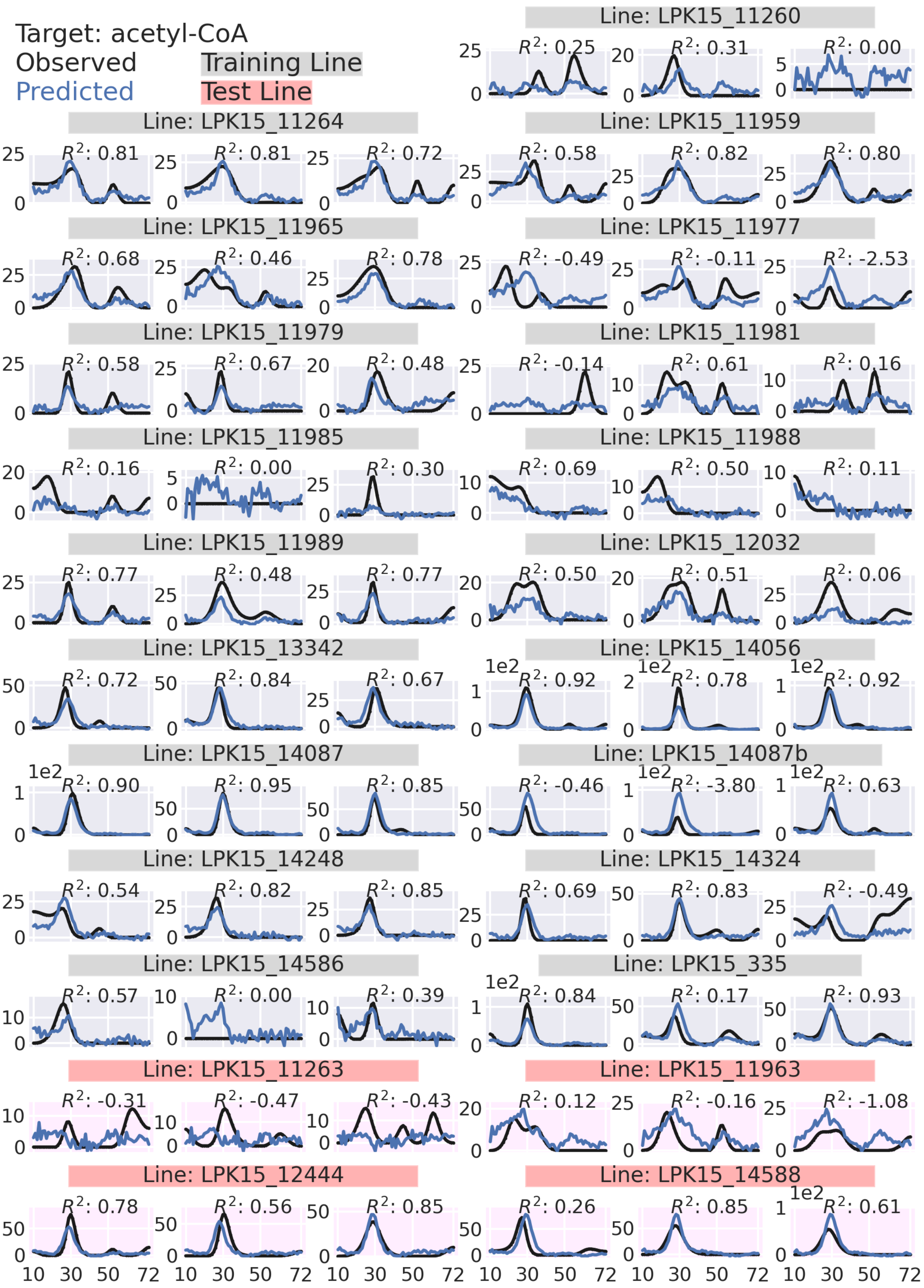

Target: acetyl-CoA
Observed
Predicted
Training Line
Test Line
Line: LPK15_11260
Line: LPK15_11264
Line: LPK15_11959
Line: LPK15_11965
Line: LPK15_11977
Line: LPK15_11979
Line: LPK15_11981
Line: LPK15_11985
Line: LPK15_11988
Line: LPK15_11989
Line: LPK15_12032
Line: LPK15_13342
Line: LPK15_14056
Line: LPK15_14087
Line: LPK15_14087b
Line: LPK15_14248
Line: LPK15_14324
Line: LPK15_14586
Line: LPK15_335
Line: LPK15_11263
Line: LPK15_11963
Line: LPK15_12444
Line: LPK15_14588

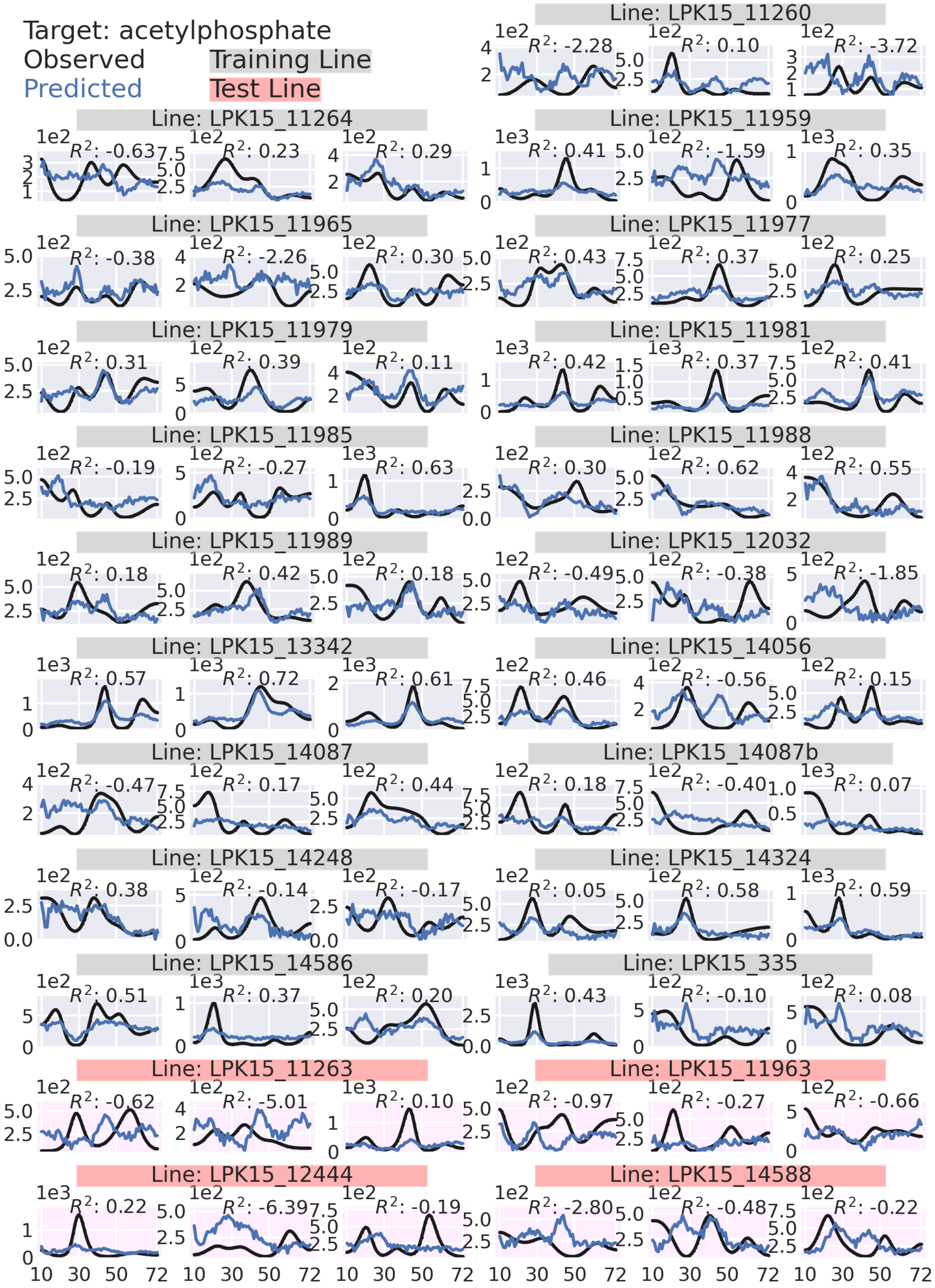
Target: acetylphosphate
Observed
Predicted
Training Line
Test Line
Line: LPK15_11260
Line: LPK15_11264
Line: LPK15_11959
Line: LPK15_11965
Line: LPK15_11977
Line: LPK15_11979
Line: LPK15_11981
Line: LPK15_11985
Line: LPK15_11988
Line: LPK15_11989
Line: LPK15_12032
Line: LPK15_13342
Line: LPK15_14056
Line: LPK15_14087
Line: LPK15_14087b
Line: LPK15_14248
Line: LPK15_14324
Line: LPK15_14586
Line: LPK15_335
Line: LPK15_11263
Line: LPK15_11963
Line: LPK15_12444
Line: LPK15_14588

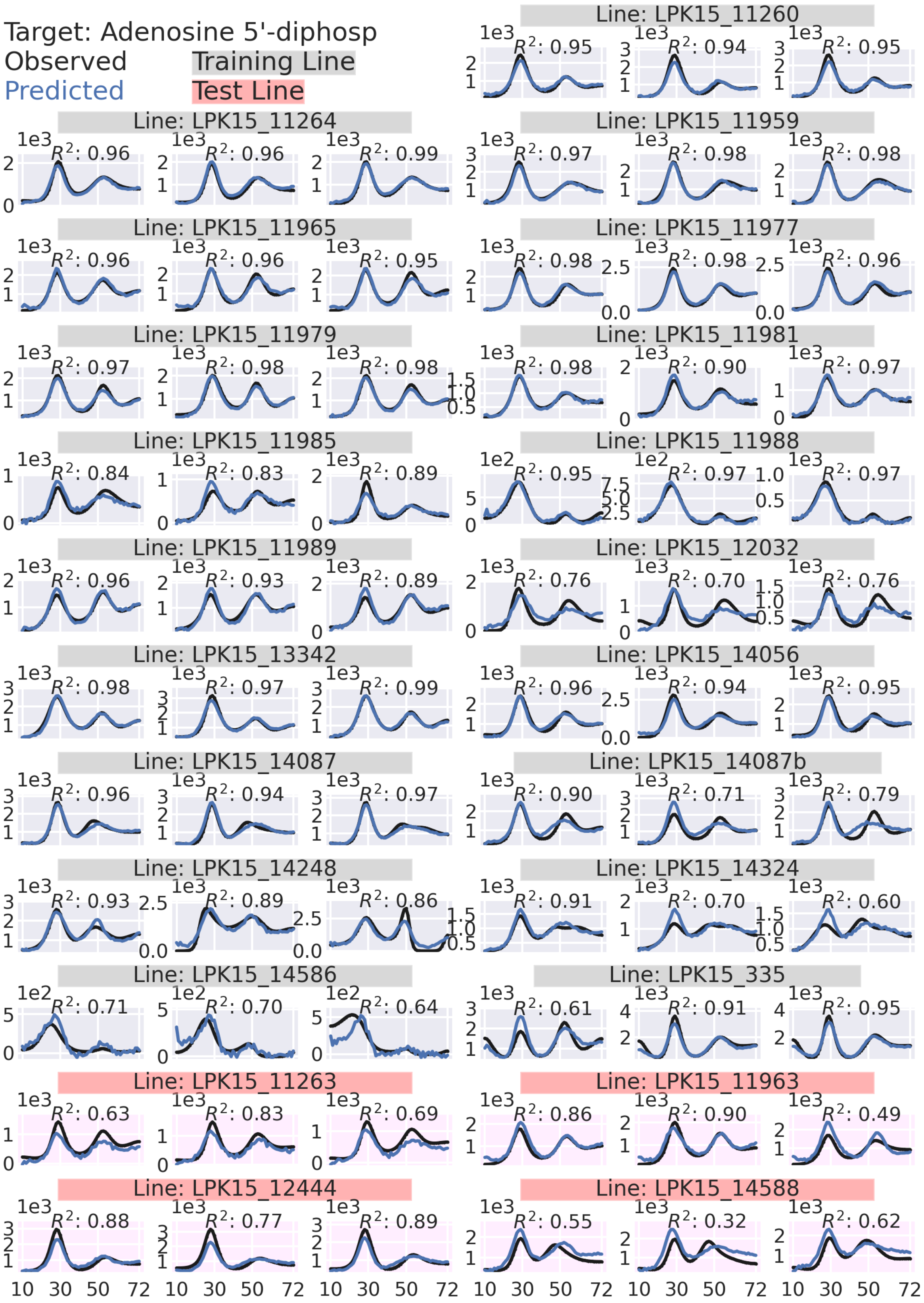

Target: Adenosine 5'-diphosp
Observed
Predicted
Training Line
Test Line
Line: LPK15_11260
Line: LPK15_11264
Line: LPK15_11959
Line: LPK15_11965
Line: LPK15_11977
Line: LPK15_11979
Line: LPK15_11981
Line: LPK15_11985
Line: LPK15_11988
Line: LPK15_11989
Line: LPK15_12032
Line: LPK15_13342
Line: LPK15_14056
Line: LPK15_14087
Line: LPK15_14087b
Line: LPK15_14248
Line: LPK15_14324
Line: LPK15_14586
Line: LPK15_335
Line: LPK15_11263
Line: LPK15_11963
Line: LPK15_12444
Line: LPK15_14588
10 30 50 72

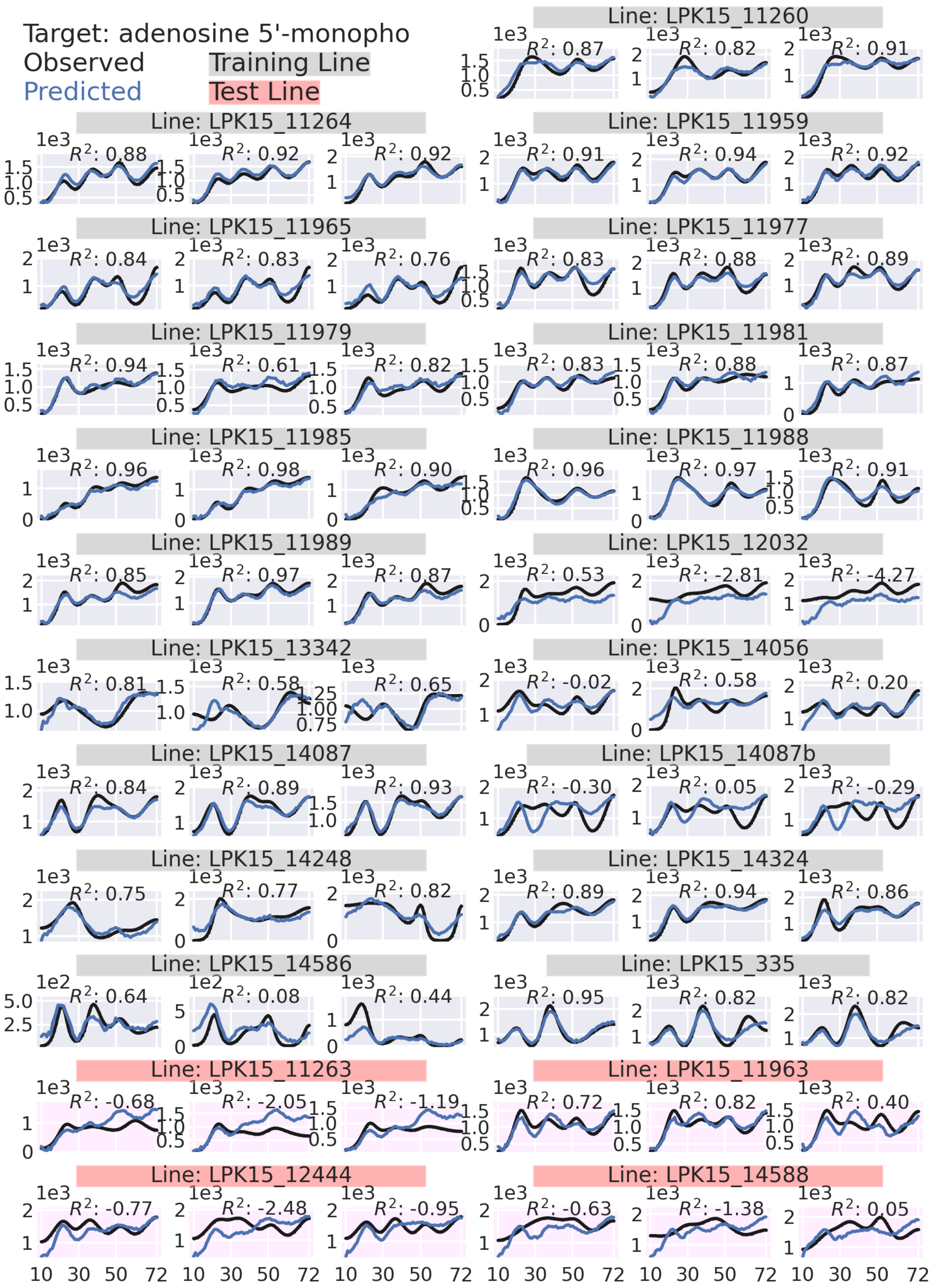

Target: adenosine 5'-monopho
Observed
Predicted
Training Line
Test Line
Line: LPK15_11260
Line: LPK15_11264
Line: LPK15_11959
Line: LPK15_11965
Line: LPK15_11977
Line: LPK15_11979
Line: LPK15_11981
Line: LPK15_11985
Line: LPK15_11988
Line: LPK15_11989
Line: LPK15_12032
Line: LPK15_13342
Line: LPK15_14056
Line: LPK15_14087
Line: LPK15_14087b
Line: LPK15_14248
Line: LPK15_14324
Line: LPK15_14586
Line: LPK15_335
Line: LPK15_11263
Line: LPK15_11963
Line: LPK15_12444
Line: LPK15_14588
$R^2$: 0.87 $R^2$: 0.82 $R^2$: 0.91
$R^2$: 0.88 $R^2$: 0.92 $R^2$: 0.92 $R^2$: 0.91 $R^2$: 0.94 $R^2$: 0.92
$R^2$: 0.84 $R^2$: 0.83 $R^2$: 0.76 $R^2$: 0.83 $R^2$: 0.88 $R^2$: 0.89
$R^2$: 0.94 $R^2$: 0.61 $R^2$: 0.82 $R^2$: 0.83 $R^2$: 0.88 $R^2$: 0.87
$R^2$: 0.96 $R^2$: 0.98 $R^2$: 0.90 $R^2$: 0.96 $R^2$: 0.97 $R^2$: 0.91
$R^2$: 0.85 $R^2$: 0.97 $R^2$: 0.87 $R^2$: 0.53 $R^2$: -2.81 $R^2$: -4.27
$R^2$: 0.81 $R^2$: 0.58 $R^2$: 0.65 $R^2$: -0.02 $R^2$: 0.58 $R^2$: 0.20
$R^2$: 0.84 $R^2$: 0.89 $R^2$: 0.93 $R^2$: -0.30 $R^2$: 0.05 $R^2$: -0.29
$R^2$: 0.75 $R^2$: 0.77 $R^2$: 0.82 $R^2$: 0.89 $R^2$: 0.94 $R^2$: 0.86
$R^2$: 0.64 $R^2$: 0.08 $R^2$: 0.44 $R^2$: 0.95 $R^2$: 0.82 $R^2$: 0.82
$R^2$: -0.68 $R^2$: -2.05 $R^2$: -1.19 $R^2$: 0.72 $R^2$: 0.82 $R^2$: 0.40
$R^2$: -0.77 $R^2$: -2.48 $R^2$: -0.95 $R^2$: -0.63 $R^2$: -1.38 $R^2$: 0.05
1e3
1e2
10 30 50 72

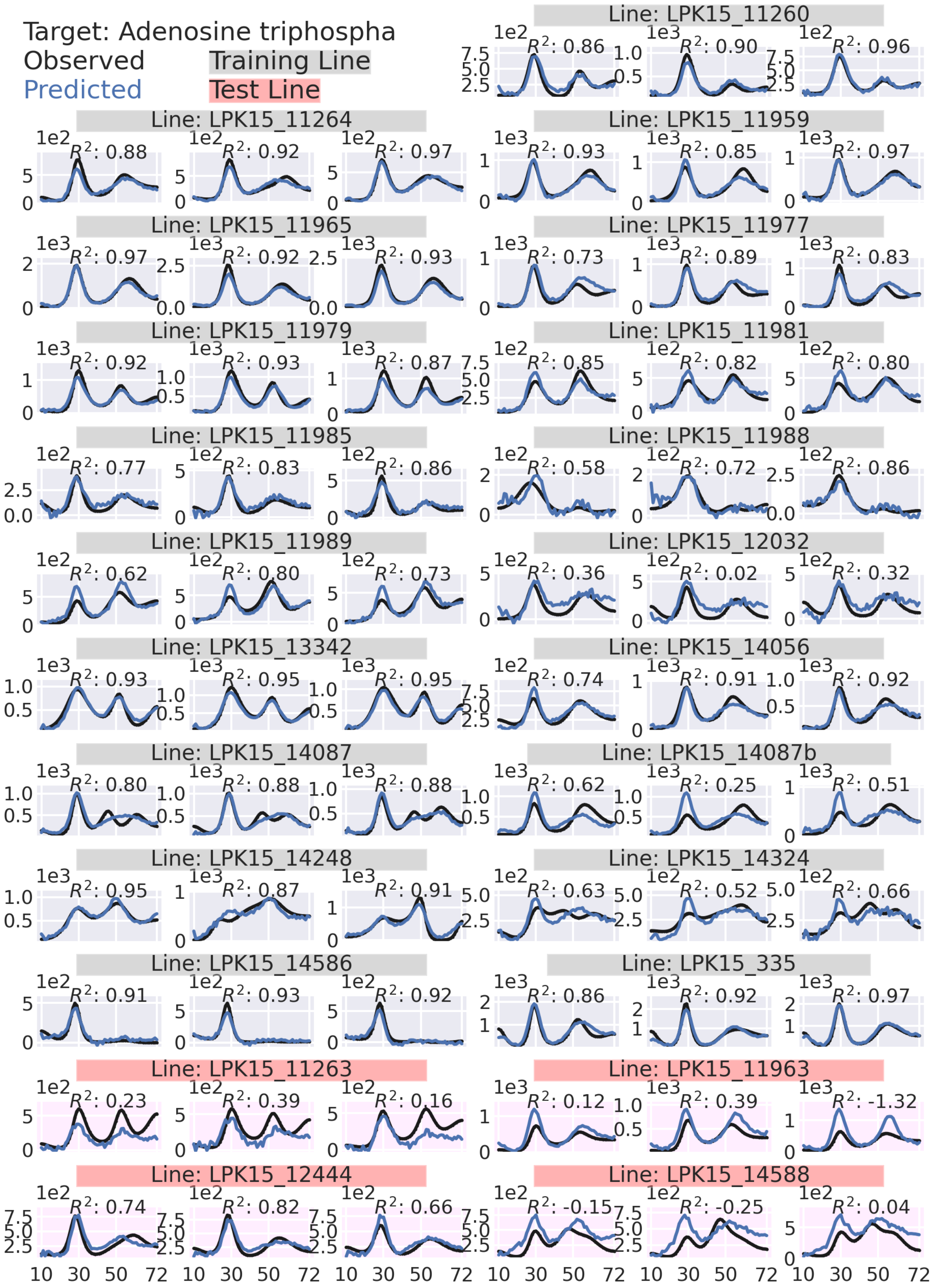

Target: Adenosine triphospha
Observed
Predicted
Training Line
Test Line
Line: LPK15_11260
Line: LPK15_11264
Line: LPK15_11959
Line: LPK15_11965
Line: LPK15_11977
Line: LPK15_11979
Line: LPK15_11981
Line: LPK15_11985
Line: LPK15_11988
Line: LPK15_11989
Line: LPK15_12032
Line: LPK15_13342
Line: LPK15_14056
Line: LPK15_14087
Line: LPK15_14087b
Line: LPK15_14248
Line: LPK15_14324
Line: LPK15_14586
Line: LPK15_335
Line: LPK15_11263
Line: LPK15_11963
Line: LPK15_12444
Line: LPK15_14588

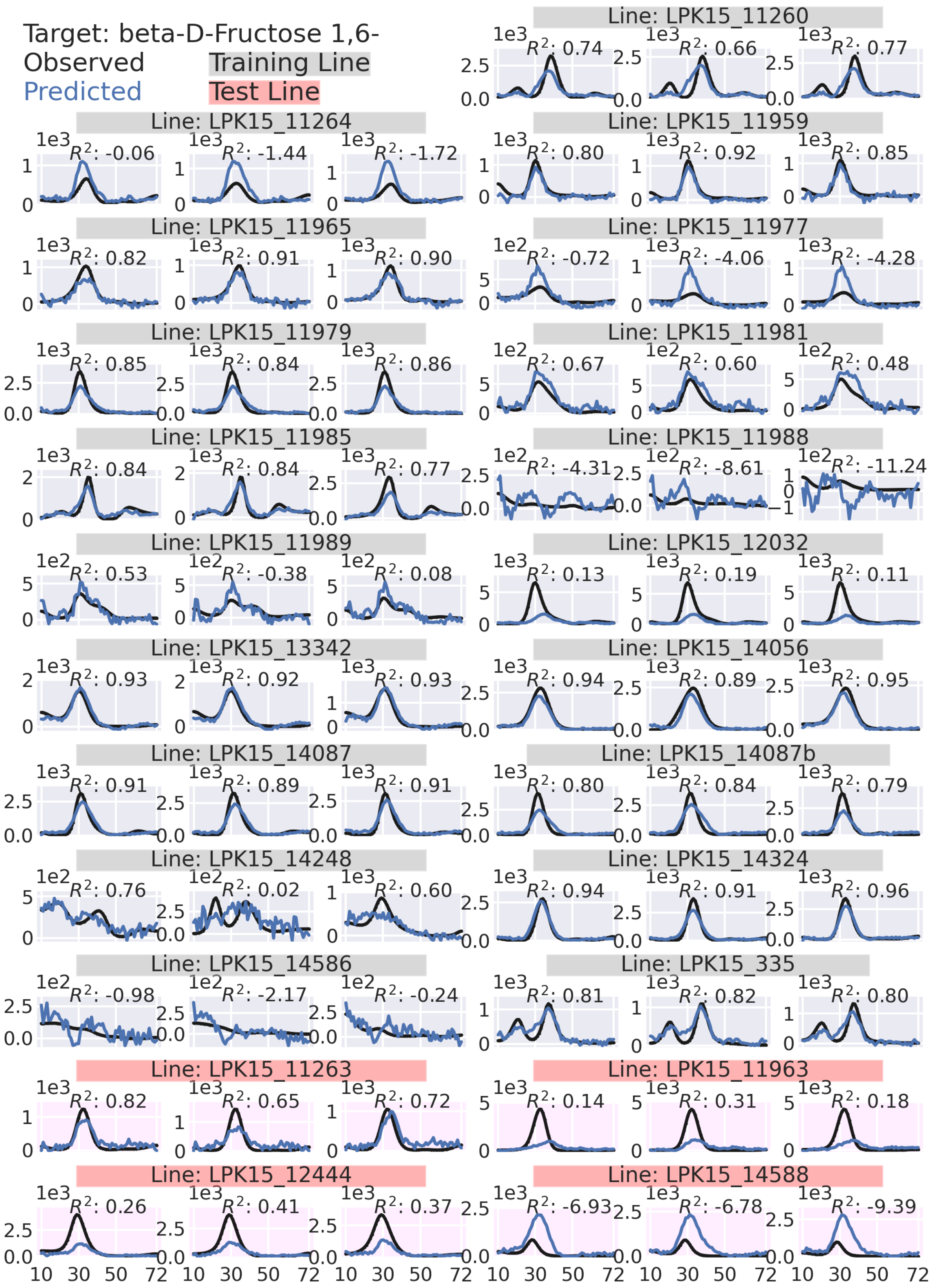

Target: beta-D-Fructose 1,6-
Observed
Predicted
Training Line
Test Line
Line: LPK15_11260
Line: LPK15_11264
Line: LPK15_11959
Line: LPK15_11965
Line: LPK15_11977
Line: LPK15_11979
Line: LPK15_11981
Line: LPK15_11985
Line: LPK15_11988
Line: LPK15_11989
Line: LPK15_12032
Line: LPK15_13342
Line: LPK15_14056
Line: LPK15_14087
Line: LPK15_14087b
Line: LPK15_14248
Line: LPK15_14324
Line: LPK15_14586
Line: LPK15_335
Line: LPK15_11263
Line: LPK15_11963
Line: LPK15_12444
Line: LPK15_14588

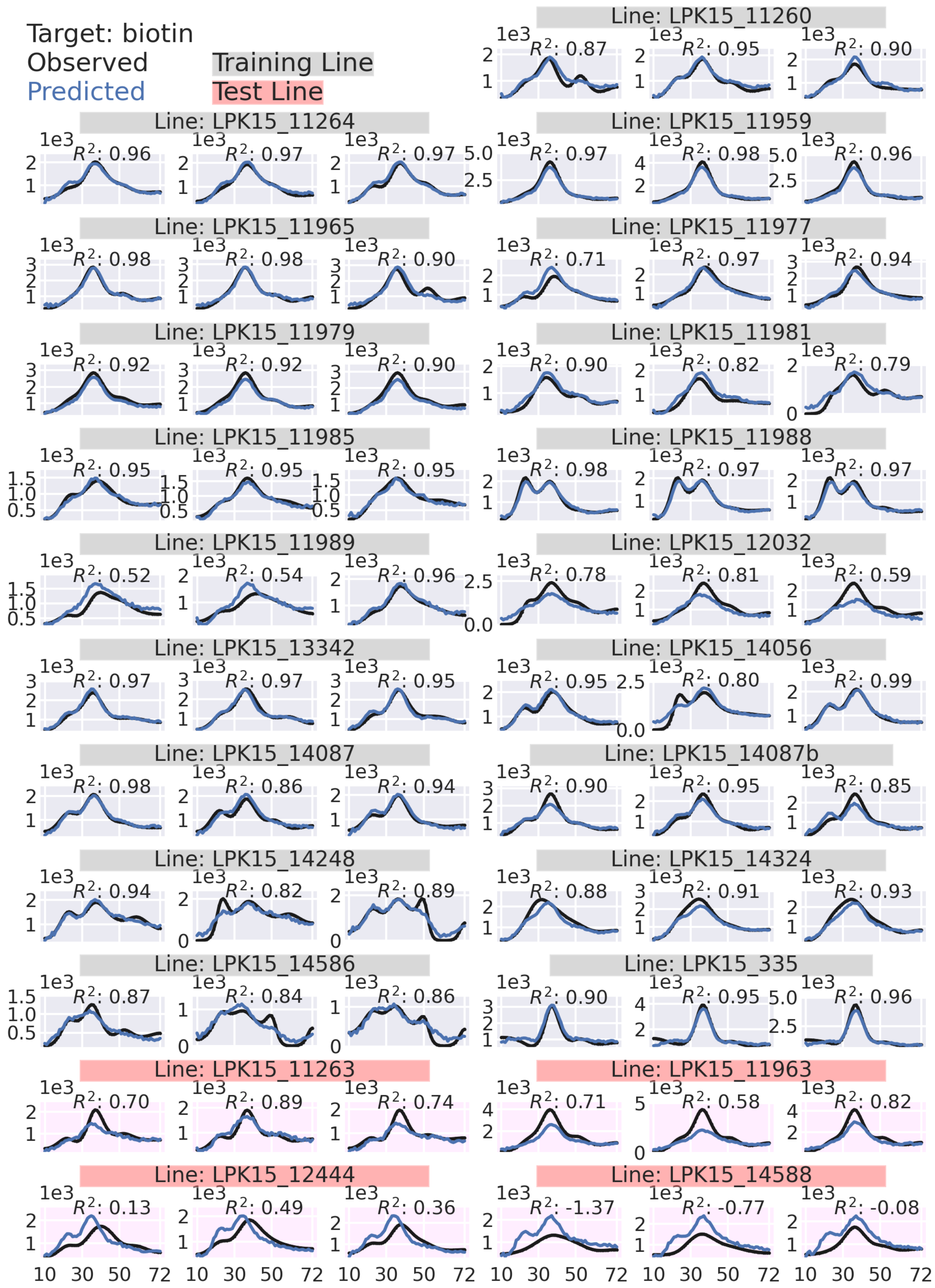

Target: biotin
Observed
Predicted
Training Line
Test Line
Line: LPK15_11260
Line: LPK15_11264
Line: LPK15_11959
Line: LPK15_11965
Line: LPK15_11977
Line: LPK15_11979
Line: LPK15_11981
Line: LPK15_11985
Line: LPK15_11988
Line: LPK15_11989
Line: LPK15_12032
Line: LPK15_13342
Line: LPK15_14056
Line: LPK15_14087
Line: LPK15_14087b
Line: LPK15_14248
Line: LPK15_14324
Line: LPK15_14586
Line: LPK15_335
Line: LPK15_11263
Line: LPK15_11963
Line: LPK15_12444
Line: LPK15_14588

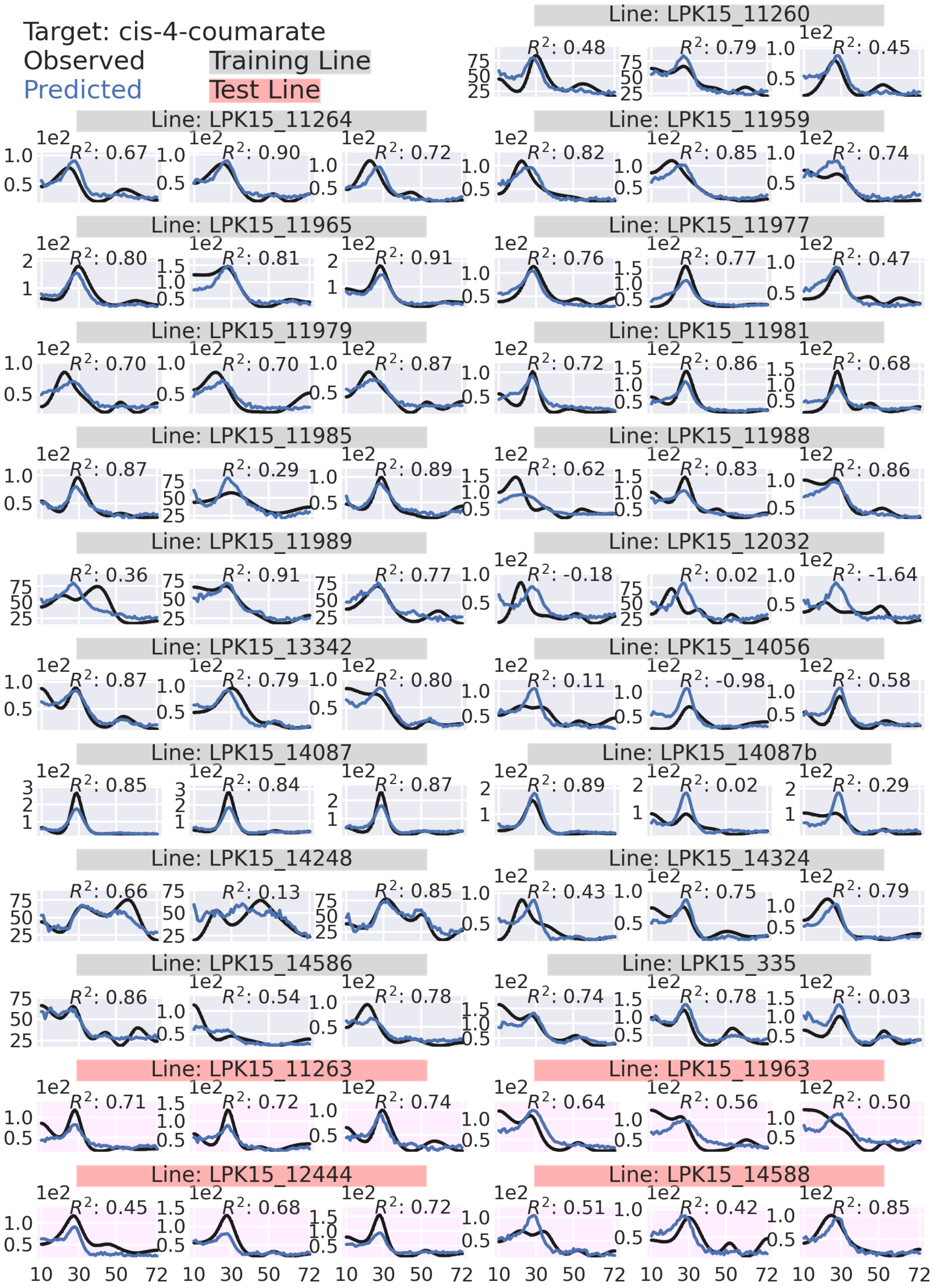

Target: cis-4-coumarate
Observed
Predicted
Training Line
Test Line
Line: LPK15_11260
$R^2$: 0.48
$R^2$: 0.79
$R^2$: 0.45
Line: LPK15_11264
$R^2$: 0.67
$R^2$: 0.90
$R^2$: 0.72
Line: LPK15_11959
$R^2$: 0.82
$R^2$: 0.85
$R^2$: 0.74
Line: LPK15_11965
$R^2$: 0.80
$R^2$: 0.81
$R^2$: 0.91
Line: LPK15_11977
$R^2$: 0.76
$R^2$: 0.77
$R^2$: 0.47
Line: LPK15_11979
$R^2$: 0.70
$R^2$: 0.70
$R^2$: 0.87
Line: LPK15_11981
$R^2$: 0.72
$R^2$: 0.86
$R^2$: 0.68
Line: LPK15_11985
$R^2$: 0.87
$R^2$: 0.29
$R^2$: 0.89
Line: LPK15_11988
$R^2$: 0.62
$R^2$: 0.83
$R^2$: 0.86
Line: LPK15_11989
$R^2$: 0.36
$R^2$: 0.91
$R^2$: 0.77
Line: LPK15_12032
$R^2$: -0.18
$R^2$: 0.02
$R^2$: -1.64
Line: LPK15_13342
$R^2$: 0.87
$R^2$: 0.79
$R^2$: 0.80
Line: LPK15_14056
$R^2$: 0.11
$R^2$: -0.98
$R^2$: 0.58
Line: LPK15_14087
$R^2$: 0.85
$R^2$: 0.84
$R^2$: 0.87
Line: LPK15_14087b
$R^2$: 0.89
$R^2$: 0.02
$R^2$: 0.29
Line: LPK15_14248
$R^2$: 0.66
$R^2$: 0.13
$R^2$: 0.85
Line: LPK15_14324
$R^2$: 0.43
$R^2$: 0.75
$R^2$: 0.79
Line: LPK15_14586
$R^2$: 0.86
$R^2$: 0.54
$R^2$: 0.78
Line: LPK15_335
$R^2$: 0.74
$R^2$: 0.78
$R^2$: 0.03
Line: LPK15_11263
$R^2$: 0.71
$R^2$: 0.72
$R^2$: 0.74
Line: LPK15_11963
$R^2$: 0.64
$R^2$: 0.56
$R^2$: 0.50
Line: LPK15_12444
$R^2$: 0.45
$R^2$: 0.68
$R^2$: 0.72
Line: LPK15_14588
$R^2$: 0.51
$R^2$: 0.42
$R^2$: 0.85
1e2
10 30 50 72

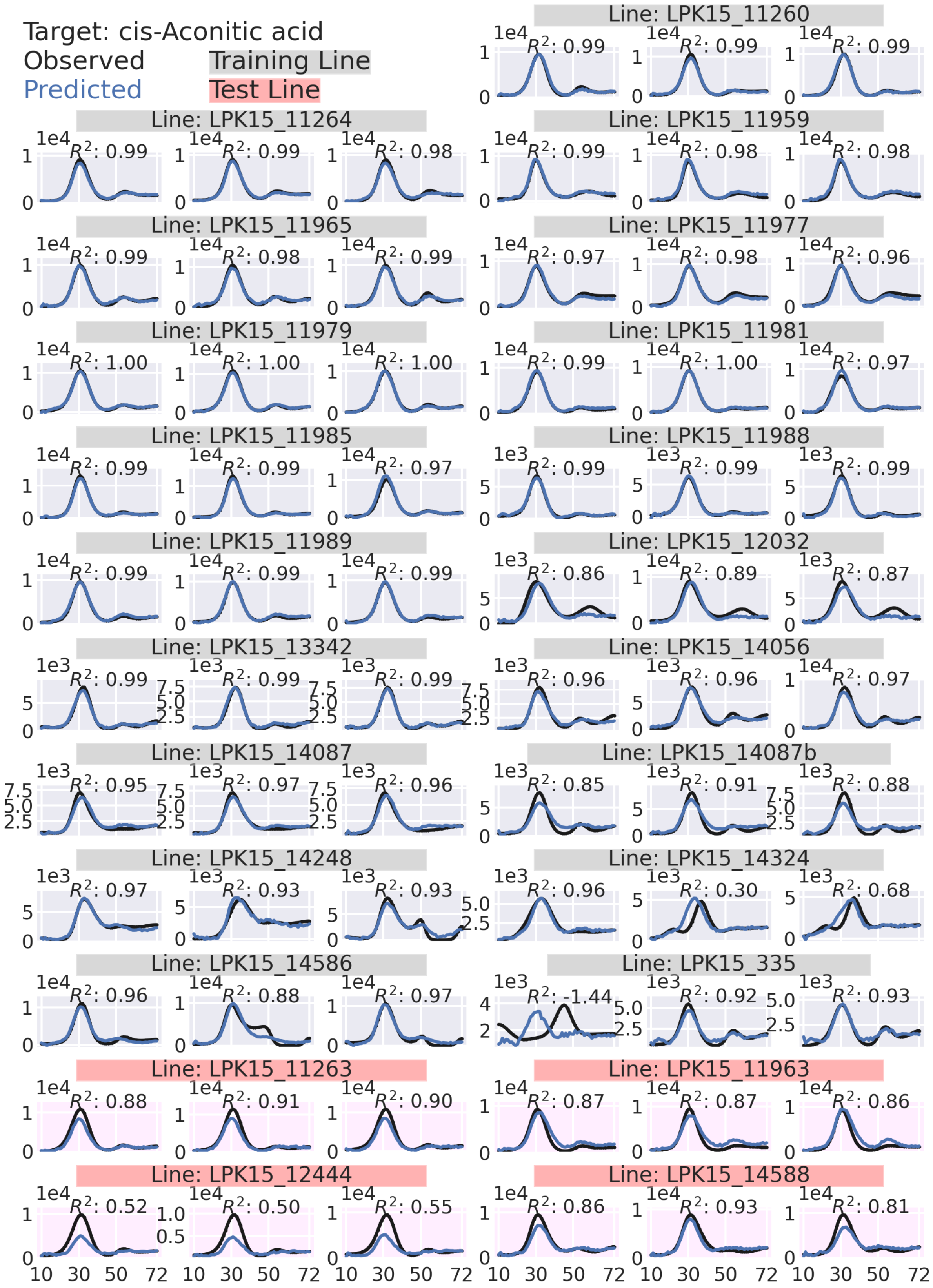
Target: cis-Aconitic acid
Observed
Predicted
Training Line
Test Line
Line: LPK15_11260
Line: LPK15_11264
Line: LPK15_11959
Line: LPK15_11965
Line: LPK15_11977
Line: LPK15_11979
Line: LPK15_11981
Line: LPK15_11985
Line: LPK15_11988
Line: LPK15_11989
Line: LPK15_12032
Line: LPK15_13342
Line: LPK15_14056
Line: LPK15_14087
Line: LPK15_14087b
Line: LPK15_14248
Line: LPK15_14324
Line: LPK15_14586
Line: LPK15_335
Line: LPK15_11263
Line: LPK15_11963
Line: LPK15_12444
Line: LPK15_14588

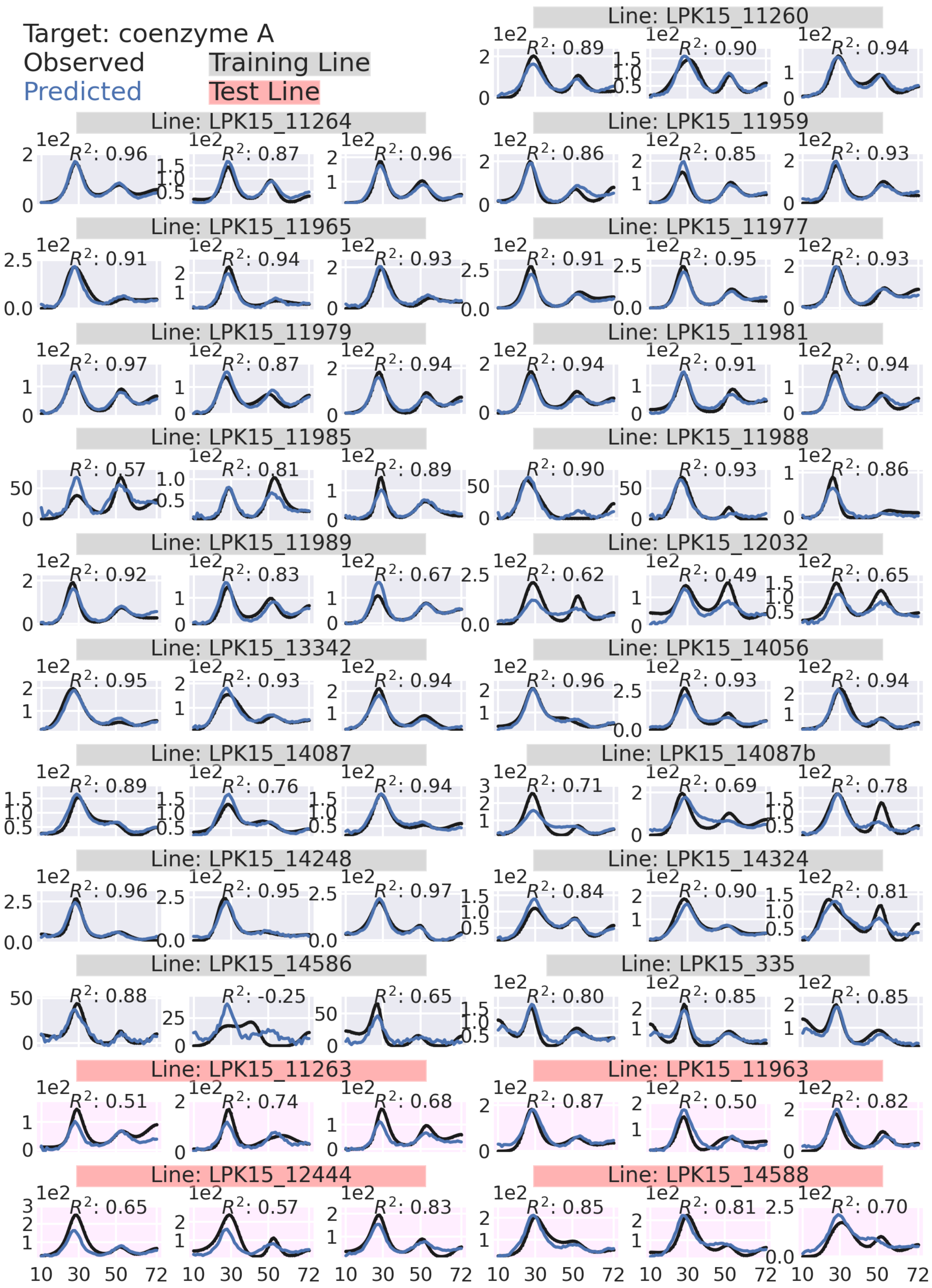

Target: coenzyme A
Observed
Predicted
Training Line
Test Line
Line: LPK15_11260
Line: LPK15_11264
Line: LPK15_11959
Line: LPK15_11965
Line: LPK15_11977
Line: LPK15_11979
Line: LPK15_11981
Line: LPK15_11985
Line: LPK15_11988
Line: LPK15_11989
Line: LPK15_12032
Line: LPK15_13342
Line: LPK15_14056
Line: LPK15_14087
Line: LPK15_14087b
Line: LPK15_14248
Line: LPK15_14324
Line: LPK15_14586
Line: LPK15_335
Line: LPK15_11263
Line: LPK15_11963
Line: LPK15_12444
Line: LPK15_14588

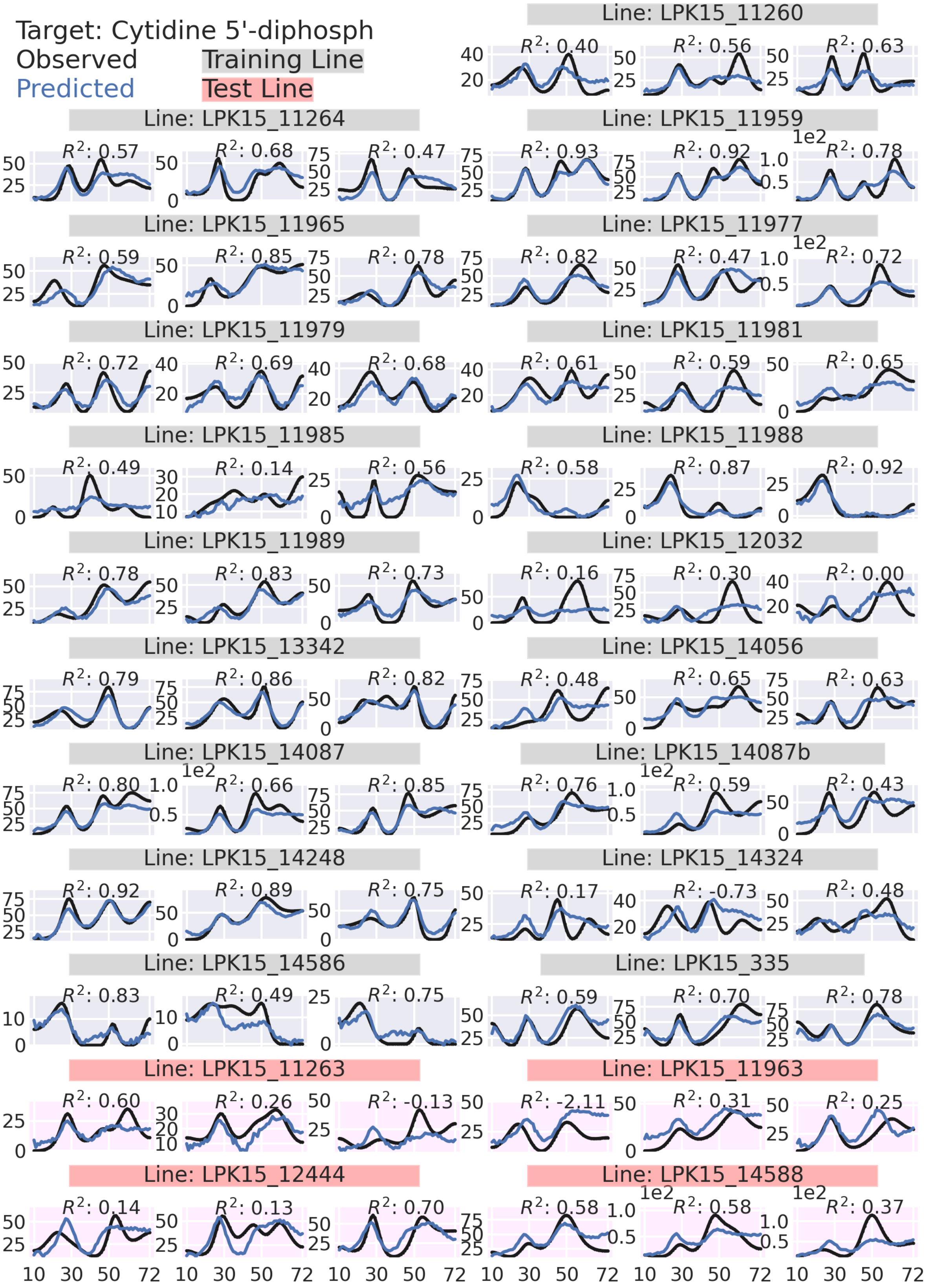

Target: Cytidine 5'-diphosph
Observed
Predicted
Training Line
Test Line
Line: LPK15_11260
R²: 0.40 R²: 0.56 R²: 0.63
Line: LPK15_11264
R²: 0.57 R²: 0.68 R²: 0.47
Line: LPK15_11959
R²: 0.93 R²: 0.92 R²: 0.78
Line: LPK15_11965
R²: 0.59 R²: 0.85 R²: 0.78
Line: LPK15_11977
R²: 0.82 R²: 0.47 R²: 0.72
Line: LPK15_11979
R²: 0.72 R²: 0.69 R²: 0.68
Line: LPK15_11981
R²: 0.61 R²: 0.59 R²: 0.65
Line: LPK15_11985
R²: 0.49 R²: 0.14 R²: 0.56
Line: LPK15_11988
R²: 0.58 R²: 0.87 R²: 0.92
Line: LPK15_11989
R²: 0.78 R²: 0.83 R²: 0.73
Line: LPK15_12032
R²: 0.16 R²: 0.30 R²: 0.00
Line: LPK15_13342
R²: 0.79 R²: 0.86 R²: 0.82
Line: LPK15_14056
R²: 0.48 R²: 0.65 R²: 0.63
Line: LPK15_14087
R²: 0.80 R²: 0.66 R²: 0.85
Line: LPK15_14087b
R²: 0.76 R²: 0.59 R²: 0.43
Line: LPK15_14248
R²: 0.92 R²: 0.89 R²: 0.75
Line: LPK15_14324
R²: 0.17 R²: -0.73 R²: 0.48
Line: LPK15_14586
R²: 0.83 R²: 0.49 R²: 0.75
Line: LPK15_335
R²: 0.59 R²: 0.70 R²: 0.78
Line: LPK15_11263
R²: 0.60 R²: 0.26 R²: -0.13
Line: LPK15_11963
R²: -2.11 R²: 0.31 R²: 0.25
Line: LPK15_12444
R²: 0.14 R²: 0.13 R²: 0.70
Line: LPK15_14588
R²: 0.58 R²: 0.58 R²: 0.37
1e2
10 30 50 72

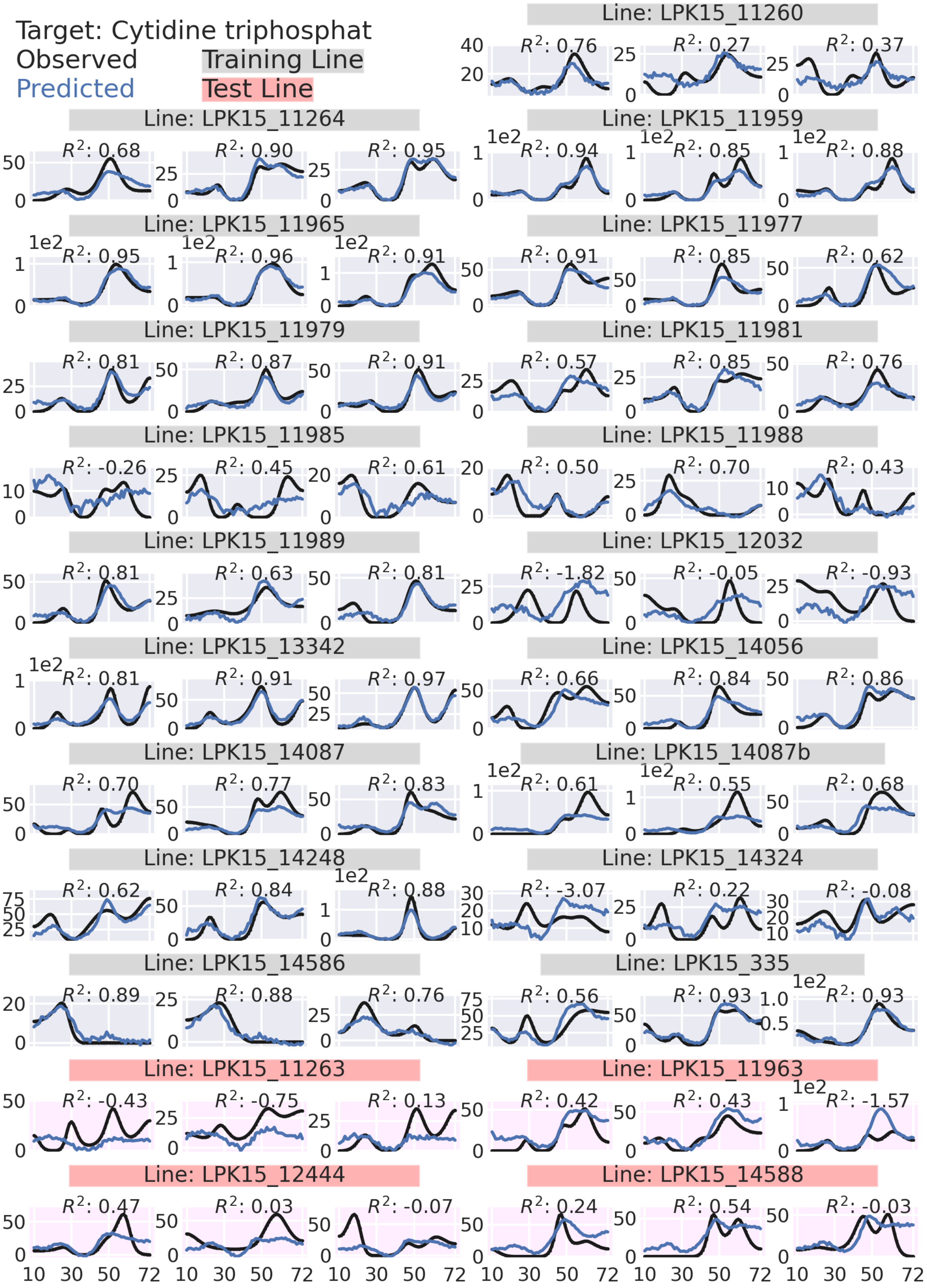

Target: Cytidine triphosphat
Observed
Predicted
Training Line
Test Line
Line: LPK15_11260
Line: LPK15_11264
Line: LPK15_11959
Line: LPK15_11965
Line: LPK15_11977
Line: LPK15_11979
Line: LPK15_11981
Line: LPK15_11985
Line: LPK15_11988
Line: LPK15_11989
Line: LPK15_12032
Line: LPK15_13342
Line: LPK15_14056
Line: LPK15_14087
Line: LPK15_14087b
Line: LPK15_14248
Line: LPK15_14324
Line: LPK15_14586
Line: LPK15_335
Line: LPK15_11263
Line: LPK15_11963
Line: LPK15_12444
Line: LPK15_14588
10 30 50 72

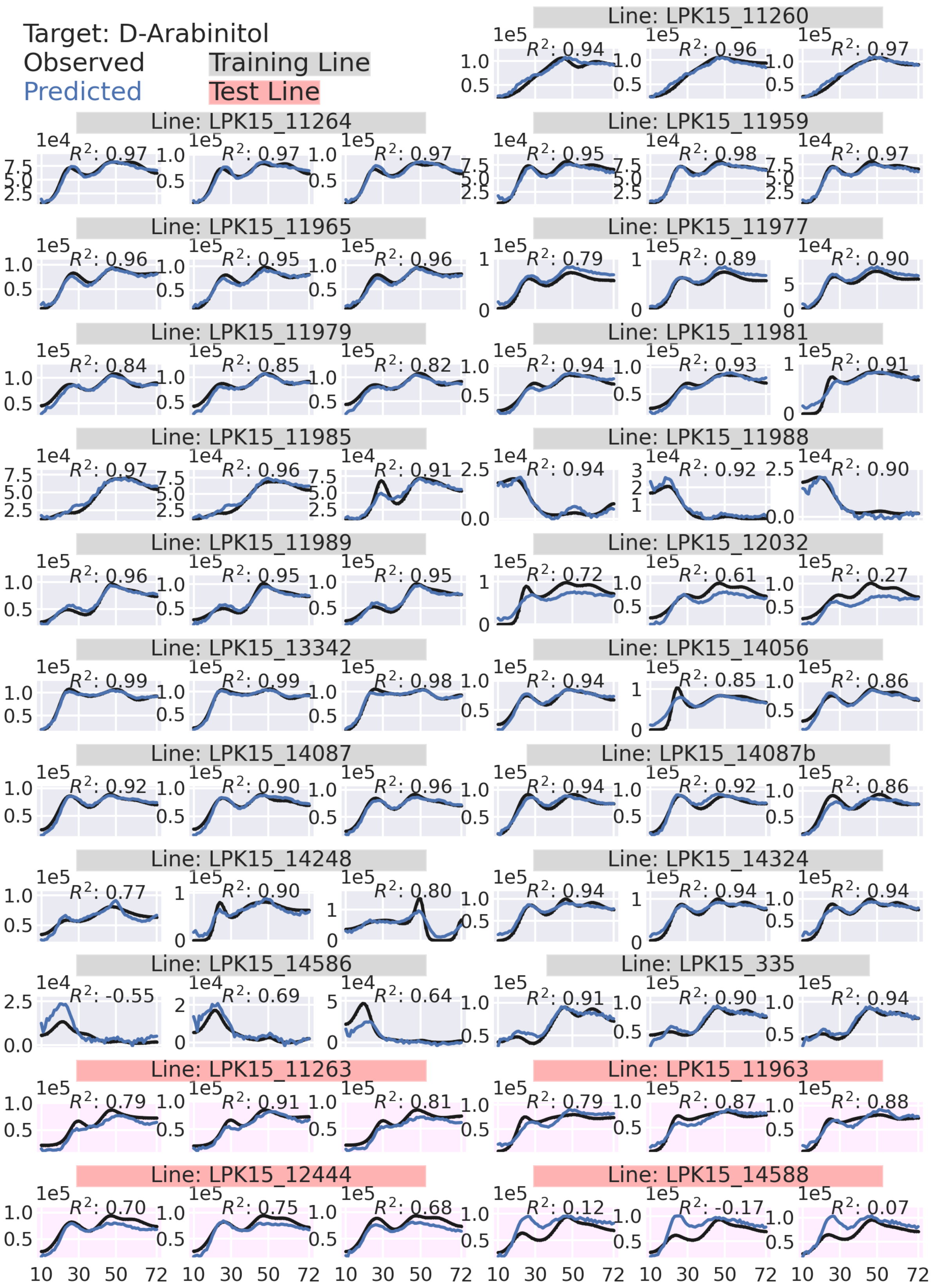

Target: D-Arabinitol
Observed
Predicted
Training Line
Test Line
Line: LPK15_11260
$R^2$: 0.94 $R^2$: 0.96 $R^2$: 0.97
Line: LPK15_11264
$R^2$: 0.97 $R^2$: 0.97 $R^2$: 0.97
Line: LPK15_11959
$R^2$: 0.95 $R^2$: 0.98 $R^2$: 0.97
Line: LPK15_11965
$R^2$: 0.96 $R^2$: 0.95 $R^2$: 0.96
Line: LPK15_11977
$R^2$: 0.79 $R^2$: 0.89 $R^2$: 0.90
Line: LPK15_11979
$R^2$: 0.84 $R^2$: 0.85 $R^2$: 0.82
Line: LPK15_11981
$R^2$: 0.94 $R^2$: 0.93 $R^2$: 0.91
Line: LPK15_11985
$R^2$: 0.97 $R^2$: 0.96 $R^2$: 0.91
Line: LPK15_11988
$R^2$: 0.94 $R^2$: 0.92 $R^2$: 0.90
Line: LPK15_11989
$R^2$: 0.96 $R^2$: 0.95 $R^2$: 0.95
Line: LPK15_12032
$R^2$: 0.72 $R^2$: 0.61 $R^2$: 0.27
Line: LPK15_13342
$R^2$: 0.99 $R^2$: 0.99 $R^2$: 0.98
Line: LPK15_14056
$R^2$: 0.94 $R^2$: 0.85 $R^2$: 0.86
Line: LPK15_14087
$R^2$: 0.92 $R^2$: 0.90 $R^2$: 0.96
Line: LPK15_14087b
$R^2$: 0.94 $R^2$: 0.92 $R^2$: 0.86
Line: LPK15_14248
$R^2$: 0.77 $R^2$: 0.90 $R^2$: 0.80
Line: LPK15_14324
$R^2$: 0.94 $R^2$: 0.94 $R^2$: 0.94
Line: LPK15_14586
$R^2$: -0.55 $R^2$: 0.69 $R^2$: 0.64
Line: LPK15_335
$R^2$: 0.91 $R^2$: 0.90 $R^2$: 0.94
Line: LPK15_11263
$R^2$: 0.79 $R^2$: 0.91 $R^2$: 0.81
Line: LPK15_11963
$R^2$: 0.79 $R^2$: 0.87 $R^2$: 0.88
Line: LPK15_12444
$R^2$: 0.70 $R^2$: 0.75 $R^2$: 0.68
Line: LPK15_14588
$R^2$: 0.12 $R^2$: -0.17 $R^2$: 0.07
10 30 50 72

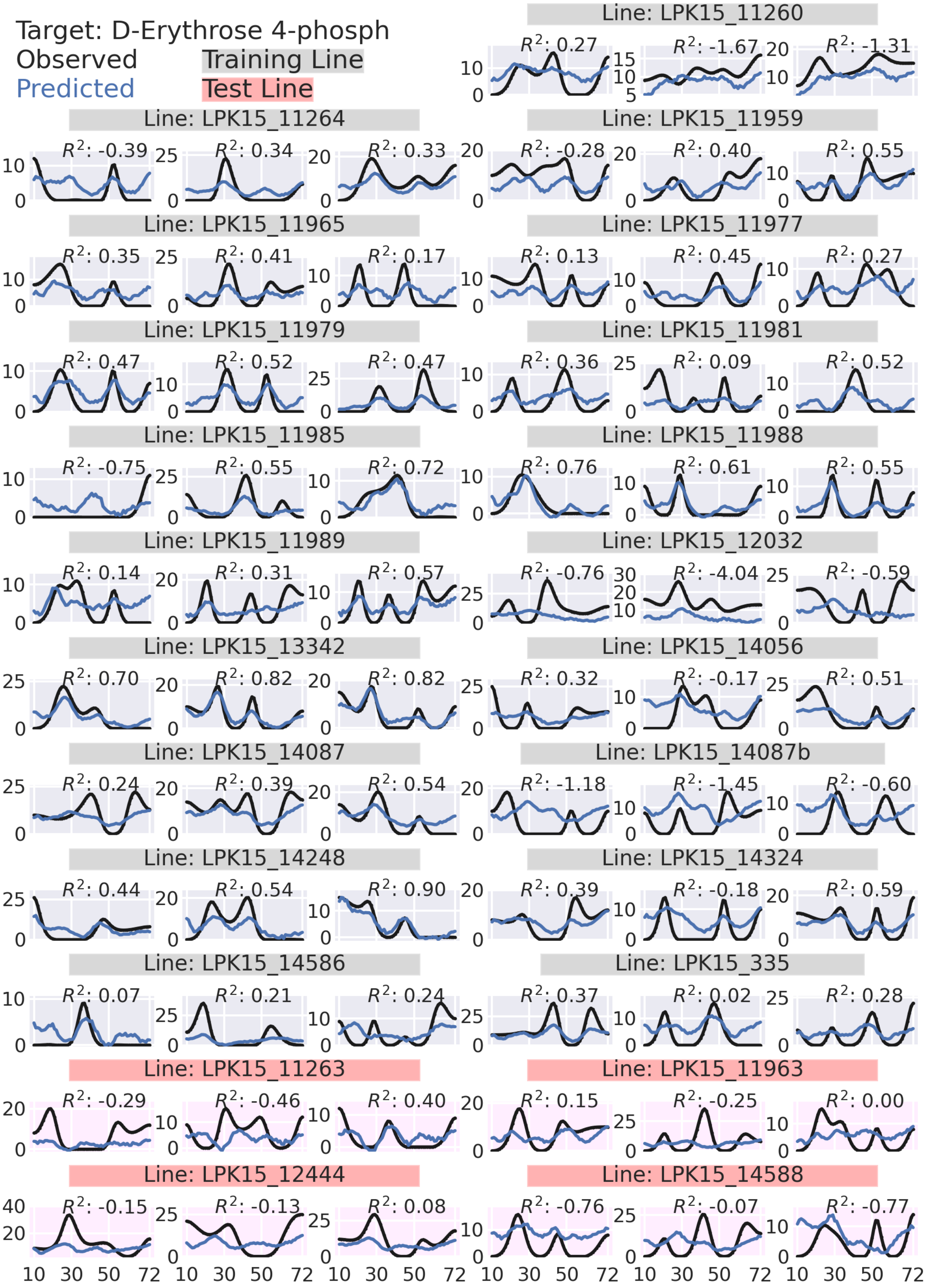

Target: D-Erythrose 4-phosph
Observed
Predicted
Training Line
Test Line
Line: LPK15_11260
R²: 0.27
R²: -1.67
R²: -1.31
Line: LPK15_11264
R²: -0.39
R²: 0.34
R²: 0.33
Line: LPK15_11959
R²: -0.28
R²: 0.40
R²: 0.55
Line: LPK15_11965
R²: 0.35
R²: 0.41
R²: 0.17
Line: LPK15_11977
R²: 0.13
R²: 0.45
R²: 0.27
Line: LPK15_11979
R²: 0.47
R²: 0.52
R²: 0.47
Line: LPK15_11981
R²: 0.36
R²: 0.09
R²: 0.52
Line: LPK15_11985
R²: -0.75
R²: 0.55
R²: 0.72
Line: LPK15_11988
R²: 0.76
R²: 0.61
R²: 0.55
Line: LPK15_11989
R²: 0.14
R²: 0.31
R²: 0.57
Line: LPK15_12032
R²: -0.76
R²: -4.04
R²: -0.59
Line: LPK15_13342
R²: 0.70
R²: 0.82
R²: 0.82
Line: LPK15_14056
R²: 0.32
R²: -0.17
R²: 0.51
Line: LPK15_14087
R²: 0.24
R²: 0.39
R²: 0.54
Line: LPK15_14087b
R²: -1.18
R²: -1.45
R²: -0.60
Line: LPK15_14248
R²: 0.44
R²: 0.54
R²: 0.90
Line: LPK15_14324
R²: 0.39
R²: -0.18
R²: 0.59
Line: LPK15_14586
R²: 0.07
R²: 0.21
R²: 0.24
Line: LPK15_335
R²: 0.37
R²: 0.02
R²: 0.28
Line: LPK15_11263
R²: -0.29
R²: -0.46
R²: 0.40
Line: LPK15_11963
R²: 0.15
R²: -0.25
R²: 0.00
Line: LPK15_12444
R²: -0.15
R²: -0.13
R²: 0.08
Line: LPK15_14588
R²: -0.76
R²: -0.07
R²: -0.77
10 30 50 72

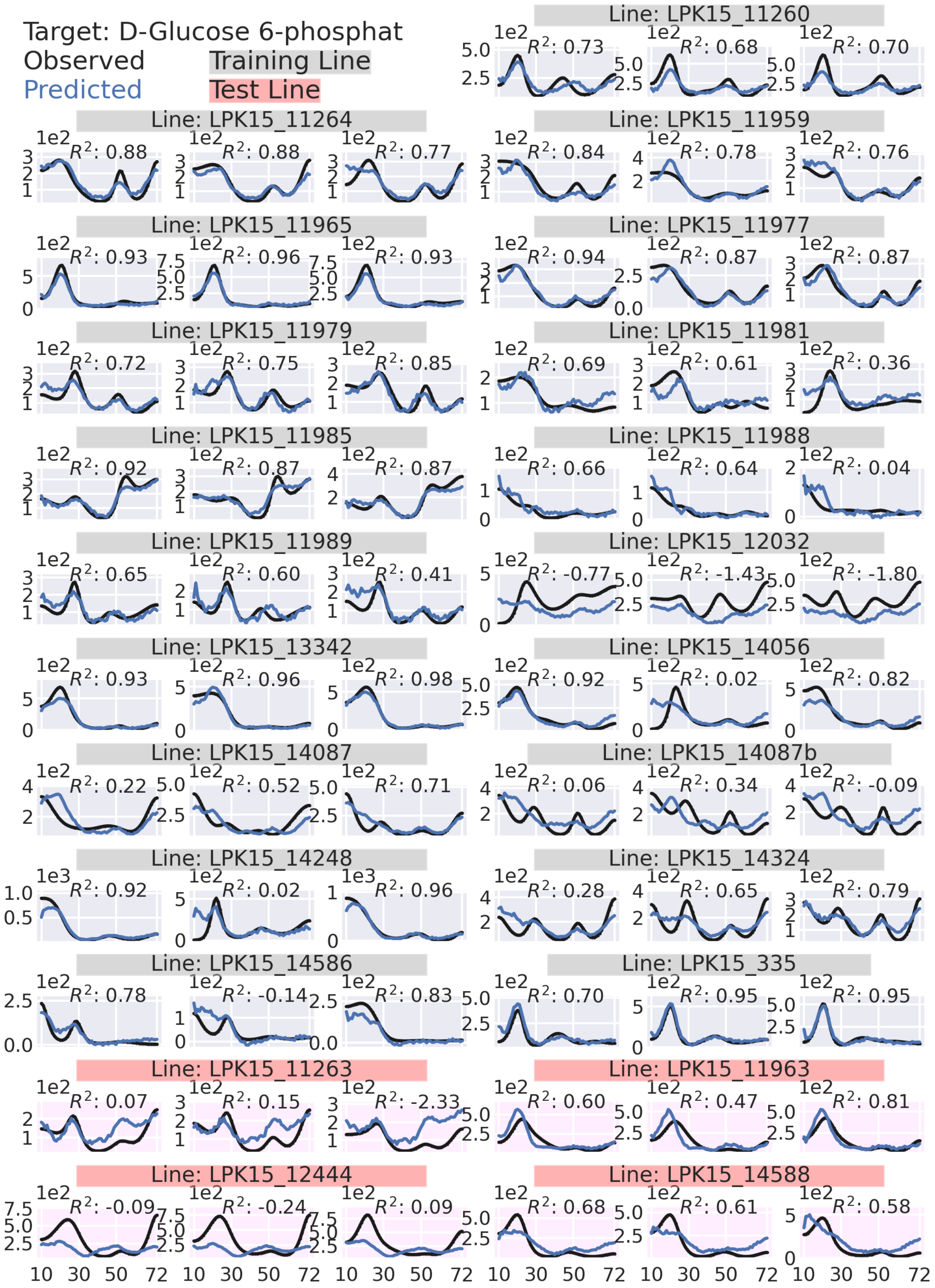

Target: D-Glucose 6-phosphat
Observed
Predicted
Training Line
Test Line
Line: LPK15_11260
R²: 0.73
R²: 0.68
R²: 0.70
Line: LPK15_11264
R²: 0.88
R²: 0.88
R²: 0.77
Line: LPK15_11959
R²: 0.84
R²: 0.78
R²: 0.76
Line: LPK15_11965
R²: 0.93
R²: 0.96
R²: 0.93
Line: LPK15_11977
R²: 0.94
R²: 0.87
R²: 0.87
Line: LPK15_11979
R²: 0.72
R²: 0.75
R²: 0.85
Line: LPK15_11981
R²: 0.69
R²: 0.61
R²: 0.36
Line: LPK15_11985
R²: 0.92
R²: 0.87
R²: 0.87
Line: LPK15_11988
R²: 0.66
R²: 0.64
R²: 0.04
Line: LPK15_11989
R²: 0.65
R²: 0.60
R²: 0.41
Line: LPK15_12032
R²: -0.77
R²: -1.43
R²: -1.80
Line: LPK15_13342
R²: 0.93
R²: 0.96
R²: 0.98
Line: LPK15_14056
R²: 0.92
R²: 0.02
R²: 0.82
Line: LPK15_14087
R²: 0.22
R²: 0.52
R²: 0.71
Line: LPK15_14087b
R²: 0.06
R²: 0.34
R²: -0.09
Line: LPK15_14248
R²: 0.92
R²: 0.02
R²: 0.96
Line: LPK15_14324
R²: 0.28
R²: 0.65
R²: 0.79
Line: LPK15_14586
R²: 0.78
R²: -0.14
R²: 0.83
Line: LPK15_335
R²: 0.70
R²: 0.95
R²: 0.95
Line: LPK15_11263
R²: 0.07
R²: 0.15
R²: -2.33
Line: LPK15_11963
R²: 0.60
R²: 0.47
R²: 0.81
Line: LPK15_12444
R²: -0.09
R²: -0.24
R²: 0.09
Line: LPK15_14588
R²: 0.68
R²: 0.61
R²: 0.58
10 30 50 72

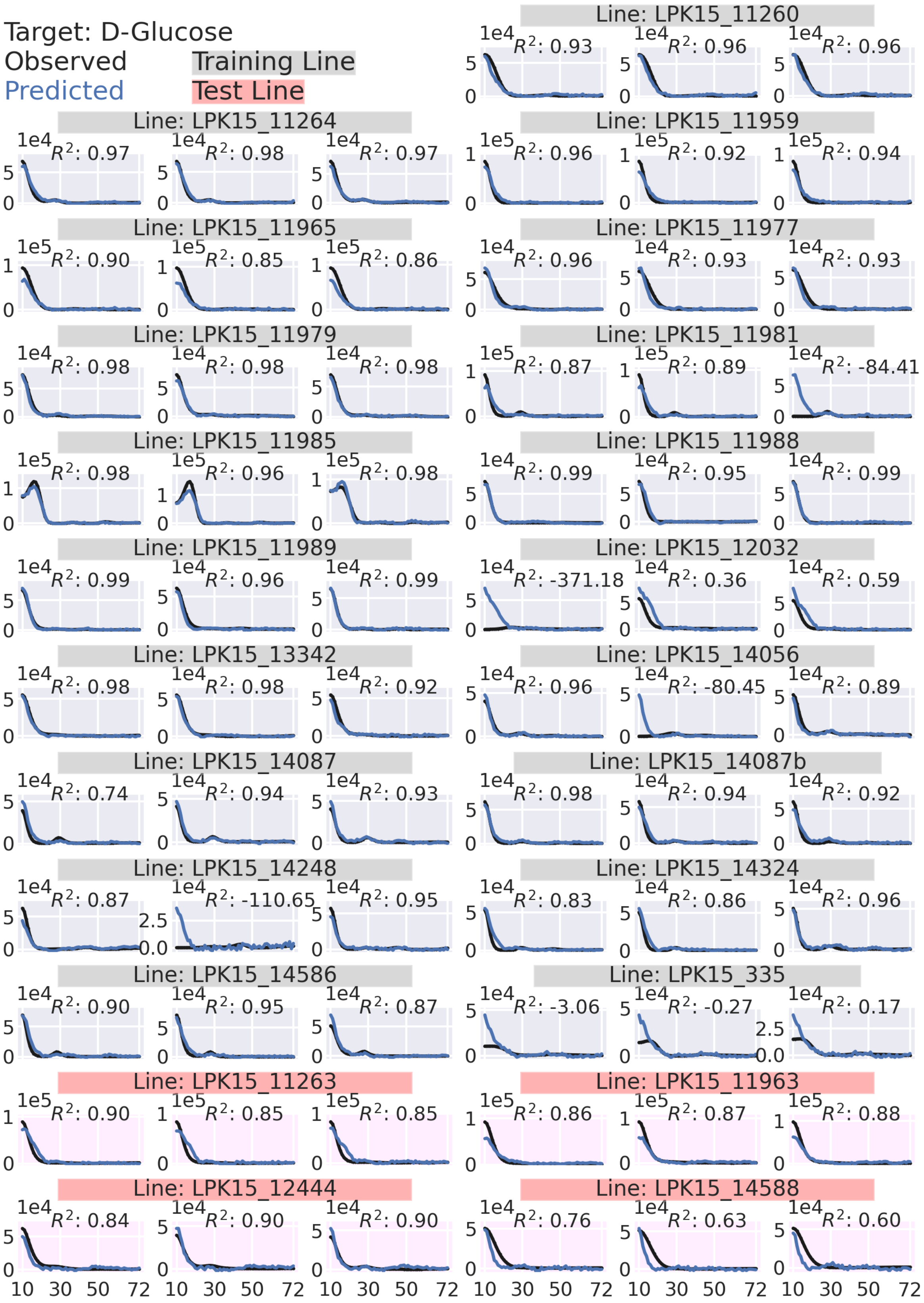

Target: D-Glucose
Observed
Predicted
Training Line
Test Line
Line: LPK15_11260
$R^2$: 0.93
$R^2$: 0.96
$R^2$: 0.96
Line: LPK15_11264
$R^2$: 0.97
$R^2$: 0.98
$R^2$: 0.97
Line: LPK15_11959
$R^2$: 0.96
$R^2$: 0.92
$R^2$: 0.94
Line: LPK15_11965
$R^2$: 0.90
$R^2$: 0.85
$R^2$: 0.86
Line: LPK15_11977
$R^2$: 0.96
$R^2$: 0.93
$R^2$: 0.93
Line: LPK15_11979
$R^2$: 0.98
$R^2$: 0.98
$R^2$: 0.98
Line: LPK15_11981
$R^2$: 0.87
$R^2$: 0.89
$R^2$: -84.41
Line: LPK15_11985
$R^2$: 0.98
$R^2$: 0.96
$R^2$: 0.98
Line: LPK15_11988
$R^2$: 0.99
$R^2$: 0.95
$R^2$: 0.99
Line: LPK15_11989
$R^2$: 0.99
$R^2$: 0.96
$R^2$: 0.99
Line: LPK15_12032
$R^2$: -371.18
$R^2$: 0.36
$R^2$: 0.59
Line: LPK15_13342
$R^2$: 0.98
$R^2$: 0.98
$R^2$: 0.92
Line: LPK15_14056
$R^2$: 0.96
$R^2$: -80.45
$R^2$: 0.89
Line: LPK15_14087
$R^2$: 0.74
$R^2$: 0.94
$R^2$: 0.93
Line: LPK15_14087b
$R^2$: 0.98
$R^2$: 0.94
$R^2$: 0.92
Line: LPK15_14248
$R^2$: 0.87
$R^2$: -110.65
$R^2$: 0.95
Line: LPK15_14324
$R^2$: 0.83
$R^2$: 0.86
$R^2$: 0.96
Line: LPK15_14586
$R^2$: 0.90
$R^2$: 0.95
$R^2$: 0.87
Line: LPK15_335
$R^2$: -3.06
$R^2$: -0.27
$R^2$: 0.17
Line: LPK15_11263
$R^2$: 0.90
$R^2$: 0.85
$R^2$: 0.85
Line: LPK15_11963
$R^2$: 0.86
$R^2$: 0.87
$R^2$: 0.88
Line: LPK15_12444
$R^2$: 0.84
$R^2$: 0.90
$R^2$: 0.90
Line: LPK15_14588
$R^2$: 0.76
$R^2$: 0.63
$R^2$: 0.60
10 30 50 72

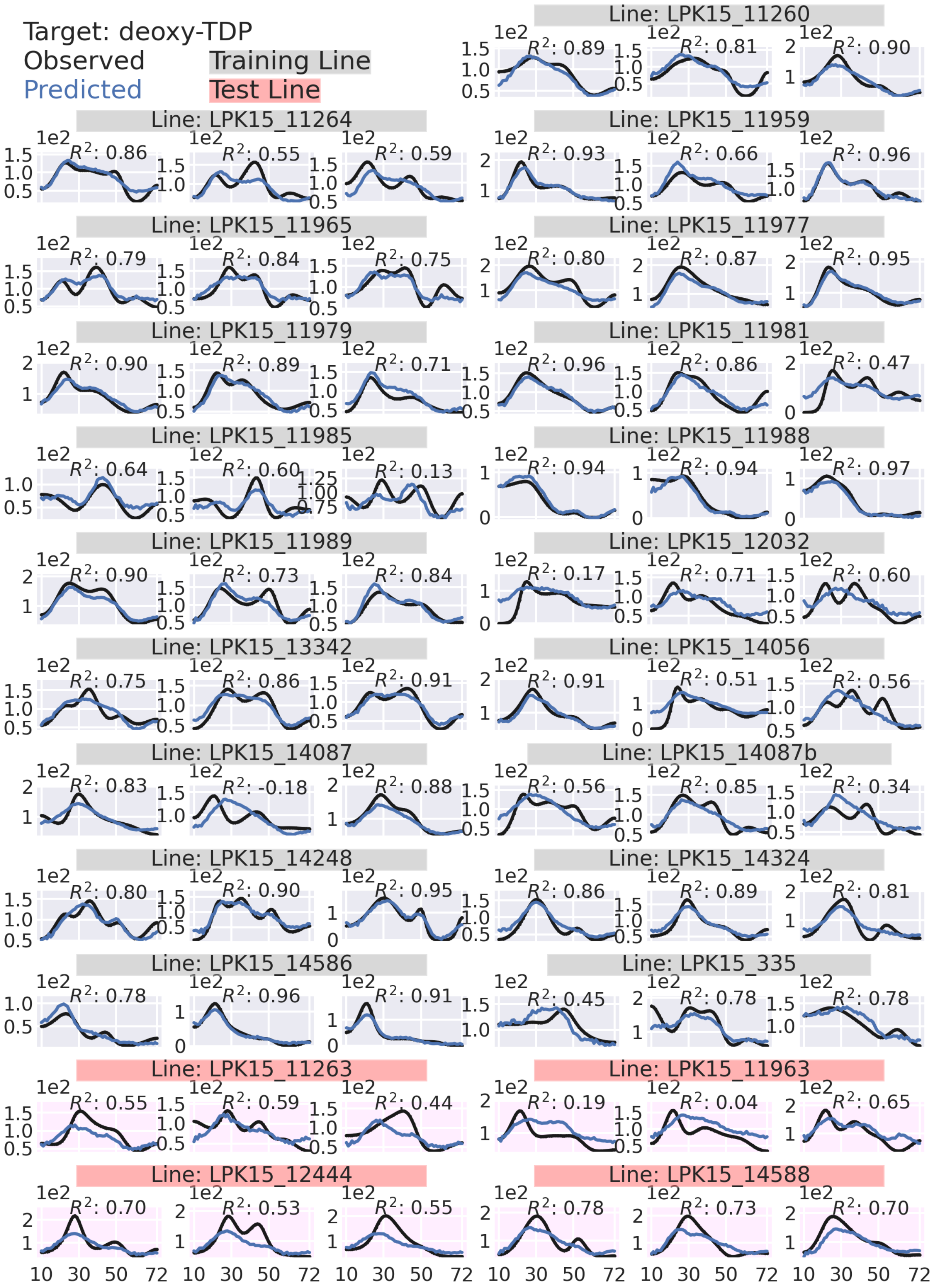
Target: deoxy-TDP
Observed
Predicted
Training Line
Test Line
Line: LPK15_11260
R²: 0.89 R²: 0.81 R²: 0.90
Line: LPK15_11264
R²: 0.86 R²: 0.55 R²: 0.59
Line: LPK15_11959
R²: 0.93 R²: 0.66 R²: 0.96
Line: LPK15_11965
R²: 0.79 R²: 0.84 R²: 0.75
Line: LPK15_11977
R²: 0.80 R²: 0.87 R²: 0.95
Line: LPK15_11979
R²: 0.90 R²: 0.89 R²: 0.71
Line: LPK15_11981
R²: 0.96 R²: 0.86 R²: 0.47
Line: LPK15_11985
R²: 0.64 R²: 0.60 R²: 0.13
Line: LPK15_11988
R²: 0.94 R²: 0.94 R²: 0.97
Line: LPK15_11989
R²: 0.90 R²: 0.73 R²: 0.84
Line: LPK15_12032
R²: 0.17 R²: 0.71 R²: 0.60
Line: LPK15_13342
R²: 0.75 R²: 0.86 R²: 0.91
Line: LPK15_14056
R²: 0.91 R²: 0.51 R²: 0.56
Line: LPK15_14087
R²: 0.83 R²: -0.18 R²: 0.88
Line: LPK15_14087b
R²: 0.56 R²: 0.85 R²: 0.34
Line: LPK15_14248
R²: 0.80 R²: 0.90 R²: 0.95
Line: LPK15_14324
R²: 0.86 R²: 0.89 R²: 0.81
Line: LPK15_14586
R²: 0.78 R²: 0.96 R²: 0.91
Line: LPK15_335
R²: 0.45 R²: 0.78 R²: 0.78
Line: LPK15_11263
R²: 0.55 R²: 0.59 R²: 0.44
Line: LPK15_11963
R²: 0.19 R²: 0.04 R²: 0.65
Line: LPK15_12444
R²: 0.70 R²: 0.53 R²: 0.55
Line: LPK15_14588
R²: 0.78 R²: 0.73 R²: 0.70
1e2
10 30 50 72

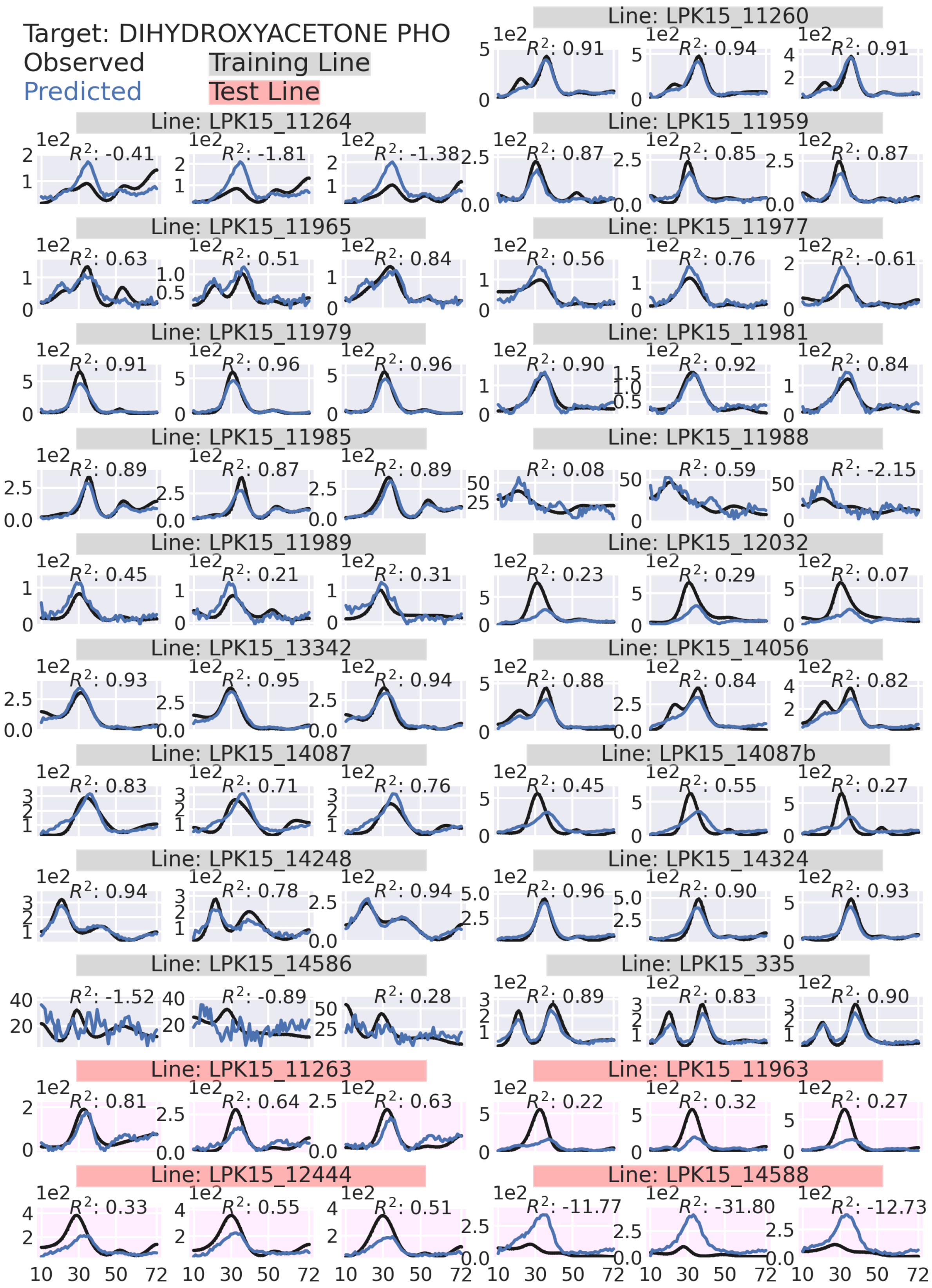

Target: DIHYDROXYACETONE PHO
Observed
Predicted
Training Line
Test Line
Line: LPK15_11260
Line: LPK15_11264
Line: LPK15_11959
Line: LPK15_11965
Line: LPK15_11977
Line: LPK15_11979
Line: LPK15_11981
Line: LPK15_11985
Line: LPK15_11988
Line: LPK15_11989
Line: LPK15_12032
Line: LPK15_13342
Line: LPK15_14056
Line: LPK15_14087
Line: LPK15_14087b
Line: LPK15_14248
Line: LPK15_14324
Line: LPK15_14586
Line: LPK15_335
Line: LPK15_11263
Line: LPK15_11963
Line: LPK15_12444
Line: LPK15_14588

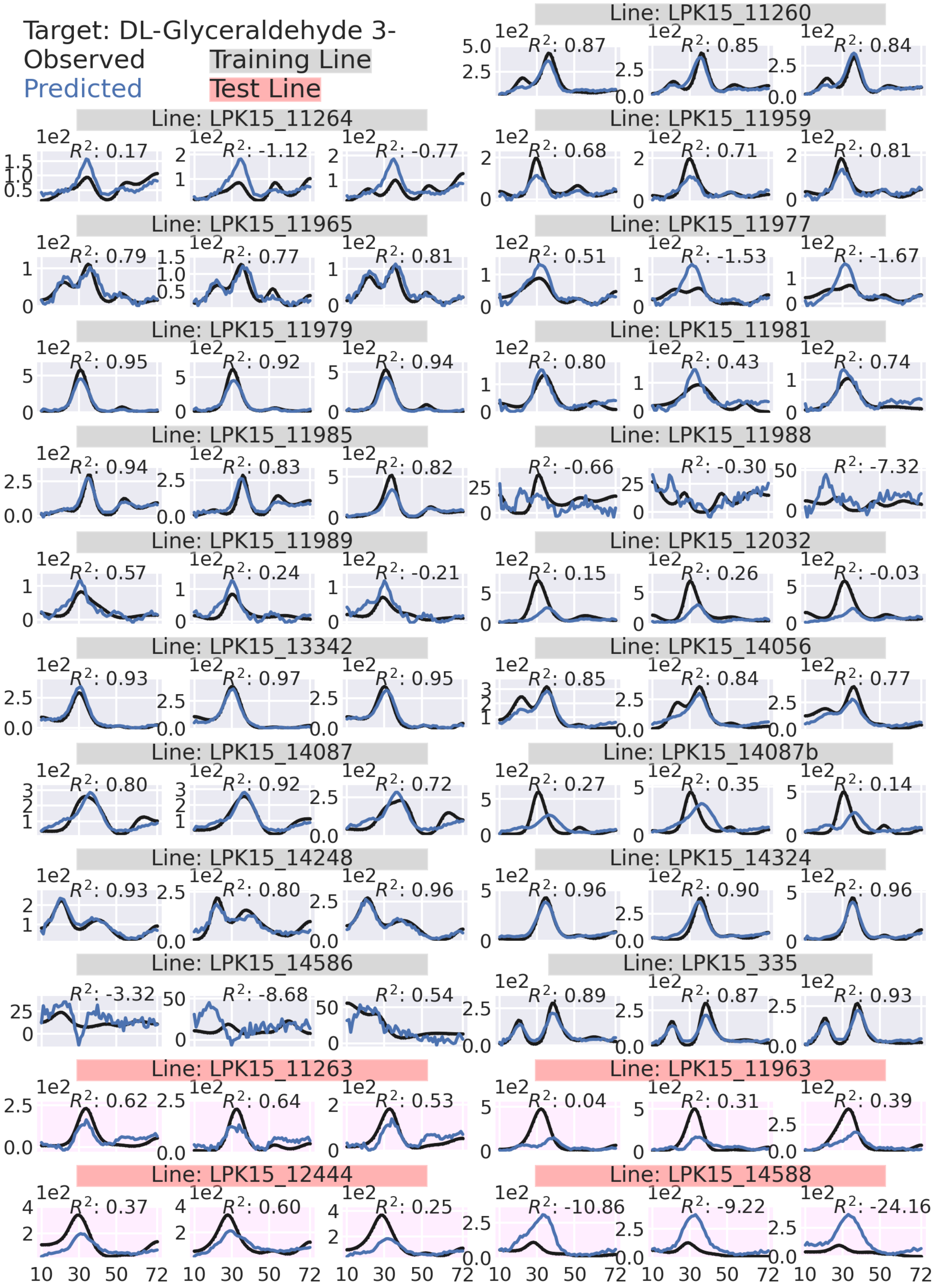
Target: DL-Glyceraldehyde 3-
Observed
Predicted
Training Line
Test Line
Line: LPK15_11260
Line: LPK15_11264
Line: LPK15_11959
Line: LPK15_11965
Line: LPK15_11977
Line: LPK15_11979
Line: LPK15_11981
Line: LPK15_11985
Line: LPK15_11988
Line: LPK15_11989
Line: LPK15_12032
Line: LPK15_13342
Line: LPK15_14056
Line: LPK15_14087
Line: LPK15_14087b
Line: LPK15_14248
Line: LPK15_14324
Line: LPK15_14586
Line: LPK15_335
Line: LPK15_11263
Line: LPK15_11963
Line: LPK15_12444
Line: LPK15_14588

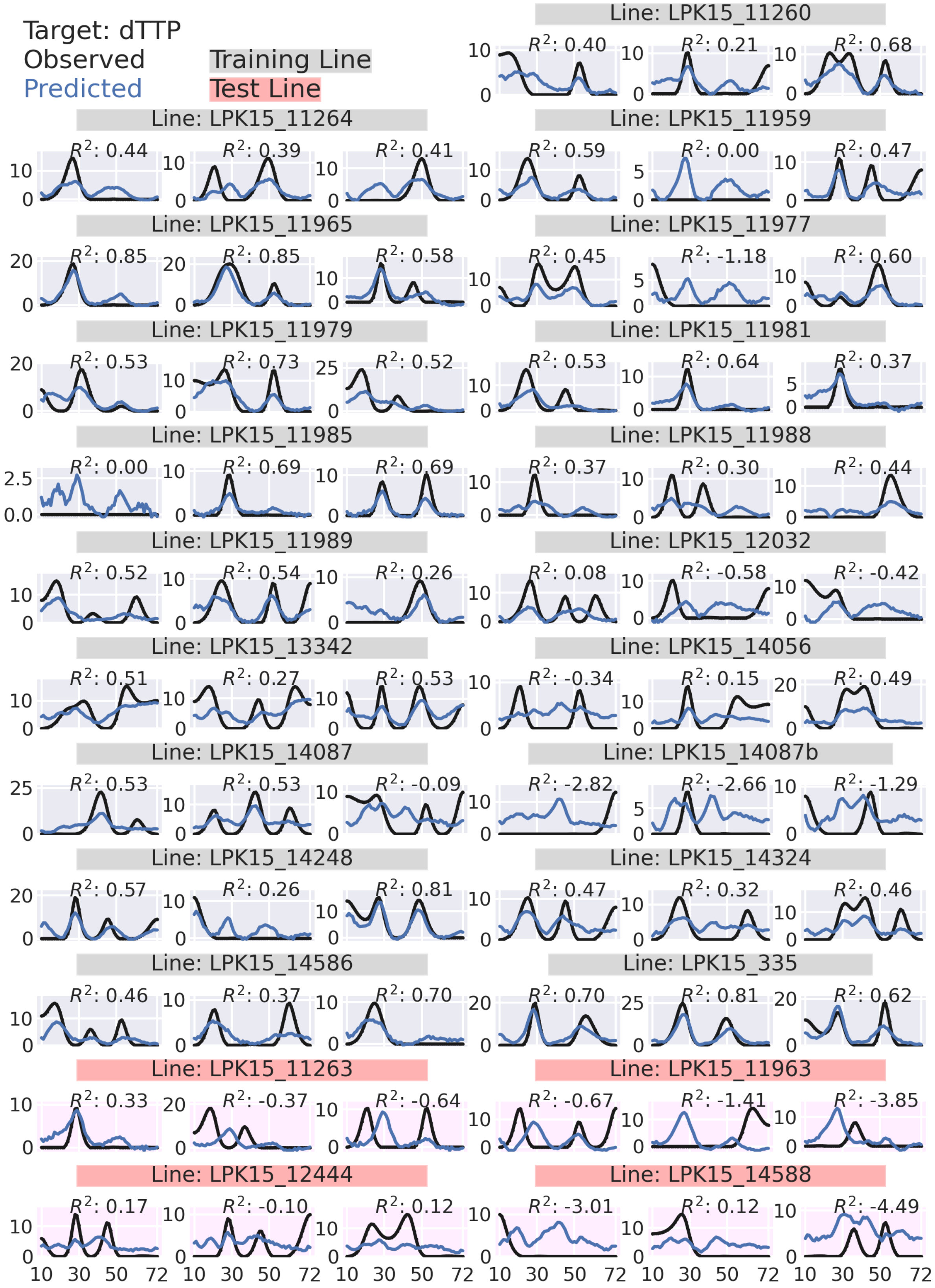

Target: dTTP
Observed
Predicted
Training Line
Test Line
Line: LPK15_11260
Line: LPK15_11264
Line: LPK15_11959
Line: LPK15_11965
Line: LPK15_11977
Line: LPK15_11979
Line: LPK15_11981
Line: LPK15_11985
Line: LPK15_11988
Line: LPK15_11989
Line: LPK15_12032
Line: LPK15_13342
Line: LPK15_14056
Line: LPK15_14087
Line: LPK15_14087b
Line: LPK15_14248
Line: LPK15_14324
Line: LPK15_14586
Line: LPK15_335
Line: LPK15_11263
Line: LPK15_11963
Line: LPK15_12444
Line: LPK15_14588

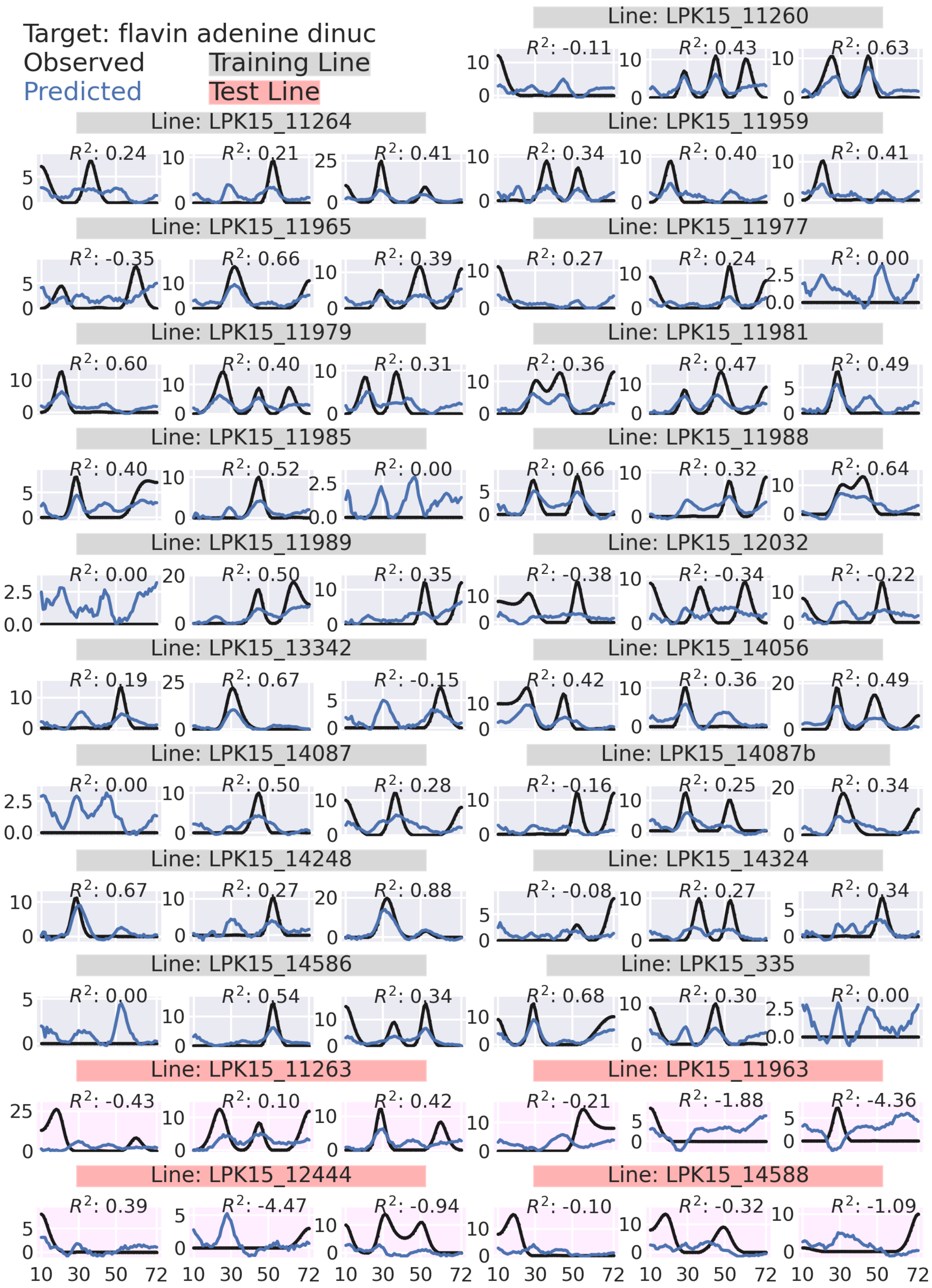
Target: flavin adenine dinuc
Observed
Predicted
Training Line
Test Line
Line: LPK15_11260
$R^2$: -0.11
$R^2$: 0.43
$R^2$: 0.63
Line: LPK15_11264
$R^2$: 0.24
$R^2$: 0.21
$R^2$: 0.41
Line: LPK15_11959
$R^2$: 0.34
$R^2$: 0.40
$R^2$: 0.41
Line: LPK15_11965
$R^2$: -0.35
$R^2$: 0.66
$R^2$: 0.39
Line: LPK15_11977
$R^2$: 0.27
$R^2$: 0.24
$R^2$: 0.00
Line: LPK15_11979
$R^2$: 0.60
$R^2$: 0.40
$R^2$: 0.31
Line: LPK15_11981
$R^2$: 0.36
$R^2$: 0.47
$R^2$: 0.49
Line: LPK15_11985
$R^2$: 0.40
$R^2$: 0.52
$R^2$: 0.00
Line: LPK15_11988
$R^2$: 0.66
$R^2$: 0.32
$R^2$: 0.64
Line: LPK15_11989
$R^2$: 0.00
$R^2$: 0.50
$R^2$: 0.35
Line: LPK15_12032
$R^2$: -0.38
$R^2$: -0.34
$R^2$: -0.22
Line: LPK15_13342
$R^2$: 0.19
$R^2$: 0.67
$R^2$: -0.15
Line: LPK15_14056
$R^2$: 0.42
$R^2$: 0.36
$R^2$: 0.49
Line: LPK15_14087
$R^2$: 0.00
$R^2$: 0.50
$R^2$: 0.28
Line: LPK15_14087b
$R^2$: -0.16
$R^2$: 0.25
$R^2$: 0.34
Line: LPK15_14248
$R^2$: 0.67
$R^2$: 0.27
$R^2$: 0.88
Line: LPK15_14324
$R^2$: -0.08
$R^2$: 0.27
$R^2$: 0.34
Line: LPK15_14586
$R^2$: 0.00
$R^2$: 0.54
$R^2$: 0.34
Line: LPK15_335
$R^2$: 0.68
$R^2$: 0.30
$R^2$: 0.00
Line: LPK15_11263
$R^2$: -0.43
$R^2$: 0.10
$R^2$: 0.42
Line: LPK15_11963
$R^2$: -0.21
$R^2$: -1.88
$R^2$: -4.36
Line: LPK15_12444
$R^2$: 0.39
$R^2$: -4.47
$R^2$: -0.94
Line: LPK15_14588
$R^2$: -0.10
$R^2$: -0.32
$R^2$: -1.09
10 30 50 72

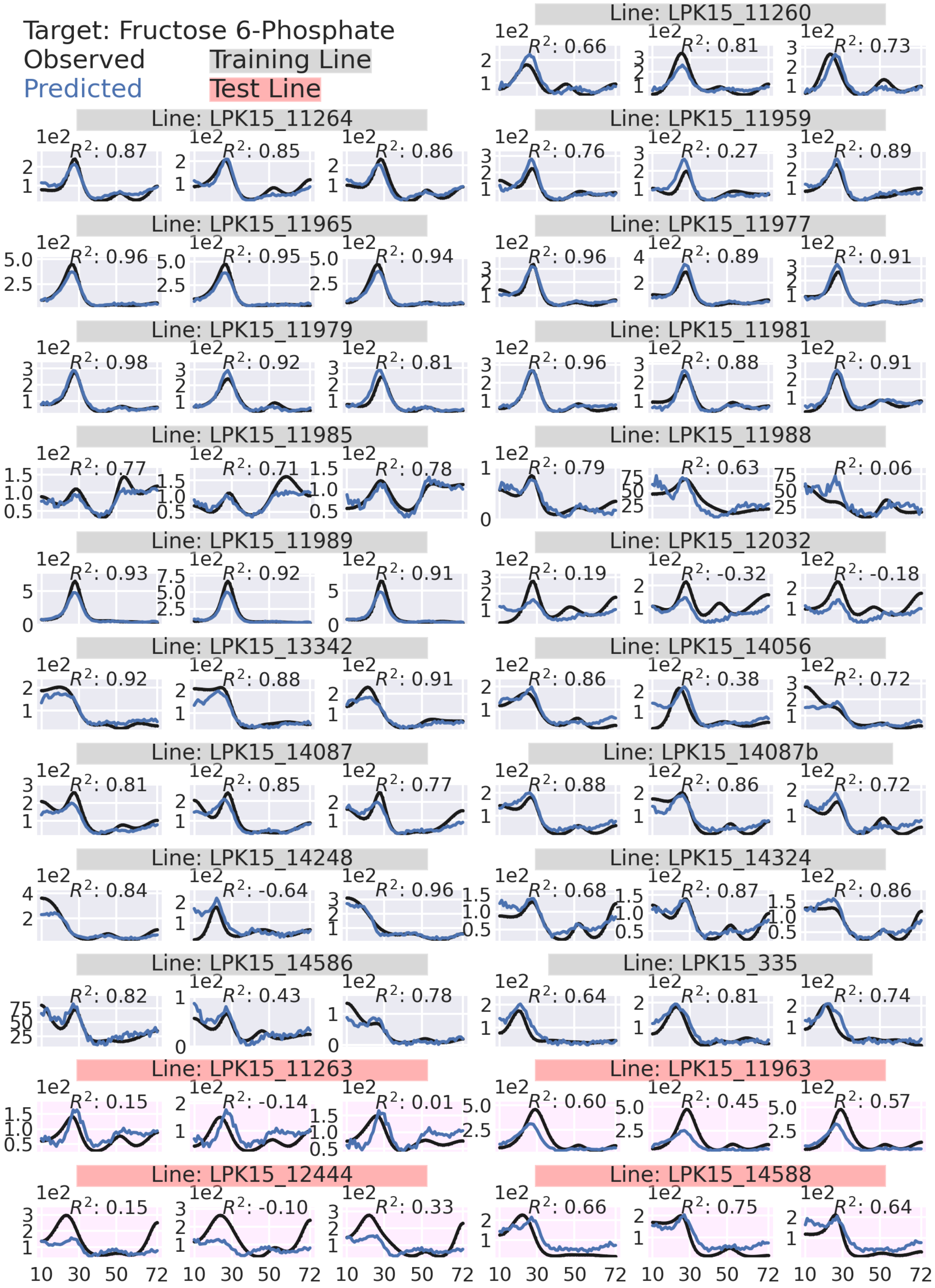
Target: Fructose 6-Phosphate
Observed
Predicted
Training Line
Test Line
Line: LPK15_11260
Line: LPK15_11264
Line: LPK15_11959
Line: LPK15_11965
Line: LPK15_11977
Line: LPK15_11979
Line: LPK15_11981
Line: LPK15_11985
Line: LPK15_11988
Line: LPK15_11989
Line: LPK15_12032
Line: LPK15_13342
Line: LPK15_14056
Line: LPK15_14087
Line: LPK15_14087b
Line: LPK15_14248
Line: LPK15_14324
Line: LPK15_14586
Line: LPK15_335
Line: LPK15_11263
Line: LPK15_11963
Line: LPK15_12444
Line: LPK15_14588
10 30 50 72

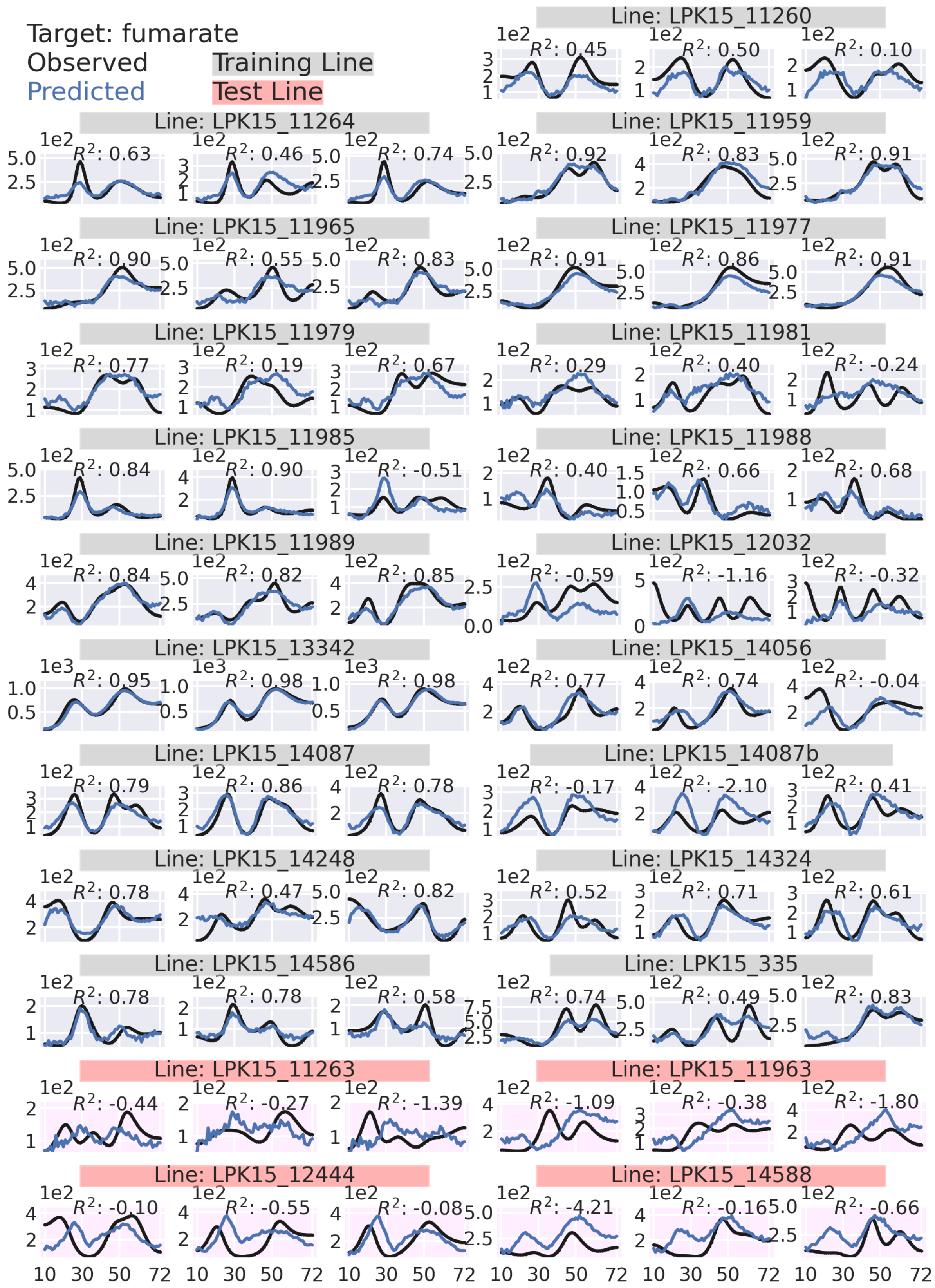

Target: fumarate
Observed
Predicted
Training Line
Test Line
Line: LPK15_11260
$R^2$: 0.45
$R^2$: 0.50
$R^2$: 0.10
Line: LPK15_11264
$R^2$: 0.63
$R^2$: 0.46
$R^2$: 0.74
Line: LPK15_11959
$R^2$: 0.92
$R^2$: 0.83
$R^2$: 0.91
Line: LPK15_11965
$R^2$: 0.90
$R^2$: 0.55
$R^2$: 0.83
Line: LPK15_11977
$R^2$: 0.91
$R^2$: 0.86
$R^2$: 0.91
Line: LPK15_11979
$R^2$: 0.77
$R^2$: 0.19
$R^2$: 0.67
Line: LPK15_11981
$R^2$: 0.29
$R^2$: 0.40
$R^2$: -0.24
Line: LPK15_11985
$R^2$: 0.84
$R^2$: 0.90
$R^2$: -0.51
Line: LPK15_11988
$R^2$: 0.40
$R^2$: 0.66
$R^2$: 0.68
Line: LPK15_11989
$R^2$: 0.84
$R^2$: 0.82
$R^2$: 0.85
Line: LPK15_12032
$R^2$: -0.59
$R^2$: -1.16
$R^2$: -0.32
Line: LPK15_13342
$R^2$: 0.95
$R^2$: 0.98
$R^2$: 0.98
Line: LPK15_14056
$R^2$: 0.77
$R^2$: 0.74
$R^2$: -0.04
Line: LPK15_14087
$R^2$: 0.79
$R^2$: 0.86
$R^2$: 0.78
Line: LPK15_14087b
$R^2$: -0.17
$R^2$: -2.10
$R^2$: 0.41
Line: LPK15_14248
$R^2$: 0.78
$R^2$: 0.47
$R^2$: 0.82
Line: LPK15_14324
$R^2$: 0.52
$R^2$: 0.71
$R^2$: 0.61
Line: LPK15_14586
$R^2$: 0.78
$R^2$: 0.78
$R^2$: 0.58
Line: LPK15_335
$R^2$: 0.74
$R^2$: 0.49
$R^2$: 0.83
Line: LPK15_11263
$R^2$: -0.44
$R^2$: -0.27
$R^2$: -1.39
Line: LPK15_11963
$R^2$: -1.09
$R^2$: -0.38
$R^2$: -1.80
Line: LPK15_12444
$R^2$: -0.10
$R^2$: -0.55
$R^2$: -0.08
Line: LPK15_14588
$R^2$: -4.21
$R^2$: -0.16
$R^2$: -0.66
10 30 50 72

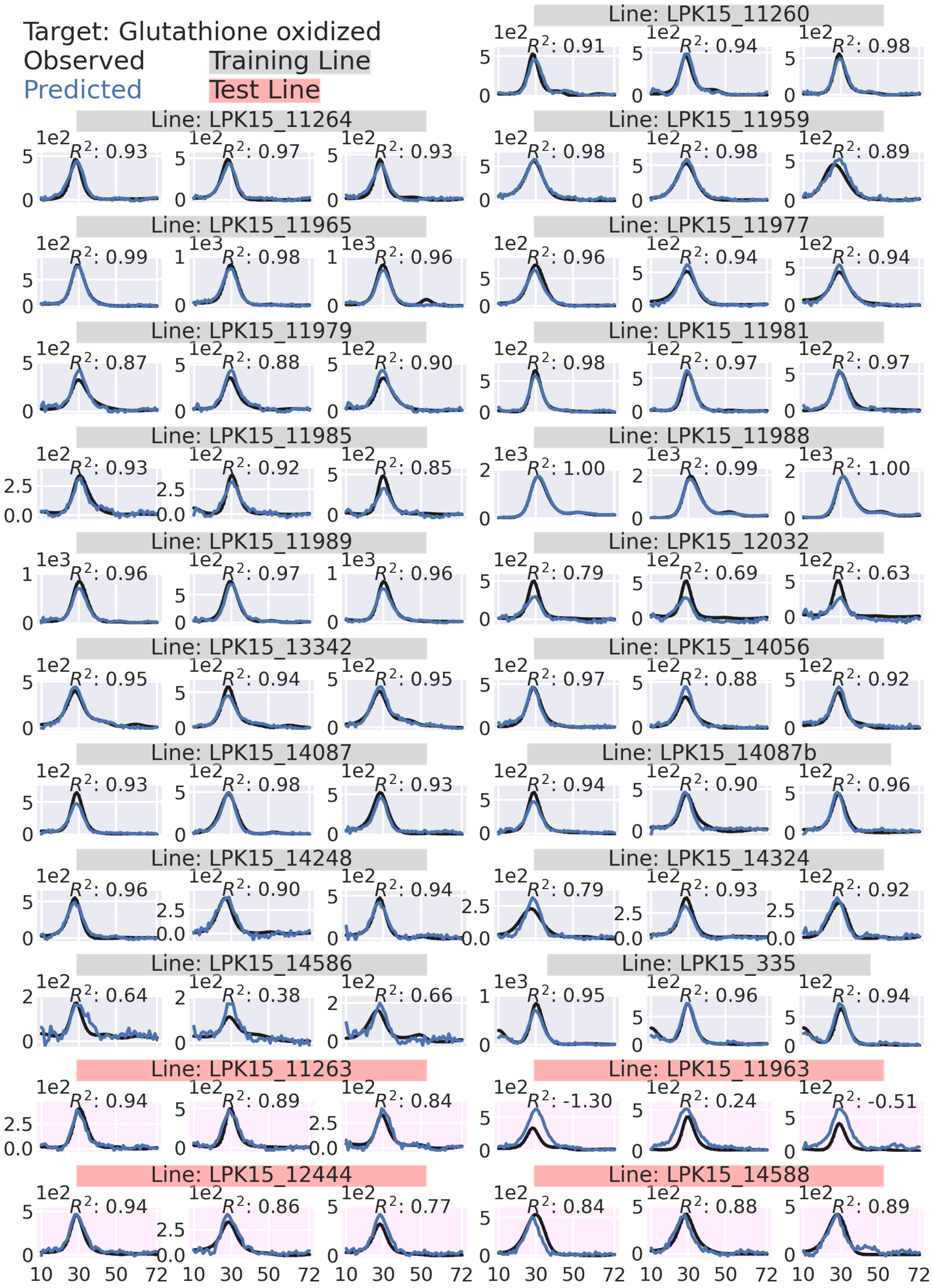
Target: Glutathione oxidized
Observed
Predicted
Training Line
Test Line
Line: LPK15_11260
Line: LPK15_11264
Line: LPK15_11959
Line: LPK15_11965
Line: LPK15_11977
Line: LPK15_11979
Line: LPK15_11981
Line: LPK15_11985
Line: LPK15_11988
Line: LPK15_11989
Line: LPK15_12032
Line: LPK15_13342
Line: LPK15_14056
Line: LPK15_14087
Line: LPK15_14087b
Line: LPK15_14248
Line: LPK15_14324
Line: LPK15_14586
Line: LPK15_335
Line: LPK15_11263
Line: LPK15_11963
Line: LPK15_12444
Line: LPK15_14588

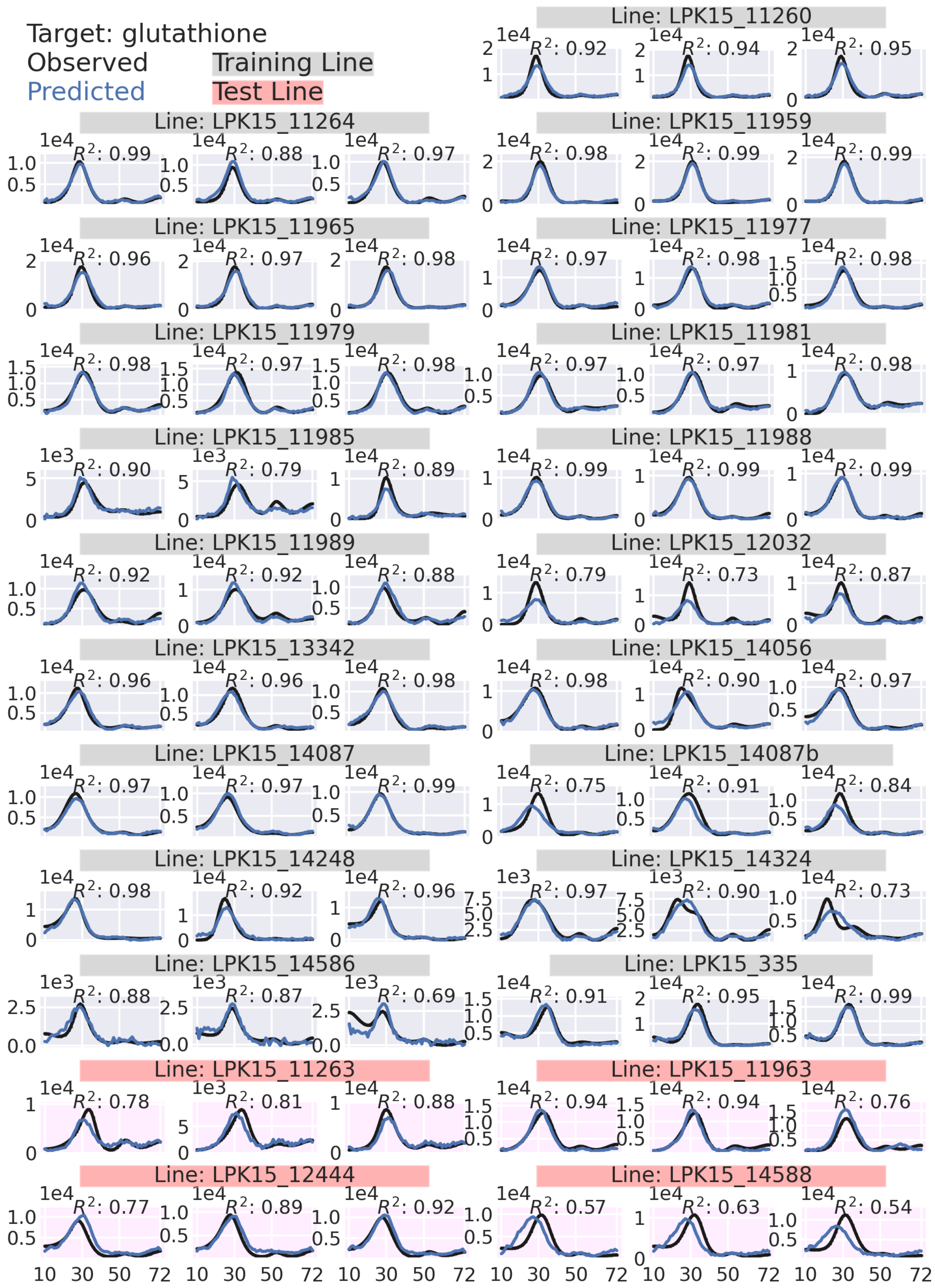
Target: glutathione
Observed
Predicted
Training Line
Test Line
Line: LPK15_11260
$R^2$: 0.92
$R^2$: 0.94
$R^2$: 0.95
Line: LPK15_11264
$R^2$: 0.99
$R^2$: 0.88
$R^2$: 0.97
Line: LPK15_11959
$R^2$: 0.98
$R^2$: 0.99
$R^2$: 0.99
Line: LPK15_11965
$R^2$: 0.96
$R^2$: 0.97
$R^2$: 0.98
Line: LPK15_11977
$R^2$: 0.97
$R^2$: 0.98
$R^2$: 0.98
Line: LPK15_11979
$R^2$: 0.98
$R^2$: 0.97
$R^2$: 0.98
Line: LPK15_11981
$R^2$: 0.97
$R^2$: 0.97
$R^2$: 0.98
Line: LPK15_11985
$R^2$: 0.90
$R^2$: 0.79
$R^2$: 0.89
Line: LPK15_11988
$R^2$: 0.99
$R^2$: 0.99
$R^2$: 0.99
Line: LPK15_11989
$R^2$: 0.92
$R^2$: 0.92
$R^2$: 0.88
Line: LPK15_12032
$R^2$: 0.79
$R^2$: 0.73
$R^2$: 0.87
Line: LPK15_13342
$R^2$: 0.96
$R^2$: 0.96
$R^2$: 0.98
Line: LPK15_14056
$R^2$: 0.98
$R^2$: 0.90
$R^2$: 0.97
Line: LPK15_14087
$R^2$: 0.97
$R^2$: 0.97
$R^2$: 0.99
Line: LPK15_14087b
$R^2$: 0.75
$R^2$: 0.91
$R^2$: 0.84
Line: LPK15_14248
$R^2$: 0.98
$R^2$: 0.92
$R^2$: 0.96
Line: LPK15_14324
$R^2$: 0.97
$R^2$: 0.90
$R^2$: 0.73
Line: LPK15_14586
$R^2$: 0.88
$R^2$: 0.87
$R^2$: 0.69
Line: LPK15_335
$R^2$: 0.91
$R^2$: 0.95
$R^2$: 0.99
Line: LPK15_11263
$R^2$: 0.78
$R^2$: 0.81
$R^2$: 0.88
Line: LPK15_11963
$R^2$: 0.94
$R^2$: 0.94
$R^2$: 0.76
Line: LPK15_12444
$R^2$: 0.77
$R^2$: 0.89
$R^2$: 0.92
Line: LPK15_14588
$R^2$: 0.57
$R^2$: 0.63
$R^2$: 0.54
10 30 50 72

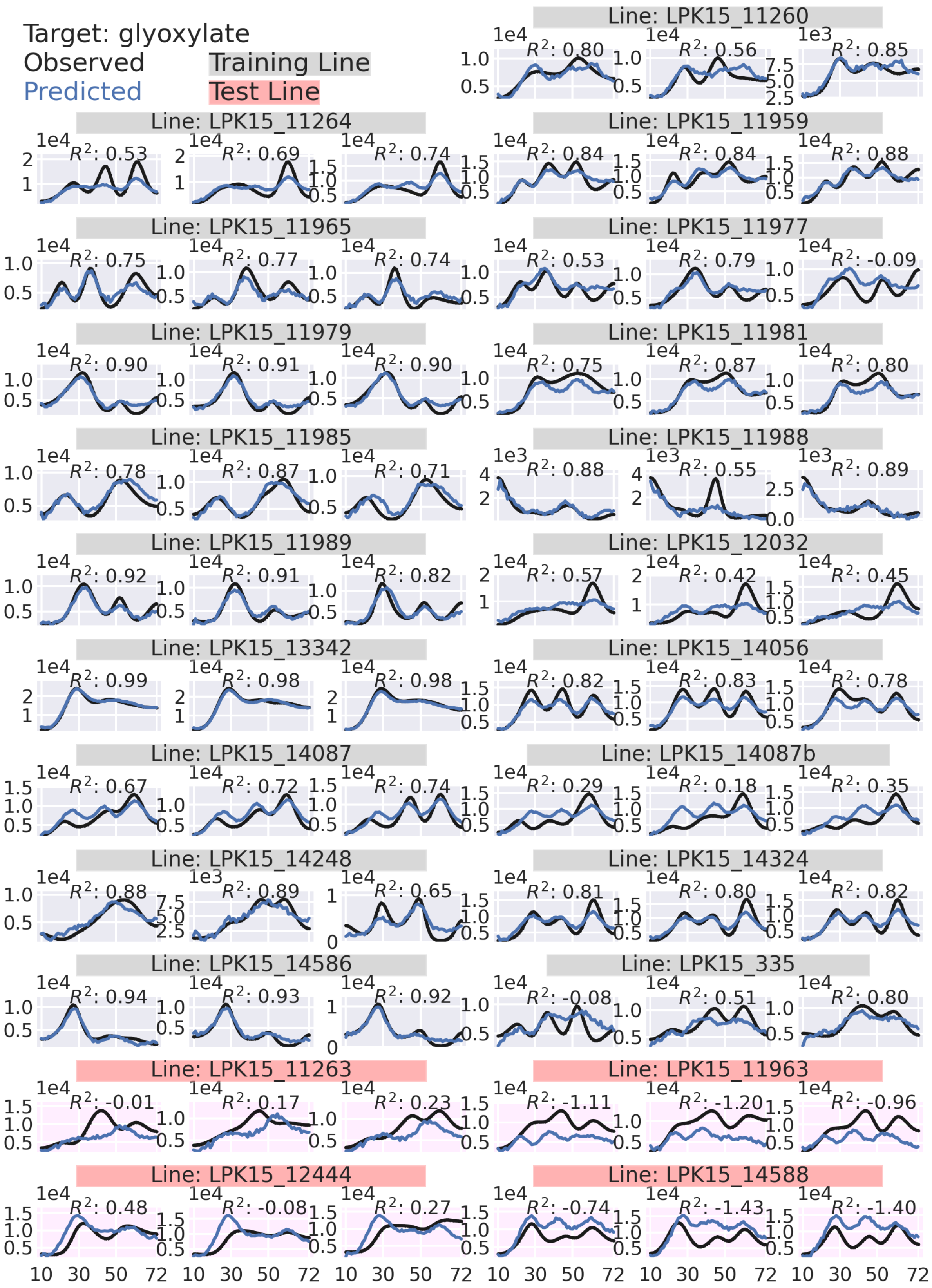

Target: glyoxylate
Observed
Predicted
Training Line
Test Line
Line: LPK15_11260
$R^2$: 0.80
$R^2$: 0.56
$R^2$: 0.85
Line: LPK15_11264
$R^2$: 0.53
$R^2$: 0.69
$R^2$: 0.74
Line: LPK15_11959
$R^2$: 0.84
$R^2$: 0.84
$R^2$: 0.88
Line: LPK15_11965
$R^2$: 0.75
$R^2$: 0.77
$R^2$: 0.74
Line: LPK15_11977
$R^2$: 0.53
$R^2$: 0.79
$R^2$: -0.09
Line: LPK15_11979
$R^2$: 0.90
$R^2$: 0.91
$R^2$: 0.90
Line: LPK15_11981
$R^2$: 0.75
$R^2$: 0.87
$R^2$: 0.80
Line: LPK15_11985
$R^2$: 0.78
$R^2$: 0.87
$R^2$: 0.71
Line: LPK15_11988
$R^2$: 0.88
$R^2$: 0.55
$R^2$: 0.89
Line: LPK15_11989
$R^2$: 0.92
$R^2$: 0.91
$R^2$: 0.82
Line: LPK15_12032
$R^2$: 0.57
$R^2$: 0.42
$R^2$: 0.45
Line: LPK15_13342
$R^2$: 0.99
$R^2$: 0.98
$R^2$: 0.98
Line: LPK15_14056
$R^2$: 0.82
$R^2$: 0.83
$R^2$: 0.78
Line: LPK15_14087
$R^2$: 0.67
$R^2$: 0.72
$R^2$: 0.74
Line: LPK15_14087b
$R^2$: 0.29
$R^2$: 0.18
$R^2$: 0.35
Line: LPK15_14248
$R^2$: 0.88
$R^2$: 0.89
$R^2$: 0.65
Line: LPK15_14324
$R^2$: 0.81
$R^2$: 0.80
$R^2$: 0.82
Line: LPK15_14586
$R^2$: 0.94
$R^2$: 0.93
$R^2$: 0.92
Line: LPK15_335
$R^2$: -0.08
$R^2$: 0.51
$R^2$: 0.80
Line: LPK15_11263
$R^2$: -0.01
$R^2$: 0.17
$R^2$: 0.23
Line: LPK15_11963
$R^2$: -1.11
$R^2$: -1.20
$R^2$: -0.96
Line: LPK15_12444
$R^2$: 0.48
$R^2$: -0.08
$R^2$: 0.27
Line: LPK15_14588
$R^2$: -0.74
$R^2$: -1.43
$R^2$: -1.40
10 30 50 72

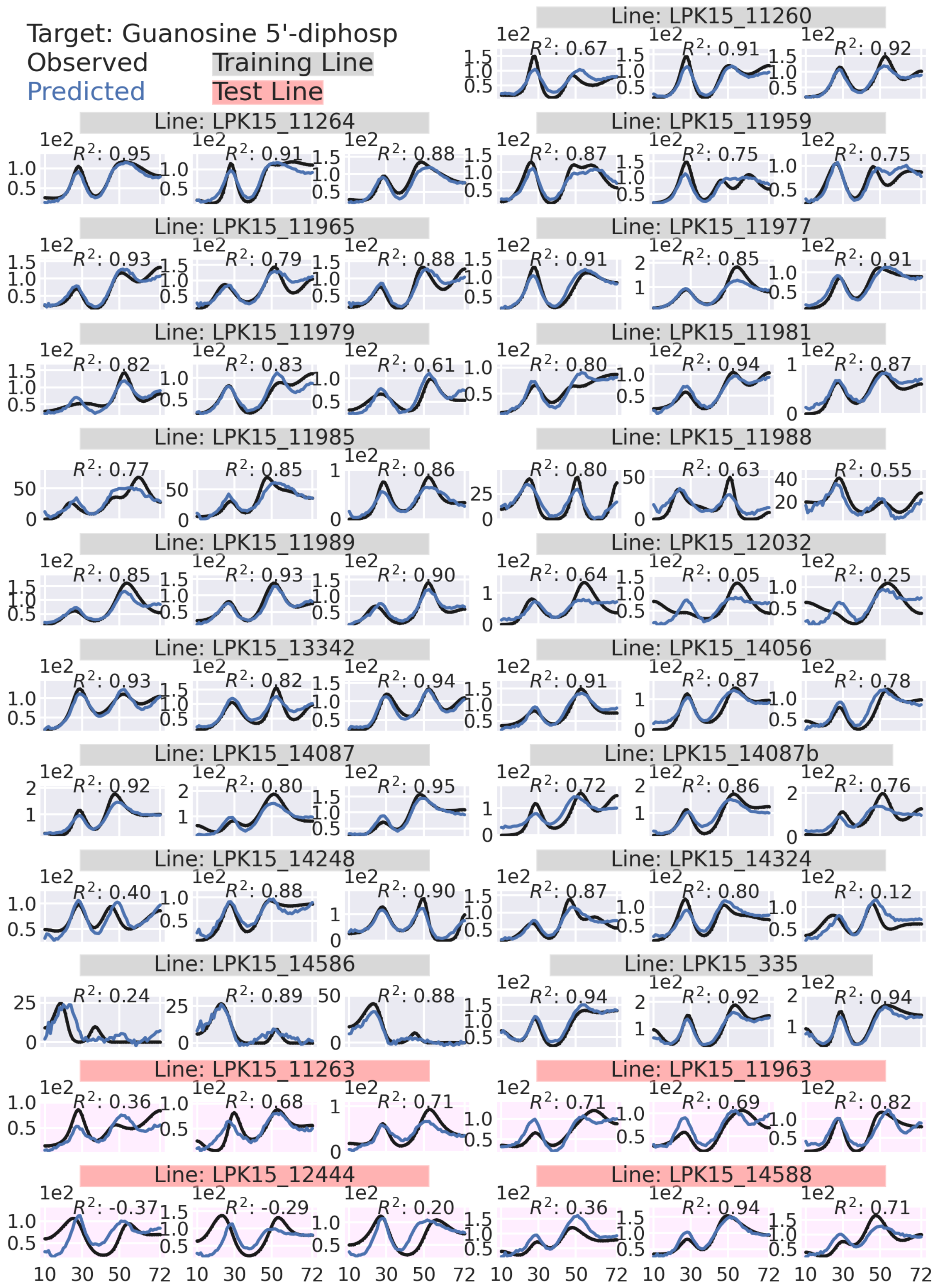

Target: Guanosine 5'-diphosp
Observed
Predicted
Training Line
Test Line
Line: LPK15_11260
Line: LPK15_11264
Line: LPK15_11959
Line: LPK15_11965
Line: LPK15_11977
Line: LPK15_11979
Line: LPK15_11981
Line: LPK15_11985
Line: LPK15_11988
Line: LPK15_11989
Line: LPK15_12032
Line: LPK15_13342
Line: LPK15_14056
Line: LPK15_14087
Line: LPK15_14087b
Line: LPK15_14248
Line: LPK15_14324
Line: LPK15_14586
Line: LPK15_335
Line: LPK15_11263
Line: LPK15_11963
Line: LPK15_12444
Line: LPK15_14588

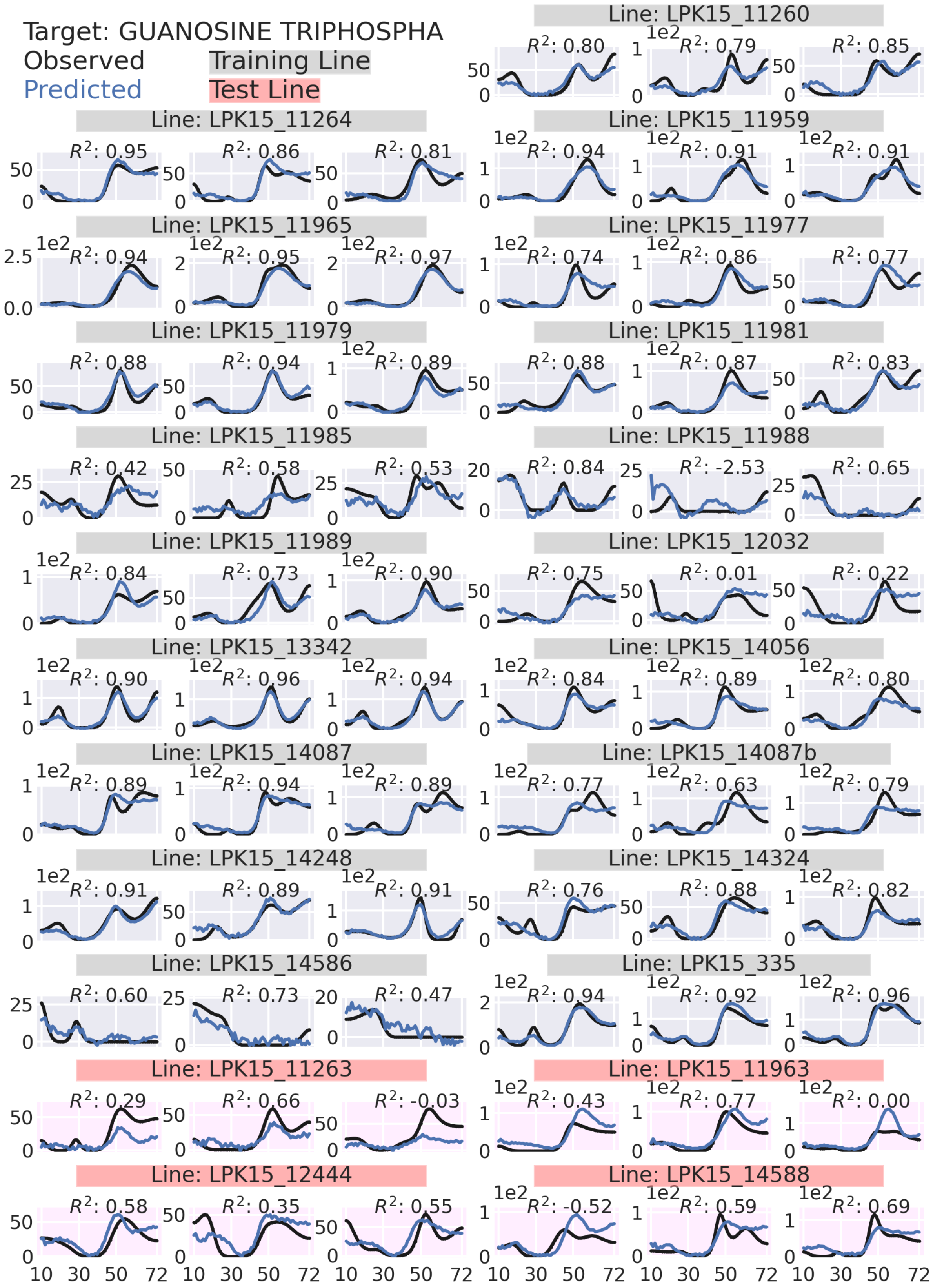

Target: GUANOSINE TRIPHOSPHA
Observed
Predicted
Training Line
Test Line
Line: LPK15_11260
Line: LPK15_11264
Line: LPK15_11959
Line: LPK15_11965
Line: LPK15_11977
Line: LPK15_11979
Line: LPK15_11981
Line: LPK15_11985
Line: LPK15_11988
Line: LPK15_11989
Line: LPK15_12032
Line: LPK15_13342
Line: LPK15_14056
Line: LPK15_14087
Line: LPK15_14087b
Line: LPK15_14248
Line: LPK15_14324
Line: LPK15_14586
Line: LPK15_335
Line: LPK15_11263
Line: LPK15_11963
Line: LPK15_12444
Line: LPK15_14588

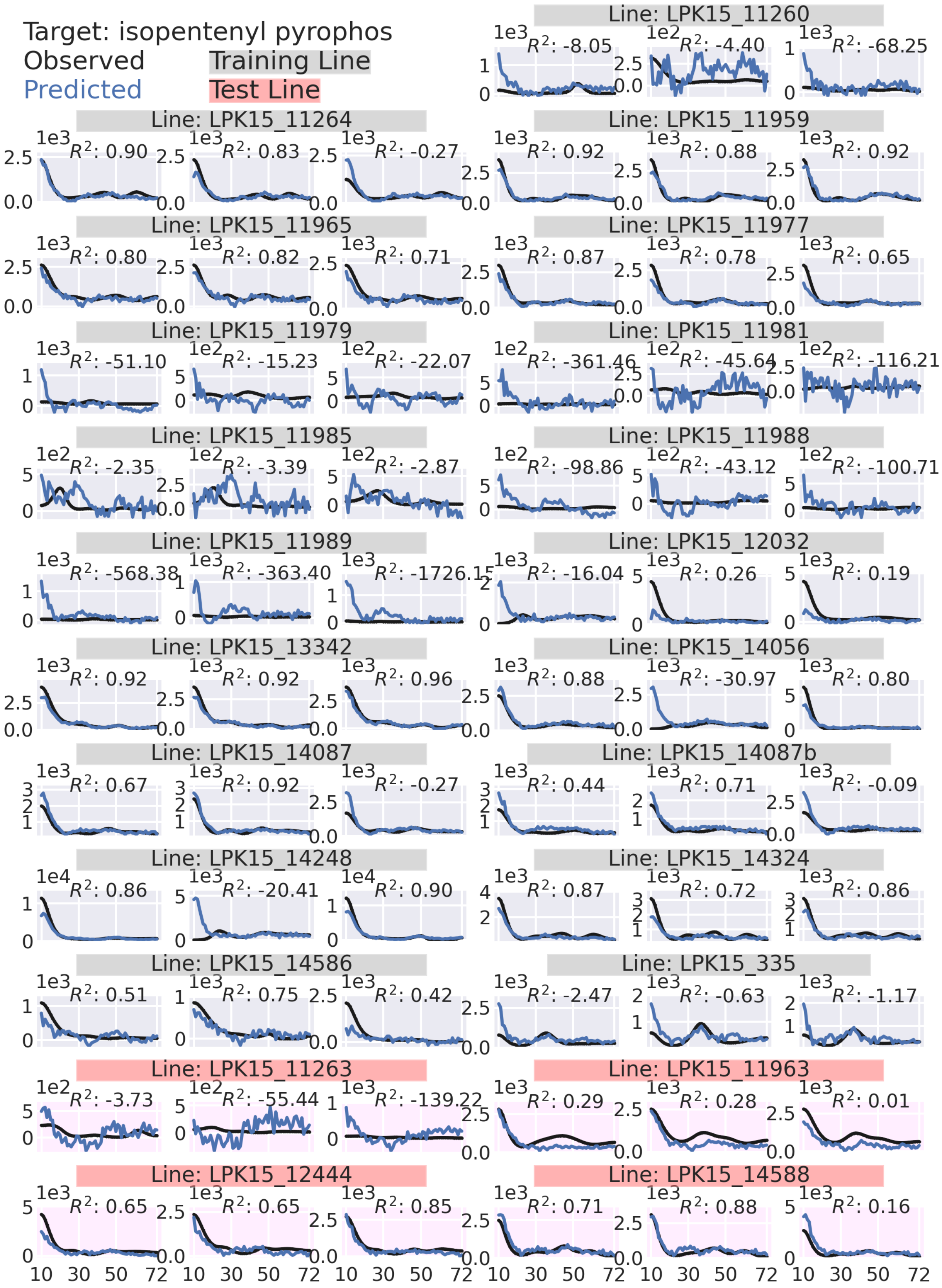
Target: isopentenyl pyrophos
Observed
Predicted
Training Line
Test Line
Line: LPK15_11260
$R^2$: -8.05
$R^2$: -4.40
$R^2$: -68.25
Line: LPK15_11264
$R^2$: 0.90
$R^2$: 0.83
$R^2$: -0.27
Line: LPK15_11959
$R^2$: 0.92
$R^2$: 0.88
$R^2$: 0.92
Line: LPK15_11965
$R^2$: 0.80
$R^2$: 0.82
$R^2$: 0.71
Line: LPK15_11977
$R^2$: 0.87
$R^2$: 0.78
$R^2$: 0.65
Line: LPK15_11979
$R^2$: -51.10
$R^2$: -15.23
$R^2$: -22.07
Line: LPK15_11981
$R^2$: -361.46
$R^2$: -45.64
$R^2$: -116.21
Line: LPK15_11985
$R^2$: -2.35
$R^2$: -3.39
$R^2$: -2.87
Line: LPK15_11988
$R^2$: -98.86
$R^2$: -43.12
$R^2$: -100.71
Line: LPK15_11989
$R^2$: -568.38
$R^2$: -363.40
$R^2$: -1726.15
Line: LPK15_12032
$R^2$: -16.04
$R^2$: 0.26
$R^2$: 0.19
Line: LPK15_13342
$R^2$: 0.92
$R^2$: 0.92
$R^2$: 0.96
Line: LPK15_14056
$R^2$: 0.88
$R^2$: -30.97
$R^2$: 0.80
Line: LPK15_14087
$R^2$: 0.67
$R^2$: 0.92
$R^2$: -0.27
Line: LPK15_14087b
$R^2$: 0.44
$R^2$: 0.71
$R^2$: -0.09
Line: LPK15_14248
$R^2$: 0.86
$R^2$: -20.41
$R^2$: 0.90
Line: LPK15_14324
$R^2$: 0.87
$R^2$: 0.72
$R^2$: 0.86
Line: LPK15_14586
$R^2$: 0.51
$R^2$: 0.75
$R^2$: 0.42
Line: LPK15_335
$R^2$: -2.47
$R^2$: -0.63
$R^2$: -1.17
Line: LPK15_11263
$R^2$: -3.73
$R^2$: -55.44
$R^2$: -139.22
Line: LPK15_11963
$R^2$: 0.29
$R^2$: 0.28
$R^2$: 0.01
Line: LPK15_12444
$R^2$: 0.65
$R^2$: 0.65
$R^2$: 0.85
Line: LPK15_14588
$R^2$: 0.71
$R^2$: 0.88
$R^2$: 0.16
10 30 50 72

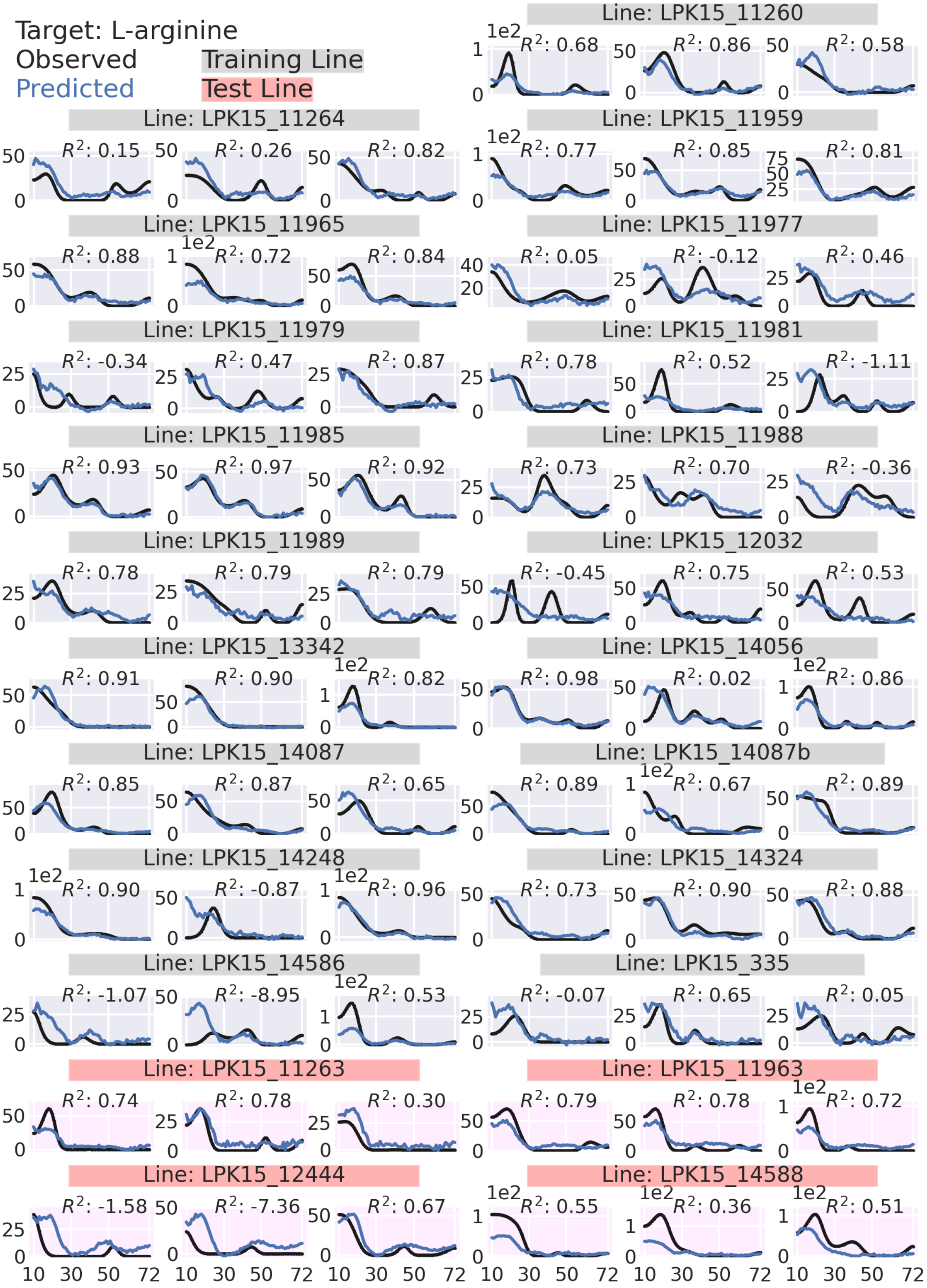

Target: L-arginine
Observed
Predicted
Training Line
Test Line
Line: LPK15_11260
Line: LPK15_11264
Line: LPK15_11959
Line: LPK15_11965
Line: LPK15_11977
Line: LPK15_11979
Line: LPK15_11981
Line: LPK15_11985
Line: LPK15_11988
Line: LPK15_11989
Line: LPK15_12032
Line: LPK15_13342
Line: LPK15_14056
Line: LPK15_14087
Line: LPK15_14087b
Line: LPK15_14248
Line: LPK15_14324
Line: LPK15_14586
Line: LPK15_335
Line: LPK15_11263
Line: LPK15_11963
Line: LPK15_12444
Line: LPK15_14588

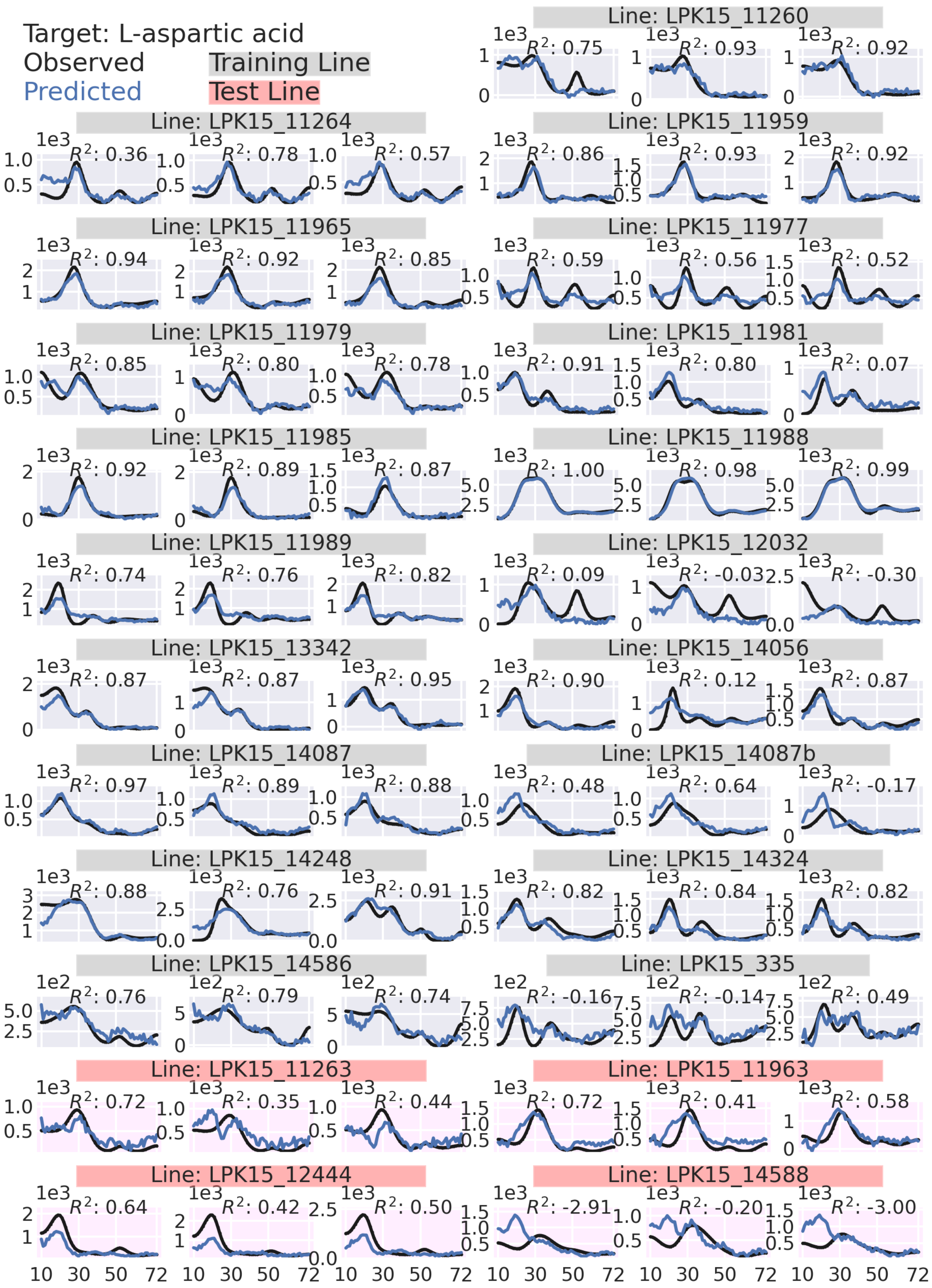

Target: L-aspartic acid
Observed
Predicted
Training Line
Test Line
Line: LPK15_11260
R²: 0.75
R²: 0.93
R²: 0.92
Line: LPK15_11264
R²: 0.36
R²: 0.78
R²: 0.57
Line: LPK15_11959
R²: 0.86
R²: 0.93
R²: 0.92
Line: LPK15_11965
R²: 0.94
R²: 0.92
R²: 0.85
Line: LPK15_11977
R²: 0.59
R²: 0.56
R²: 0.52
Line: LPK15_11979
R²: 0.85
R²: 0.80
R²: 0.78
Line: LPK15_11981
R²: 0.91
R²: 0.80
R²: 0.07
Line: LPK15_11985
R²: 0.92
R²: 0.89
R²: 0.87
Line: LPK15_11988
R²: 1.00
R²: 0.98
R²: 0.99
Line: LPK15_11989
R²: 0.74
R²: 0.76
R²: 0.82
Line: LPK15_12032
R²: 0.09
R²: -0.03
R²: -0.30
Line: LPK15_13342
R²: 0.87
R²: 0.87
R²: 0.95
Line: LPK15_14056
R²: 0.90
R²: 0.12
R²: 0.87
Line: LPK15_14087
R²: 0.97
R²: 0.89
R²: 0.88
Line: LPK15_14087b
R²: 0.48
R²: 0.64
R²: -0.17
Line: LPK15_14248
R²: 0.88
R²: 0.76
R²: 0.91
Line: LPK15_14324
R²: 0.82
R²: 0.84
R²: 0.82
Line: LPK15_14586
R²: 0.76
R²: 0.79
R²: 0.74
Line: LPK15_335
R²: -0.16
R²: -0.14
R²: 0.49
Line: LPK15_11263
R²: 0.72
R²: 0.35
R²: 0.44
Line: LPK15_11963
R²: 0.72
R²: 0.41
R²: 0.58
Line: LPK15_12444
R²: 0.64
R²: 0.42
R²: 0.50
Line: LPK15_14588
R²: -2.91
R²: -0.20
R²: -3.00
10 30 50 72

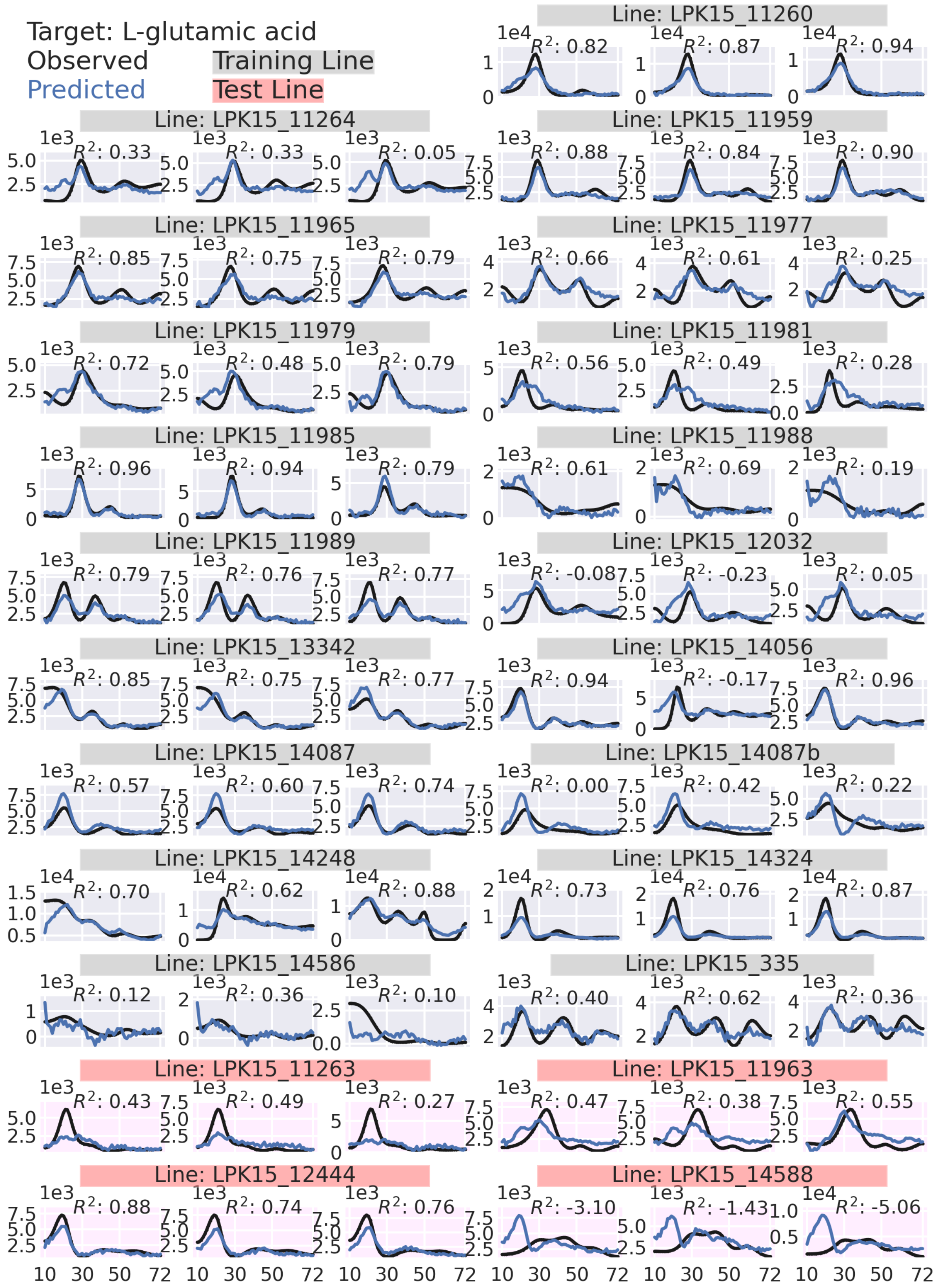
Target: L-glutamic acid
Observed
Predicted
Training Line
Test Line
Line: LPK15_11260
R²: 0.82 R²: 0.87 R²: 0.94
Line: LPK15_11264
R²: 0.33 R²: 0.33 R²: 0.05
Line: LPK15_11959
R²: 0.88 R²: 0.84 R²: 0.90
Line: LPK15_11965
R²: 0.85 R²: 0.75 R²: 0.79
Line: LPK15_11977
R²: 0.66 R²: 0.61 R²: 0.25
Line: LPK15_11979
R²: 0.72 R²: 0.48 R²: 0.79
Line: LPK15_11981
R²: 0.56 R²: 0.49 R²: 0.28
Line: LPK15_11985
R²: 0.96 R²: 0.94 R²: 0.79
Line: LPK15_11988
R²: 0.61 R²: 0.69 R²: 0.19
Line: LPK15_11989
R²: 0.79 R²: 0.76 R²: 0.77
Line: LPK15_12032
R²: -0.08 R²: -0.23 R²: 0.05
Line: LPK15_13342
R²: 0.85 R²: 0.75 R²: 0.77
Line: LPK15_14056
R²: 0.94 R²: -0.17 R²: 0.96
Line: LPK15_14087
R²: 0.57 R²: 0.60 R²: 0.74
Line: LPK15_14087b
R²: 0.00 R²: 0.42 R²: 0.22
Line: LPK15_14248
R²: 0.70 R²: 0.62 R²: 0.88
Line: LPK15_14324
R²: 0.73 R²: 0.76 R²: 0.87
Line: LPK15_14586
R²: 0.12 R²: 0.36 R²: 0.10
Line: LPK15_335
R²: 0.40 R²: 0.62 R²: 0.36
Line: LPK15_11263
R²: 0.43 R²: 0.49 R²: 0.27
Line: LPK15_11963
R²: 0.47 R²: 0.38 R²: 0.55
Line: LPK15_12444
R²: 0.88 R²: 0.74 R²: 0.76
Line: LPK15_14588
R²: -3.10 R²: -1.43 R²: -5.06
10 30 50 72

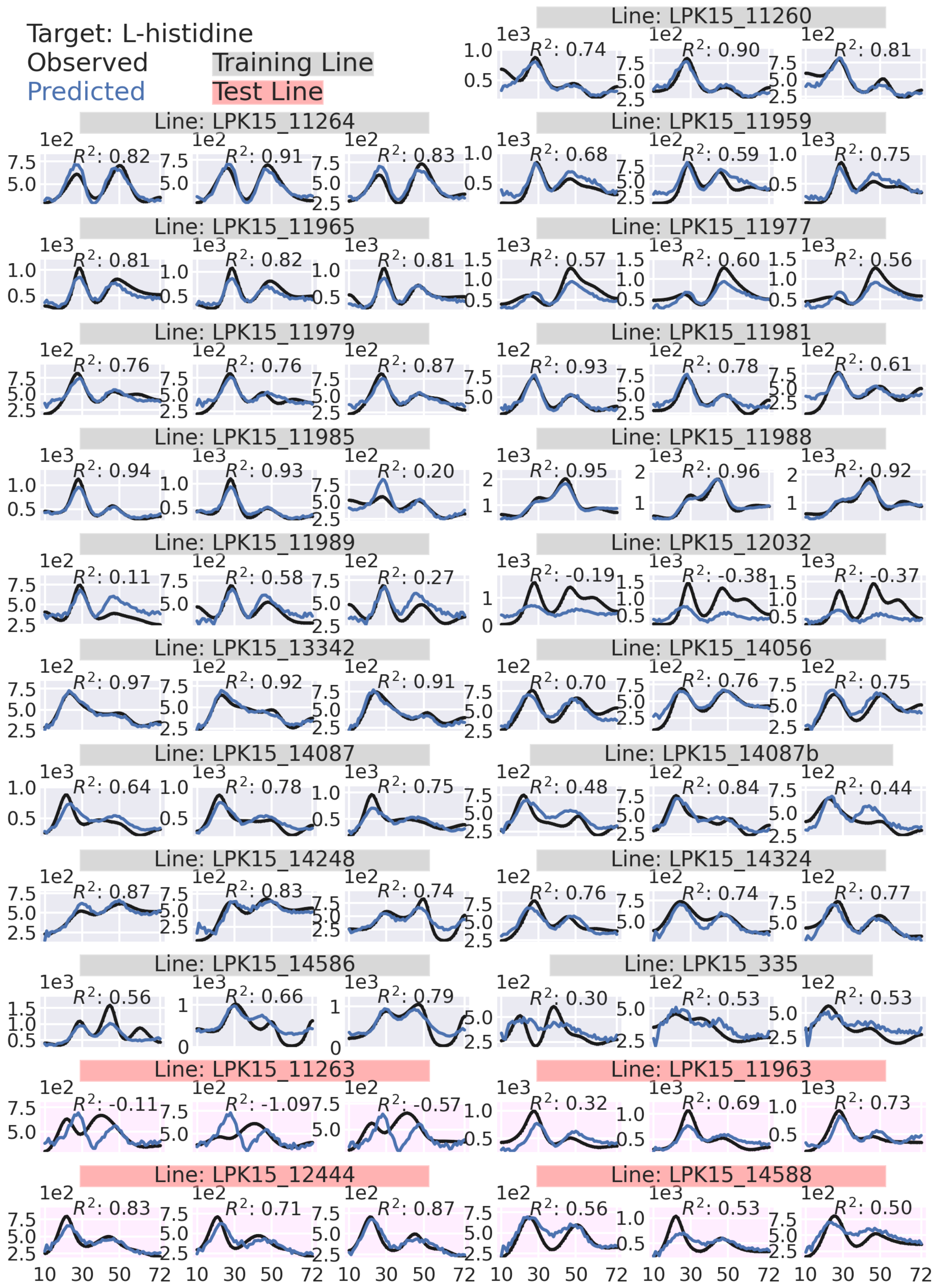

Target: L-histidine
Observed
Predicted
Training Line
Test Line
Line: LPK15_11260
Line: LPK15_11264
Line: LPK15_11959
Line: LPK15_11965
Line: LPK15_11977
Line: LPK15_11979
Line: LPK15_11981
Line: LPK15_11985
Line: LPK15_11988
Line: LPK15_11989
Line: LPK15_12032
Line: LPK15_13342
Line: LPK15_14056
Line: LPK15_14087
Line: LPK15_14087b
Line: LPK15_14248
Line: LPK15_14324
Line: LPK15_14586
Line: LPK15_335
Line: LPK15_11263
Line: LPK15_11963
Line: LPK15_12444
Line: LPK15_14588

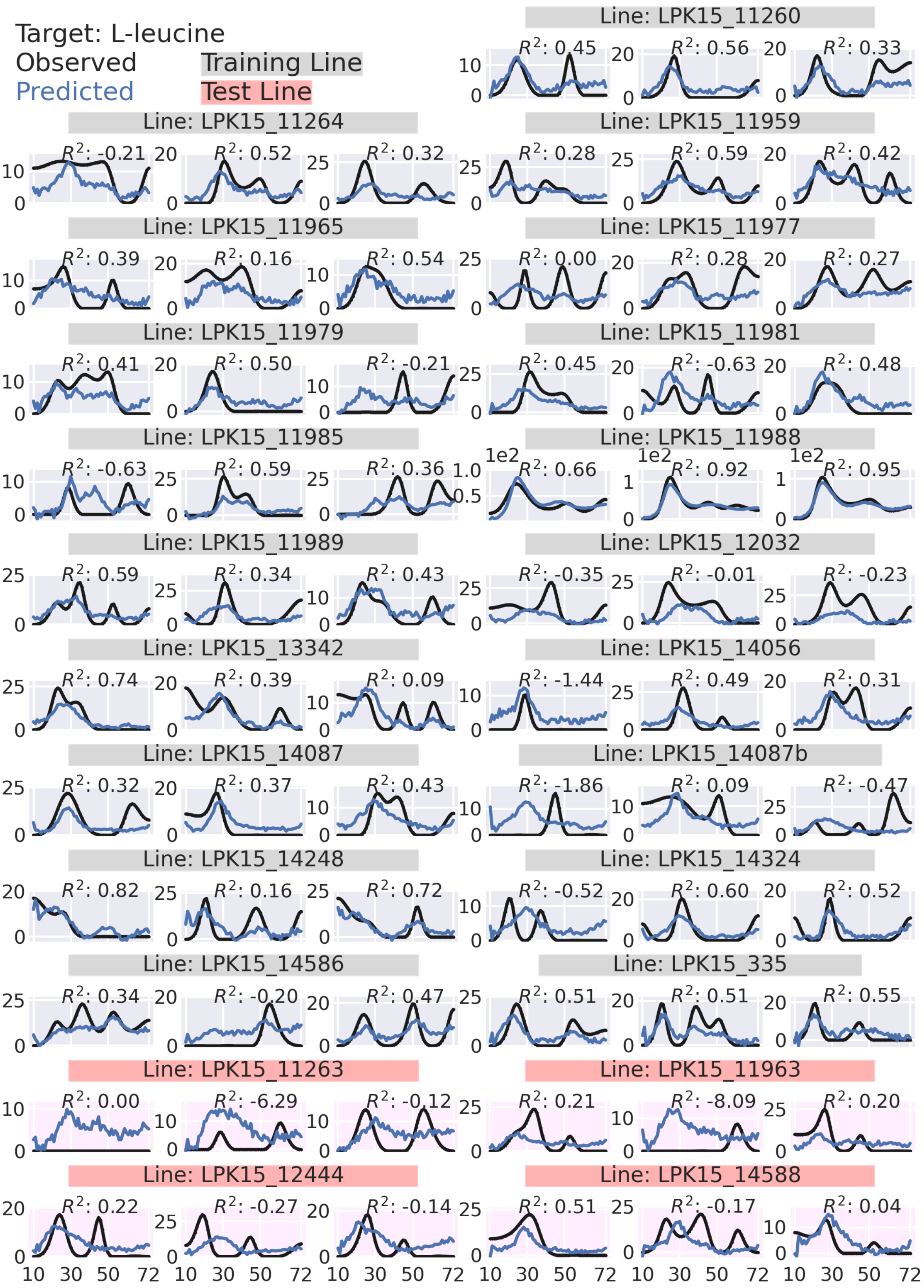
Target: L-leucine
Observed
Predicted
Training Line
Test Line
Line: LPK15_11260
Line: LPK15_11264
Line: LPK15_11959
Line: LPK15_11965
Line: LPK15_11977
Line: LPK15_11979
Line: LPK15_11981
Line: LPK15_11985
Line: LPK15_11988
Line: LPK15_11989
Line: LPK15_12032
Line: LPK15_13342
Line: LPK15_14056
Line: LPK15_14087
Line: LPK15_14087b
Line: LPK15_14248
Line: LPK15_14324
Line: LPK15_14586
Line: LPK15_335
Line: LPK15_11263
Line: LPK15_11963
Line: LPK15_12444
Line: LPK15_14588

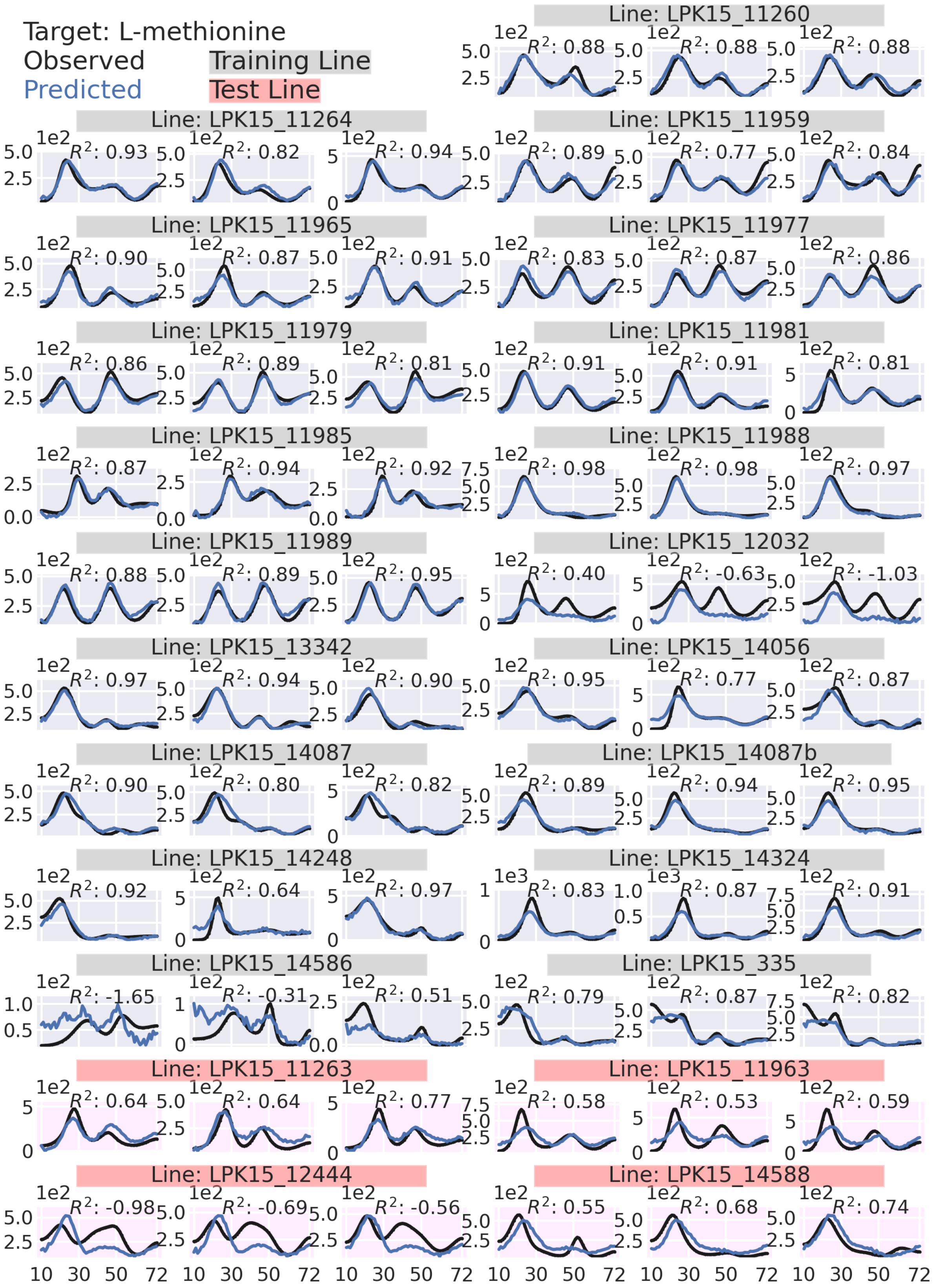

Target: L-methionine
Observed
Predicted
Training Line
Test Line
Line: LPK15_11260
R²: 0.88 R²: 0.88 R²: 0.88
Line: LPK15_11264
R²: 0.93 R²: 0.82 R²: 0.94
Line: LPK15_11959
R²: 0.89 R²: 0.77 R²: 0.84
Line: LPK15_11965
R²: 0.90 R²: 0.87 R²: 0.91
Line: LPK15_11977
R²: 0.83 R²: 0.87 R²: 0.86
Line: LPK15_11979
R²: 0.86 R²: 0.89 R²: 0.81
Line: LPK15_11981
R²: 0.91 R²: 0.91 R²: 0.81
Line: LPK15_11985
R²: 0.87 R²: 0.94 R²: 0.92
Line: LPK15_11988
R²: 0.98 R²: 0.98 R²: 0.97
Line: LPK15_11989
R²: 0.88 R²: 0.89 R²: 0.95
Line: LPK15_12032
R²: 0.40 R²: -0.63 R²: -1.03
Line: LPK15_13342
R²: 0.97 R²: 0.94 R²: 0.90
Line: LPK15_14056
R²: 0.95 R²: 0.77 R²: 0.87
Line: LPK15_14087
R²: 0.90 R²: 0.80 R²: 0.82
Line: LPK15_14087b
R²: 0.89 R²: 0.94 R²: 0.95
Line: LPK15_14248
R²: 0.92 R²: 0.64 R²: 0.97
Line: LPK15_14324
R²: 0.83 R²: 0.87 R²: 0.91
Line: LPK15_14586
R²: -1.65 R²: -0.31 R²: 0.51
Line: LPK15_335
R²: 0.79 R²: 0.87 R²: 0.82
Line: LPK15_11263
R²: 0.64 R²: 0.64 R²: 0.77
Line: LPK15_11963
R²: 0.58 R²: 0.53 R²: 0.59
Line: LPK15_12444
R²: -0.98 R²: -0.69 R²: -0.56
Line: LPK15_14588
R²: 0.55 R²: 0.68 R²: 0.74
10 30 50 72

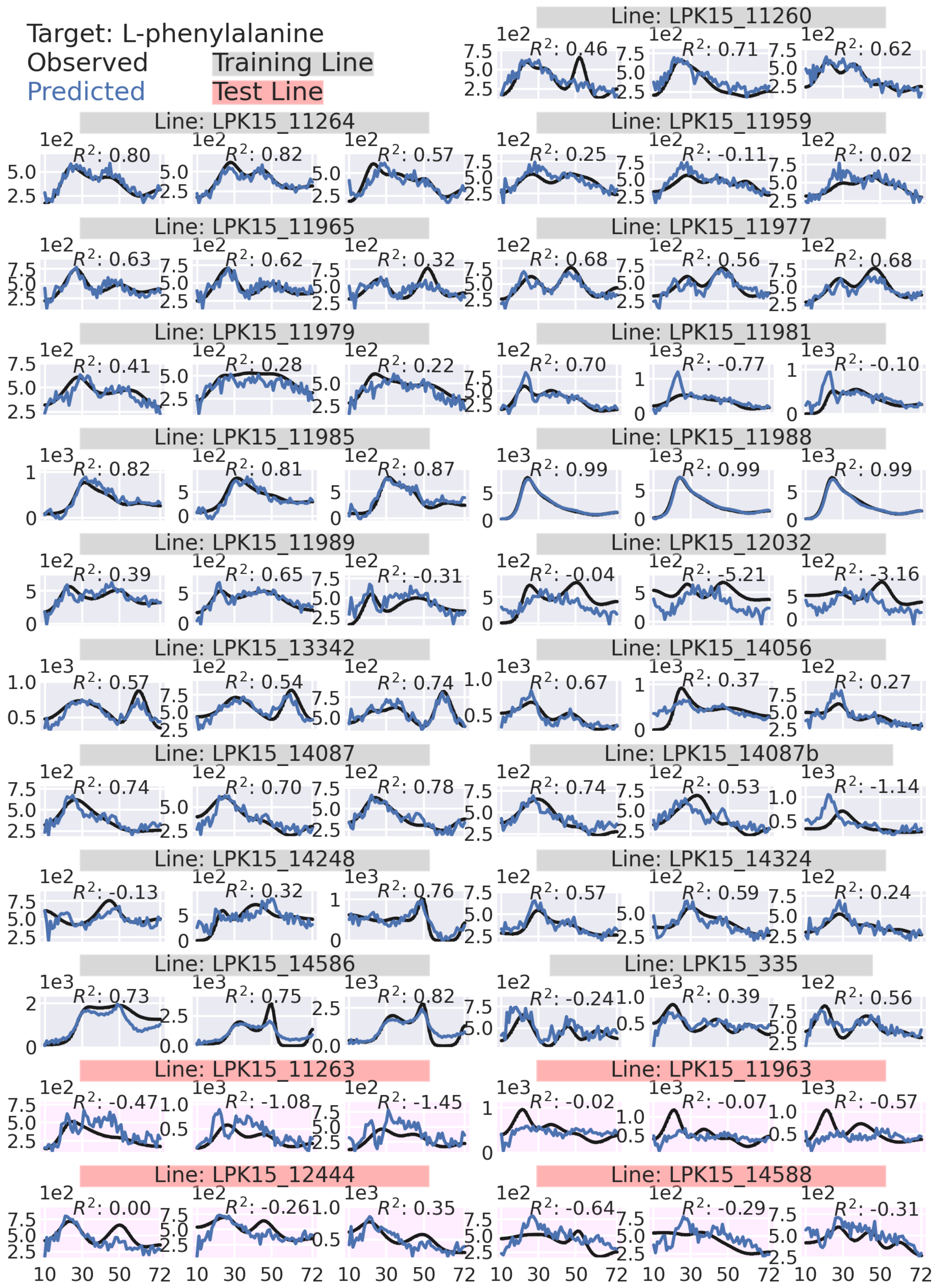

Target: L-phenylalanine
Observed
Predicted
Training Line
Test Line
Line: LPK15_11260
Line: LPK15_11264
Line: LPK15_11959
Line: LPK15_11965
Line: LPK15_11977
Line: LPK15_11979
Line: LPK15_11981
Line: LPK15_11985
Line: LPK15_11988
Line: LPK15_11989
Line: LPK15_12032
Line: LPK15_13342
Line: LPK15_14056
Line: LPK15_14087
Line: LPK15_14087b
Line: LPK15_14248
Line: LPK15_14324
Line: LPK15_14586
Line: LPK15_335
Line: LPK15_11263
Line: LPK15_11963
Line: LPK15_12444
Line: LPK15_14588

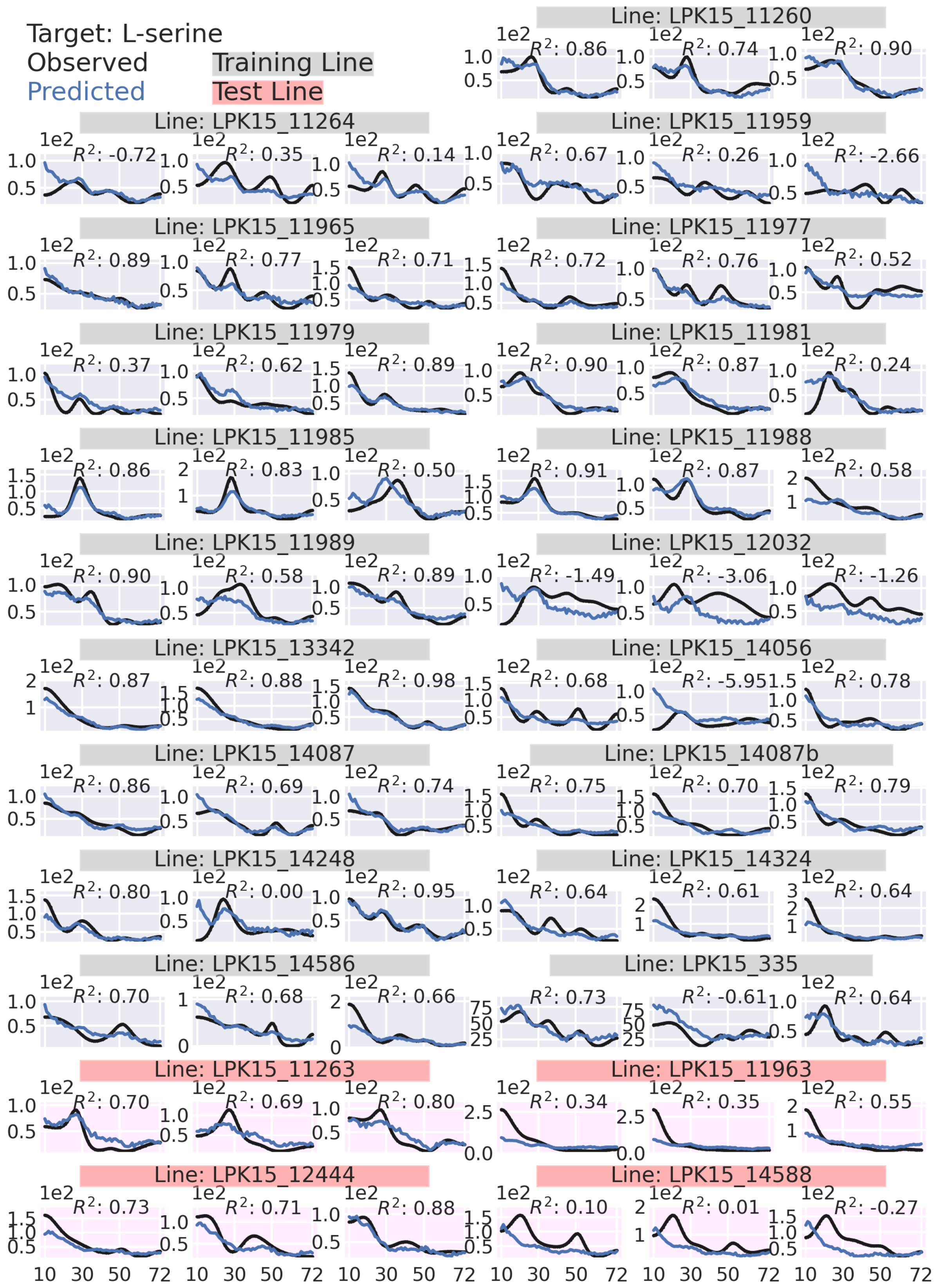

Target: L-serine
Observed
Predicted
Training Line
Test Line
Line: LPK15_11260
Line: LPK15_11264
Line: LPK15_11959
Line: LPK15_11965
Line: LPK15_11977
Line: LPK15_11979
Line: LPK15_11981
Line: LPK15_11985
Line: LPK15_11988
Line: LPK15_11989
Line: LPK15_12032
Line: LPK15_13342
Line: LPK15_14056
Line: LPK15_14087
Line: LPK15_14087b
Line: LPK15_14248
Line: LPK15_14324
Line: LPK15_14586
Line: LPK15_335
Line: LPK15_11263
Line: LPK15_11963
Line: LPK15_12444
Line: LPK15_14588
$R^2$: 0.86 $R^2$: 0.74 $R^2$: 0.90
$R^2$: -0.72 $R^2$: 0.35 $R^2$: 0.14 $R^2$: 0.67 $R^2$: 0.26 $R^2$: -2.66
$R^2$: 0.89 $R^2$: 0.77 $R^2$: 0.71 $R^2$: 0.72 $R^2$: 0.76 $R^2$: 0.52
$R^2$: 0.37 $R^2$: 0.62 $R^2$: 0.89 $R^2$: 0.90 $R^2$: 0.87 $R^2$: 0.24
$R^2$: 0.86 $R^2$: 0.83 $R^2$: 0.50 $R^2$: 0.91 $R^2$: 0.87 $R^2$: 0.58
$R^2$: 0.90 $R^2$: 0.58 $R^2$: 0.89 $R^2$: -1.49 $R^2$: -3.06 $R^2$: -1.26
$R^2$: 0.87 $R^2$: 0.88 $R^2$: 0.98 $R^2$: 0.68 $R^2$: -5.95 $R^2$: 0.78
$R^2$: 0.86 $R^2$: 0.69 $R^2$: 0.74 $R^2$: 0.75 $R^2$: 0.70 $R^2$: 0.79
$R^2$: 0.80 $R^2$: 0.00 $R^2$: 0.95 $R^2$: 0.64 $R^2$: 0.61 $R^2$: 0.64
$R^2$: 0.70 $R^2$: 0.68 $R^2$: 0.66 $R^2$: 0.73 $R^2$: -0.61 $R^2$: 0.64
$R^2$: 0.70 $R^2$: 0.69 $R^2$: 0.80 $R^2$: 0.34 $R^2$: 0.35 $R^2$: 0.55
$R^2$: 0.73 $R^2$: 0.71 $R^2$: 0.88 $R^2$: 0.10 $R^2$: 0.01 $R^2$: -0.27
1e2
10 30 50 72

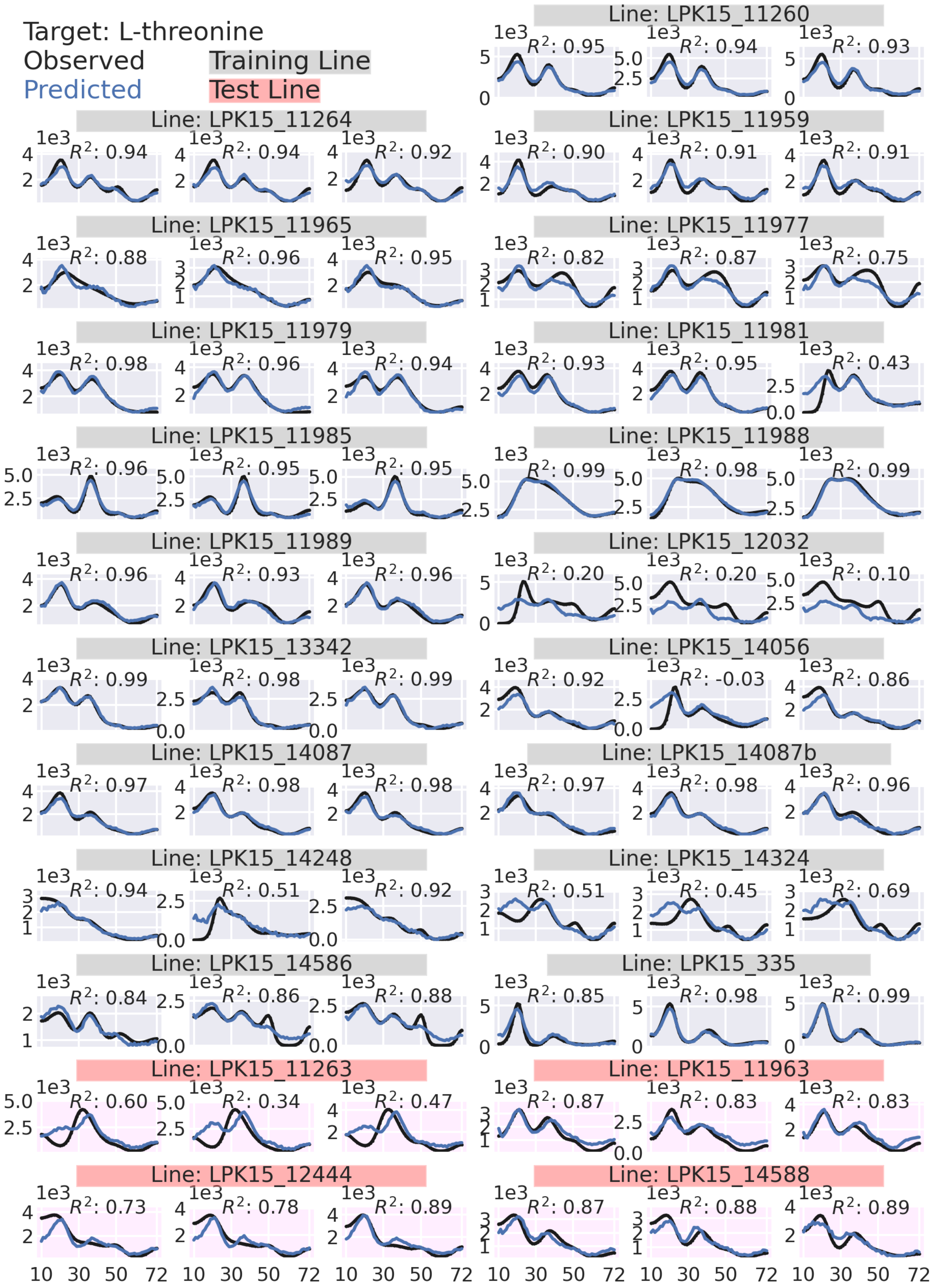

Target: L-threonine
Observed
Predicted
Training Line
Test Line
Line: LPK15_11260
$R^2$: 0.95
$R^2$: 0.94
$R^2$: 0.93
Line: LPK15_11264
$R^2$: 0.94
$R^2$: 0.94
$R^2$: 0.92
Line: LPK15_11959
$R^2$: 0.90
$R^2$: 0.91
$R^2$: 0.91
Line: LPK15_11965
$R^2$: 0.88
$R^2$: 0.96
$R^2$: 0.95
Line: LPK15_11977
$R^2$: 0.82
$R^2$: 0.87
$R^2$: 0.75
Line: LPK15_11979
$R^2$: 0.98
$R^2$: 0.96
$R^2$: 0.94
Line: LPK15_11981
$R^2$: 0.93
$R^2$: 0.95
$R^2$: 0.43
Line: LPK15_11985
$R^2$: 0.96
$R^2$: 0.95
$R^2$: 0.95
Line: LPK15_11988
$R^2$: 0.99
$R^2$: 0.98
$R^2$: 0.99
Line: LPK15_11989
$R^2$: 0.96
$R^2$: 0.93
$R^2$: 0.96
Line: LPK15_12032
$R^2$: 0.20
$R^2$: 0.20
$R^2$: 0.10
Line: LPK15_13342
$R^2$: 0.99
$R^2$: 0.98
$R^2$: 0.99
Line: LPK15_14056
$R^2$: 0.92
$R^2$: -0.03
$R^2$: 0.86
Line: LPK15_14087
$R^2$: 0.97
$R^2$: 0.98
$R^2$: 0.98
Line: LPK15_14087b
$R^2$: 0.97
$R^2$: 0.98
$R^2$: 0.96
Line: LPK15_14248
$R^2$: 0.94
$R^2$: 0.51
$R^2$: 0.92
Line: LPK15_14324
$R^2$: 0.51
$R^2$: 0.45
$R^2$: 0.69
Line: LPK15_14586
$R^2$: 0.84
$R^2$: 0.86
$R^2$: 0.88
Line: LPK15_335
$R^2$: 0.85
$R^2$: 0.98
$R^2$: 0.99
Line: LPK15_11263
$R^2$: 0.60
$R^2$: 0.34
$R^2$: 0.47
Line: LPK15_11963
$R^2$: 0.87
$R^2$: 0.83
$R^2$: 0.83
Line: LPK15_12444
$R^2$: 0.73
$R^2$: 0.78
$R^2$: 0.89
Line: LPK15_14588
$R^2$: 0.87
$R^2$: 0.88
$R^2$: 0.89
1e3
10 30 50 72

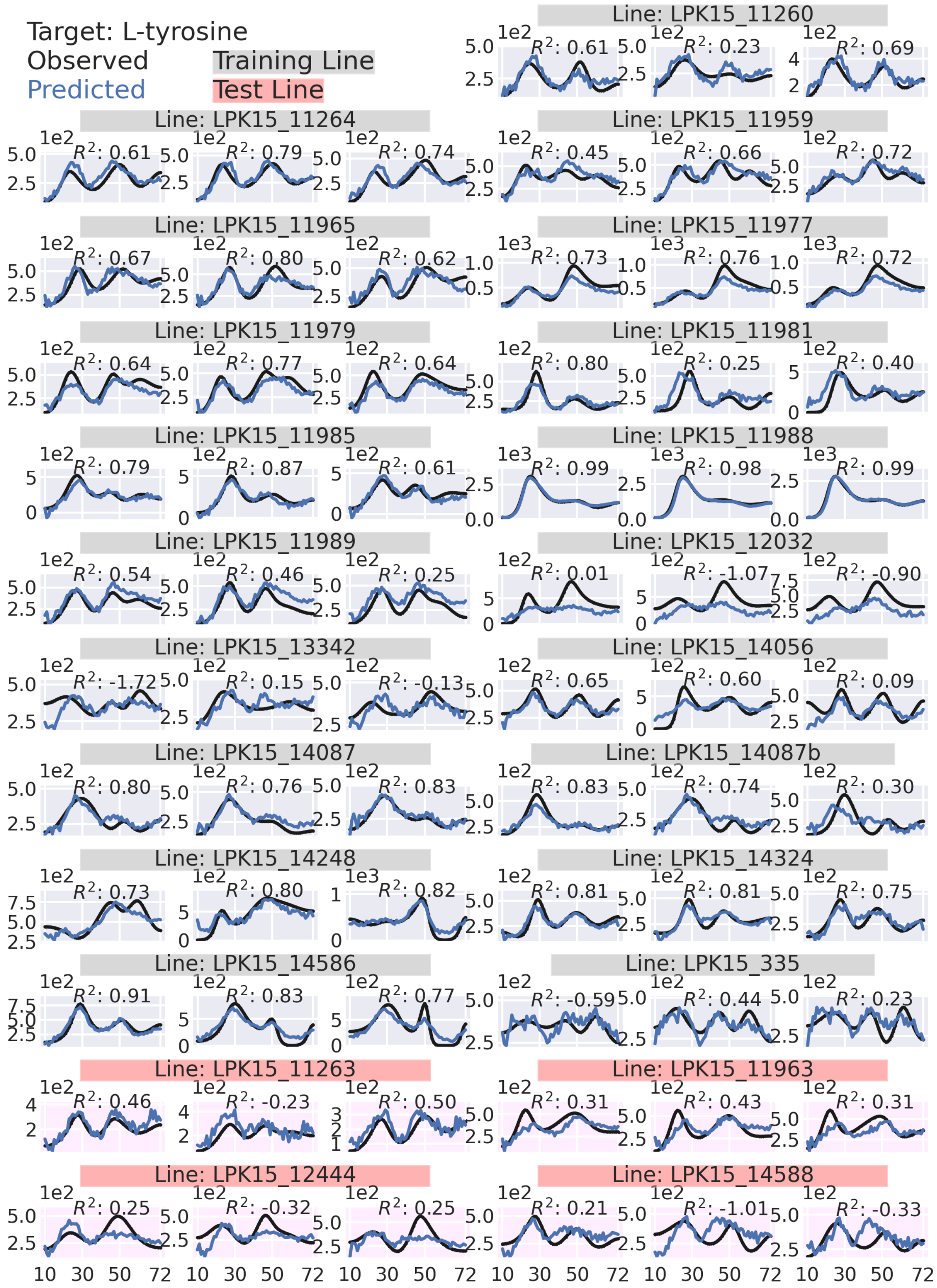

Target: L-tyrosine
Observed
Predicted
Training Line
Test Line
Line: LPK15_11260
Line: LPK15_11264
Line: LPK15_11959
Line: LPK15_11965
Line: LPK15_11977
Line: LPK15_11979
Line: LPK15_11981
Line: LPK15_11985
Line: LPK15_11988
Line: LPK15_11989
Line: LPK15_12032
Line: LPK15_13342
Line: LPK15_14056
Line: LPK15_14087
Line: LPK15_14087b
Line: LPK15_14248
Line: LPK15_14324
Line: LPK15_14586
Line: LPK15_335
Line: LPK15_11263
Line: LPK15_11963
Line: LPK15_12444
Line: LPK15_14588

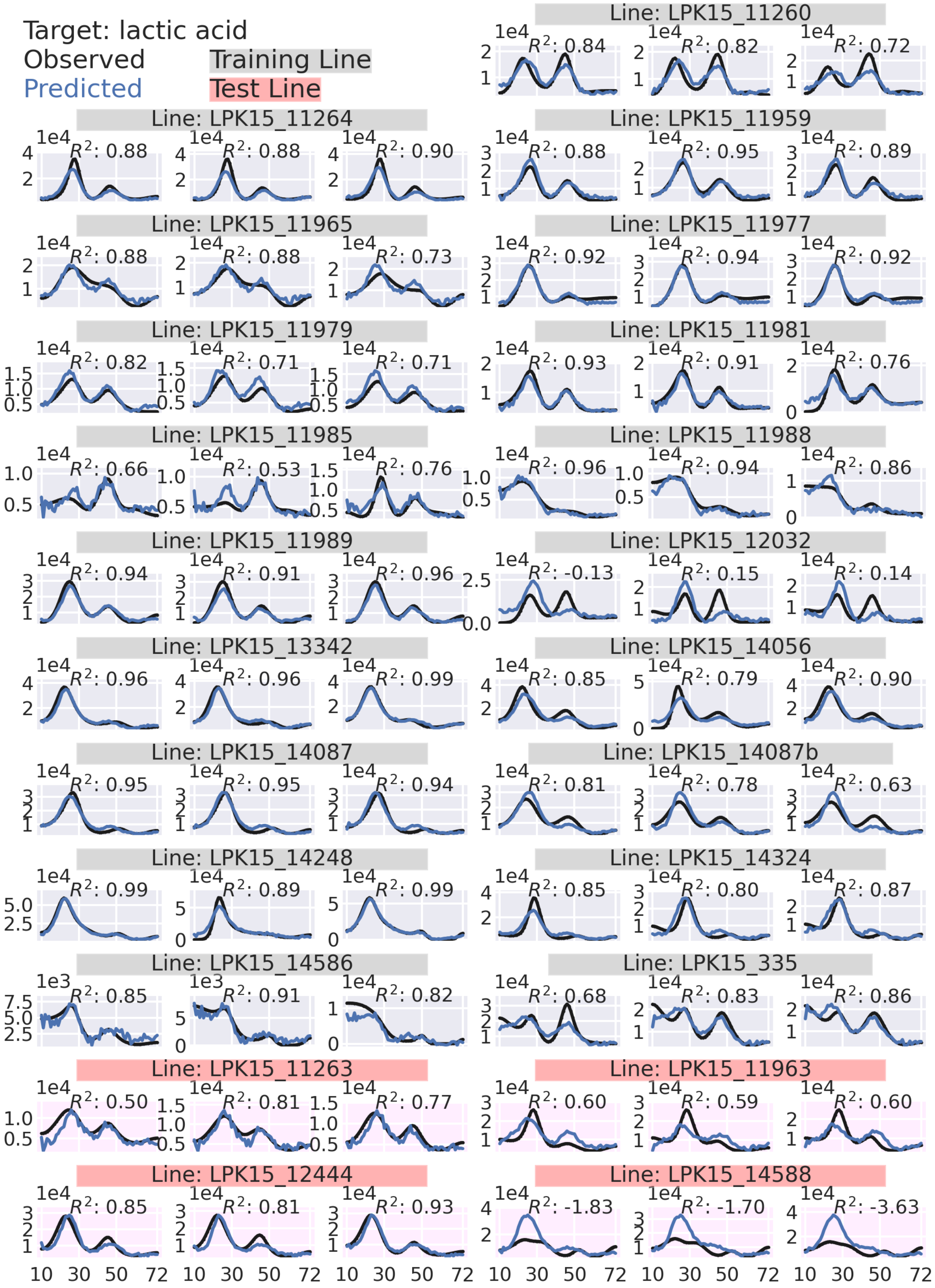

Target: lactic acid
Observed
Predicted
Training Line
Test Line
Line: LPK15_11260
Line: LPK15_11264
Line: LPK15_11959
Line: LPK15_11965
Line: LPK15_11977
Line: LPK15_11979
Line: LPK15_11981
Line: LPK15_11985
Line: LPK15_11988
Line: LPK15_11989
Line: LPK15_12032
Line: LPK15_13342
Line: LPK15_14056
Line: LPK15_14087
Line: LPK15_14087b
Line: LPK15_14248
Line: LPK15_14324
Line: LPK15_14586
Line: LPK15_335
Line: LPK15_11263
Line: LPK15_11963
Line: LPK15_12444
Line: LPK15_14588

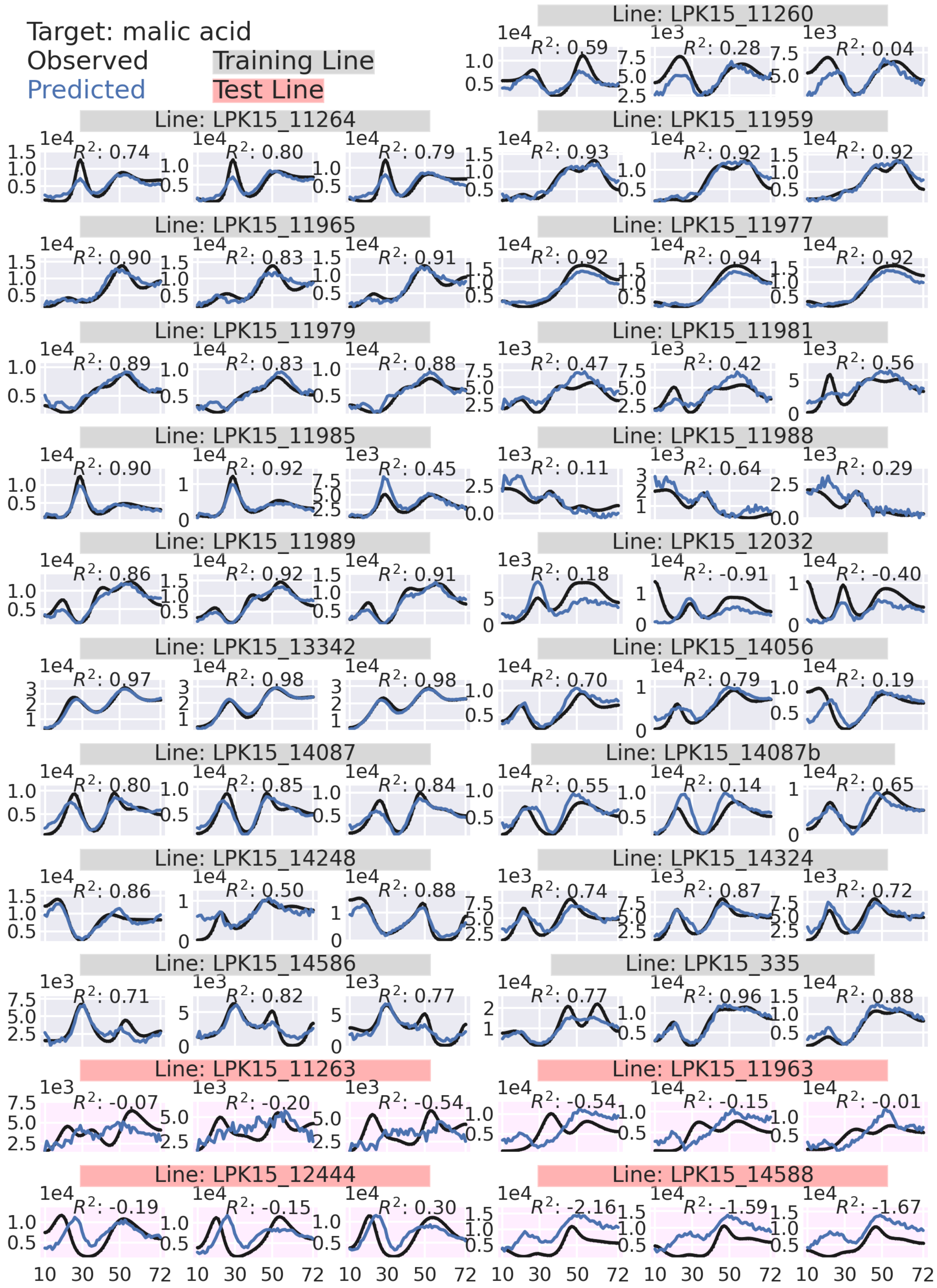
Target: malic acid
Observed
Predicted
Training Line
Test Line
Line: LPK15_11260
Line: LPK15_11264
Line: LPK15_11959
Line: LPK15_11965
Line: LPK15_11977
Line: LPK15_11979
Line: LPK15_11981
Line: LPK15_11985
Line: LPK15_11988
Line: LPK15_11989
Line: LPK15_12032
Line: LPK15_13342
Line: LPK15_14056
Line: LPK15_14087
Line: LPK15_14087b
Line: LPK15_14248
Line: LPK15_14324
Line: LPK15_14586
Line: LPK15_335
Line: LPK15_11263
Line: LPK15_11963
Line: LPK15_12444
Line: LPK15_14588
10 30 50 72

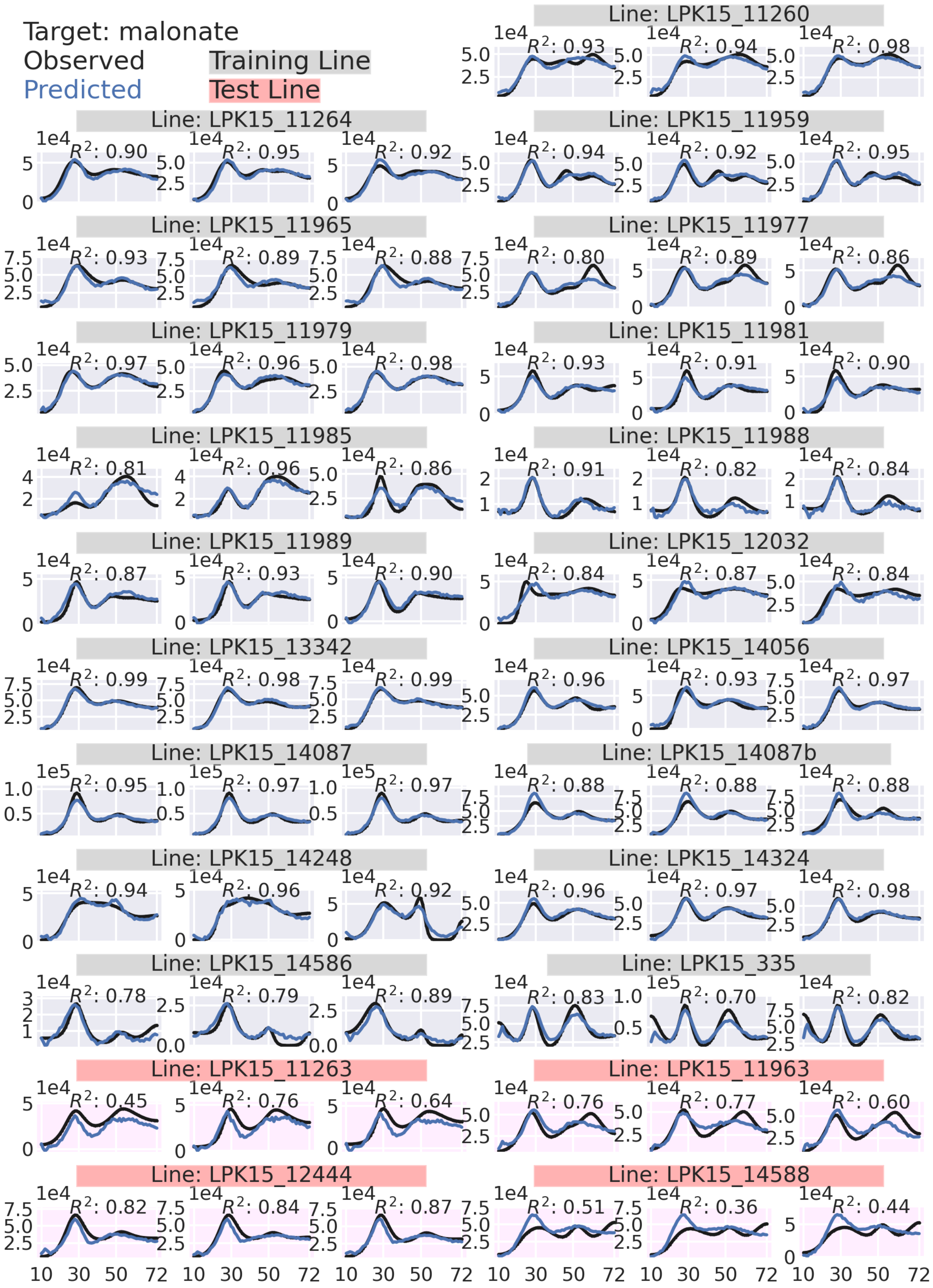

Target: malonate
Observed
Predicted
Training Line
Test Line
Line: LPK15_11260
R²: 0.93
R²: 0.94
R²: 0.98
Line: LPK15_11264
R²: 0.90
R²: 0.95
R²: 0.92
Line: LPK15_11959
R²: 0.94
R²: 0.92
R²: 0.95
Line: LPK15_11965
R²: 0.93
R²: 0.89
R²: 0.88
Line: LPK15_11977
R²: 0.80
R²: 0.89
R²: 0.86
Line: LPK15_11979
R²: 0.97
R²: 0.96
R²: 0.98
Line: LPK15_11981
R²: 0.93
R²: 0.91
R²: 0.90
Line: LPK15_11985
R²: 0.81
R²: 0.96
R²: 0.86
Line: LPK15_11988
R²: 0.91
R²: 0.82
R²: 0.84
Line: LPK15_11989
R²: 0.87
R²: 0.93
R²: 0.90
Line: LPK15_12032
R²: 0.84
R²: 0.87
R²: 0.84
Line: LPK15_13342
R²: 0.99
R²: 0.98
R²: 0.99
Line: LPK15_14056
R²: 0.96
R²: 0.93
R²: 0.97
Line: LPK15_14087
R²: 0.95
R²: 0.97
R²: 0.97
Line: LPK15_14087b
R²: 0.88
R²: 0.88
R²: 0.88
Line: LPK15_14248
R²: 0.94
R²: 0.96
R²: 0.92
Line: LPK15_14324
R²: 0.96
R²: 0.97
R²: 0.98
Line: LPK15_14586
R²: 0.78
R²: 0.79
R²: 0.89
Line: LPK15_335
R²: 0.83
R²: 0.70
R²: 0.82
Line: LPK15_11263
R²: 0.45
R²: 0.76
R²: 0.64
Line: LPK15_11963
R²: 0.76
R²: 0.77
R²: 0.60
Line: LPK15_12444
R²: 0.82
R²: 0.84
R²: 0.87
Line: LPK15_14588
R²: 0.51
R²: 0.36
R²: 0.44
10 30 50 72

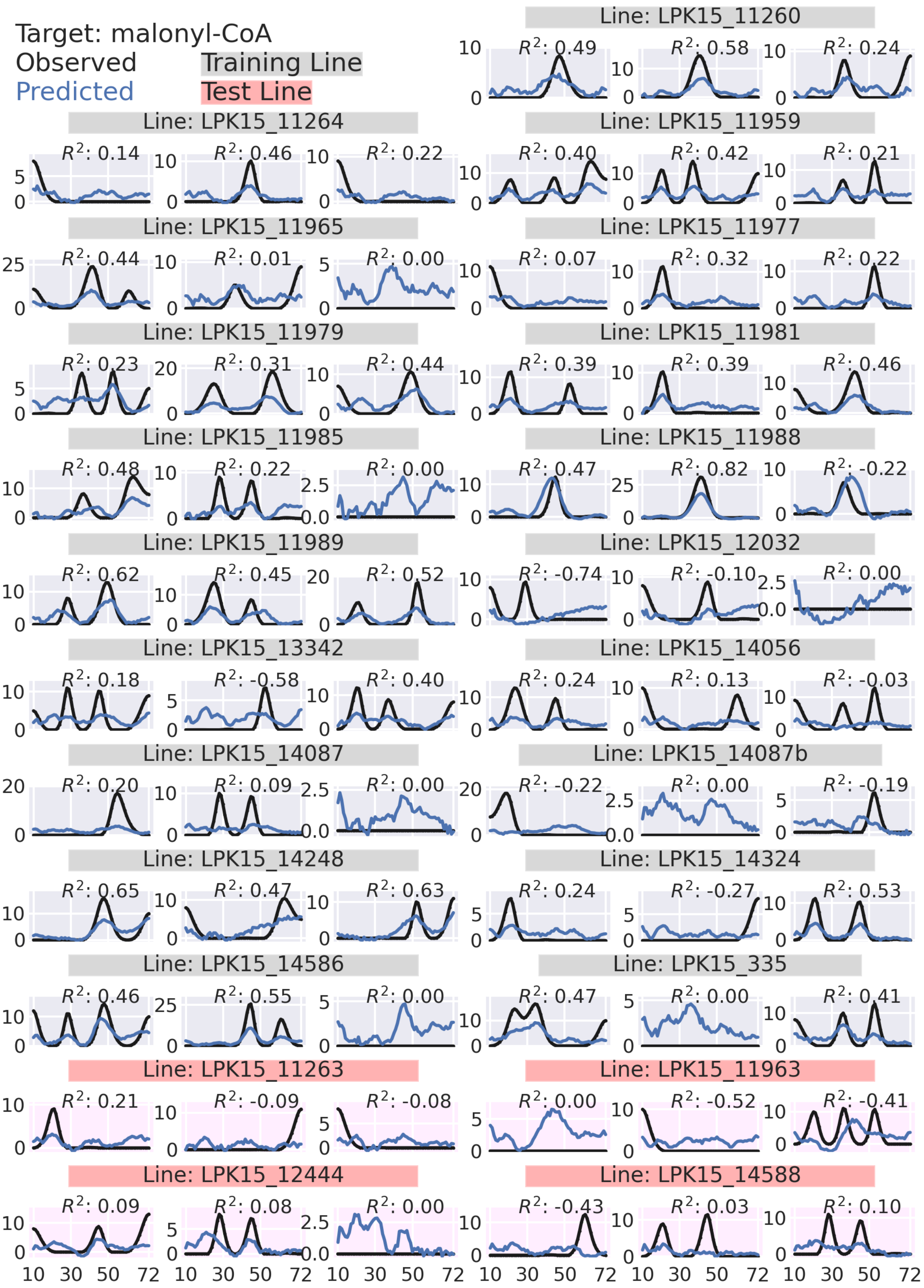

Target: malonyl-CoA
Observed
Predicted
Training Line
Test Line
Line: LPK15_11260
Line: LPK15_11264
Line: LPK15_11959
Line: LPK15_11965
Line: LPK15_11977
Line: LPK15_11979
Line: LPK15_11981
Line: LPK15_11985
Line: LPK15_11988
Line: LPK15_11989
Line: LPK15_12032
Line: LPK15_13342
Line: LPK15_14056
Line: LPK15_14087
Line: LPK15_14087b
Line: LPK15_14248
Line: LPK15_14324
Line: LPK15_14586
Line: LPK15_335
Line: LPK15_11263
Line: LPK15_11963
Line: LPK15_12444
Line: LPK15_14588

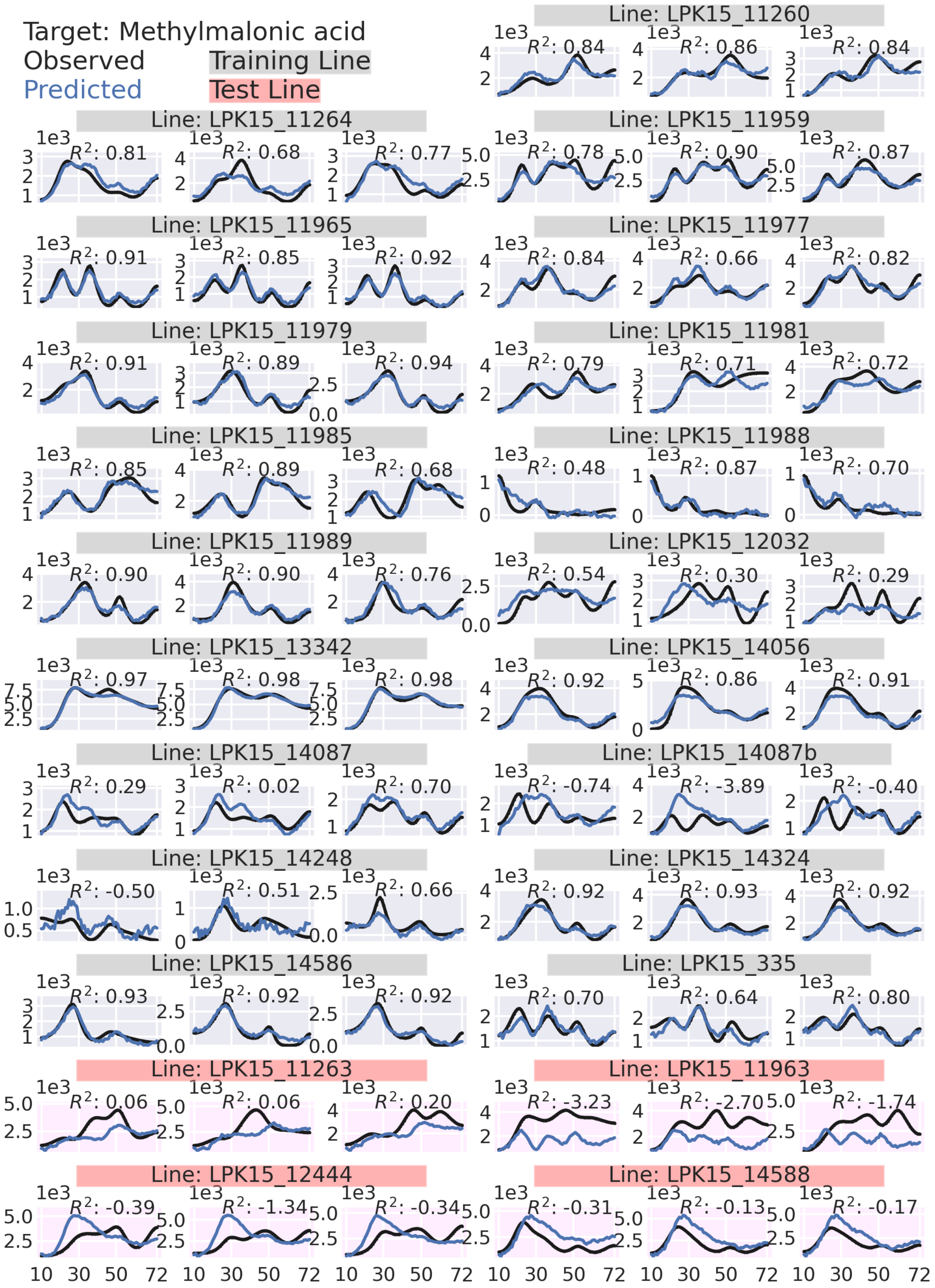

Target: Methylmalonic acid
Observed
Predicted
Training Line
Test Line
Line: LPK15_11260
R²: 0.84 R²: 0.86 R²: 0.84
Line: LPK15_11264
R²: 0.81 R²: 0.68 R²: 0.77
Line: LPK15_11959
R²: 0.78 R²: 0.90 R²: 0.87
Line: LPK15_11965
R²: 0.91 R²: 0.85 R²: 0.92
Line: LPK15_11977
R²: 0.84 R²: 0.66 R²: 0.82
Line: LPK15_11979
R²: 0.91 R²: 0.89 R²: 0.94
Line: LPK15_11981
R²: 0.79 R²: 0.71 R²: 0.72
Line: LPK15_11985
R²: 0.85 R²: 0.89 R²: 0.68
Line: LPK15_11988
R²: 0.48 R²: 0.87 R²: 0.70
Line: LPK15_11989
R²: 0.90 R²: 0.90 R²: 0.76
Line: LPK15_12032
R²: 0.54 R²: 0.30 R²: 0.29
Line: LPK15_13342
R²: 0.97 R²: 0.98 R²: 0.98
Line: LPK15_14056
R²: 0.92 R²: 0.86 R²: 0.91
Line: LPK15_14087
R²: 0.29 R²: 0.02 R²: 0.70
Line: LPK15_14087b
R²: -0.74 R²: -3.89 R²: -0.40
Line: LPK15_14248
R²: -0.50 R²: 0.51 R²: 0.66
Line: LPK15_14324
R²: 0.92 R²: 0.93 R²: 0.92
Line: LPK15_14586
R²: 0.93 R²: 0.92 R²: 0.92
Line: LPK15_335
R²: 0.70 R²: 0.64 R²: 0.80
Line: LPK15_11263
R²: 0.06 R²: 0.06 R²: 0.20
Line: LPK15_11963
R²: -3.23 R²: -2.70 R²: -1.74
Line: LPK15_12444
R²: -0.39 R²: -1.34 R²: -0.34
Line: LPK15_14588
R²: -0.31 R²: -0.13 R²: -0.17
1e3
10 30 50 72

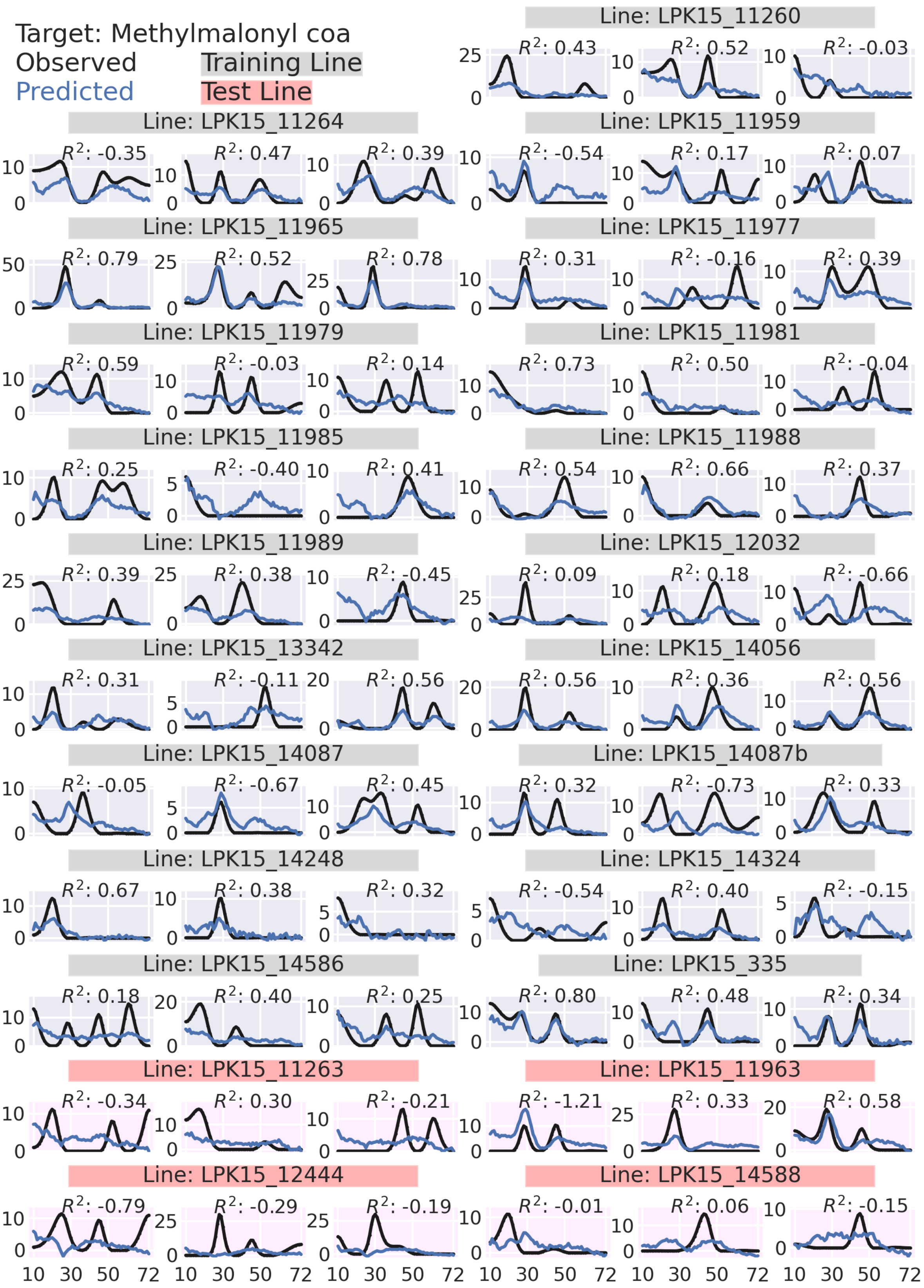

Target: Methylmalonyl coa
Observed
Predicted
Training Line
Test Line
Line: LPK15_11260
Line: LPK15_11264
Line: LPK15_11959
Line: LPK15_11965
Line: LPK15_11977
Line: LPK15_11979
Line: LPK15_11981
Line: LPK15_11985
Line: LPK15_11988
Line: LPK15_11989
Line: LPK15_12032
Line: LPK15_13342
Line: LPK15_14056
Line: LPK15_14087
Line: LPK15_14087b
Line: LPK15_14248
Line: LPK15_14324
Line: LPK15_14586
Line: LPK15_335
Line: LPK15_11263
Line: LPK15_11963
Line: LPK15_12444
Line: LPK15_14588

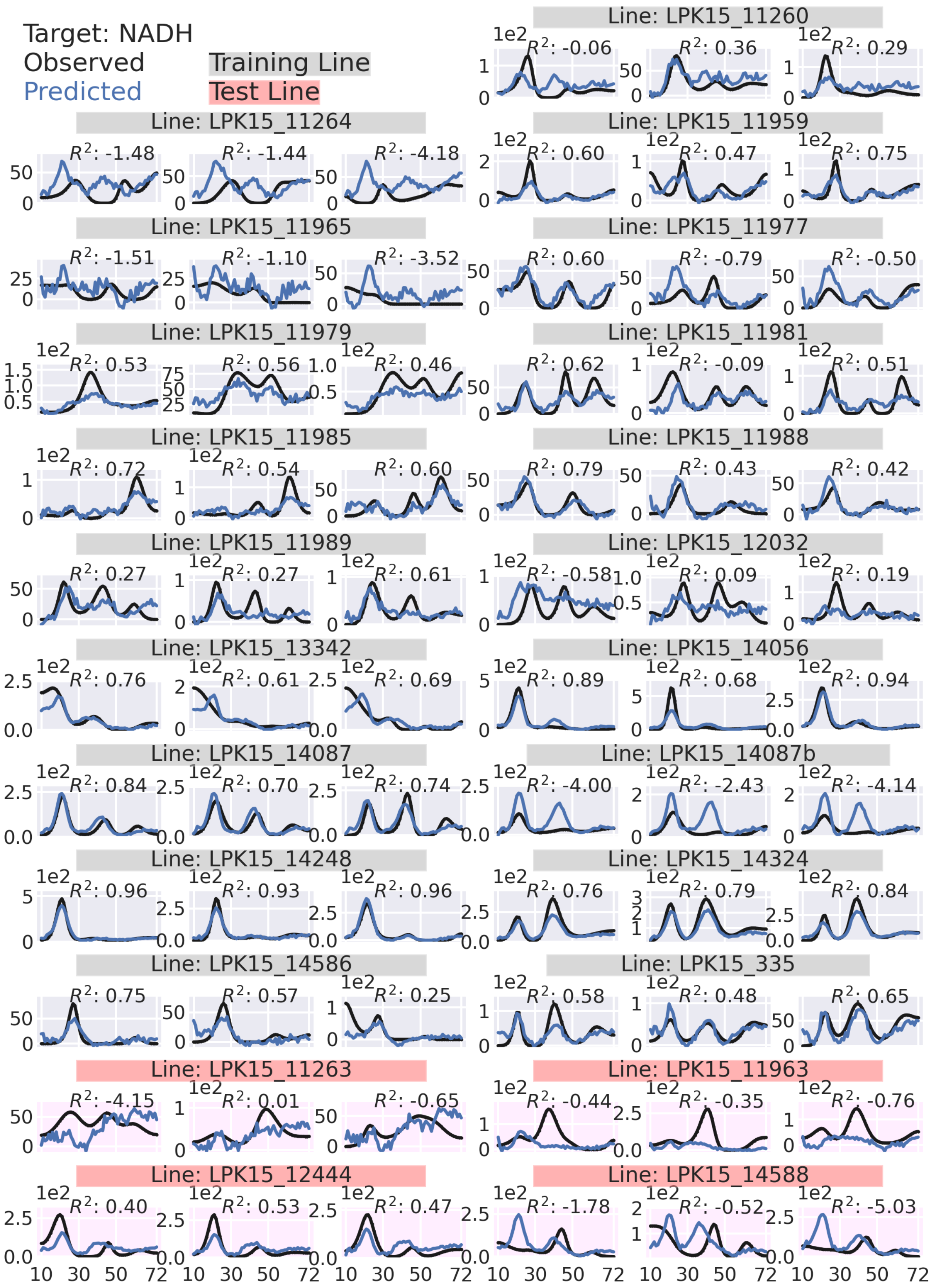

Target: NADH
Observed
Predicted
Training Line
Test Line
Line: LPK15_11260
Line: LPK15_11264
Line: LPK15_11959
Line: LPK15_11965
Line: LPK15_11977
Line: LPK15_11979
Line: LPK15_11981
Line: LPK15_11985
Line: LPK15_11988
Line: LPK15_11989
Line: LPK15_12032
Line: LPK15_13342
Line: LPK15_14056
Line: LPK15_14087
Line: LPK15_14087b
Line: LPK15_14248
Line: LPK15_14324
Line: LPK15_14586
Line: LPK15_335
Line: LPK15_11263
Line: LPK15_11963
Line: LPK15_12444
Line: LPK15_14588

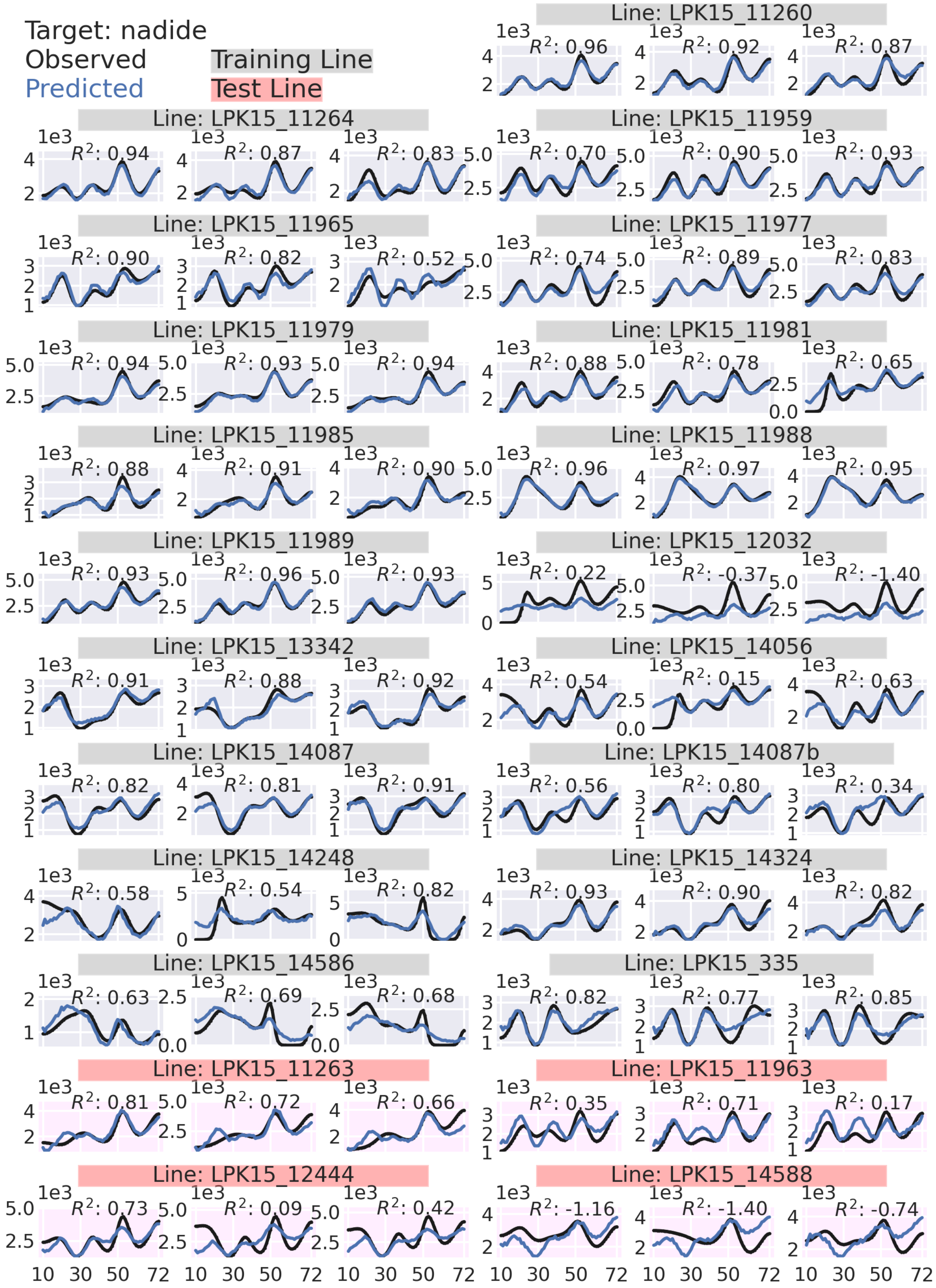

Target: nadide
Observed
Predicted
Training Line
Test Line
Line: LPK15_11260
Line: LPK15_11264
Line: LPK15_11959
Line: LPK15_11965
Line: LPK15_11977
Line: LPK15_11979
Line: LPK15_11981
Line: LPK15_11985
Line: LPK15_11988
Line: LPK15_11989
Line: LPK15_12032
Line: LPK15_13342
Line: LPK15_14056
Line: LPK15_14087
Line: LPK15_14087b
Line: LPK15_14248
Line: LPK15_14324
Line: LPK15_14586
Line: LPK15_335
Line: LPK15_11263
Line: LPK15_11963
Line: LPK15_12444
Line: LPK15_14588

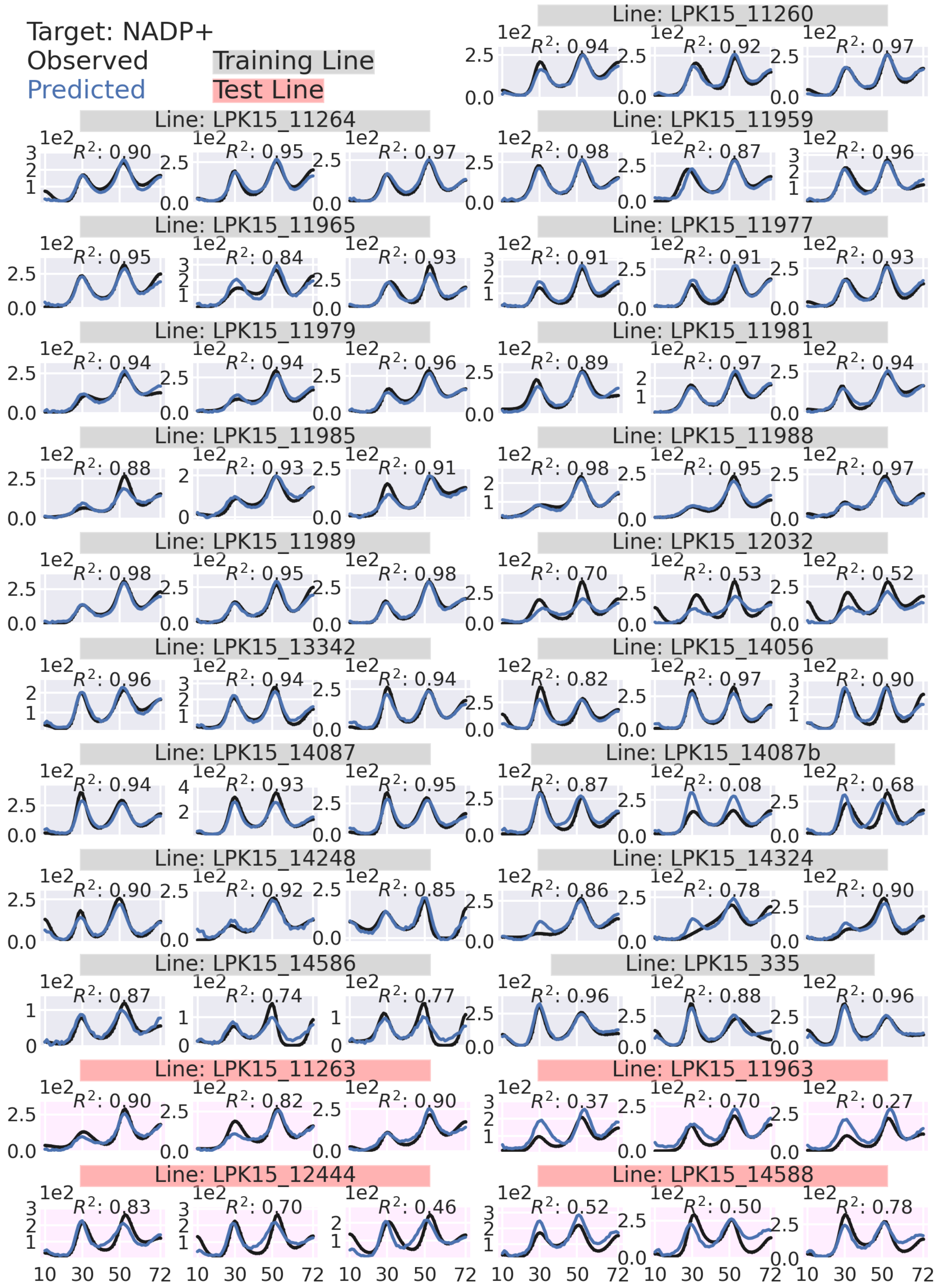
Target: NADP+
Observed
Predicted
Training Line
Test Line
Line: LPK15_11260
Line: LPK15_11264
Line: LPK15_11959
Line: LPK15_11965
Line: LPK15_11977
Line: LPK15_11979
Line: LPK15_11981
Line: LPK15_11985
Line: LPK15_11988
Line: LPK15_11989
Line: LPK15_12032
Line: LPK15_13342
Line: LPK15_14056
Line: LPK15_14087
Line: LPK15_14087b
Line: LPK15_14248
Line: LPK15_14324
Line: LPK15_14586
Line: LPK15_335
Line: LPK15_11263
Line: LPK15_11963
Line: LPK15_12444
Line: LPK15_14588

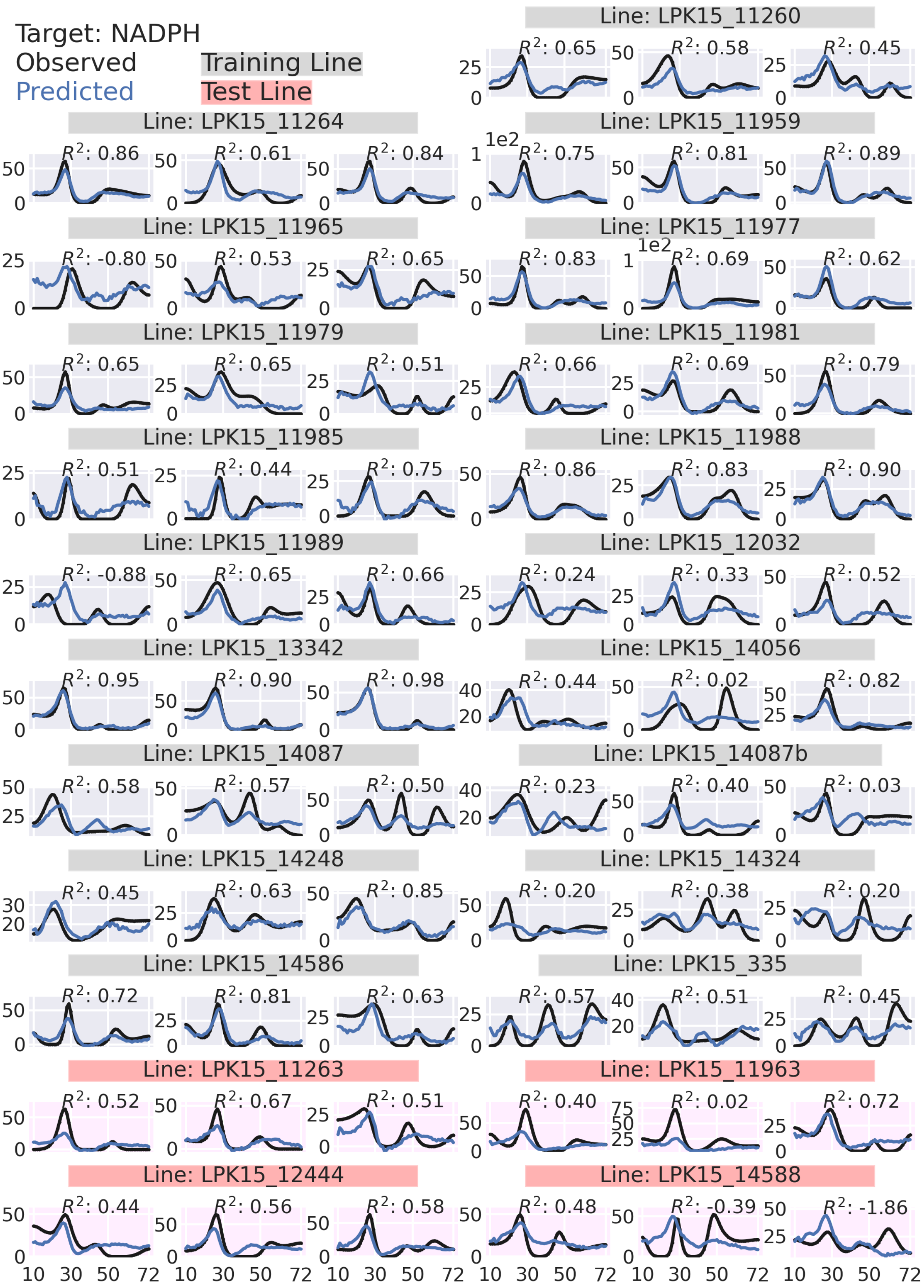
Target: NADPH
Observed
Predicted
Training Line
Test Line
Line: LPK15_11260
Line: LPK15_11264
Line: LPK15_11959
Line: LPK15_11965
Line: LPK15_11977
Line: LPK15_11979
Line: LPK15_11981
Line: LPK15_11985
Line: LPK15_11988
Line: LPK15_11989
Line: LPK15_12032
Line: LPK15_13342
Line: LPK15_14056
Line: LPK15_14087
Line: LPK15_14087b
Line: LPK15_14248
Line: LPK15_14324
Line: LPK15_14586
Line: LPK15_335
Line: LPK15_11263
Line: LPK15_11963
Line: LPK15_12444
Line: LPK15_14588

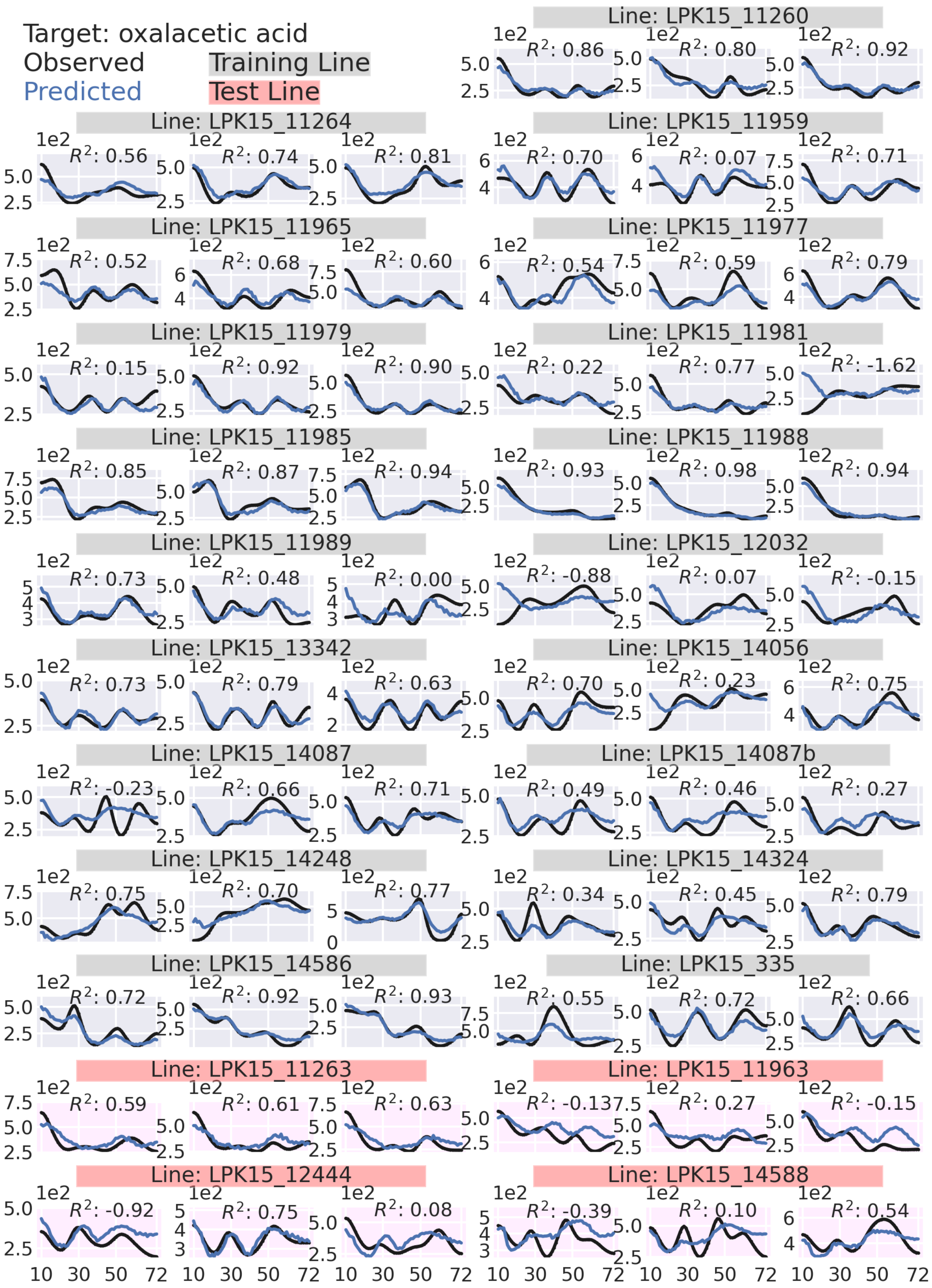

Target: oxalacetic acid
Observed
Predicted
Training Line
Test Line
Line: LPK15_11260
$R^2$: 0.86
$R^2$: 0.80
$R^2$: 0.92
Line: LPK15_11264
$R^2$: 0.56
$R^2$: 0.74
$R^2$: 0.81
Line: LPK15_11959
$R^2$: 0.70
$R^2$: 0.07
$R^2$: 0.71
Line: LPK15_11965
$R^2$: 0.52
$R^2$: 0.68
$R^2$: 0.60
Line: LPK15_11977
$R^2$: 0.54
$R^2$: 0.59
$R^2$: 0.79
Line: LPK15_11979
$R^2$: 0.15
$R^2$: 0.92
$R^2$: 0.90
Line: LPK15_11981
$R^2$: 0.22
$R^2$: 0.77
$R^2$: -1.62
Line: LPK15_11985
$R^2$: 0.85
$R^2$: 0.87
$R^2$: 0.94
Line: LPK15_11988
$R^2$: 0.93
$R^2$: 0.98
$R^2$: 0.94
Line: LPK15_11989
$R^2$: 0.73
$R^2$: 0.48
$R^2$: 0.00
Line: LPK15_12032
$R^2$: -0.88
$R^2$: 0.07
$R^2$: -0.15
Line: LPK15_13342
$R^2$: 0.73
$R^2$: 0.79
$R^2$: 0.63
Line: LPK15_14056
$R^2$: 0.70
$R^2$: 0.23
$R^2$: 0.75
Line: LPK15_14087
$R^2$: -0.23
$R^2$: 0.66
$R^2$: 0.71
Line: LPK15_14087b
$R^2$: 0.49
$R^2$: 0.46
$R^2$: 0.27
Line: LPK15_14248
$R^2$: 0.75
$R^2$: 0.70
$R^2$: 0.77
Line: LPK15_14324
$R^2$: 0.34
$R^2$: 0.45
$R^2$: 0.79
Line: LPK15_14586
$R^2$: 0.72
$R^2$: 0.92
$R^2$: 0.93
Line: LPK15_335
$R^2$: 0.55
$R^2$: 0.72
$R^2$: 0.66
Line: LPK15_11263
$R^2$: 0.59
$R^2$: 0.61
$R^2$: 0.63
Line: LPK15_11963
$R^2$: -0.13
$R^2$: 0.27
$R^2$: -0.15
Line: LPK15_12444
$R^2$: -0.92
$R^2$: 0.75
$R^2$: 0.08
Line: LPK15_14588
$R^2$: -0.39
$R^2$: 0.10
$R^2$: 0.54
1e2
10 30 50 72

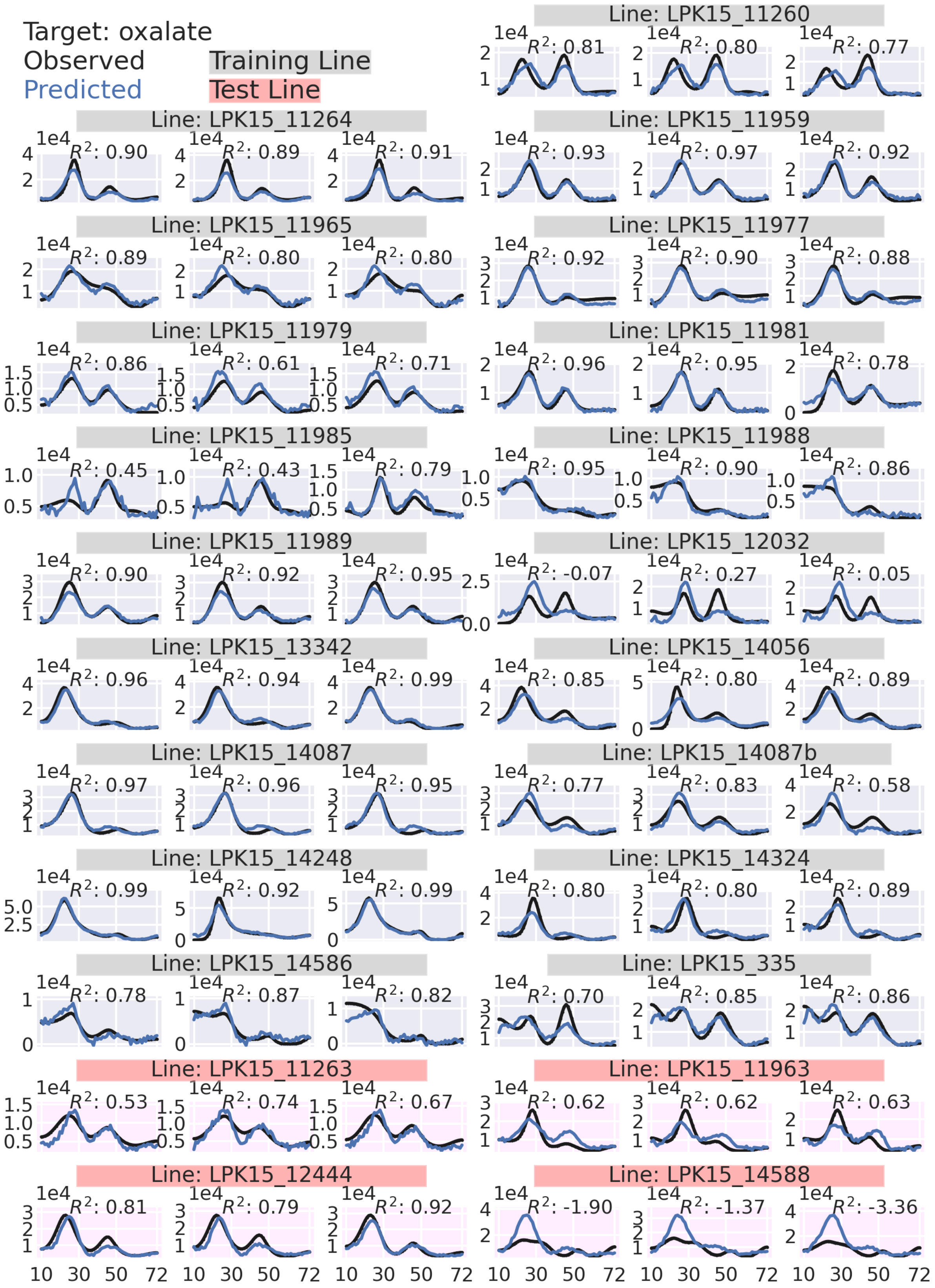

Target: oxalate
Observed
Predicted
Training Line
Test Line
Line: LPK15_11260
R²: 0.81 R²: 0.80 R²: 0.77
Line: LPK15_11264
R²: 0.90 R²: 0.89 R²: 0.91
Line: LPK15_11959
R²: 0.93 R²: 0.97 R²: 0.92
Line: LPK15_11965
R²: 0.89 R²: 0.80 R²: 0.80
Line: LPK15_11977
R²: 0.92 R²: 0.90 R²: 0.88
Line: LPK15_11979
R²: 0.86 R²: 0.61 R²: 0.71
Line: LPK15_11981
R²: 0.96 R²: 0.95 R²: 0.78
Line: LPK15_11985
R²: 0.45 R²: 0.43 R²: 0.79
Line: LPK15_11988
R²: 0.95 R²: 0.90 R²: 0.86
Line: LPK15_11989
R²: 0.90 R²: 0.92 R²: 0.95
Line: LPK15_12032
R²: -0.07 R²: 0.27 R²: 0.05
Line: LPK15_13342
R²: 0.96 R²: 0.94 R²: 0.99
Line: LPK15_14056
R²: 0.85 R²: 0.80 R²: 0.89
Line: LPK15_14087
R²: 0.97 R²: 0.96 R²: 0.95
Line: LPK15_14087b
R²: 0.77 R²: 0.83 R²: 0.58
Line: LPK15_14248
R²: 0.99 R²: 0.92 R²: 0.99
Line: LPK15_14324
R²: 0.80 R²: 0.80 R²: 0.89
Line: LPK15_14586
R²: 0.78 R²: 0.87 R²: 0.82
Line: LPK15_335
R²: 0.70 R²: 0.85 R²: 0.86
Line: LPK15_11263
R²: 0.53 R²: 0.74 R²: 0.67
Line: LPK15_11963
R²: 0.62 R²: 0.62 R²: 0.63
Line: LPK15_12444
R²: 0.81 R²: 0.79 R²: 0.92
Line: LPK15_14588
R²: -1.90 R²: -1.37 R²: -3.36
1e4
10 30 50 72

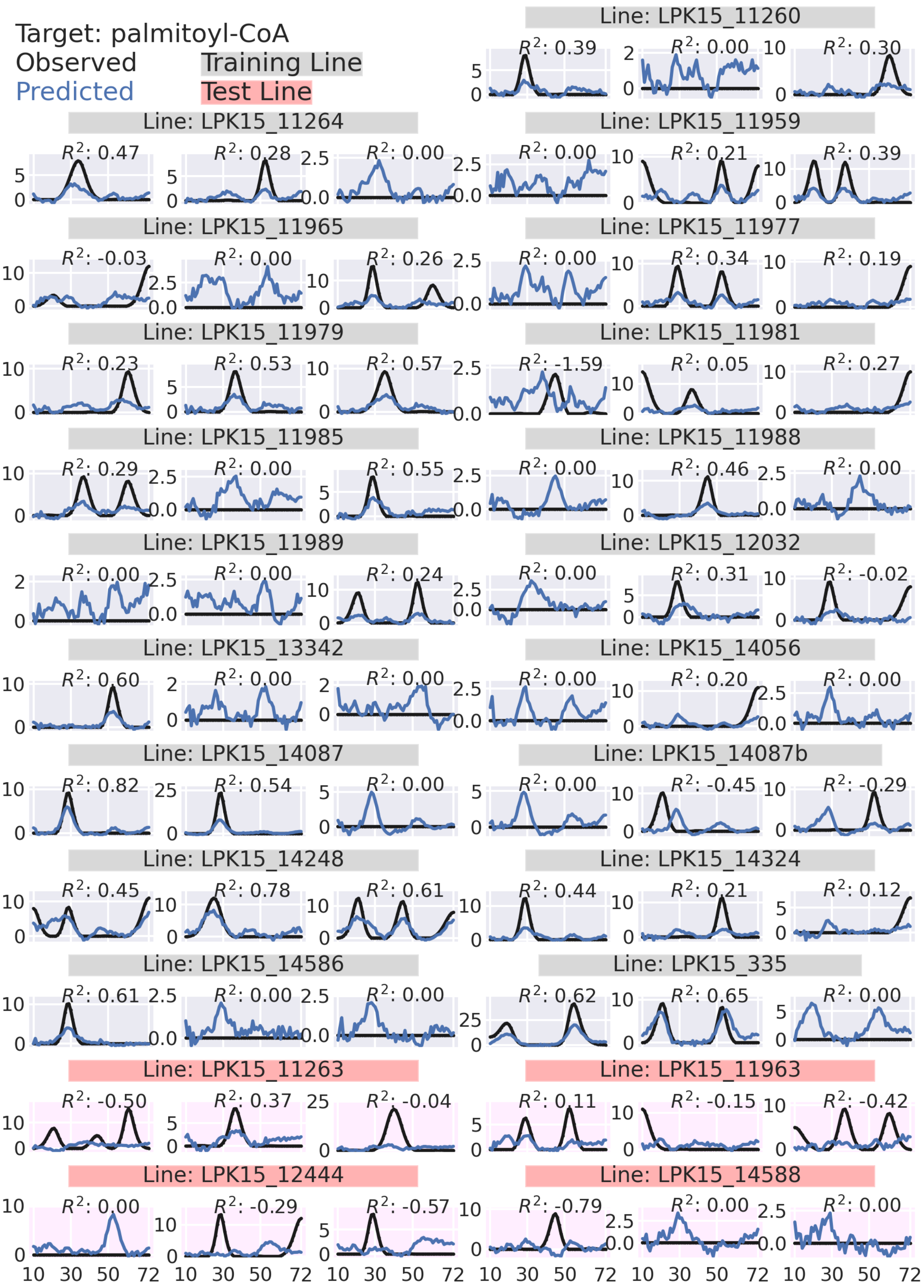
Target: palmitoyl-CoA
Observed
Predicted
Training Line
Test Line
Line: LPK15_11260
Line: LPK15_11264
Line: LPK15_11959
Line: LPK15_11965
Line: LPK15_11977
Line: LPK15_11979
Line: LPK15_11981
Line: LPK15_11985
Line: LPK15_11988
Line: LPK15_11989
Line: LPK15_12032
Line: LPK15_13342
Line: LPK15_14056
Line: LPK15_14087
Line: LPK15_14087b
Line: LPK15_14248
Line: LPK15_14324
Line: LPK15_14586
Line: LPK15_335
Line: LPK15_11263
Line: LPK15_11963
Line: LPK15_12444
Line: LPK15_14588

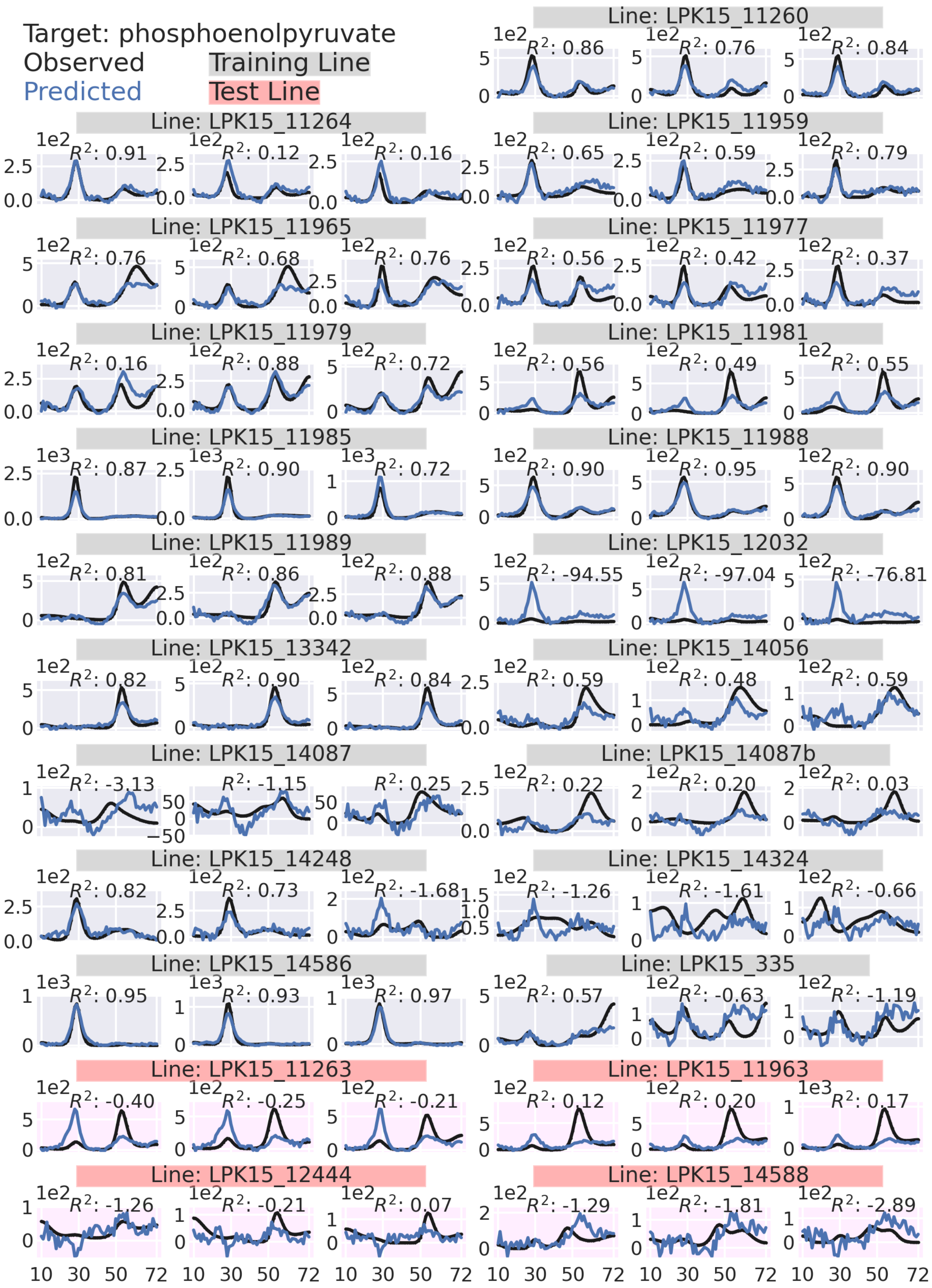

Target: phosphoenolpyruvate
Observed
Predicted
Training Line
Test Line
Line: LPK15_11260
Line: LPK15_11264
Line: LPK15_11959
Line: LPK15_11965
Line: LPK15_11977
Line: LPK15_11979
Line: LPK15_11981
Line: LPK15_11985
Line: LPK15_11988
Line: LPK15_11989
Line: LPK15_12032
Line: LPK15_13342
Line: LPK15_14056
Line: LPK15_14087
Line: LPK15_14087b
Line: LPK15_14248
Line: LPK15_14324
Line: LPK15_14586
Line: LPK15_335
Line: LPK15_11263
Line: LPK15_11963
Line: LPK15_12444
Line: LPK15_14588

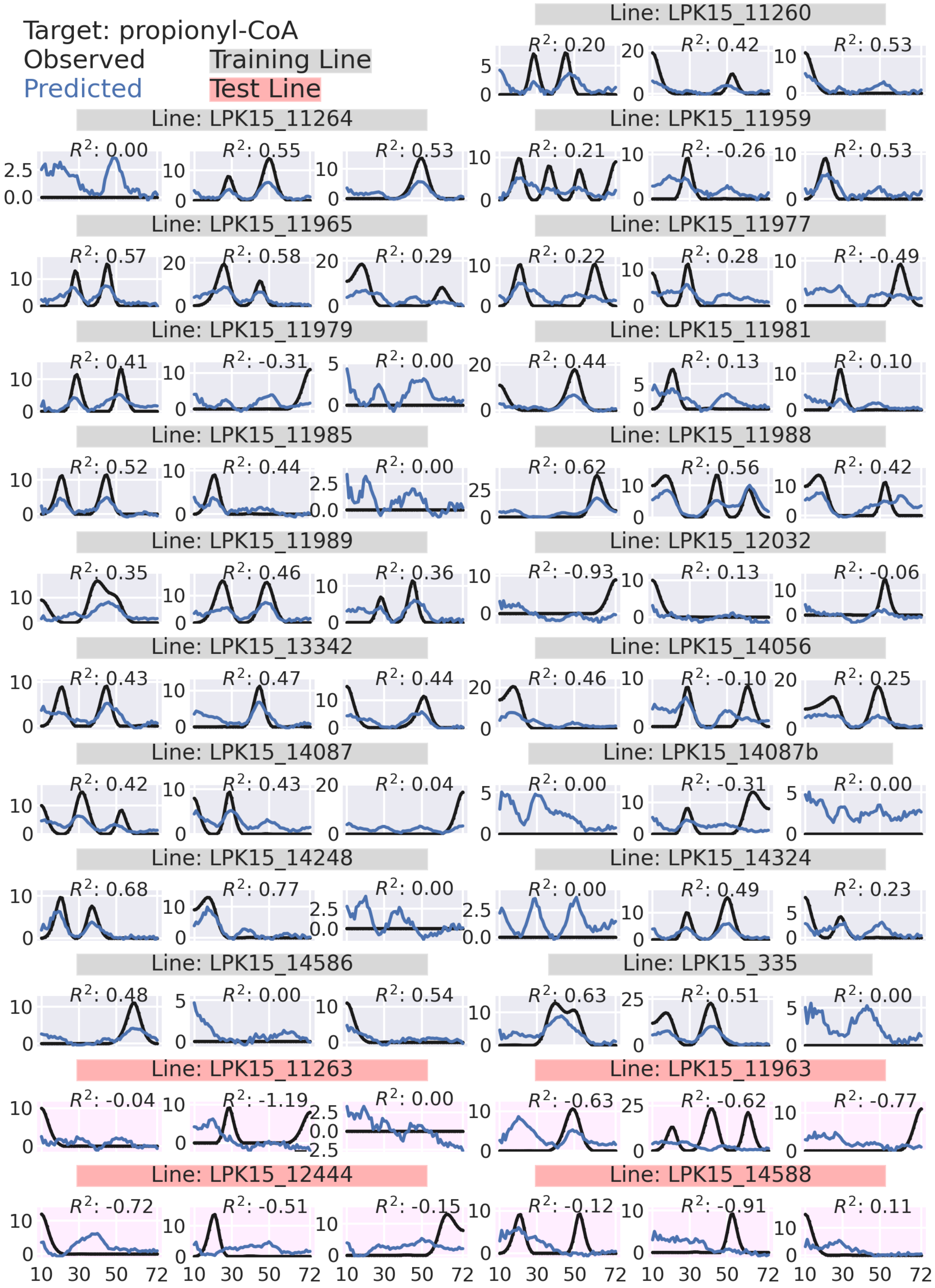
Target: propionyl-CoA
Observed
Predicted
Training Line
Test Line
Line: LPK15_11260
Line: LPK15_11264
Line: LPK15_11959
Line: LPK15_11965
Line: LPK15_11977
Line: LPK15_11979
Line: LPK15_11981
Line: LPK15_11985
Line: LPK15_11988
Line: LPK15_11989
Line: LPK15_12032
Line: LPK15_13342
Line: LPK15_14056
Line: LPK15_14087
Line: LPK15_14087b
Line: LPK15_14248
Line: LPK15_14324
Line: LPK15_14586
Line: LPK15_335
Line: LPK15_11263
Line: LPK15_11963
Line: LPK15_12444
Line: LPK15_14588

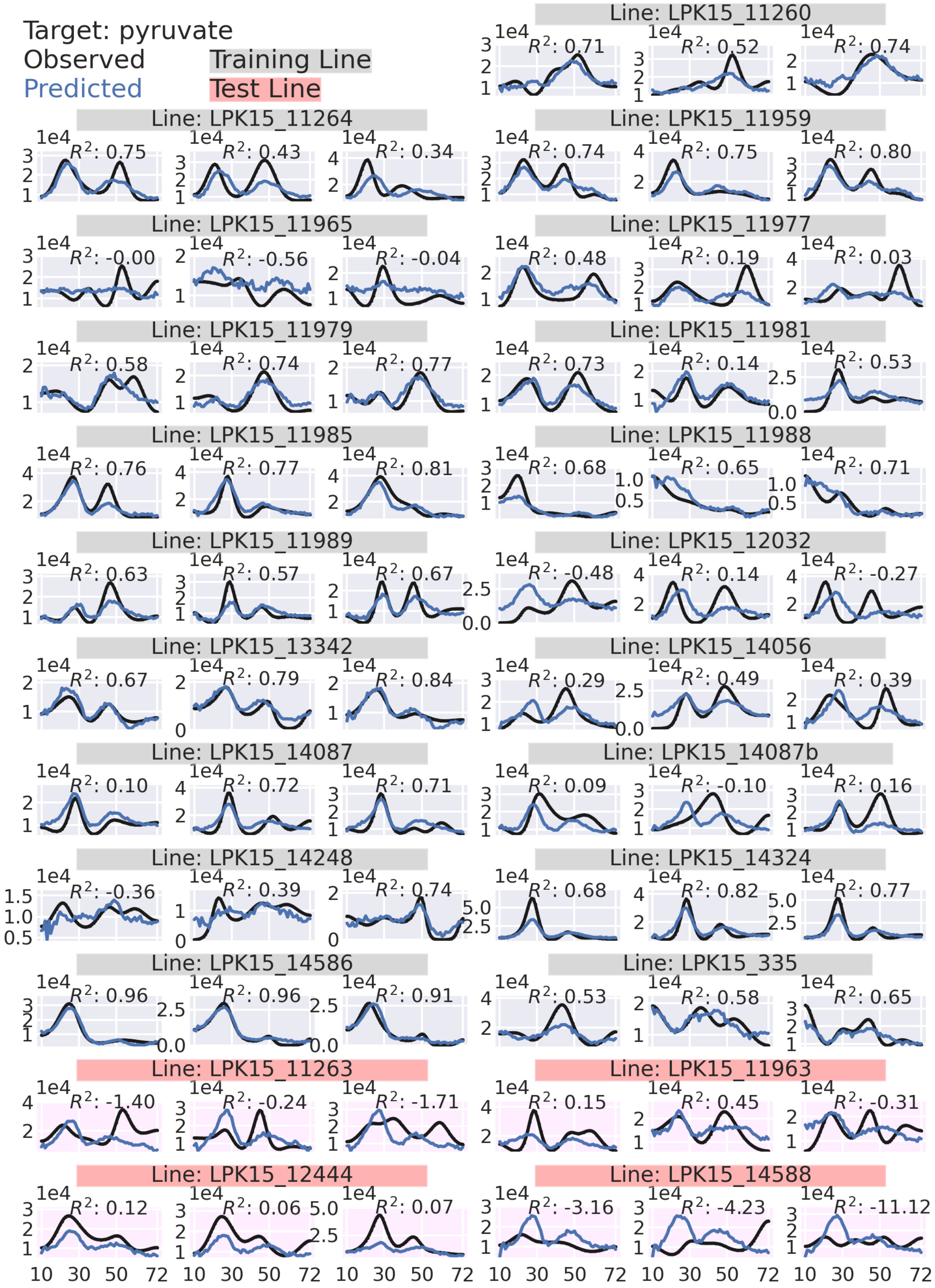

Target: pyruvate
Observed
Predicted
Training Line
Test Line
Line: LPK15_11260
Line: LPK15_11264
Line: LPK15_11959
Line: LPK15_11965
Line: LPK15_11977
Line: LPK15_11979
Line: LPK15_11981
Line: LPK15_11985
Line: LPK15_11988
Line: LPK15_11989
Line: LPK15_12032
Line: LPK15_13342
Line: LPK15_14056
Line: LPK15_14087
Line: LPK15_14087b
Line: LPK15_14248
Line: LPK15_14324
Line: LPK15_14586
Line: LPK15_335
Line: LPK15_11263
Line: LPK15_11963
Line: LPK15_12444
Line: LPK15_14588

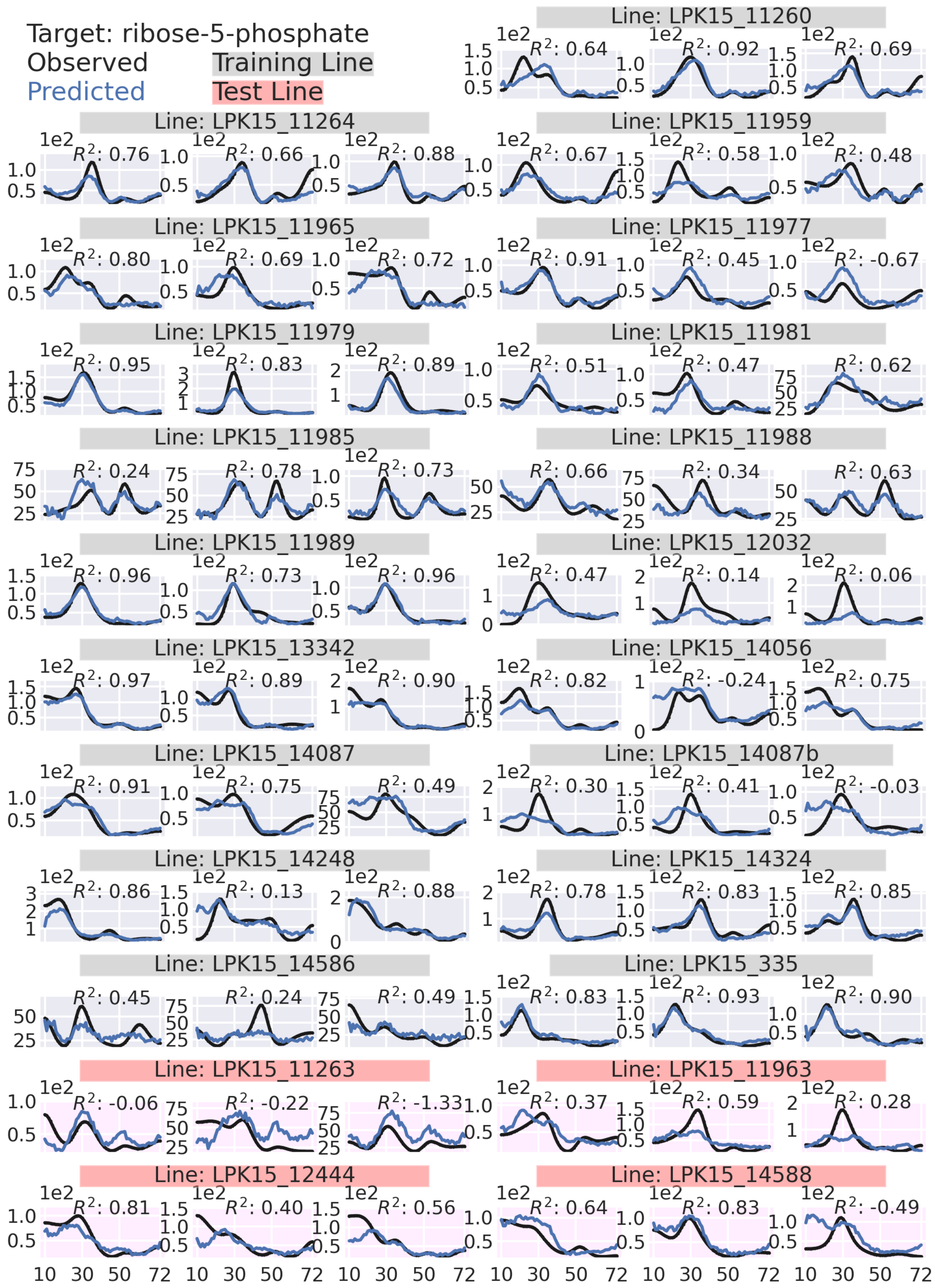

Target: ribose-5-phosphate
Observed
Predicted
Training Line
Test Line
Line: LPK15_11260
Line: LPK15_11264
Line: LPK15_11959
Line: LPK15_11965
Line: LPK15_11977
Line: LPK15_11979
Line: LPK15_11981
Line: LPK15_11985
Line: LPK15_11988
Line: LPK15_11989
Line: LPK15_12032
Line: LPK15_13342
Line: LPK15_14056
Line: LPK15_14087
Line: LPK15_14087b
Line: LPK15_14248
Line: LPK15_14324
Line: LPK15_14586
Line: LPK15_335
Line: LPK15_11263
Line: LPK15_11963
Line: LPK15_12444
Line: LPK15_14588

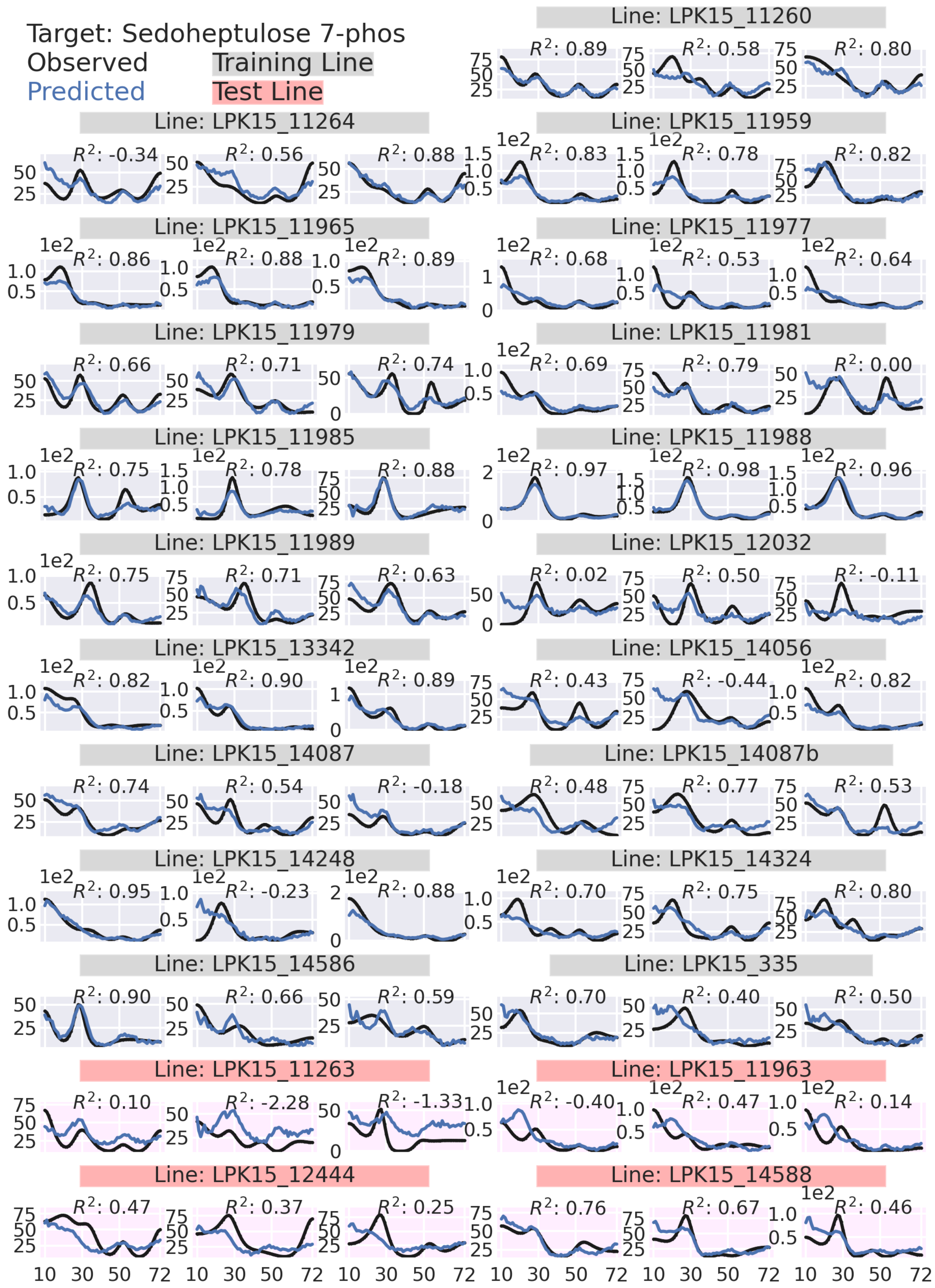

Target: Sedoheptulose 7-phos
Observed
Predicted
Training Line
Test Line
Line: LPK15_11260
Line: LPK15_11264
Line: LPK15_11959
Line: LPK15_11965
Line: LPK15_11977
Line: LPK15_11979
Line: LPK15_11981
Line: LPK15_11985
Line: LPK15_11988
Line: LPK15_11989
Line: LPK15_12032
Line: LPK15_13342
Line: LPK15_14056
Line: LPK15_14087
Line: LPK15_14087b
Line: LPK15_14248
Line: LPK15_14324
Line: LPK15_14586
Line: LPK15_335
Line: LPK15_11263
Line: LPK15_11963
Line: LPK15_12444
Line: LPK15_14588

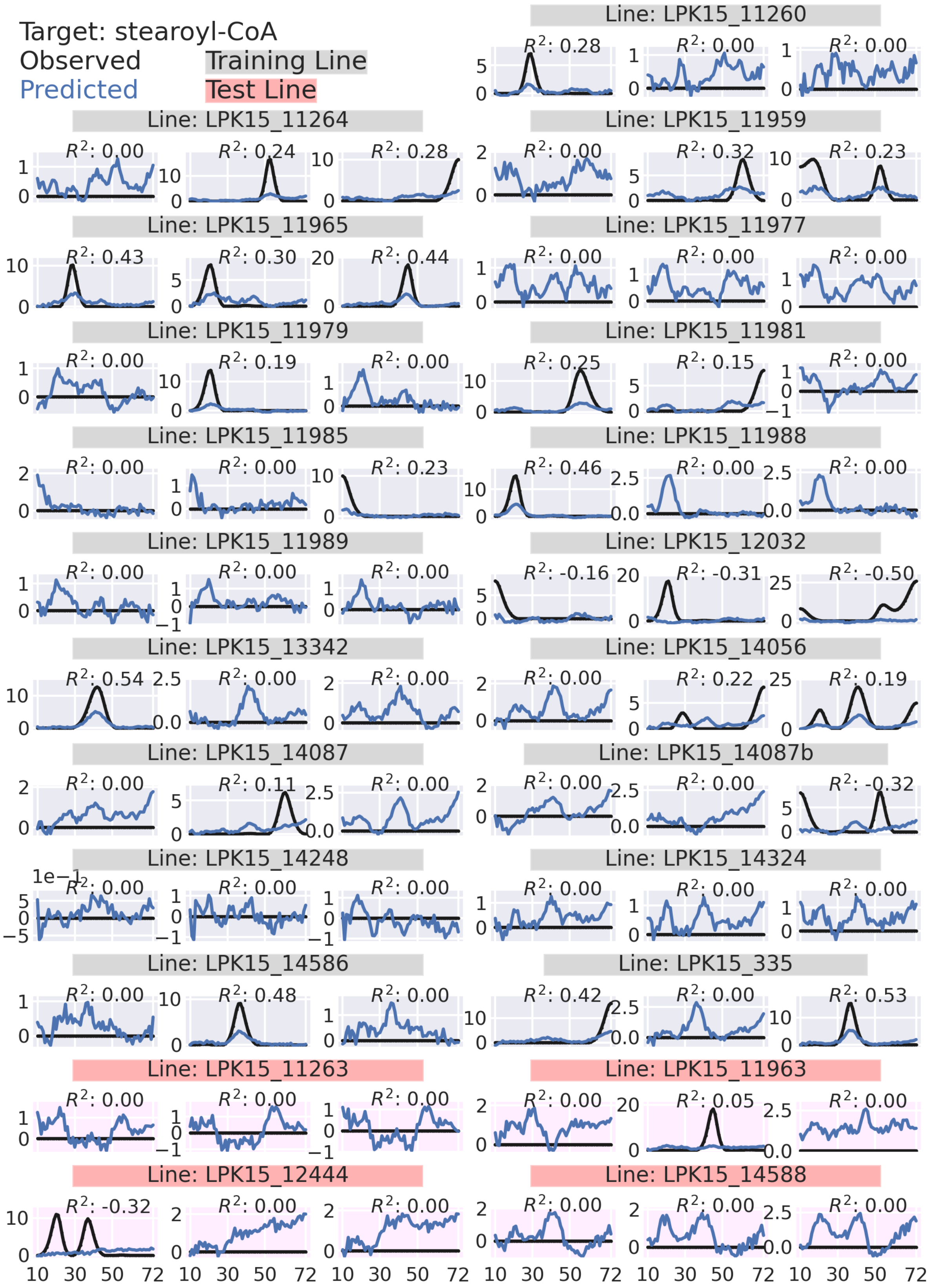
Target: stearoyl-CoA
Observed
Predicted
Training Line
Test Line
Line: LPK15_11260
Line: LPK15_11264
Line: LPK15_11959
Line: LPK15_11965
Line: LPK15_11977
Line: LPK15_11979
Line: LPK15_11981
Line: LPK15_11985
Line: LPK15_11988
Line: LPK15_11989
Line: LPK15_12032
Line: LPK15_13342
Line: LPK15_14056
Line: LPK15_14087
Line: LPK15_14087b
Line: LPK15_14248
Line: LPK15_14324
Line: LPK15_14586
Line: LPK15_335
Line: LPK15_11263
Line: LPK15_11963
Line: LPK15_12444
Line: LPK15_14588

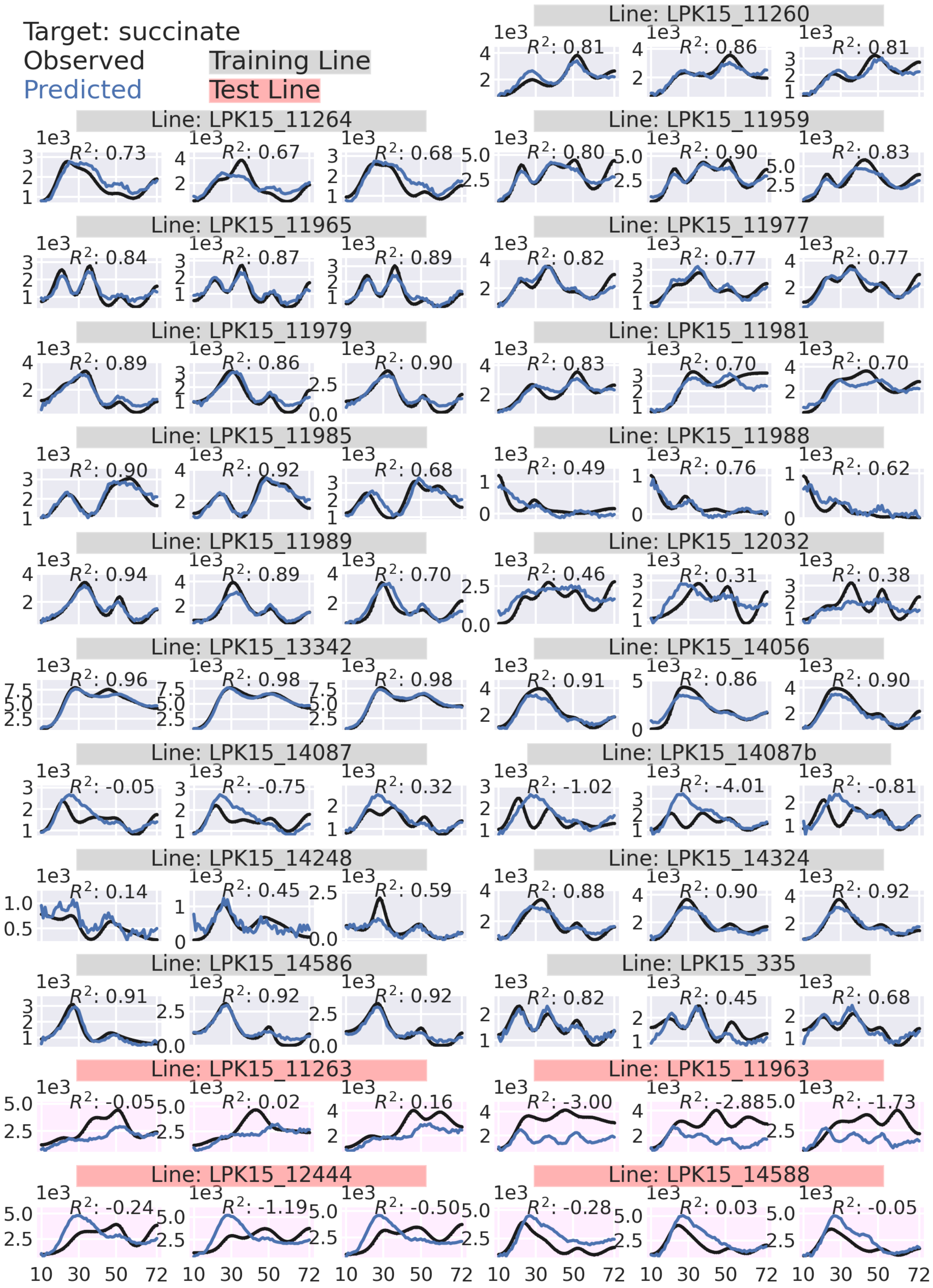

Target: succinate
Observed
Predicted
Training Line
Test Line
Line: LPK15_11260
Line: LPK15_11264
Line: LPK15_11959
Line: LPK15_11965
Line: LPK15_11977
Line: LPK15_11979
Line: LPK15_11981
Line: LPK15_11985
Line: LPK15_11988
Line: LPK15_11989
Line: LPK15_12032
Line: LPK15_13342
Line: LPK15_14056
Line: LPK15_14087
Line: LPK15_14087b
Line: LPK15_14248
Line: LPK15_14324
Line: LPK15_14586
Line: LPK15_335
Line: LPK15_11263
Line: LPK15_11963
Line: LPK15_12444
Line: LPK15_14588

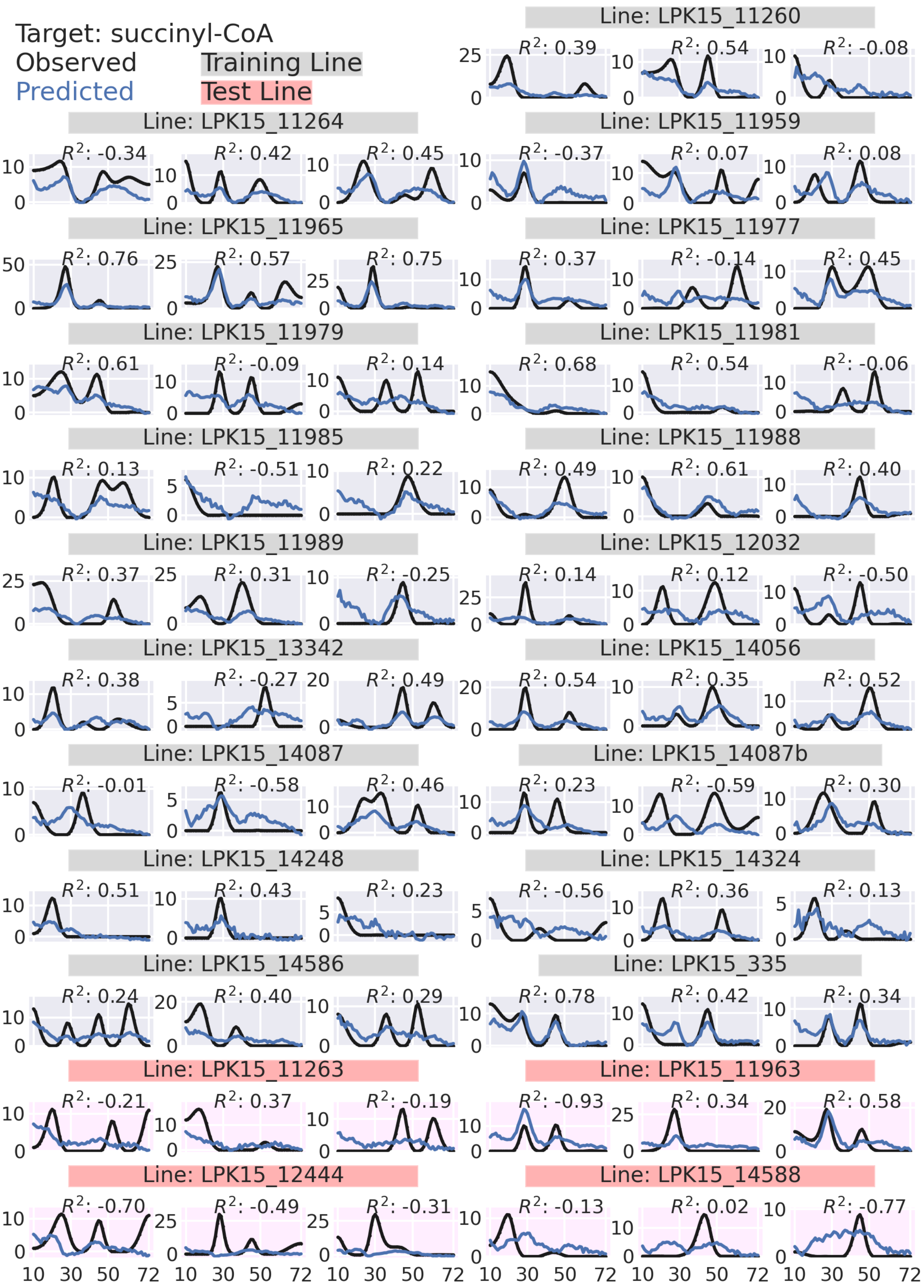

Target: succinyl-CoA
Observed
Predicted
Training Line
Test Line
Line: LPK15_11260
$R^2$: 0.39
$R^2$: 0.54
$R^2$: -0.08
Line: LPK15_11264
$R^2$: -0.34
$R^2$: 0.42
$R^2$: 0.45
Line: LPK15_11959
$R^2$: -0.37
$R^2$: 0.07
$R^2$: 0.08
Line: LPK15_11965
$R^2$: 0.76
$R^2$: 0.57
$R^2$: 0.75
Line: LPK15_11977
$R^2$: 0.37
$R^2$: -0.14
$R^2$: 0.45
Line: LPK15_11979
$R^2$: 0.61
$R^2$: -0.09
$R^2$: 0.14
Line: LPK15_11981
$R^2$: 0.68
$R^2$: 0.54
$R^2$: -0.06
Line: LPK15_11985
$R^2$: 0.13
$R^2$: -0.51
$R^2$: 0.22
Line: LPK15_11988
$R^2$: 0.49
$R^2$: 0.61
$R^2$: 0.40
Line: LPK15_11989
$R^2$: 0.37
$R^2$: 0.31
$R^2$: -0.25
Line: LPK15_12032
$R^2$: 0.14
$R^2$: 0.12
$R^2$: -0.50
Line: LPK15_13342
$R^2$: 0.38
$R^2$: -0.27
$R^2$: 0.49
Line: LPK15_14056
$R^2$: 0.54
$R^2$: 0.35
$R^2$: 0.52
Line: LPK15_14087
$R^2$: -0.01
$R^2$: -0.58
$R^2$: 0.46
Line: LPK15_14087b
$R^2$: 0.23
$R^2$: -0.59
$R^2$: 0.30
Line: LPK15_14248
$R^2$: 0.51
$R^2$: 0.43
$R^2$: 0.23
Line: LPK15_14324
$R^2$: -0.56
$R^2$: 0.36
$R^2$: 0.13
Line: LPK15_14586
$R^2$: 0.24
$R^2$: 0.40
$R^2$: 0.29
Line: LPK15_335
$R^2$: 0.78
$R^2$: 0.42
$R^2$: 0.34
Line: LPK15_11263
$R^2$: -0.21
$R^2$: 0.37
$R^2$: -0.19
Line: LPK15_11963
$R^2$: -0.93
$R^2$: 0.34
$R^2$: 0.58
Line: LPK15_12444
$R^2$: -0.70
$R^2$: -0.49
$R^2$: -0.31
Line: LPK15_14588
$R^2$: -0.13
$R^2$: 0.02
$R^2$: -0.77
10 30 50 72

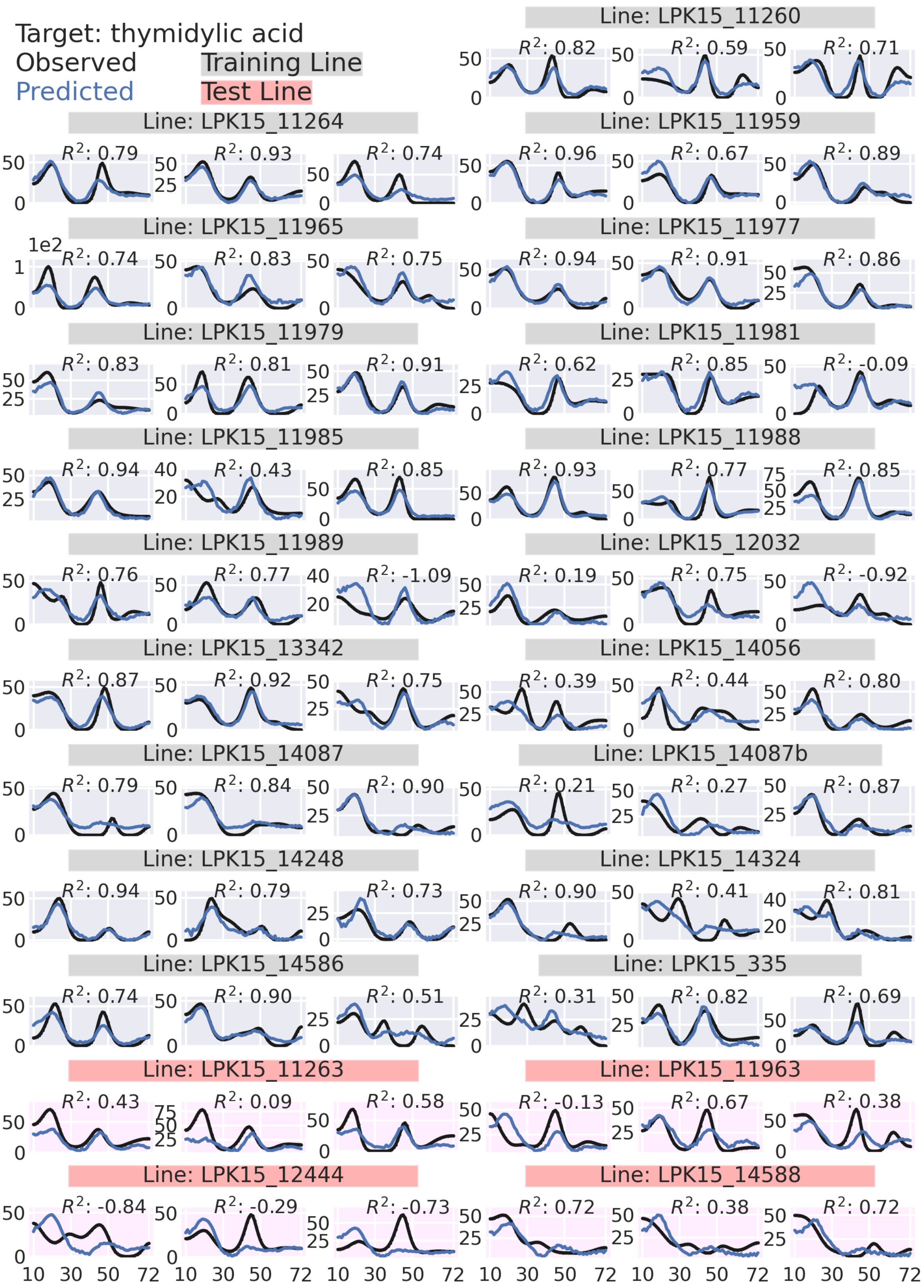
Target: thymidylic acid
Observed
Predicted
Training Line
Test Line
Line: LPK15_11260
$R^2$: 0.82
$R^2$: 0.59
$R^2$: 0.71
Line: LPK15_11264
$R^2$: 0.79
$R^2$: 0.93
$R^2$: 0.74
Line: LPK15_11959
$R^2$: 0.96
$R^2$: 0.67
$R^2$: 0.89
Line: LPK15_11965
1e2
$R^2$: 0.74
$R^2$: 0.83
$R^2$: 0.75
Line: LPK15_11977
$R^2$: 0.94
$R^2$: 0.91
$R^2$: 0.86
Line: LPK15_11979
$R^2$: 0.83
$R^2$: 0.81
$R^2$: 0.91
Line: LPK15_11981
$R^2$: 0.62
$R^2$: 0.85
$R^2$: -0.09
Line: LPK15_11985
$R^2$: 0.94
$R^2$: 0.43
$R^2$: 0.85
Line: LPK15_11988
$R^2$: 0.93
$R^2$: 0.77
$R^2$: 0.85
Line: LPK15_11989
$R^2$: 0.76
$R^2$: 0.77
$R^2$: -1.09
Line: LPK15_12032
$R^2$: 0.19
$R^2$: 0.75
$R^2$: -0.92
Line: LPK15_13342
$R^2$: 0.87
$R^2$: 0.92
$R^2$: 0.75
Line: LPK15_14056
$R^2$: 0.39
$R^2$: 0.44
$R^2$: 0.80
Line: LPK15_14087
$R^2$: 0.79
$R^2$: 0.84
$R^2$: 0.90
Line: LPK15_14087b
$R^2$: 0.21
$R^2$: 0.27
$R^2$: 0.87
Line: LPK15_14248
$R^2$: 0.94
$R^2$: 0.79
$R^2$: 0.73
Line: LPK15_14324
$R^2$: 0.90
$R^2$: 0.41
$R^2$: 0.81
Line: LPK15_14586
$R^2$: 0.74
$R^2$: 0.90
$R^2$: 0.51
Line: LPK15_335
$R^2$: 0.31
$R^2$: 0.82
$R^2$: 0.69
Line: LPK15_11263
$R^2$: 0.43
$R^2$: 0.09
$R^2$: 0.58
Line: LPK15_11963
$R^2$: -0.13
$R^2$: 0.67
$R^2$: 0.38
Line: LPK15_12444
$R^2$: -0.84
$R^2$: -0.29
$R^2$: -0.73
Line: LPK15_14588
$R^2$: 0.72
$R^2$: 0.38
$R^2$: 0.72
10 30 50 72

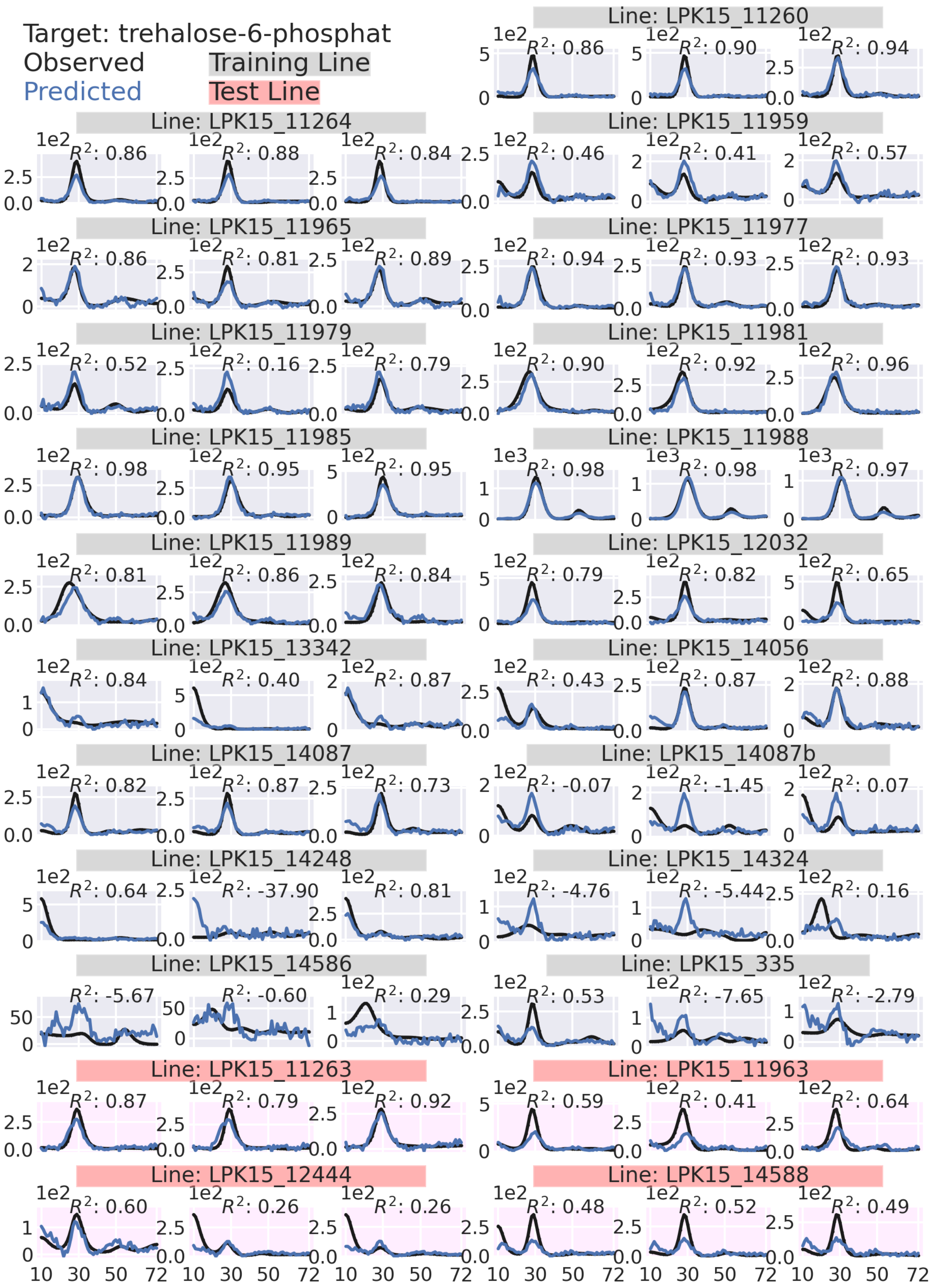

Target: trehalose-6-phosphat
Observed
Predicted
Training Line
Test Line
Line: LPK15_11260
R²: 0.86
R²: 0.90
R²: 0.94
Line: LPK15_11264
R²: 0.86
R²: 0.88
R²: 0.84
Line: LPK15_11959
R²: 0.46
R²: 0.41
R²: 0.57
Line: LPK15_11965
R²: 0.86
R²: 0.81
R²: 0.89
Line: LPK15_11977
R²: 0.94
R²: 0.93
R²: 0.93
Line: LPK15_11979
R²: 0.52
R²: 0.16
R²: 0.79
Line: LPK15_11981
R²: 0.90
R²: 0.92
R²: 0.96
Line: LPK15_11985
R²: 0.98
R²: 0.95
R²: 0.95
Line: LPK15_11988
R²: 0.98
R²: 0.98
R²: 0.97
Line: LPK15_11989
R²: 0.81
R²: 0.86
R²: 0.84
Line: LPK15_12032
R²: 0.79
R²: 0.82
R²: 0.65
Line: LPK15_13342
R²: 0.84
R²: 0.40
R²: 0.87
Line: LPK15_14056
R²: 0.43
R²: 0.87
R²: 0.88
Line: LPK15_14087
R²: 0.82
R²: 0.87
R²: 0.73
Line: LPK15_14087b
R²: -0.07
R²: -1.45
R²: 0.07
Line: LPK15_14248
R²: 0.64
R²: -37.90
R²: 0.81
Line: LPK15_14324
R²: -4.76
R²: -5.44
R²: 0.16
Line: LPK15_14586
R²: -5.67
R²: -0.60
R²: 0.29
Line: LPK15_335
R²: 0.53
R²: -7.65
R²: -2.79
Line: LPK15_11263
R²: 0.87
R²: 0.79
R²: 0.92
Line: LPK15_11963
R²: 0.59
R²: 0.41
R²: 0.64
Line: LPK15_12444
R²: 0.60
R²: 0.26
R²: 0.26
Line: LPK15_14588
R²: 0.48
R²: 0.52
R²: 0.49
10 30 50 72

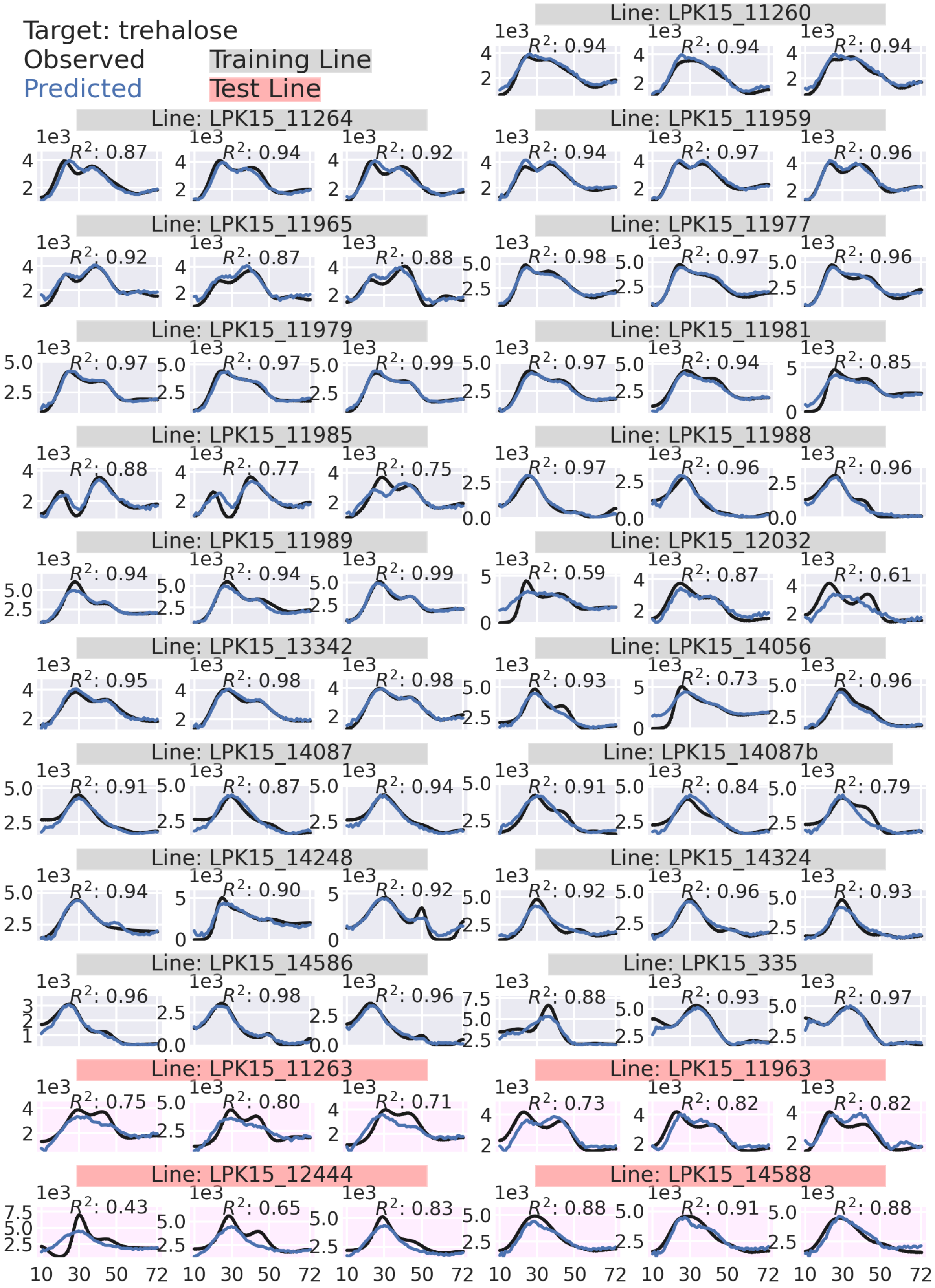

Target: trehalose
Observed
Predicted
Training Line
Test Line
Line: LPK15_11260
R²: 0.94
R²: 0.94
R²: 0.94
Line: LPK15_11264
R²: 0.87
R²: 0.94
R²: 0.92
Line: LPK15_11959
R²: 0.94
R²: 0.97
R²: 0.96
Line: LPK15_11965
R²: 0.92
R²: 0.87
R²: 0.88
Line: LPK15_11977
R²: 0.98
R²: 0.97
R²: 0.96
Line: LPK15_11979
R²: 0.97
R²: 0.97
R²: 0.99
Line: LPK15_11981
R²: 0.97
R²: 0.94
R²: 0.85
Line: LPK15_11985
R²: 0.88
R²: 0.77
R²: 0.75
Line: LPK15_11988
R²: 0.97
R²: 0.96
R²: 0.96
Line: LPK15_11989
R²: 0.94
R²: 0.94
R²: 0.99
Line: LPK15_12032
R²: 0.59
R²: 0.87
R²: 0.61
Line: LPK15_13342
R²: 0.95
R²: 0.98
R²: 0.98
Line: LPK15_14056
R²: 0.93
R²: 0.73
R²: 0.96
Line: LPK15_14087
R²: 0.91
R²: 0.87
R²: 0.94
Line: LPK15_14087b
R²: 0.91
R²: 0.84
R²: 0.79
Line: LPK15_14248
R²: 0.94
R²: 0.90
R²: 0.92
Line: LPK15_14324
R²: 0.92
R²: 0.96
R²: 0.93
Line: LPK15_14586
R²: 0.96
R²: 0.98
R²: 0.96
Line: LPK15_335
R²: 0.88
R²: 0.93
R²: 0.97
Line: LPK15_11263
R²: 0.75
R²: 0.80
R²: 0.71
Line: LPK15_11963
R²: 0.73
R²: 0.82
R²: 0.82
Line: LPK15_12444
R²: 0.43
R²: 0.65
R²: 0.83
Line: LPK15_14588
R²: 0.88
R²: 0.91
R²: 0.88
1e3
10 30 50 72

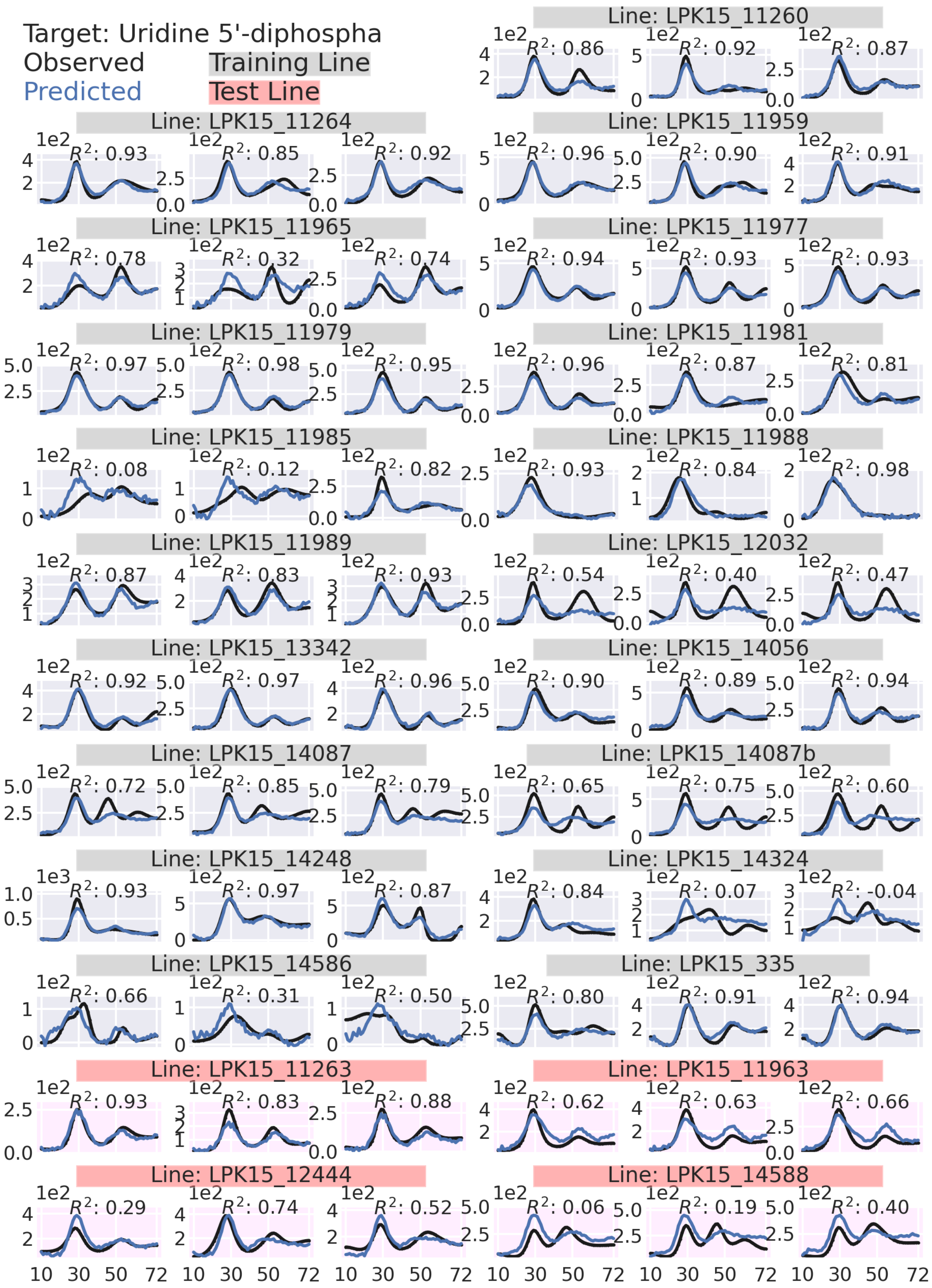
Target: Uridine 5'-diphospha
Observed
Predicted
Training Line
Test Line
Line: LPK15_11260
$R^2$: 0.86
$R^2$: 0.92
$R^2$: 0.87
Line: LPK15_11264
$R^2$: 0.93
$R^2$: 0.85
$R^2$: 0.92
Line: LPK15_11959
$R^2$: 0.96
$R^2$: 0.90
$R^2$: 0.91
Line: LPK15_11965
$R^2$: 0.78
$R^2$: 0.32
$R^2$: 0.74
Line: LPK15_11977
$R^2$: 0.94
$R^2$: 0.93
$R^2$: 0.93
Line: LPK15_11979
$R^2$: 0.97
$R^2$: 0.98
$R^2$: 0.95
Line: LPK15_11981
$R^2$: 0.96
$R^2$: 0.87
$R^2$: 0.81
Line: LPK15_11985
$R^2$: 0.08
$R^2$: 0.12
$R^2$: 0.82
Line: LPK15_11988
$R^2$: 0.93
$R^2$: 0.84
$R^2$: 0.98
Line: LPK15_11989
$R^2$: 0.87
$R^2$: 0.83
$R^2$: 0.93
Line: LPK15_12032
$R^2$: 0.54
$R^2$: 0.40
$R^2$: 0.47
Line: LPK15_13342
$R^2$: 0.92
$R^2$: 0.97
$R^2$: 0.96
Line: LPK15_14056
$R^2$: 0.90
$R^2$: 0.89
$R^2$: 0.94
Line: LPK15_14087
$R^2$: 0.72
$R^2$: 0.85
$R^2$: 0.79
Line: LPK15_14087b
$R^2$: 0.65
$R^2$: 0.75
$R^2$: 0.60
Line: LPK15_14248
$R^2$: 0.93
$R^2$: 0.97
$R^2$: 0.87
Line: LPK15_14324
$R^2$: 0.84
$R^2$: 0.07
$R^2$: -0.04
Line: LPK15_14586
$R^2$: 0.66
$R^2$: 0.31
$R^2$: 0.50
Line: LPK15_335
$R^2$: 0.80
$R^2$: 0.91
$R^2$: 0.94
Line: LPK15_11263
$R^2$: 0.93
$R^2$: 0.83
$R^2$: 0.88
Line: LPK15_11963
$R^2$: 0.62
$R^2$: 0.63
$R^2$: 0.66
Line: LPK15_12444
$R^2$: 0.29
$R^2$: 0.74
$R^2$: 0.52
Line: LPK15_14588
$R^2$: 0.06
$R^2$: 0.19
$R^2$: 0.40
1e2
1e3
10 30 50 72

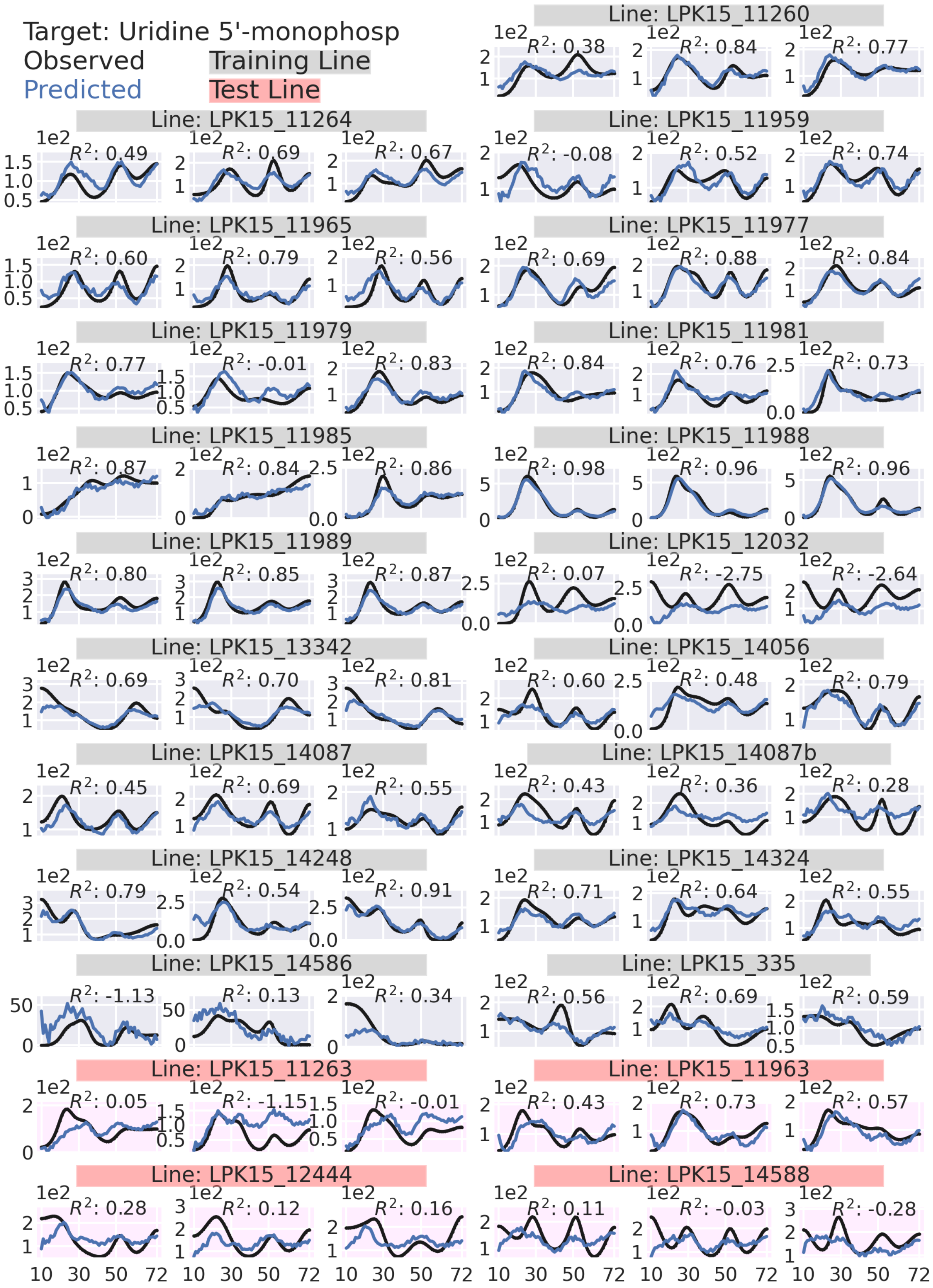

Target: Uridine 5'-monophosp
Observed
Predicted
Training Line
Test Line
Line: LPK15_11260
Line: LPK15_11264
Line: LPK15_11959
Line: LPK15_11965
Line: LPK15_11977
Line: LPK15_11979
Line: LPK15_11981
Line: LPK15_11985
Line: LPK15_11988
Line: LPK15_11989
Line: LPK15_12032
Line: LPK15_13342
Line: LPK15_14056
Line: LPK15_14087
Line: LPK15_14087b
Line: LPK15_14248
Line: LPK15_14324
Line: LPK15_14586
Line: LPK15_335
Line: LPK15_11263
Line: LPK15_11963
Line: LPK15_12444
Line: LPK15_14588

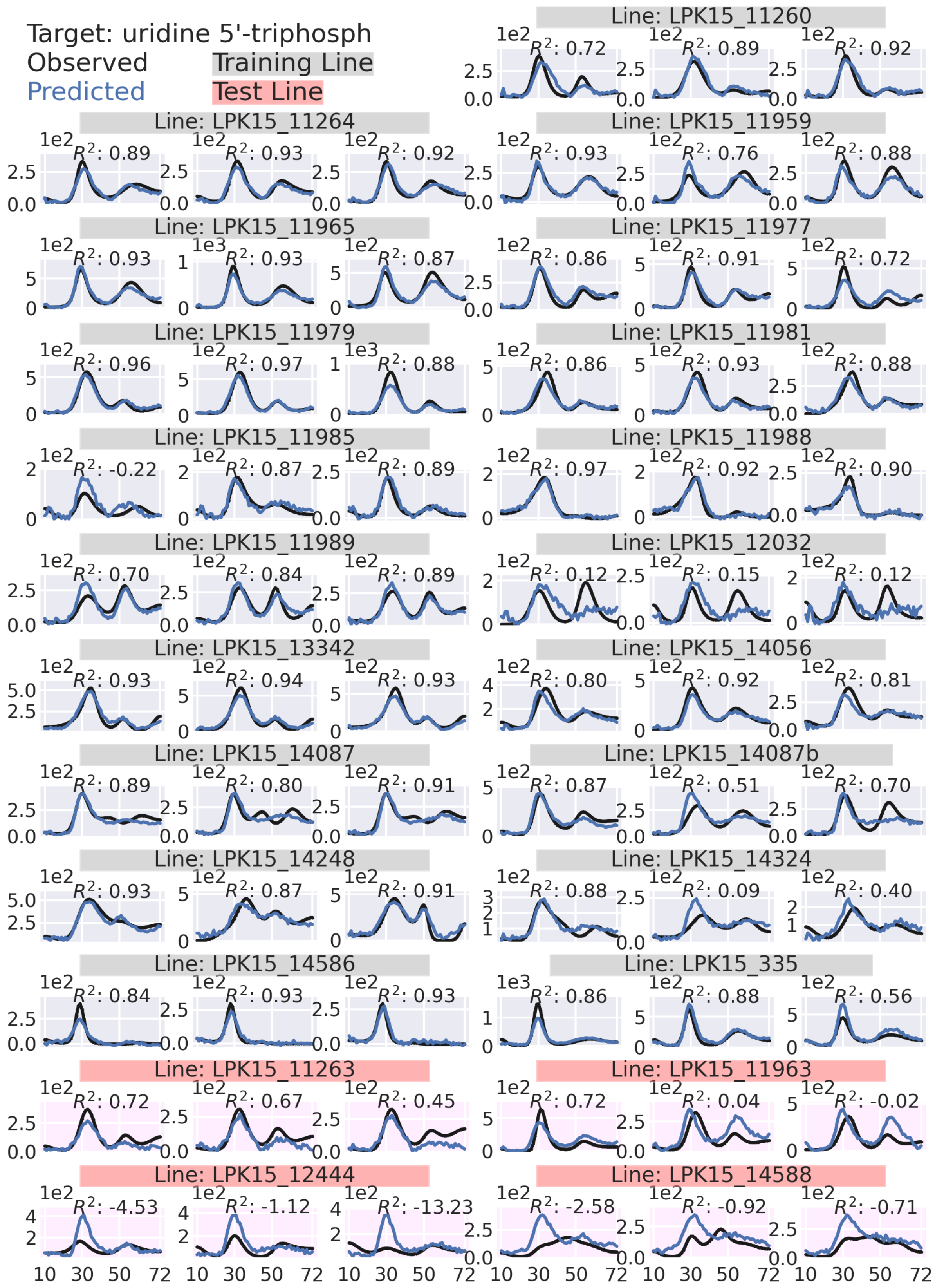

Target: uridine 5'-triphosph
Observed
Predicted
Training Line
Test Line
Line: LPK15_11260
Line: LPK15_11264
Line: LPK15_11959
Line: LPK15_11965
Line: LPK15_11977
Line: LPK15_11979
Line: LPK15_11981
Line: LPK15_11985
Line: LPK15_11988
Line: LPK15_11989
Line: LPK15_12032
Line: LPK15_13342
Line: LPK15_14056
Line: LPK15_14087
Line: LPK15_14087b
Line: LPK15_14248
Line: LPK15_14324
Line: LPK15_14586
Line: LPK15_335
Line: LPK15_11263
Line: LPK15_11963
Line: LPK15_12444
Line: LPK15_14588
